# Identifying Novel Candidate Defense Genes Against Rice Blast By Disease-Resistance Transcriptome Analysis

**DOI:** 10.1101/2022.10.05.510921

**Authors:** Ramil Mauleon, Kouji Satoh, Violeta Bartolome, Marietta Baraoidan, Emily Deomano, Rita P. Laude, Shoshi Kikuchi, Hei Leung

## Abstract

A blast-resistance rice mutant, GR978, generated by gamma-irradiation of *indica* cultivar IR64 was used to characterize the disease resistance transcriptome of rice to gain a better understanding of genes or chromosomal regions contributing to broad-spectrum disease resistance. GR978 was selected from the IR64 mutant collection at IRRI. To facilitate phenotypic characterization of the collection, a set of controlled vocabularies (CV) documenting mutant phenotypes in ∼3,700 entries was developed. In collaboration with the Tos17 rice mutant group at National Institute of Agrobiological Sciences, Japan, a merged CV set with 91 descriptions that map onto public ontology databases (PO, TO, OBO) is implemented in the IR64 mutant database.

To better characterize the disease resistance transcriptome of rice, gene expression data from a blast resistant cultivar, SHZ-2, was incorporated in the analysis. Disease resistance transcriptome parameters, including differentially expressed genes (DEGs), regions of correlated gene expression (RCEs), and associations between DEGs and RCEs were determined statistically within and between genotypes using MAANOVA, correlation, and fixed ratio analysis. Twelve DEGs were found within the inferred physical location of the recessive gene locus on a ∼3.8MB region of chromosome 12 defined by genetic analysis of GR978. Highly expressed DEGs (≥ 2fold difference) in GR978 or SHZ-2 and in common between the two, are mostly defense-response related, suggesting that most of the DEGs participate in causing the resistance phenotype.

Comparing RCEs between SHZ-2 and GR978 showed that most RCEs between genotypes did not overlap. However, an 8-gene RCE in chromosome 11 was in common between SHZ2 and GR978. Gene annotations and GO enrichment analysis showed a high association with resistance response. This region has no DEGs nor is it associated with known blast resistance QTLs. Association analyses between RCEs and DEGs show that there was no enrichment of DEGs in the RCEs within a genotype and across genotypes as well.

Association analysis of blast-resistance QTL (Bl-QTLs) regions (assembled from published literature; data courtesy of R. Wisser, pers comm., Cornell University) with DEGs and RCEs showed that while Bl- QTLs are not significantly associated with DEGs, they are associated with genotype-specific RCEs; GR978- RCEs are enriched within Bl-QTLs. The analysis suggested that examining patterns of correlated gene expression patterns in a chromosomal context (rather than the expression levels of individual genes) can yield additional insights into the causal relationship between gene expression and phenotype. Based on these results, we put forward a hypothesis that QTLs with small or moderate effects are represented by genomic regions in which the genes show correlated expression. It implies that gene expression within such a region is regulated by a common mechanism, and that coordinated expression of the region contributes to phenotypic effects. This hypothesis is testable by co segregation analysis of the expression patterns in well-characterized backcross and recombinant inbred lines.

## INTRODUCTION

Rice (*Oryza sativa* L) has emerged as the new model plant for functional genomic studies. It is an ideal plant for plant biology studies, with a relatively short generation time, prolific in production of seeds, and is easy to cultivate under greenhouse conditions. Genetic work on rice is easy since it has a relatively small genome and is a diploid with twelve chromosomes. Its self-pollinating nature can be considered an advantage and a drawback for breeders, and there are well-established breeding techniques to get around this. Along with all these features is the fact that rice is the primary staple for more than half of the world’s population (Maclean et al., 2002), which has made it the focus of one of the most (if not the most) intense and continuously publicly funded crop improvement programs worldwide. Due to public interest in many developed and developing nations, the major focus of improvement has been to genetically equip rice to become raise yield efficiency, more tolerant or resistant to biotic and abiotic stresses, and improve nutrition quality, in no particular order.

In 2002, the “race” to publish the first draft sequence of the *Oryza sativa* genome can be declared a draw, with the simultaneous publication of the draft *indica* subspecies (cultivar 93-11) and *japonica* ssp. (cv. Nipponbare) sequences (Goff et al., 2002, Yu et al., 2002).

The draft sequence of the *indica* rice genome was finished using whole-genome shotgun sequencing. The genome was 466 megabases in size, with an estimated 46,022 to 55,615 genes from a 92.0% functional coverage in the assembled sequences. Approximately 42.2% of the genome was in exact 20-nucleotide oligomer repeats, and most of the transposons were in the intergenic regions. In terms of homology to *Arabidopsis thaliana* genes, 80.6% of *A. thaliana* predicted genes had a homolog in rice, while only 49.4% of predicted rice genes had a homolog in *A. thaliana*. The authors attributed the large proportion of rice genes with no recognizable homologs (NH genes) as the result of a gradient in the GC content of rice coding sequences (Yu et al. 2002).

For the *japonica* genome, the whole-genome shotgun sequencing was also employed, with the assembled sequence covering 93% of the 420-megabase genome. Gene predictions suggest that there were 32,000 to 50,000 genes. Synteny and gene homology (98%) between rice and the other cereal genomes are extensive, whereas synteny with *Arabidopsis* is limited, though assignment of candidate rice orthologs to *Arabidopsis* genes is possible in many cases (Goff et al. 2002).

In terms of expressed rice genes, Kikuchi et al. (2003) collected and completely sequenced 28,469 full-length complementary DNA (FL-cDNA) clones from the *japonica* cultivar, Nipponbare (the same *japonica* variety used for genome sequencing). Alignment of the FL-cDNA sequences with publicly available sequence data showed homology of 21,596 clones, and tentative protein functions based on these alignments were assigned to these clones. Mapping the FL-cDNA clones back to the *japonica* genome shows there are 19,000 to 20,500 transcription units in the rice genome. Alignment of the translated protein with InterPro (2001) database showed the existence of proteins present only in rice but not in the *Arabidopsis* proteome, while 64% of the FL-cDNAs are homologous to *Arabidopsis* proteome.

Jabbari et al. (2004) questioned the large number of genes predicted by both groups and concentrated on further characterizing the roughly half of the protein-coding genes in rice which, apparently, had no significant homologs in the previously sequenced genome of *Arabidopsis*, or, in most cases, in the sequenced DNA of any other taxa (the NH genes) They noted that these genes could be new type(s) of anomalous but functional proteins, possibly specific to Gramineae or monocots, with highly unusual compositional and structural features. However, their suspicion is that a third or more of the published *ab initio* coding predictions for rice are incorrect, and that more stringent gene prediction criteria should be adapted.

With the publication of the complete genome sequences of both *indica* (93-11, Yu et al 2002) and *japonica* rice cultivars (Nipponbarre, Goff et al. 2002), and the subsequent completely annotated Nipponbarre *japonica* genome from The International Rice Genome Sequencing Project (IRGSP release 4.0) and The Institute for Genomic Research (TIGR rice pseudomolecule rel 4.0) comes the potential of dissecting and understanding gene functions and metabolic pathways in order to further improve the crop. Knowledge of the gene constitution of the rice genome, freely provided by the two groups mentioned, enables the researcher to examine all at once, the expression of thousands of genes in response to specific conditions within specific cultivars of interest. DNA microarray technology has enabled high throughput examination of gene expression profiles, and there are many sources of microarray platforms (commercial or non-profit) available for this type of study such as the Beijing Genomics Institute rice oligochip with 44,000 rice genes, the US NSF Rice Oligonucleotide Array Project oligochip with 45,000 genes, and the Agilent Technologies rice catalog oligoarray with ∼22,000 genes, all of which use short nucleotide signatures of the informative genes from the sequenced rice genomes. Costs of facilities, equipment, and consumables to conduct microarray-based gene expression laboratory work still remain high, though. The prevailing spirit of collaboration among rice researchers however, is a solution to the cost constraint. The computing power needed to manage and analyze the tremendous amount of data generated by such high throughput procedures is available cheaply in ordinary desktop personal computers, though.

Almost all of the researches for genetic improvement of rice can utilize microarray technology, taking advantage of the high throughput nature of the method. However, the power of microarrays are highly dependent on the target material being compared at once; for basic rice biology studies of very particular traits, comparing highly diverse rice cultivars directly can generate a lot of expression “noise”. The use of rice mutants to which gene disruptions are well known or characterized makes the dissection and exploration of individual genes and their interactions more tenable using gene expression data. Mutant resources have been used to great success in characterizing mutant genes and the associated altered gene expression caused by the disrupted gene (Pan, 2004, NPR1 gene in *Arabidopsis*, Zeng et al. 2004, in *spl11* rice mutant).

One of the enduring problems in rice research is breeding for durable resistance against fungal pathogens such as rice blast caused by the fungus *Magnaporthe grisea*. The diversity of rice blast populations in the field results in the rapid “breakdown” of newly introduced rice varieties that are bred specifically for blast resistance (Zeigler et al. 1994). Rice blast disease is the most widespread and damaging disease of cultivated rice, and it can be found in both the tropics as well as the temperate rice growing regions. The disease is caused by the fungus *Pyricularia grisea* Sacc., the anamorph of *Magnaporthe grisea* (Hebert) Barr, (Zeigler et al, 1994). Rice plants are most seriously damaged when the leaves (during early vegetative) and panicles (during reproductive stage) are attacked, although all above-ground parts are vulnerable to infection. Recent reports describe infection of the roots as well (Sesma and Osbourn 2004). Rice breeders have for so long focused on breeding for resistance in order to control rice blast, typically by identification of pathogenic races of the blast fungus and incorporating corresponding resistance genes from various rice germplasm, resulting in a new resistant variety (Ou 1985). Numerous failures to this approach are well documented, with varieties becoming susceptible to blast after a few short seasons of planting. The widely accepted explanation for this “breakdown” of resistance is the diversity of blast populations in the field, which implies pathogenic variability. The study of blast population structures was started with the discovery of neutral, repetitive DNA sequences (designated as MGR for *Magnaporthe grisea* repeat) by Hamer et al. (1989) in the rice blast genome in 1989, and this sequence provided a method of studying populations independently of the pathogenicity of the fungal isolates. MGR “fingerprints” generated by RFLP and PCR-based DNA-analysis of the blast genome provided an estimate of their relatedness of different isolates. Levy et al. (1993) showed that there was a direct relationship between MGR fingerprint type (now known as lineages) and pathogenic races. Zeigler et al. (1994) reported some success in breeding for blast resistance by selecting resistance genes that work against blast lineages, not just with individual pathogenic races. The lack of gene information in the host and pathogen side, however, has made this strategy difficult, especially when in rice growing regions with a very complex pathogen population. With the advent of genome sequencing, gene information has become available to the researcher.

Genomic sequence information has been rapidly increasing since the 1990s, and with the publication of the rice blast genome, putative mechanisms for virulence and pathogenic variability can be identified (Dean et al. 2005). Preliminary analysis of the draft blast genome reveals (1) a very diverse set of proteins involved in extra cellular perception and signal transduction via a family of novel G-protein-coupled receptors, (2) an extensive array of secreted proteins and secondary metabolites, specifically adapted regulatory pathways controlling infection-related development, and (3) repetitive DNA and repeat-induced point mutations, which implies generation of genetic variation even without sexual reproduction. Novel targets for disease control using candidate host genes can be inferred from this information.

The DNA microarray is the latest in a line of techniques to exploit a potent feature of the DNA duplex—the sequence complementarity of the two strands (Southern et al 1999). Sequence hybridization has been utilized since the 1960s, when Gillespie and Spiegelman (1965), observed that single stranded DNA binds strongly to nitrocellulose membranes such that it prevents the strands from reannealing with each other, but permits hybridization to complementary RNA. Almost immediately, the method was expanded to investigate copy number of rRNA and tRNA species into what became known as a dot blot (Kafatos et al., 1979). DNA microarrays evolved from dot blots through the use of an impermeable, rigid substrate, such as glass, which has a number of practical advantages over porous membranes and gel pads. Schena et al. reported the first cDNA array fabrication (via robotic printing) and use in 1995, reporting the differential expression measurements of 45 *Arabidopsis* genes by means of simultaneous, two-color fluorescence hybridization.

Since then, the use of DNA microarrays for gene expression monitoring and quantification has been growing by leaps and bound, with almost every genomic study reporting the use of this technology. In plants, the first focus of studies was with the model dicot *Arabidopsis*, owing to the availability of the genome sequence. Riechmann et al. (2000) studied *Arabidopsis* transcription factors and compared them with other eukaryote via microarray. They found that *Arabidopsis* dedicates over 5% of its genome to code for more than 1,500 transcription factors, about 45% of which are from families specific to plants. *Arabidopsis* transcription factors that belong to families common to all eukaryotes do not share significant similarity with those of the other kingdoms beyond the conserved DNA binding domains. Cheong et al. (2002) used Affymetrix™ *Arabidopsis* gene chips to profile gene expression in wounded plants, under different stress and hormone exposures, revealing novel interactions between wounding, pathogen, abiotic stress, and hormonal responses. It appears that wounding activate novel genes that signal other signal transduction pathways. Expression profiling of transcription factors in *Arabidopsis* by Chen et al. (2002) reveal that transcription factor genes apparently involved in stress responses exhibit nonspecific and specific alterations in expression profiles, and that they affect salicylic acid, jasmonic acid, and ethylene signaling pathways. Furthermore, stress response and senescence in *Arabidopsis* may share overlapping signaling pathways. Brodersen et al. (2002) used a cDNA microarray platform developed by Ruan et al. (1998) to monitor 9,861 cDNAs expressed throughout *Arabidopsis* development, in order to compare expression profiles of *acd11* mutants and with wildtype seedling. Pan et al. (2004) produced their own *Arabidopsis* cDNA array and applied various data mining techniques combined with sequence motif information in the promoter region of genes to discover functional genes that are involved in the defense mechanism of systemic acquired resistance. Marathe et al (2004) used the commercially available Affymetrix ATH1 gene chip representing the entire genome of *Arabidopsis* to profile the RCY1-mediated response of *Arabidopsis* ecotype C24 against the viral pathogen CMV-Y.

For rice, Oh et al. (2005) developed transgenic rice plants that constitutively expressed CBF3/DREB1A (CBF3) and ABF3, *Arabidopsis* genes that function in abscisic acid-independent and abscisic acid-dependent stress-response pathways, respectively, with the objective of finding native rice genes that were induced by the transgenes, as the plants were subjected to drought or freezing conditions. Their platform for gene expression study was the GreenGene 60K rice oligo chip, which shares the same sequences as the BGI 60k oligochip. Twelve rice genes were affected by CBF3 while seven were affected by ABF3. The transgenic plants became more tolerant to the stress conditions without exhibiting stunting of growth. Salzmann et al. (2005) used an in-house cDNA platform from sorghum ESTs to profile gene transcription in sorghum exposed to signaling compounds that are involved in plant defense systems, namely salicylic acid, methyl jasmonate (MeJA), and the ethylene precursor aminocyclopropane carboxylic acid. They found not only cooperative regulation of the other pathogenesis and defense-response genes but also mutual and one-way antagonisms as well, indicating that a subset of genes coregulated by SA and JA may comprise a uniquely evolved sector of plant signaling responsive cascades. Cooper et al. (2003) used a custom Affymetrix GeneChip Rice Genome Array representing 21,000 predicted ORFs in ∼42,000 contigs consisting the Syngenta japonica rice genome assembly. They combined this microarray expression data under a variety of conditions with protein–protein interactions, localized rice genes to regions within QTLs, and evaluated mutant plants. They discovered sets of genes modulated by stressful and developmental stimuli and identified cognate proteins governed by associations with other proteins involved in similar stress and developmental responses. With this integrated set of genomics data, they ascribed functions to >200 rice genes and verified these functions with evaluation of plants with mutations in these genes. More interestingly, they were able to construct a network of rice genes associated with biotic and abiotic stress response and seed development.

Walia et al. (2005) used the commercial GeneChip® Rice Genome Array from Affymetrix, the design of which was created within the Affymetrix GeneChip® Consortia Program. This chip contains probe sets designed to query 51,279 transcripts representing indica (1,260 transcripts) and japonica (48,564 transcripts) rice cultivars. They compared the transcriptome of two contrasting indica genotypes (salt-tolerant FL478, and salt-sensitive IR29) under salt stress and reported that there is an induction in a number of genes involved in the flavonoid biosynthesis pathway in IR29 but not in FL478.

The use of Agilent Technologies rice oligoarray system for genome-wide expression profiling was reported by Yazaki et al. in 2004. In their study, the used a custom oligoarray composed of ∼20,000 genes to profile rice callus treated with either absiscic acid (ABA) or gibberellic acid (GABA). The oligoarray utilized 60-mer oligos designed from the FL-cDNA collection of Kikuchi et al. (2003). They identified 200 ABA-responsive and 301 GA-responsive genes, many of which are not annotated as responsive to either chemical in prior expression studies.

A major innovation that could be brought by microarray technology would be to determine and understand the co-regulation of genes expressed during blast resistance response. The use of multiple-pathogen gain of resistance rice mutants for microarray gene expression studies will enable dissection of gene expression profiles associated with broad-spectrum disease resistance. This can pave the way for the discovery of new candidate genes important for broad spectrum and durable resistance against rice blast and other disease affecting rice, as well as the understanding of the plant-pathogen interaction at the gene level. These are important research frontiers where microarray technology is the tool of choice, and the results can be used in breeding programs for improving rice disease resistance.

Several key developments that enable functional genomic research of defense response (DR) and broad-spectrum resistance (BSR) in rice are:

1. *Knowledge of candidate genes implicated in BSR.* A wealth of experimentally validated information regarding genes involved in DR against biotic and abiotic stresses has been compiled and is publicly available. These genes range from those of *Arabidopsis* and most interestingly, rice genes (Ramalingam et al. 2003, Glazebrook 2001, Glazebrook et al, 2003, Cooper et al. 2003). Focusing on the disease response genes annotated in the japonica sequence, a degree of conservation between *Arabidopsis* and rice was observed. Disease resistance (R) genes function for early and specific recognition of pathogen attack and initiating signal transduction, enabling the host plant to deploy their defense mechanisms (Dangl and Jones 2001). DR genes are categorized into two major and three minor structural classes. The largest class of known R gene products contains characteristic nucleotide binding sites, leucine-rich repeats (NB-LRRs), and an apoptosis-resistance-conserved (ARC) domain, and the rice genome has ∼600 genes homologous to this class. However, a striking difference in the rice orthologs is the absence of obvious genes that encode TIR (Toll–interleukin 1 receptor resistance) motifs at their NH2-termini; these are very common in the dicot orthologs, but homology in the rice NBS-LRR are very weak. These domains in rice may function like TIR domains in dicots but are highly diverged in terms of sequence, and are likely evolved after the divergence of monocots and dicots. Another major class of DR genes code for extracellular LRRs either with short cytoplasmic tails or COOH-terminal serine-threonine protein kinase domains. In rice, there are ∼450 extracellular LRR genes, and ∼50% of them encode COOH-terminal protein kinase domains. This structural class differs from the NB-LRR genes since it is known to include proteins with functions not related to disease resistance (Dangl and Jones 2001). For the minor classes of DR genes, these include genes encoding the cytoplasmic serine-threonine kinases Pto and PBS1, each have 14 rice homologs; Hs1pro-1, with one rice homolog; and RPW8, with one rice homolog. Most of the known *Arabidopsis* genes known to control disease resistance responses have putative orthologs in rice, suggesting extensive conservation of disease resistance signaling between monocots and dicots. One rice homolog corresponds for each of the *Arabidopsis* disease signal transduction genes NDR1, PAD4, and EDS1, as well as the barley gene RAR1. Three rice homologs corresponded to COI1, a gene required for responses to the signal molecule jasmonic acid (JA); six for NPR1, a gene required for responses to the signal molecule salicylic acid (SA); and six for LSD1, a gene required for control of programmed cell death (PCD). Many rice homologs for the *Arabidopsis* mitogen-activated protein kinase (MAPK) gene MPK4 and the MAPK kinase gene EDR1 were found, preventing the assignment of putative orthologs. No rice gene similar to Arabidopsis SNI1 was found (Goff et al. 2002).

Numerous studies have identified rice and *Arabidopsis* defense response genes against biotic stresses such as fungal and bacterial diseases. The general model of host plant resistance is that plants carry specific R genes that can recognize pathogens carrying corresponding avirulence (avr) genes, triggering a rapid defense response that generally includes programmed plant cell death in the area in contact with the pathogen. This phenomenon is called the hypersensitive response (HR - van der Biezen 1998). New findings by Leister and Katagiri (2000) suggest that R proteins ‘guard’ plant proteins (i.e. ‘guardees’) that are the actual targets of pathogen avr proteins. The HR response is then triggered when the avr– guardee interactions are detected]. They reported the detection of an *in vivo* complex containing an R protein, an avr protein, and an unidentified plant protein. To date, there are four models for defense response, namely: R-gene signal transduction (which includes the PBS, NBS-LRR-TIR, and the leucine zipper containing genes), SA-dependent signaling resistance, JA/ethylene-dependent signaling resistance, and induced systemic resistance (Glazebrook 1999, 2001). Additional genes affecting SA-dependent signaling as well as for JA-dependent signaling and for induced systemic resistance have been identified.

In rice, gene sequences from defense-related pathways from other crops were used to genetically map chromosomal regions associated with the resistance phenotype. Ramalingam et al. (2003) used candidate genes from rice, barley, and maize involved in both recognition (resistance gene analogs [RGAs] – such as aldose reductase, oxalate oxidase, JAMyb (a jasmonic acid-induced Myb transcription factor), and peroxidase) and general plant defense (putative defense response [DR) as molecular markers (either STS or RFLP) to test for association with resistance in rice to blast, bacterial blight (BB), sheath blight, and brown plant-hopper (BPH). Many RGAs and DR genes detected a single locus with variable copy number and mapped on different chromosomes. Clusters of RGAs were observed in regions where many known blast and BB resistance genes and quantitative trait loci (QTL) for blast, BB, sheath blight, and BPH were located. Major resistance genes and QTL for blast and BB resistance located on different chromosomes were associated with several candidate genes. Using a similar approach, Bin et al. (2004) used 5 heterologous defense response genes (encoding putative oxalate oxidase, dehydrin, PR-1, chitinase, and 14-3-3 protein) to map five QTLs and a major gene for rice blast resistance in the cultivar SHZ-2. These results give a hint of the conserved roles of the rice heterologs of these DR/RGA genes from other crops.

Brodersen et al. (2002) described a lethal recessive *accelerated-cell-death11 Arabidopsis* mutant (*acd11*) that causes constitutive expression of defense-related genes and PCD associated with HR triggered by avirulent pathogen. These responses are SA-dependent and requires mediation by PAD4 and EDS1. The expression of *acd11* mutation depends on the SA signal transducer NPR11.

Pan et al. (2004) studied the expression profiles of genes altered by the mutation of a key regulator gene of systemic acquired resistance (SAR) response in *A. thaliana*, *NPR1*. Comparisons of the wild-type with *npr1-3* mutant plus a combination of novel data mining techniques were used to reveal in two highly resolved gene groups, implicated in the SAR phenotype. In these groups, there were 24 candidate genes (12 were down regulated and 12 were up-regulated genes) in the npr1-3 mutant plants in response to treatment with SA. Promoter analysis of each group shows that the down-regulated genes were highly enriched with W-box and Wy-box motifs, while that of up-regulated genes are deprived of W-box. The ASF-1 motif appears to be correlated with the phenotype: throughout the entire genome of *A. thaliana*, the motif appears to be under represented, and it is further so in the up-regulated group of genes. However, a group of 8 most informative down regulated genes are enriched with the ASF-1 motif as compared with its genome wide distribution.

Marathe et al. (2004) analyzed the *RCY1* -mediated resistance response in *Arabidopsis* against the viral pathogen Cucumber Mosaic Virus-Yellow strain by comparing expression profiles of mock and virus inoculated *A. thaliana* ecotype C24. *RCY1* conditions hypersensitive response in *Arabidopsis*. They identified 444 putative factors belonging to 9 different functional classes associated with the *RCY1* effect, and 80 of these factors are defense responsive genes.

Zeng et al. (2004) isolated a rice gene, *Spl11*, facilitated by the identification of *spl11* mutant alleles in IR64; plants with these mutant alleles show enhanced resistance against rice fungal and bacterial pathogens. In vitro assays indicate that the SPL11 protein possesses E3 ubiquitin ligase activity, which suggest the role of the ubiquitination system in controlling PCD and defense response.

2. *Ideal biological resources for the study of BSR.* In order to uncover the functions of genes predicted by sequence analysis of the rice genome, rice plants with altered gene functions need to be available for phenotype – gene reverse genetics study. Currently, several institutions offer rice mutants with gene knockouts generated using various insertion and/or deletion methods. Hirochika et al. (2004) summarized the status of rice mutant resources and listed ten centers with available mutants. Different cultivars were used by each center for mutagenesis (Nipponbare, IR64, Tainung 67, Dongjin, Hwayoung, Dongjinbyeo, and Zhonghua 11). Depending on the method of mutagenesis, target site specificity of each insertion element and subsequent knockout efficiency varies. Available data indicate that insertions of Ac/Ds, T-DNA and Tos17 are biased toward low-copy, gene-rich regions (Enoki et al., 1999; Miyao et al., 2003; An et al., 2003; Kolesnik et al., 2004; Sallaud et al. 2004). Zhang et al. (2006) reported the Rice Mutant Database (RMD) which contains information regarding mutant phenotypes, reporter-gene expression patterns, flanking sequences of T-DNA insertional sites, seed availability, among others, for approximately 129,000 rice T-DNA insertion lines (enhancer trap lines, generated by an enhancer trap system using japonica cultivars Zhonghua 11 and 15)

Chemical and irradiation mutagenesis, while causing a larger number of mutated loci per genome, offers the advantage of producing a large number of functional variants in any genotype, potentially enabling researchers to observe and study more mutant traits. IRRI has a large collection of deletion mutants produced from the agronomically important indica cultivar, IR64. The mutants are screened and characterized for observable agronomic phenotypes. This is currently the most comprehensive indica mutant resource to date and seeds of the mutants have been distributed widely to many rice research groups (Leung et al., 2001, Wu et al., 2005).

IRRI has a collection of rice mutants derived from *indica* variety IR64, and generated by various chemical- and irradiation mutagenesis (Leung et al., 2001, Wu et al., 2005). Some of these mutants exhibit enhancement of resistance, with corresponding differential expression of DR-related genes vs. the wild type. Mapping populations are also available for these mutants. Furthermore, advanced lines with mapped resistance genes, developed from previous genetic studies (such as SHZ-2/LTH varietal cross from Bin et al., 2004) are available for functional genomics research.

The particular focus of this study for gene expression analysis is the rice mutant designated as GR978, which exhibits gain of resistance against rice blast. To facilitate accurate and unambiguous characterization of observable mutant traits in the mutant collection where GR978 was obtained, a controlled vocabulary set was developed to describe the mutant phenotypes.

3. *Bioinformatics and statistical methods/tools for gene expression analysis.* DNA microarrays have been touted, from its inception in 1995, as having the potential to shed light on cellular processes by identifying groups of genes that appear to be co expressed (Quackenbush 2003). Most microarray experiments focus on identifying patterns of gene expression in a particular system, for example, analyzing gene expression responses to a stress challenge. Consequently, there is a greater demand for statistical assessment of the conclusions drawn from microarray experiments (Churchill 2002). Fundamental issues such as how to design an experiment to ensure that the resulting data are amenable to statistical analysis have led to more sophisticated microarray experimental designs encompassing increasingly broad surveys of diversity within a system. As a result, genes whose patterns of expression are identified as being statistically significant can be assigned a greater degree of confidence. Yang and Speed (2002) discussed three main issues in the design of microarray experiments, and devised a checklist that enables researchers to answer the scientific questions that microarray experiments intend to address while keeping in touch with the practical (logistic, economic) aspect.

Expression data pre-processing is another key issue in analyzing microarray experimental results. Data transformation in order to normalize the dataset is required so that meaningful biological comparisons can be made. There are a number of reasons why data must be normalized, including unequal quantities of starting RNA, differences in labeling or detection efficiencies between the fluorescent dyes used, and systematic biases in the measured expression levels (Quackenbush 2002). Three of the most used transformation and normalization categories are LOWESS, intensity-dependent filtering, and replicate filtering. After data pre-processing, the next statistical tests intend to establish which genes are differentially expressed between the RNA species being compared at a statistically determined significance level. Pan (2002) compared 3 methods: the t-test, a regression modeling approach, and a mixture model approach, and also did comparisons with a Bayesian approach and the Significance Analysis of Microarray (SAM) method by Tusher (2001). The choice of statistical analysis is determined by several factors, such as the availability of experimental replications, as well as the underlying biological nature of the problem such as the actual number of genes involved in the contrasting phenotypes observed (small vs. large number) and should be considered beforehand.

The true power of microarray analysis does not come from the analysis of single experiments, but rather, from the analysis of many hybridizations to identify common patterns of gene expression (Quackenbush 2002). For cellular processes, genes that are contained in a particular pathway or that respond to a common environmental challenge, should be co-regulated and consequently, should show similar patterns of expression. The identification of genes that show similar patterns of expression can be achieved using a large group of statistical methods, generally referred to as ‘CLUSTER ANALYSIS’ (Quackenbush 2001. Two broad categories of clustering algorithms exist; the first is the UNSUPERVISED method. Hierarchical clustering (HCL) is a very popular algorithm under this category. Eisen et al. (1998) used this to cluster gene expression patterns in the budding yeast *Saccharomyces cerevisiae*, and found that clustering gene expression data groups together efficiently genes of known similar function, and also found a similar tendency in human data. Hence, HCL is the most widely used clustering method in use today. In the presence of advanced knowledge about the number of clusters that should be represented in the data, k-means clustering is a good alternative to HCL, and this was used by Soukas et al. (2000) in adipose tissue gene expression studies. Principal components analysis is another algorithm that was used by Raychaudhuri et al. (2000) for time series experiments. Self-organizing maps (SOM) is another approach; this is a NEURAL-NETWORK-based divisive clustering approach (Kohonen 1982). For gene expression clustering, a SOM assigns genes to a series of partitions on the basis of the similarity of their expression vectors to reference vectors that are defined for each partition (Tamayo et al. 1999).

SUPERVISED clustering methods are a powerful alternative to the previous methods mentioned, such that users can incorporate biological information into the algorithm. The most popular algorithm in use is the Support Vector Machine (SVM). SVMs use a training set, with prior biological information, in which genes known to be related by, for example function, are provided as positive examples and genes known not to be members of that class are negative examples. The SVM can then be used to recognize and classify the genes in the experimental data set on the basis of their expression (Brown et al. 2000). Another algorithm that utilizes prior biological information about the gene set under investigation is Expression Analysis Systematic Explorer (EASE-Hosack et al. 2003). EASE utilizes annotation information attached to each gene in the set (from Gene Ontology, Interpro, KEGG, Genbank, or other biological databases) and groups genes into biological themes, based on a statistical significance (either Fisher exact test, or EASE score). The most attractive feature of EASE is that gene groups are named according to the higher order biological theme it belongs to, which is easy for biologists to make sense of.

The use of Gene Ontology (GO - Gene Ontology Consortium, 2004) categories has become very prevalent in characterizing gene expression themes obtained from a list of differentially expressed gene. The GO project is a concerted effort to provide standardized, controlled vocabularies (ontologies) for genome annotation systems. It provides three structured ontologies describing gene products in terms of their associated biological processes, cellular components and molecular functions in a species-independent manner. A GO-annotated genome enables the classification of a subset gene list from the genome with higher-level general functions (biological themes) owing to the organization of each GO term as directed acyclic graphs wherein each term has parent term(s), so navigating to a previous node (the parent) gives less specialized information about a gene. In the GO website (http://www.geneontology.org/GO.doc.shtml), this particular example is given (verbatim): “the biological process term hexose biosynthesis has two parents, hexose metabolism and monosaccharide biosynthesis. This is because biosynthesis is a subtype of metabolism, and a hexose is a type of monosaccharide. When any gene involved in hexose biosynthesis is annotated to this term, it is automatically annotated to both hexose metabolism and monosaccharide biosynthesis, because every GO term must obey the true path rule: if the child term describes the gene product, then all its parent terms must also apply to that gene product”. Therefore, a list of genes may be the child of the same non-specialized GO term, and this unifies this gene list under the same biological description as this parent term.

Other means of obtaining biological meaning from gene expression data is localizing the set of significantly expressed genes into biologically relevant regions of the genome. Spellman and Rubin (2002) analyzed numerous *Drosophila* microarray experiments and found that the genome has over 200 groups of adjacent genes with a common expression pattern (by correlation analysis) but with no obvious functional relationships. These groups are composed of 10 to 30 genes, not related to each other by homology, and span hundreds of kilobases. Ma et al. (2005) did the same analysis using an oligo microarray in rice and found similar domains of co-regulation of many gene models, about 100kb in length.

Segal et al. (2003) presented a probabilistic method for identifying regulatory modules from gene expression datasets. They took a gene expression data set and a large precompiled set of candidate regulatory genes that they curated from yeast, and their algorithm simultaneously searches for a partition of genes into modules and for a regulatory program. The combination of previously known biological information and the probabilistic nature of the method enabled them to identify functionally coherent modules and the correct regulators. Furthermore, the proposed modules and regulators can be statistically tested.

Wisser et al. (2005) used QTL regions in the rice genome associated with disease response to determine significant association of differentially expressed genes with the regions. They integrated publicly available data on QTLs and qualitative R genes with rice genome information and disease-related EST information from rice and other cereal crops in order to nominate statistically tested chromosome regions associated with broad-spectrum quantitative disease resistance (BS-QDR) in rice. The BS-QDR regions are useful biological information to overlay in gene expression studies involving rice: a set of differentially expressed genes of known genome location can be tested for association with these regions and this information can be used to classify gene sets identified in expression studies involving disease resistance in rice.

New statistical models and bioinformatics tools/software that implement new computational models from human and yeast studies have been developed and are freely available via Open Source. These applications were specifically written to handle the problems of immense data quantity generated by microarray experiments, determination significantly expressed genes, and determine association/correlation of a gene or set of genes leading to inference of gene expression networks or metabolic pathways. Furthermore, ordinary high-powered desktop personal computers are now cheap and this has enabled researchers to do complicated and resource-intensive computations on their own computers at minimal cost.

Functional genomics study of rice through transcriptome analysis using the Agilent Technologies 22K rice oligoarray platform was carried out in order to determine the expression profile of the gain-of-resistance rice mutant GR978 in comparison with the wild-type IR64 undergoing a broad-spectrum resistance (BSR) response against rice blast infection. Gene expression profiling of GR978 undergoing disease response against rice blast may result in the (1) discovery of candidate genes responsible for this resistance response, and (2) dissection of the transcription profile associated with BSR (the disease resistance transcriptome).

Gene expression results of a similarly stressed rice variety (SHZ-2) were data-mined and an integrated 2^nd^ order analysis of gene expression data was conducted to characterize the disease resistance transcriptome of rice. The specific objectives of the study are to:

1. Develop a controlled vocabulary describing mutant phenotypes in the mutant collection
2. Identify and characterize significantly expressed genes involved in GR978 BSR
3. Discover previously uncharacterized genes implicated in GR978 BSR
4. Infer the topology of the gene regulatory network of these candidate genes that are associated with the BSR response in GR978, and
5. Determine the associations and inter-relationships of significantly expressed individual genes/co-regulated group of genes across genotypes.

The study was conducted in different locations and laboratories, due to the various capabilities and equipment required. These locations are at the Genetics and Molecular Biology Laboratory at the Institute of Biological Sciences, College of Arts and Sciences, University of the Philippines Los Baños (for genetic and preliminary work), at the Entomology and Plant Pathology Division and Biometrics and Bioinformatics Unit, International Rice Research Institute (IRRI, for greenhouse and field experiments and primary portion of the gene expression experiments, and final gene expression analysis), and the National Institutes of Agrobiological Sciences at Tsukuba City, Japan (for DNA microarray gene expression experiments and preliminary data analysis). The experiments were conducted starting June 25, 2004 (first greenhouse experiment) and completed in December 20, 2005 (last microarray experiment). Data analysis was completed on 25^th^ of May 2006.

## MATERIALS AND METHODS

A combination of greenhouse, laboratory experiments, and bioinformatics were done in order to generate the robust biological dataset needed for in-depth gene expression analysis of the study.

### Plant Materials Used

The rice mutant used in this study, available under the accession ID GR978-18 (GR stands for Gamma Ray induced mutant), referred hereon as GR978, is one of the interesting mutants found in the IR64 mutant bank at IRRI. The original progenitor seeds, designated as IR64-21, were mutagenized by gamma-irradiation and maintained in the mutant collection as described by Wu et al. (2005). M2 families were initially screened for bacterial blight (*Xanthomonas oryzae pv. oryzae, Xoo*) resistance, and M2 family #18 was subsequently found to have qualitative resistance against *Xoo* and moderate, quantitative resistance against leaf blast, using isolate Ca89 for blast resistance screening. Genetic analysis revealed that a recessive gene mutation mapped onto a 30.8 cM region on chromosome 12 conditions the broad-spectrum resistance. (Madamba et al (2009).

The current batch of GR978-18 stocks is at more than M6 generation already, and batches of seed stocks are tested for resistance against isolate Ca89 prior to any use for resistance characterization. All gene expression experiments used this >M6 stock of GR978.

The susceptible variety used for contrast in all gene expression profiling experiments was descended from the original IR64-21 wildtype stock used for generating GR978.

### Treatments with Blast Inoculation (Greenhouse Experiments to Subject IR64 and GR978 Under Water and Rice Blast Stress)

Rice blast interaction was examined in IR64 (wildtype) and GR978 using 21-day old seedlings inoculated with leaf blast fungus under greenhouse conditions. A complete biological replication (biorep) consisted of two blocks, each block with two rice entries subjected to one of the following treatments: (1) inoculation with rice blast isolate Ca89, an isolate that causes a susceptible disease reaction in IR64 but to which GR978 shows moderate resistance, or (2) inoculation with blank. Table 1 summarizes the combination of entries and treatments for each biorep.

**Table 1.**
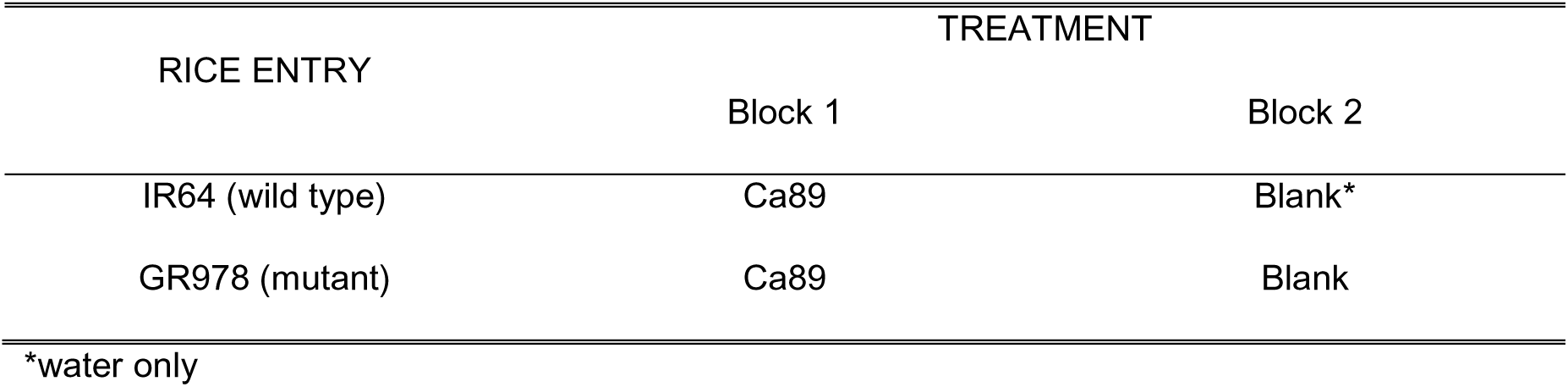
The combination of treatment and rice entries for each biorep made

| RICE ENTRY | TREATMENT |  |
| --- | --- | --- |
|  | Block 1 | Block 2 |
| IR64 (wild type) | Ca89 | Blank* |
| GR978 (mutant) | Ca89 | Blank |

\*water only

Plants were grown in two blocks, each block was a 34 × 27 × 12 cm plastic tray containing approximately 7 kg of sterilized Maahas Clay supplemented with ammonium sulfate incorporated at 6g/kg soil. For each block, two rice entries were planted, each with four rows, and each row with ten plants. The rows were completely randomized. At each sowing three biological replications (six blocks or trays) were made. The first sowing date was June 25, 2004, while the second sowing date was on October 20, 2005. This simple design was adapted owing to the high variation of gene expression attributed to each individual plant.

For each biorep, the two blocks were sprayed separately with either blank inoculum (water and Tween 20) or with Ca89 spore suspension. Inoculum preparation was carried out on the same day of seed sowing, following the protocol described by Chen et al. (1995). Spore suspension was adjusted to a concentration of 2.5 x10^4^ conidia/ml using a hemacytometer. For inoculation, 100 ml suspension was sprayed using an atomizer connected to an air pump for each tray mounted on a rotating platform. After spraying, the trays placed in a dew chamber for 24 hours, and then transferred to the mist room with environment conditions maintained specifically to enhance blast disease development.

Plant tissue was sampled per biorep at 24h after inoculation, by pooling five tillers (cut about 2 cm from the base of each plant) selected at random from different plants of the same treatment-entry combination. Each sample pool was wrapped in aluminum foil and immediate immersed in liquid nitrogen, then transferred to −70°C freezer and kept until the time of RNA extraction. The same sampling protocol was repeated at 48h after inoculation. For each sampling time, twelve sample pools were obtained (1 sample pool per entry × 2 entries per block × 2 blocks (or treatments) per biorep × 3 bioreps). To ensure that samples were obtained from plants exhibiting the correct disease resistance phenotype, disease reaction was assessed per biorep at three to five days after inoculation. Disease reaction of each entry was evaluated using a 0 - 5 scoring system described by Bonman et al. (1986) with 0 - 3 considered as resistant reaction (non-sporulating lesions of pinprick size to pinhead size) and 4 - 5 (sporulating lesions with gray centers) as susceptible reaction.

### RNA Extraction from Leaf Tissue Samples

Total RNA was extracted from each sample pool collected at 24h and 48h after inoculation. Each leaf sample pool was extracted for total RNA using the Trizol™ (Life Technologies Inc., Gaithersburg, MD, USA) protocol. Tissue homogenization was carried out using a mortar and pestle with liquid nitrogen. The leaf tissues were kept frozen until addition of Trizol. Five ml of leaf tissue powder was used for RNA extraction. RNA samples were determined for concentration and quality-checked by electrophoresis in 2% agarose gel in1X MOPS buffer. RNA samples were stored separately in −70°C until needed for microarray hybridization. Eight RNA samples were obtained for each biorep: (1) GR978 inoculated-24h, (2) GR978 water-24h, (3) GR978 inoculated-48h, (4) GR978 water-48h, (5) IR64 inoculated-24h, (6) IR64 water-24h, (7) IR64 inoculated-48h, and (8) IR64 water-48h.

### Microarray Experimental Design: Contrasting Experiments

The two timepoints selected (24h and 48h post inoculation) for sampling RNA from each rice entry subjected to rice blast conditions were empirically determined by Nobuko Sugiyama (pers comm., IRRI) to observe the greatest number of differentially expressed genes was observed between GR978 and IR64. This prior experiment was conducted using a cDNA microarray containing genes implicated in defense response in rice and related cereals (1k cDNA defense response array, IRRI-KSU, results not shown).

In order to maximize the power of statistically detecting differentially expressed genes, the microarray experiments diagrammed in Figure 1 were done.

**Figure 1.**
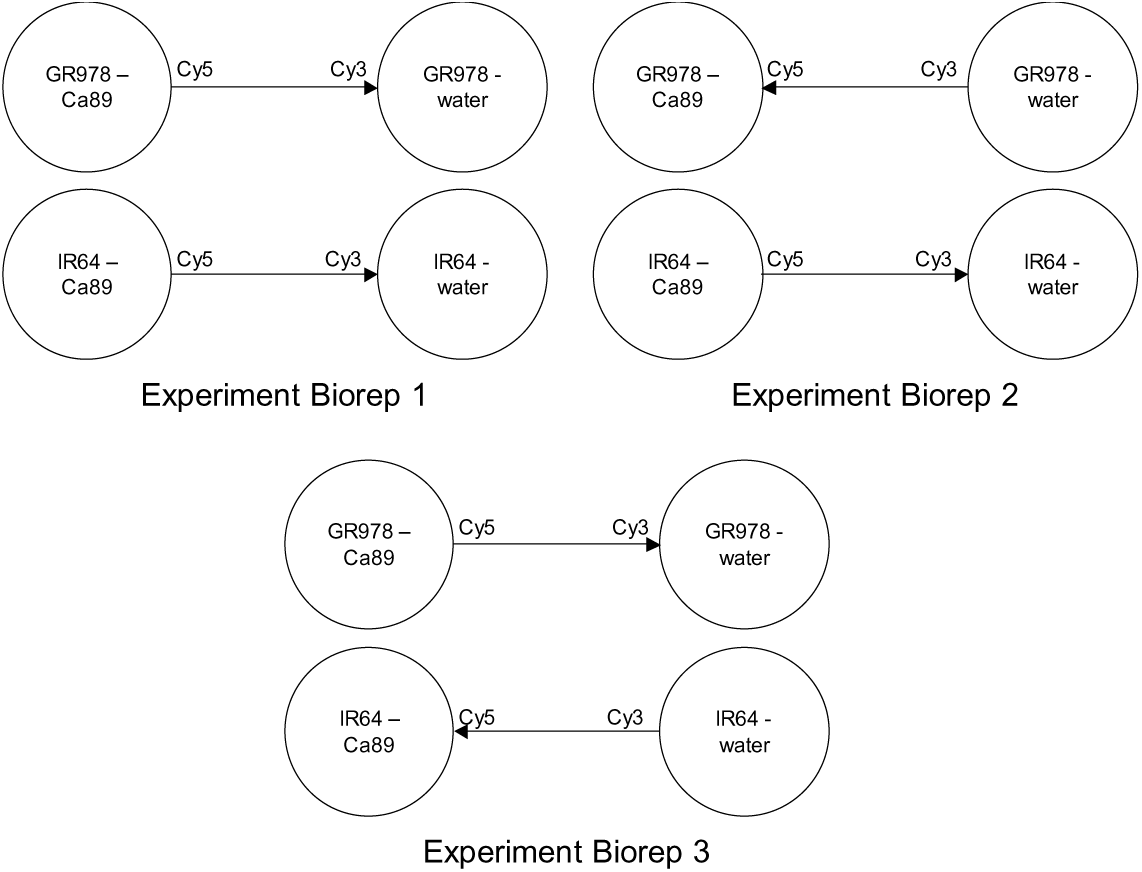
RNA hybridization plan and experimental design. The same design was used for the 24h and 48h timepoint experiments.

One complete experiment consists of two hybridizations (2 slides): one is the GR978 contrasting treatments (GR978-inoc vs GR978-water), and the other is the IR64 contrasting treatments (IR64-inoc vs IR64-water). Two circles connected by an arrow denote one hybridization (1 microarray slide). In experiment biorep1, both GR978 and IR64 hybridizations are straight-labeled (inoculated sample is cy5 labeled). Reverse dye labeling was conducted across replicated experiments, so in experiment biorep2, GR978 hybridization is reverse labeled and IR64 hybridization is straight-labeled; in experiment biorep 3, IR64 hybridization is reverse labeled and GR978 hybridization is straight-labeled. From these experiments, statistical determination of the constitutively differentially expressed genes between rice genotypes, as well as the induced genes, in response to rice blast infection is possible. Effects of gene × dye interaction and slide variation cannot be directly estimated but still can be quantified due to being confounded with biological replication.

### Microarray Wet-Lab Experiments

Microarray experiments were conducted in collaboration with Drs. K. Satoh and S. Kikuchi at National Institute of Agrobiological Sciences (NIAS) in Japan. RNA samples were brought to the gene expression microarray experiment facility located at the Laboratory of Gene Expression (Kikuchi and Satoh, pers comm.) at NIAS. Each RNA sample was prepared by precipitating with two volumes of 100% ethanol and 1/10 volume NaOAc and keeping in −20C for 15 to 30 mins, then spun for 10 mins at 10,000 RPM using a refrigerated microcentrifuge. The supernate was discarded and the pellet was suspended in 0.5 ml of 100% ethanol. The samples were packed in dry ice and transported. At NIAS, the RNA samples were reconstituted in DNAse-free water and prepared for DNA oligoarray hybridization. RNA concentration was determined using the NanoDrop^TM^ Spectrophotometer (NanoDrop Technologies, Wilmington, Delaware USA), following the instruments’ protocol. A working RNA solution of 5ng/uL was prepared for each sample.

The subsequent procedures used were optimized for the Agilent™ (Agilent Technologies Inc., USA) 22K rice oligo microarray system. Being a 2-dye system, the Agilent system requires the differential labeling of the two contrasting RNAs with cyanine (cy) 3 and cyanine 5, and the subsequent competitive hybridization and detection of the two RNA species on a single rice oligoarray slide. Pre and post-labelling RNA quality check was done using the RNA 6000 Nano Assay Protocol and analyzed using the Agilent 2100 Bioanalyzer. cRNA amplification and fluorescent labeling was carried out using Agilent Low RNA Input Fluorescent Linear Amplification Kit (Product number 5184-3523) which is optimized to work with Agilent’s oligoarrays. This protocol first generates cDNA from total RNA by reverse transcription using the T7 promoter primer and MMLV reverse transcriptase, and then synthesizes cRNA with cy 5 or cy 3 dye labels incorporated directly. This method routinely results in at least a 100-fold RNA amplification and the amplification is unbiased by RNA transcript size. Microarray hybridization and stringency washing were done following the Agilent 60-mer oligo microarray processing protocol (6-screw chamber/SSC, manual part number G4140-90010). Array signal detection/scanning was done using the Agilent dual laser microarray scanner (Agilent Technologies DNA Microarray Scanner, Model G2565B) using 532 and 633 nm laser lines) at a 10 micron resolution. Image analysis and spot quantification was done using the Agilent Feature Extraction (FE) software v 8.1.

The DNA oligomers printed on the Agilent Rice Oligo Microarray kit (product number G4138A) are 60 bases long, and were designed from the 28,000 FL-cDNA collection of Kikuchi et al. (2003). Most of the oligos uniquely represent roughly ∼20,000 genes, which is around 50% of the expected transcriptome of rice. With this platform, a successful hybridization experiment is expected to yield expression data for half of all the genes involved in disease resistance response in GR978.

### Data Management System

All raw data from Agilent FE software was preprocessed by simple removal of extraneous headers then stored and managed using a relational database management system (RDBMS). The RDBMS of choice was custom relational tables implemented under either Microsoft Access or MySQL, and databases were implemented for both platforms. To facilitate all database-related activities, the assigned oligo names for each of the ∼22,000 genes/features on the 22K oligoarray was used as unique gene ID; this is also the same Genbank ID used to identify the corresponding FL-cDNA where each oligo was uniquely designed from. This gene ID will subsequently be referred to as KOME ID and each gene in the oligoarray as KOME gene. The minimum contents of the RDBMS are the gene expression table and the KOME gene annotation table.

### First-Order Gene Expression Data Analysis: Characterization of the Resistance Transcriptome Within Genotypes

Characterization of the resistance transcriptome was done within the genotype under study (GR978) and across disease-resistant genotypes/varieties in which data is available for analysis, such as cultivar SHZ-2. The first-order analysis refers to methods that use the gene expression within one microarray experiment only. Second-order analysis combines data from several microarray experiments (GR978-SHZ2 combined dataset), as well as secondary data such as genome positions for analysis. The methods in this section refer to the general methods used for each type of analysis. The specific problems to which these methods were applied to are discussed in detail in the Results and Discussion section of the manuscript. At this point, all activities were computer-based, using appropriate software tools (preferably free or Open Source software) for data handling and analysis. Custom applications/scripts were written using Perl and SQL as needed, and these are mentioned in other sections.

#### Data pre-processing and transformation

The raw quantified data from image analysis was extracted from the RDBMS and pre-processed to prepare the data for subsequent statistical analyses. The first step was spot/feature quality check, where all genes considered as bad spot (missing, saturated, too irregular spot shape, too high background, too low intensity, flagged by the Agilent FE spot quantification output file in the following manner: not true (0) for IsSaturated, IsFeatNonUnifOL, IsBGNonUnifOL, and IsFeatPopnOL, and true (1) for IsPosAndSignif and IsWellAboveBG) is given a single bad spot flag symbol (X). Other extraneous information not needed for expression analysis is likewise filtered out. This filtering is implemented using a custom SQL script (Appendix script 1).

Data transformation of the raw signal intensities is the next step in data pre-processing. This was again done by the Agilent Technologies FE Software v 8.1 using local LOWESS (locally weighted linear regression) normalization, which is performed prior to outputting the processed signal intensities of the red and green channels (cy5 and cy3 labeled samples). The LOWESS dye normalization factor is calculated by fitting the locally weighted linear regression curve to the chosen normalization features. The amount of dye bias is determined from the curve at each feature’s intensity. Each feature gets a different LOWESS dye normalization factor per channel. The expression dataset is reduced into informative spots from this step, and is stored either as TIGR Multiple Experiment Viewer formatted intensity files (Saeed et al. 2003), or log2 ratio matrix file (Stanford format), both of which are considered as standard gene expression file formats, for each experiment, which ready for further statistical analyses.

#### Statistical determination of significantly expressed genes

Determination of significantly differentially expressed genes is the most critical step in any microarray experiment analysis, especially when dealing with 22,000 genes at once; a type 1 error for a p-value cutoff of 0.01 still translates to hundreds of genes being erroneously listed as significantly expressed. Differentially expressed genes (DEGs) were determined individually at each time point using two methods: (1) independent t-tests (Pan 2002) and (2) mixed model ANOVA using a split plot design implemented by Microarray Analysis of Variance (MAANOVA, Wu et al. 2002). The independent t-test for each experiment is able to detect genes to be significantly expressed even with low expression ratios, because unknown error variance components are estimated based on a gene-specific variance for each gene. For the independent t-tests, the significant DEGs were first determined for each rice entry-treatment comparison (eg. GR978 inoculated vs water, 3 bioreps) using one-class t-test for each gene. The mean of gene expression (log2 ratio) for 3 bioreps is tested for significant difference against zero (no difference in gene expression between samples compared). Significant DEGs between the resistant and susceptible entries were then tested using two-class t-test, with GR978 as class A and IR64 as class B, using 3 bioreps. Each t-test generates a list of significant and non-significant genes. At each timepoint, from the DEGs set, the subsetsof genes that were significantly differentially induced (DI genes) by blast inoculation in GR978 were determined by the Boolean join:

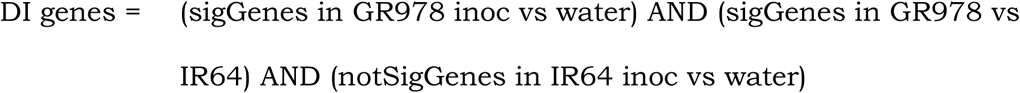

To determine the constitutively differentially expressed genes (CDE genes), the following Boolean join was used:

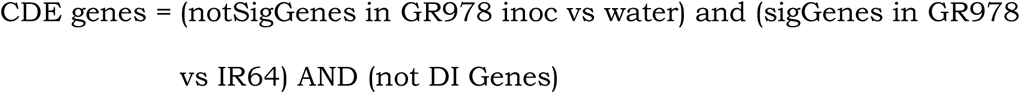

All t-tests were done using significance α= 0.01 with adjusted Bonferroni correction, and p-value based on 1,000 times permutation, as implemented by TIGR Multiple Experiment Viewer v 3.1, available at http://www.tm4.org/. Table joins to determine the CDE and DI genes were implemented using custom SQL statements either in Microsoft Access or MySQL.

Significance analysis in MAANOVA uses a weighted combination of the global variance (mean variance of all genes) and gene-specific variance (Cui 2003, 2005). Thus, low expression genes may not be detected by MAANOVA and the subsequent gene list tends to be more conservative. Mixed model MAANOVA following the proposed model of Wolfinger (2001) was modeled as:

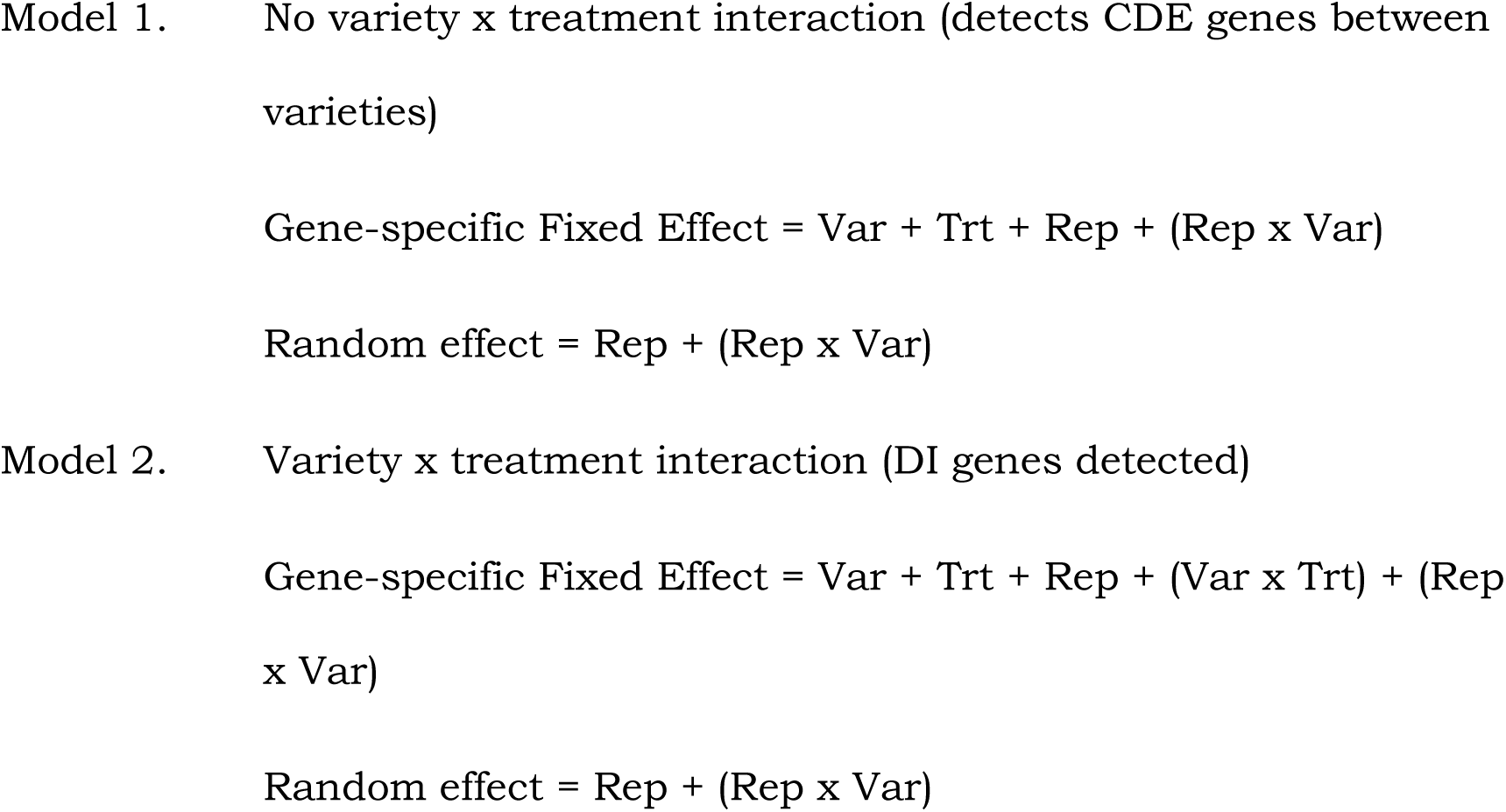

Where *Var* is the two varieties contrasted (IR64 wild type, GR978), *Trt* is the treatment contrasts (inoculated, water), and *Rep* is the biological replication, which also represents the array slides. No specific term for dye was used since it was confounded with the array, due to occurrence of dye swaps with bioreps. Note that the MAANOVA models directly compute for the CDE and DI genes. MAANOVA was implemented in the R Environment using the R/MAANOVA package, downloaded from http://www.jax.org/staff/churchill/labsite/software/Rmaanova/index.html and the script implementing the analysis is given at Appendix script 2.

#### Determining gene coexpression by purely computational methods: hierarchical clustering

In the absence of strong *a-priori* information about the genes under study, the short-listed genes from significance analysis can be grouped for similar expression profiles using unsupervised approaches to data analysis. Hierarchical clustering is the most popular of these methods which, for a short gene list, can be very effective in inferring function of genes otherwise annotated as having an unknown function, by occurrence of these unknowns in a common cluster with genes of known function. DEGs were analyzed by HCL using log2 ratios for the 24h and 48h timepoints, using the HCL module of TMEV 3.1 software; the distance metric used was Euclidean distance, and the linkage method selected was average linkage.

#### Biological themes in the differentially expressed gene list using GO terms

Biological interpretation of the short list of differentially expressed genes is demanding and laborious, especially if this is manually done one gene at a time. One systematic approach to elucidating biological themes from a gene list is by examining the short list against a predefined set of genes representing a known biological theme or metabolic pathways and determined whether any set is over-represented in the short list compared with the whole gene list (Tian et al., 2005). Such over-representation or enrichment analysis is at the core of many ontological analysis tools available to test significant relationship.

The GO consortium (2004) has manually annotated genes in *Arabidopsis* for higher-order functional categories, using controlled vocabulary. The GO annotation of the reference set of the *Arabidopsis* proteome and homology mapping of the KOME genes to the *Arabidopsis* proteome provides the reference set to which biological themes can be inferred automatically from a short list of KOME genes from gene expression data obtained from the Agilent 22K oligoarray platform. Enrichment of a particular GO category can be computed for a short list of KOME genes against the whole GO annotation for the entire ∼22,000 genes in the oligoarray using the one-tailed Fisher exact test, as described by Hosack et al. (2003) and implemented in the EASE stand-alone system (http://david.niaid.nih.gov/david/ease/help.htm). The EASE system was customized to work with the KOME gene set in the 22K oligoarray by loading two files: the master list of KOME IDs represented in the oligoarray (21, 495 in all) and a file that maps each KOME ID to a particular GO vocabulary (available at http://cdna01.dna.affrc.go.jp/cDNA/). GO category enrichment was computed by inputting the subset list of genes of interest and using the GO-KOME category system to compare against.

#### Inferring regulatory modules in the highly expressed genes: motif enrichment analysis

For the short list of genes, investigating upstream regulatory *cis*-elements might reveal higher order grouping such as sets of gene under the control of common regulatory motifs, as well as putative gene interactions effected by these regulatory motifs. The 1 kb upstream sequences of all KOME genes in the oligoarray were obtained via the TIGR rice pseudomolecule assembly rel. 3.0, using the mapping information of each KOME FLcDNA with the corresponding TIGR gene model. This information can be downloaded using the rice genome browser at the same website. All known cis acting promoter elements in the release 24 of the Plant Cis-acting Regulatory DNA Elements database (PLACE) were downloaded. Each of the 1kb upstream sequences was scanned for perfect match alignments with all the motif sequences found PLACE database using the program *tfscan*, a part of the EMBOSS software suite (Rice et al., 2000).

For a subset list of DEGs, enrichment of a particular motif was determined by a test for fixed ratio using the Chi square test. For a particular motif, the ratio of the count of this motif (*m’*) to the total count of motifs (*M’*) within the DEG list was computed and this was compared against the ratio of count of the same motif (m) to the total count of motifs in the genome (M). This analysis was repeated for each motif found in the DEG list. The details of the enrichment analysis are described in the *Association analysis general methodology* part of this section.

The annotation of the enriched motifs were further inspected and categorized according to relationship to stress responses, based on literature mining of previous published experiments. After motif enrichment analysis, KOME genes to motif association was determined by inspection to establish the set of gene under the control of common motifs, as well as the set of motifs that control a common gene, thus inferring the regulatory modules of the resistance transcriptome.

#### Determining the structure of correlation of gene expression along the rice genome

The correlation of gene expression profiles was analyzed during biotic stress response of GR978 against rice blast for a moving genome window using the method described by Spellman and Rubin (2002) for identifying coregulated, adjacent gene models. Six independent microarray experiments comparing blast infected vs water-inoculated GR978 were used for analysis. Adjustments made to the method include the selection of appropriate window size for correlation analyses from two genes up to 15 genes per window, for the whole KOME geneset in the 22K oligoarray. For the optimally determined gene window size, the number of genes and gene groups showing co-expression were calculated at p values 0.0001, 0.0005, 0.001 and 0.01. Sample data and the original Perl scripts implementing the method are available at http://www.fruitfly.org/expression/dse/data.shtml.

### Second-order Analysis of Gene Expression Data: Characterizing the Resistance Transcriptome Beyond one Genotype

The resistance transcriptome was further characterized using gene expression data from other materials undergoing resistance response, as well second-order data such as genome locations for QTLs, regions of correlated expression, extended gene annotations, among others.

#### Gene expression data sets from other plant materials

Gene expression raw data was obtained from two gene expression studies. The first study is gene expression profiling of a resistant rice cultivar, Sanhuangzhan 2 (SHZ-2) contrasted with the susceptible cultivar Lijiangxin-tuan-heigu (LTH) in response to rice blast infection. SHZ-2 is a durably blast resistant variety; five QTLs and a major gene were characterized by Bin et al. (2004) to contribute to the disease resistance phenotype. Access to phenotype and gene expression data was provided courtesy of Dr. Liu Bin of the International Rice Research Institute. All microarray experiments done for SHZ-2 used the same Agilent Technologies 22K oligoarray platform and system. The gene expression experiments follows the same design, greenhouse and laboratory protocols described in the sections pertaining to microarray experimentation, with the exception of the rice blast isolate used, which was isolate PO6-6.

The second study used for data mining is the Generation Challenge Program project number SP2 9, entitled “Systematic evaluation of rice mutant collections for conditional phenotypes with emphasis on stress tolerance”. Access to partial gene expression data was provided courtesy of Dr. Hirohiko Hirochika of the National Institute of Agrobiological Science at Tsukuba City, Japan. In this study, the same Agilent 22K oligoarray system was used to identify genes involved in Benzothiadiazole-induced defense response. BTH is effective in enhancing resistance of rice against *Rhizoctonia solani* and *Magnaporthe grisea* and the gene expression profiles in BTH-induced Nipponbarre cultivar were following BTH application to rice plants

#### Interesting genome regions for association analysis of the resistance transcriptome

Blast resistance QTLs were located in the TIGR release 3.0 of the rice genome pseudomolecule using the positions of disease resistance genes provided by Randall J. Wisser (pers comm., Cornell University) as follow-up to their paper in 2005 where they characterized regions of the rice genome associated with broad-spectrum quantitative disease resistance. From an original number of 210 disease-resistance QTLs compiled, 94 rice-blast specific QTLs (Bl-QTL) were selected, this collection represents 8 rice blast QTL studies. Since the intervals were defined by the location of flanking markers in the pseudomolecule, most of the Bl-QTLs were small regions. Each interval was increased to an approximate interval size of 5cM (1.215mb equivalent), and the new interval sizes were manually inspected for overlaps and proximity. Intervals that are sufficiently close to any adjacent interval (distance < 1 mb) were merged as one meta-interval. This was repeatedly done for all intervals until all resulting intervals were spaced at least 1 mb. The merged Bl-QTLs represent 40 regions spanning ∼102mb (Table 2) with a resolution of ∼5 cM.

**Table 2.** The Bl-QTL meta-intervals defining blast-resistance QTLs in the rice genome

| CHR | BL-QTL REGION | QTL START | QTL STOP |
| --- | --- | --- | --- |
| 1 | 1 | 0 | 5,700,756 |
| 1 | 2 | 6,836,178 | 10,264,018 |
| 1 | 3 | 13,996,510 | 15,211,510 |
| 1 | 4 | 19,971,770 | 21,186,770 |
| 1 | 5 | 28,482,780 | 33,641,697 |
| 1 | 6 | 37,086,761 | 40,275,724 |
| 2 | 7 | 21,502,272 | 22,717,272 |
| 2 | 8 | 26,392,777 | 27,607,777 |
| 2 | 9 | 33,957,861 | 36,220,610 |
| 3 | 10 | 0 | 2,013,580 |
| 3 | 11 | 22,181,443 | 25,697,260 |
| 3 | 12 | 27,598,156 | 28,813,156 |
| 4 | 13 | 7,820,818 | 9,035,818 |
| 4 | 14 | 10,465,251 | 11,680,251 |
| 4 | 15 | 19,131,926 | 21,385,381 |
| 4 | 16 | 28,106,567 | 29,321,567 |
| 4 | 17 | 30,480,112 | 32,454,582 |
| 5 | 18 | 5,423,448 | 6,638,448 |
| 5 | 19 | 20,837,495 | 22,052,495 |
| 6 | 20 | 1,158,592 | 2,373,592 |
| 6 | 21 | 6,111,224 | 7,326,224 |
| 6 | 22 | 8,929,630 | 10,144,630 |
| 6 | 23 | 23,768,532 | 24,983,532 |
| 6 | 24 | 28,080,383 | 31,509,048 |

Table 2. continued.
| CHR | BL-QTL REGION | QTL START | QTL STOP |
| --- | --- | --- | --- |
| 7 | 25 | 929,530 | 2,144,530 |
| 7 | 26 | 6,624,174 | 7,839,174 |
| 7 | 27 | 15,413,507 | 27,494,770 |
| 7 | 28 | 28,862,907 | 30,077,907 |
| 8 | 29 | 3,492,913 | 8,232,952 |
| 8 | 30 | 15,633,829 | 16,848,829 |
| 9 | 31 | 7,686,488 | 12,892,156 |
| 9 | 32 | 13,988,708 | 15,203,708 |
| 9 | 33 | 17,059,850 | 22,471,720 |
| 10 | 34 | 10,414,976 | 13,072,332 |
| 11 | 35 | 15,803,079 | 20,094,087 |
| 11 | 36 | 22,130,956 | 25,143,546 |
| 12 | 37 | 0 | 2,202,570 |
| 12 | 38 | 5,150,860 | 8,390,150 |
| 12 | 39 | 10,487,232 | 11,702,232 |
| 12 | 40 | 17,333,230 | 19,470,748 |

#### Combining expression data sets and analysis results using GR978, SHZ-2 and BTH-Nipponbarre data

All first order analyses were also done to the SHZ-2 expression data set in order to come up with a combined list of DEGs and common regions of correlated expression.

In the case of combined DEGs, DEGs from GR978, SHZ-2, and BTH-induced Nipponbarre studies were computed independently for each study using the methods mentioned before. Within the same study, DEGs detected at different time points were combined into a single list by a union operation. DEGs from different studies were combined by an intersection operation.

Determination of regions of correlated expression in GR978 and SHZ-2 was carried out by combining the log2ratio expression data matrices from GR978 and SHZ-2 studies. The combined dataset was analyzed as a single data set.

#### Association analysis: general methodology

Many cases of association analysis were encountered in this study. The motif enrichment analysis described in a prior section utilizes association analysis. Consider this case: we wanted to know if DEGs are enriched (associated positively) within a specific genome region of interest, for example QTL regions. In this case, the question was re-phrased as:

Question: In a QTL region with *n* genes within that region, is the proportion of DEGs in that region *n* significantly different than proportion of all DEGs in the whole genome under study?

We need to know four things: (1) how many genes are in the genome-wide set, assign this to *N*; (2) how many DEGs in all, genome-wide, assign this to *D*, (3) how many genes in the subset under comparison (QTLs), assign to *n*, and (4) how many DEGs within this subset, assign to *d*. The test of fixed ratio applies to this case, and the 2 × 2 contingency table (grayed area) is set-up as:

**Table 3.**
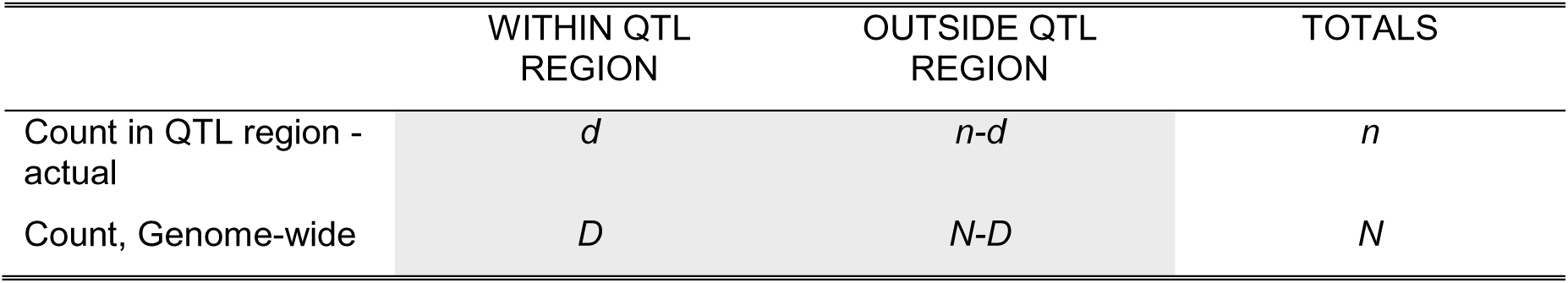
The 2 × 2 contingency table (grayed area) for the association analysis of DEGs within a QTL region

Since we are testing for fixed ratios, we adjust the genome-wide counts so that the totals in the row for genome-wide count equals *n*. We recompute for the adjusted row values; the new *D* value, assign as *D’* is:

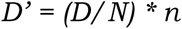

Therefore, the new 2 × 2 contingency table is:

**Table 4.** The 2 × 2 contingency table (grayed area) for the association analysis of DEGs within a QTL region, with adjusted genome-wide counts

|  | WITHIN REGION | OUTSIDE REGION | TOTALS |
| --- | --- | --- | --- |
| Count in QTL region - actual | $d$ | $n-d$ | $n$ |
| Genome-wide count, adjusted | $D'$ | $n - D'$ | $n$ |

Then we do Fisher exact test or Chi square test for fixed ratios in the QTL region vs genome-wide ratio (*d : n-d vs D’ : n-D’*). A low p value (below 0.05) indicates that the two ratios are significantly different. The enrichment value is then computed from the ratio of *d/D’*, obviously there is enrichment within the subset region if the ratio is greater than 1.

In the case of motif enrichment analysis, the counts of motifs (in the gene list of interest and at the genome level) were used for the ratios compared. The method is also easily applied to the following cases:

a. whether DEGs are statistically associated with the regions of correlated expression in the genome (the posed question is: is the ratio of DEG in correlated gene group subset the same as the ratio of all DEGs to the whole genome?),
b. whether correlated regions of expression are associated with QTL regions (is the ratio of correlated genes within QTL region the same as the ratio of correlated genes to the whole genome?).

A Fisher exact test is implemented and available online at http://www.matforsk.no/ola/fisher.htm.

## I. PHENOTYPIC EVALUATION OF RICE MUTANTS AND THE IMPLEMENTATION OF A RICE MUTANT CONTROLLED VOCABULARY

### Results

The use of mutants with discrete genetic lesions has been essential in determining gene functions and dissecting biochemical pathways (Wu et al. 2005). With the completion of the genome sequences of indica and japonica rice cultivars, and the most recent release of the rice gene models by TIGR (release 4.0), mutant resources will prove to be invaluable in assigning gene functions to many of these gene models with yet unknown functions.

The generation and availability of IR64 mutants from the International Rice Research Institute is described by Leung et al. (2003) and Wu et al. (2005). A range of visible agronomic mutations were observed through more than 11 seasons of planting in the field, as well as some interesting conditional mutants. GR978 is one of the interesting mutants obtained from this resource, exhibiting broad spectrum gain of resistance to bacterial and fungal pathogens (Madamba et al. 2009).

For each season of planting and evaluation of the mutants, entries with observable agronomic mutant phenotypes (deviating from wild-type IR64) were described either by using Standard Evaluation System descriptors or specialized terms used by rice mutant researchers (A. Bordeos and M. Baraoidan, pers comm., IRRI). Through the many seasons of screening for observable agronomic mutant traits, a relatively small set of agronomic mutant phenotypes were observed to occur repeatedly (Baraoidan et al. pers comm.). At season 11 of planting and agronomic evaluation, all observed mutant phenotypes for ∼30,000 mutant entries planted were compiled into a list, and 3,732 entries were found to exhibit at least one observable agronomic mutation. The list of mutant phenotypes was inspected for redundancy, occurrence of synonyms and evaluated for ambiguity.

Synonymous or redundant terms determined to be describing the same phenotype were grouped together, and assigned a temporary description. The new phenotype list was further rationalized until all descriptions in the list were found to be unique from one another. In order to properly and consistently describe the mutant phenotypes in the rationalized list, through the different seasons of screening, as well as recognize synonymous mutations observed in different mutant studies, a set of controlled vocabulary (CV) for IRRI internal use, documenting the observed mutant phenotypes was established. The set of vocabulary in use from the agronomic mutant observations were compiled and curated, and 86 distinct agronomic mutant phenotypes were listed.

In 2003, in collaboration with the TOS17 mutant group (Miyao, pers. comm.), the list of controlled vocabulary from the TOS17 mutant phenotypes was merged with the IRRI IR64 mutant phenotype controlled vocabulary. Fifty-six (56) distinct mutant phenotype observations from TOS17 mutants and the 86 IR64 CV terms were rationalized and each term was further refined to be more accurate and compliant to published plant CVs. The merger resulted in a new set of 91 CV terms (Table 6). The composition of the CV is illustrated in the following Venn diagram (Figure 2).

**Figure 2.**
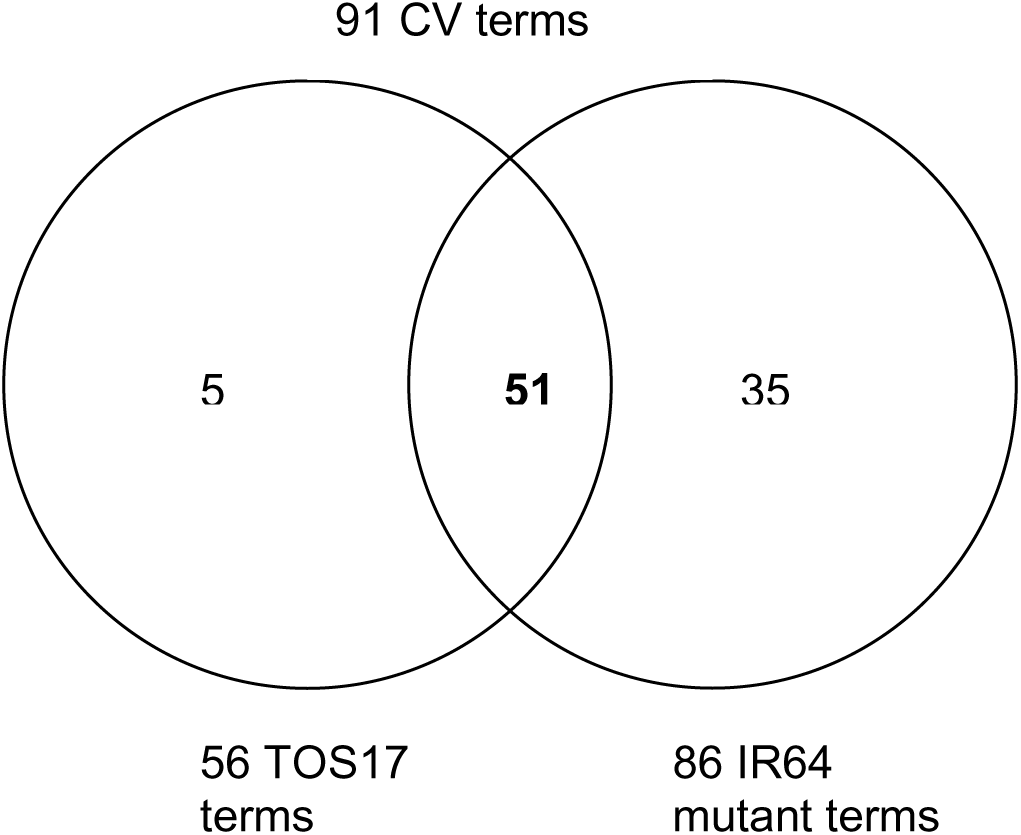
Count of common and distinct TOS17 and IR64 mutant CVs for the merged 91 CV terms

A particular CV term describing a mutant phenotype consists of trait term and trait value term; for example in the CV term *early heading time*, the trait is *heading time*, and the trait value is *early*. Terms used in these trait and trait-value pairs were obtained from mutant descriptions used by previously published work on rice mutants, the Standard Evaluation System for rice (SES, IRRI), and the Trait Ontology database. Novel mutant phenotypes that were discovered are described using combinations of pre-existing Trait Ontology or SES terms, and new terms (nouns or adjectives) were used if there is no previous report of the term describing the observation. Each CV is assigned a unique ID and mapped back to the original synonymous terms in the 56 TOS 17 mutant descriptions in use by Miyao et al. (2003). This unique ID provides the mapping of synonymous mutant descriptions to this public rice mutant database, an important step in avoiding ambiguity in describing mutant traits.

The mutant CV can be informally divided into two classes: the first class consists of CV terms that describe common agronomic traits in reference to the IR64 wild type appearance or a qualitative measurement relative to IR64 wild type. For example, the CV *red basal leaf sheath color* (ID=7) is an appearance that is described relative to the wildtype IR64 green basal leaf sheath color. An example of a mutant phenotype that is a measurement compared relative to IR64 is *long seed length* (ID=67), it is not reported as a quantitative measurement but a comparison to the average IR64 wildtype seed length. In this CV class, 51 entries were observed: 27 of which are found in SES resource, and 24 are non-SES descriptions, 19 of which map to the Trait Ontology database (at least the trait maps, but not necessarily the trait value).

The second class of CVs consists of 40 terms that were lifted from mutant-specific studies in rice. The constitution of this class is shown in Table 5.

**Table 5.** The count of trait values constituting the mutant-specific CV

| CV | COUNT OF TRAIT VALUES |
| --- | --- |
| Leaf blade color | 11 |
| Seed-related trait | 8 |
| Leaf-related trait | 6 |
| Mimic response | 4 |
| Plant height | 3 |
| Stem strength | 2 |
| Panicle-related trait | 2 |
| Vegetative vigor | 1 |
| Germination rate | 1 |
| Growth | 1 |
| Flower/floret organ | 1 |

The top five most numerous in the CV reflect the number of rice mutant studies conducted for each of these mutant traits. Many studies have described leaf color mutations with very unique terms such as chlorina, virescent, zebra, among others. One of the more studied mutant traits being studied for associated gene function is the mimic response trait.

The detailed description of the mutant phenotype CV implemented at IRRI is shown in Table 6. The terms under columns <TRAIT> and <TRAIT value> are the CV pairs recorded as observations for the mutant entries. Each trait CV is decomposed as an *observable* and an *attribute*, to enable mapping to the Plant Ontology (PO) and Open Biological Ontologies project (OBO) databases with the corresponding IDs. Each trait is also mapped to the Trait Ontology (TO) database, and when a synonym is found, the corresponding TO ID is associated as well. The phenotype ID from the TOS17 database is given as well for terms that are synonymous with the IRRI CV.

**Table 6.** The mutant phenotype CV posted at the IRIS-Mutant website

| PHENOTYPE ID | OBSERVABLE | PO TERM ID | ATTRIBUTE | OBO TERM ID | TRAIT | TO TERM ID | TRAIT VALUE | TOS17 ID | TOS17 CLASS | TOS17 PHENOTYPE |
| --- | --- | --- | --- | --- | --- | --- | --- | --- | --- | --- |
| 1 | Awning | PO:0006032 | shape | PATO:0000130 | Awning (An) | TO:0000072 | long and partly awned | 104 | Seed | Others |
| 2 | Awning | PO:0006032 | shape | PATO:0000130 | Awning (An) | TO:0000072 | short and partly awned | 104 | Seed | Others |
| 3 | Awning | PO:0006032 | shape | PATO:0000130 | Awning (An) | TO:0000072 | awned | 104 | Seed | Others |
| 4 | Awning | PO:0006032 | shape | PATO:0000130 | Awning (An) | TO:0000072 | long awn | 104 | Seed | Others |
| 5 | Awning | PO:0006032 | shape | PATO:0000130 | Awning (An) | TO:0000072 | short awn | 104 | Seed | Others |
| 6 | Basal leaf sheath | PO:0020040 | color | PATO:0000033 | Basal leaf sheath color (BLSC) | TO:0000367 | purple | 0 | -- | -- |
| 7 | Basal leaf sheath | PO:0020040 | color | PATO:0000033 | Basal leaf sheath color (BLSC) | TO:0000367 | red | 0 | -- | -- |
| 8 | Basal leaf sheath | PO:0020040 | color | PATO:0000033 | Basal leaf sheath color (BLSC) | TO:0000367 | dark brown | 0 | -- | -- |
| 9 | Culm | PO:0009047 | angle | PATO:0000124 | Culm angle (CmA) | TO:0000427 | erect | 0 | -- | -- |
| 10 | Culm | PO:0009047 | angle | PATO:0000124 | Culm angle (CmA) | TO:0000427 | spreading | 63 | Tillering | Lazy |
| 11 | Culm | PO:0009047 | angle | PATO:0000124 | Culm angle (CmA) | TO:0000427 | procumbent | 0 | -- | -- |
| 12 | Culm | PO:0009047 | diameter | Unknown | Culm diameter | Unknown | big | 101 | Seed | Large grain |
| 13 | Culm | PO:0009047 | diameter | Unknown | Culm diameter | Unknown | fine | 0 | -- | -- |
| 14 | Culm | PO:0009047 | senescence | Unknown | Culm senescence | Unknown | premature | 0 | NA | NULL |
| 15 | Flag leaf | PO:0020103 | length | PATO:0010077 | Flag leaf length | Unknown | long | 104 | Seed | Others |
| 16 | Heading | GRO:0007044 | relative time | PATO:0000147 | Heading time | TO:0000137 | early | 65 | Heading date | Early heading |

Table 6. continued...
| PHENOTYPE ID | OBSERVABLE | PO TERM ID | ATTRIBUTE | OBO TERM ID | TRAIT | TO TERM ID | TRAIT VALUE | TOS17 ID | TOS17 CLASS | TOS17 PHENOTYPE |
| --- | --- | --- | --- | --- | --- | --- | --- | --- | --- | --- |
| 17 | Heading | GRO:0007044 | relative time | PATO:0000147 | Heading time | TO:0000137 | late | 66 | Heading date | Late heading |
| 18 | Internode | PO:0020142 | length | PATO:0010077 | (Uppermost) internode length | TO:0000145 | short | 0 | -- | -- |
| 19 | Leaf | PO:0009025 | angle | PATO:0000124 | Leaf angle (LA) | TO:0000206 | erect | 0 | -- | -- |
| 20 | Leaf blade | PO:0020039 | color | PATO:0000033 | Leaf blade color (LBC) | TO:0000326 | albino | 11 | Leaf color | Albino |
| 21 | Leaf blade | PO:0020039 | color | PATO:0000033 | Leaf blade color (LBC) | TO:0000326 | chlorina | 12 | Leaf color | Yellow |
| 22 | Leaf blade | PO:0020039 | color | PATO:0000033 | Leaf blade color (LBC) | TO:0000326 | dark green | 13 | Leaf color | Deep green leaf |
| 23 | Leaf blade | PO:0020039 | color | PATO:0000033 | Leaf blade color (LBC) | TO:0000326 | light green | 14 | Leaf color | Pale green leaf |
| 24 | Leaf blade | PO:0020039 | color | PATO:0000033 | Leaf blade color (LBC) | TO:0000326 | virescent | 15 | Leaf color | Virescent |
| 25 | Leaf blade | PO:0020039 | color | PATO:0000033 | Leaf blade color (LBC) | TO:0000326 | fine stripe | 16 | Leaf color | Stripe |
| 26 | Leaf blade | PO:0020039 | color | PATO:0000033 | Leaf blade color (LBC) | TO:0000326 | zebra | 17 | Leaf color | Zebra |
| 27 | Leaf blade | PO:0020039 | color | PATO:0000033 | Leaf blade color (LBC) | TO:0000326 | white stripe | 18 | Leaf color | Others |
| 28 | Leaf blade | PO:0020039 | color | PATO:0000033 | Leaf blade color (LBC) | TO:0000326 | yellow stripe | 18 | Leaf color | Others |
| 29 | Leaf blade | PO:0020039 | color | PATO:0000033 | Leaf blade color (LBC) | TO:0000326 | white midrib | 18 | Leaf color | Others |

Table 6. continued...
| PHENOTYPE ID | OBSERVABLE | PO TERM ID | ATTRIBUTE | OBO TERM ID | TRAIT | TO TERM ID | TRAIT VALUE | TOS17 ID | TOS17 CLASS | TOS17 PHENOTYPE |
| --- | --- | --- | --- | --- | --- | --- | --- | --- | --- | --- |
| 30 | Leaf blade | PO:0020039 | color | PATO:0000033 | Leaf blade color (LBC) | TO:0000326 | almost white | 18 | Leaf color | Others |
| 31 | Leaf blade | PO:0020039 | color | PATO:0000033 | Leaf blade color (LBC) | TO:0000326 | light green stripe | 18 | Leaf color | Others |
| 32 | Leaf blade | PO:0020039 | color | PATO:0000033 | Leaf blade color (LBC) | TO:0000326 | white edge | 18 | Leaf color | Others |
| 33 | Leaf | PO:0009025 | length | PATO:0010077 | Leaf length | TO:0000492 | long | 104 | Seed | Others |
| 34 | Leaf | PO:0009025 | length | PATO:0010077 | Leaf length | TO:0000492 | short | 0 | -- | -- |
| 35 | Leaf | PO:0009025 | related trait | Unknown | Leaf related trait | Unknown | dried leaves | 30 | Leaf shape | Withering |
| 36 | Leaf | PO:0009025 | related trait | Unknown | Leaf related trait | Unknown | rolled | 26 | Leaf shape | Rolled leaf |
| 37 | Leaf | PO:0009025 | related trait | Unknown | Leaf related trait | Unknown | semi-rolled | 0 | -- | -- |
| 38 | Leaf | PO:0009025 | related trait | Unknown | Leaf related trait | Unknown | crinkled | 0 | -- | -- |
| 39 | Leaf | PO:0009025 | related trait | Unknown | Leaf related trait | Unknown | wilted upper half | 30 | Leaf shape | Withering |
| 40 | Leaf | PO:0009025 | senescence | Unknown | Leaf senescence (Sen) | TO:0000249 | intermediate | 0 | -- | -- |
| 41 | Leaf | PO:0009025 | width | PATO:0000084 | Leaf width | TO:0000492 | wide | 102 | Seed | Small grain |
| 42 | Leaf | PO:0009025 | width | PATO:0000084 | Leaf width | TO:0000492 | narrow | 22 | Leaf shape | Narrow leaf |
| 43 | Floret | PO:0006318 | color | PATO:0000033 | Lemma and palea color (LmPC) | TO:0000264 | brown | 104 | Seed | Others |
| 44 | Floret | PO:0006318 | color | PATO:0000033 | Lemma and palea color (LmPC) | TO:0000264 | gold | 104 | Seed | Others |
| 45 | Floret | PO:0006318 | color | PATO:0000033 | Lemma and palea color (LmPC) | TO:0000264 | brown spots on straw | 104 | Seed | Others |

Table 6. continued...
| PHENOTYPE ID | OBSERVABLE | PO TERM ID | ATTRIBUTE | OBO TERM ID | TRAIT | TO TERM ID | TRAIT VALUE | TOS17 ID | TOS17 CLASS | TOS17 PHENOTYPE |
| --- | --- | --- | --- | --- | --- | --- | --- | --- | --- | --- |
| 46 | Plant | PO:0009011 | response, mimic | Unknown | mimic response | TO:0000063 | lesion mimic-spl6 type | 51 | Mimic response | Spl/Lesion mimic |
| 47 | Plant | PO:0009011 | response, mimic | Unknown | mimic response | TO:0000063 | lesion mimic-uncharacterized | 51 | Mimic response | Spl/Lesion mimic |
| 48 | Plant | PO:0009011 | response, mimic | Unknown | mimic response | TO:0000063 | lesion mimic-spl11 type | 51 | Mimic response | Spl/Lesion mimic |
| 49 | Plant | PO:0009011 | response, mimic | Unknown | mimic response | TO:0000063 | lesion mimic-spl1 type | 51 | Mimic response | Spl/Lesion mimic |
| 50 | Germination | GRO:0007043 | rate | PATO:0007147 | Germination rate | TO:0000430 | slow | 1 | Germination | Germination rate |
| 51 | Shoot | PO:0009006 | abnormal | PATO:0000004 | Shoot growth | TO:0000077 | abnormal | 72 | Flower | Abnormal floral organ |
| 52 | Leaf | PO:0009025 | shape | PATO:0000130 | Leaf shape | TO:0000492 | -- | 29 | Leaf shape | Lamina joint |
| 53 | Floret | PO:0006318 | shape | PATO:0000130 | Abnormal floral organ | TO:0000499 | abnormal | 72 | Flower | Abnormal floral organ |
| 54 | Panicle | PO:0009049 | status | Unknown | Panicle status | Unknown | viviparous | 85 | Panicle | Vivipary |
| 55 | Panicle | PO:0009049 | axis | Unknown | Panicle axis (PnA) | TO:0000342 | straight | 0 | -- | -- |
| 56 | Panicle | PO:0009049 | exsertion | Unknown | Panicle exsertion (Exs) | TO:0000165 | partly exserted | 87 | Panicle | Neck leaf |
| 57 | Panicle | PO:0009049 | length | PATO:0010077 | Panicle length (PnL) | TO:0000040 | short | 0 | -- | -- |
| 58 | Panicle | PO:0009049 | related trait | Unknown | panicle related trait | Unknown | leafy head | 0 | -- | -- |
| 59 | Panicle | PO:0009049 | threshability | PATO:0300094 | Panicle threshability (Thr) | TO:0000406 | loose | 86 | Panicle | Shattering |
| 60 | Panicle | PO:0009049 | threshability | PATO:0300094 | Panicle threshability (Thr) | TO:0000406 | easy | 86 | Panicle | Shattering |
| 61 | Panicle | PO:0009049 | type | Unknown | Panicle type (PnT) | TO:0000089 | open | 83 | Panicle | Lax panicle |

Table 6. continued...
| PHENOTYPE ID | OBSERVABLE | PO TERM ID | ATTRIBUTE | OBO TERM ID | TRAIT | TO TERM ID | TRAIT VALUE | TOS17 ID | TOS17 CLASS | TOS17 PHENOTYPE |
| --- | --- | --- | --- | --- | --- | --- | --- | --- | --- | --- |
| 62 | Panicle | PO:0009049 | type | Unknown | Panicle type (PnT) | TO:0000089 | compact | 84 | Panicle | Dense panicle |
| 63 | Plant | PO:0009011 | height | PATO:0000074 | Plant height (Ht) | TO:0000207 | semidwarf | 41 | Height | Semidwarf |
| 64 | Plant | PO:0009011 | height | PATO:0000074 | Plant height (Ht) | TO:0000207 | dwarf | 42 | Height | Dwarf |
| 65 | Plant | PO:0009011 | height | PATO:0000074 | Plant height (Ht) | TO:0000207 | extreme dwarf | 43 | Height | Extremely dwarf |
| 66 | Plant | PO:0009011 | height | PATO:0000074 | Plant height (Ht) | TO:0000207 | tall | 44 | Height | Long culm |
| 67 | Seed | PO:0009010 | length | PATO:0010077 | Seed length | TO:0000146 | long | 104 | Seed | Others |
| 68 | Seed | PO:0009010 | length | PATO:0010077 | Seed length | TO:0000146 | short | 0 | -- | -- |
| 69 | Seed | PO:0009010 | related trait | Unknown | Seed related trait | TO:0000184 | open hull | 71 | Flower | Abnormal hull |
| 70 | Seed | PO:0009010 | related trait | Unknown | Seed related trait | TO:0000184 | beaked hull | 71 | Flower | Abnormal hull |
| 71 | Seed | PO:0009010 | related trait | Unknown | Seed related trait | TO:0000184 | half hull | 71 | Flower | Abnormal hull |
| 72 | Seed | PO:0009010 | related trait | Unknown | Seed related trait | TO:0000184 | double palea | 71 | Flower | Abnormal hull |
| 73 | Seed | PO:0009010 | related trait | Unknown | Seed related trait | TO:0000184 | multiple lemma and palea | 71 | Flower | Abnormal hull |
| 74 | Seed | PO:0009010 | related trait | Unknown | Seed related trait | TO:0000184 | depressed palea | 71 | Flower | Abnormal hull |
| 75 | Seed | PO:0009010 | related trait | Unknown | Seed related trait | TO:0000184 | triangular | 71 | Flower | Abnormal hull |
| 76 | Seed | PO:0009010 | related trait | Unknown | Seed related trait | TO:0000184 | extra glume | 71 | Flower | Abnormal hull |
| 77 | Seed | PO:0009010 | size | PATO:0000058 | Seed size | TO:0000391 | big | 101 | Seed | Large grain |
| 78 | Seed | PO:0009010 | size | PATO:0000058 | Seed size | TO:0000391 | small | 0 | -- | -- |
| 79 | Seed | PO:0009010 | width | PATO:0000084 | Seed width | TO:0000149 | wide | 102 | Seed | Small grain |

Table 6. continued.
| PHENOTYPE ID | OBSERVABLE | PO TERM ID | ATTRIBUTE | OBO TERM ID | TRAIT | TO TERM ID | TRAIT VALUE | TOS17 ID | TOS17 CLASS | TOS17 PHENOTYPE |
| --- | --- | --- | --- | --- | --- | --- | --- | --- | --- | --- |
| 80 | Seedling/vegetative stage | GRO:0007047 | viability | PATO:00000168 | Seedling/vegetative vigor (Vg) | TO:0000296 | yellow lethal | 2 | Growth | Lethal |
| 81 | Seedling/vegetative stage | GRO:0007047 | viability | PATO:00000168 | Seedling/vegetative vigor (Vg) | TO:0000296 | weak | 4 | Growth | Weak |
| 82 | Spikelet | PO:0009051 | fertility | PATO:00000171 | Spikelet fertility (SpFert) | TO:0000180 | completely sterile | 91 | Sterility | Sterile |
| 83 | Spikelet | PO:0009051 | fertility | PATO:00000171 | Spikelet fertility (SpFert) | TO:0000180 | highly sterile | 91 | Sterility | Sterile |
| 84 | Spikelet | PO:0009051 | fertility | PATO:00000171 | Spikelet fertility (SpFert) | TO:0000180 | partly sterile | 92 | Sterility | Low Fertile |
| 85 | Stem | PO:0009047 | strength | Unknown | Stem strength | TO:0000051 | brittle | 28 | Leaf shape | Brittle |
| 86 | Stem | PO:0009047 | strength | Unknown | Stem strength | TO:0000051 | stiff | 0 | -- | -- |
| 87 | Sterile lemma | PO:0009040 | length | PATO:0010077 | Sterile lemma length (SLmL) | TO:0000240 | long | 104 | Seed | Others |
| 88 | Tiller | PO:0006343 | yield | PATO:00000153 | Tillering ability (TI) | TO:0000329 | very high | 61 | Tillering | High tillering |
| 89 | Tiller | PO:0006343 | yield | PATO:00000153 | Tillering ability (TI) | TO:0000329 | low | 62 | Tillering | Low tillering |
| 90 | Tiller | PO:0006343 | yield | PATO:00000153 | Tillering ability (TI) | TO:0000329 | very low | 62 | Tillering | Low tillering |
| 91 | Internode | PO:0020142 | length | PATO:0010077 | (Uppermost) internode length | TO:0000145 | elongated | 88 | Panicle | Abnormal panicle shape |

### Discussion

In its current release, the phenotype CV is posted as a table in the IRIS-mutant website; it has mapped most of the CVs to public term ontology databases. Its purpose is not to create a new ontology but rather utilize existing ontological classifications. All future evaluations of mutants planted will be done using the phenotype CV terms to describe observed agronomic mutations. In case of novel mutant phenotypes discovered in upcoming mutant field screenings, appropriate CV entries will be developed and submitted to the appropriate ontology databases (eg. OBO, PO, and TO databases) for incorporation into existing publicly accessible resources. This resource does not propose to be the standard to which mutant phenotype characterization should be based upon but rather serve as an inter-database resource enabling clear and unambiguous queries of mutant traits across the various existing rice mutant stock resources. By using the CVs to describe observed mutant traits in your own experiments, synonymous mutants can easily be determined with high certainty from other databases, and associated mutant gene information can be mined. Specific mutant trait mapping is currently lacking in many mutant resources database available online.

The use of CVs in describing agronomic phenotypes and the ability to accurately map this description into other mutant resource databases is an important first step in gene discovery experiments. An approach in discovering genes for a particular trait of interest could be the use of mutants with either enhancement or knockout of the same or a correlated trait. A researcher could look at publicly available mutant resources available and infer the mutant gene(s) responsible for the trait observed if a particular mutant plant has the same controlled vocabulary description from a mutant database with known gene mutations. As an example, the mutant trait *loose panicle* (ID=59) as described using the IR64 phenotype CV is mapped to the TOS17 mutant trait *shattering panicle* (TOS17 ID=86). The TOS17 mutant database can potentially be queried using this corresponding TOS17-specific CV to give a list of disrupted genes associated with the trait. The accurate determination of mutant phenotypes across mutant resources can enrich information available to all mutant resources in two ways: (1) the list and annotation of mutant genes associated with one resource (e.g. TOS17 database) can be associated to another mutant resource with less available gene information (eg. IR64 mutant database) through mapping of mutant traits from one resource to another via CV, and (2) a broader list of mutant materials will be available from one resource if the same trait CV of interest is properly mapped across other mutant resources. Currently the Rice Mutant Database website hosted by the National Center of Plant Gene Research (Wuhan, China), available at http://rmd.ncpgr.cn/, shows the controlled vocabulary they implemented to describe the mutant phenotype in their collection, but no mapping to other CV in use by other mutant database resources is provided.

The IR64 mutant database is now publicly available at the website http://www.iris.irri.org/action/mutant?method=viewTerm, where information about all IR64 mutants and the associated agronomic mutant characters are compiled and described by CVs.

## II. GENE EXPRESSION PROFILING OF GR978 UNDERGOING BLAST RESISTANCE RESPONSE

### Results

The location of the recessive gene locus conferring broad resistance against *Xoo* and rice blast in GR978 was mapped using SSR markers (Madamba et al., 2009), and the physical location of the mutation was estimated *in-silico* from marker position to be at 7.4 to 11.2 Mb in chromosome 12 of the TIGR rice pseudomolecule release 3.0. Inspection of the gene models predicted in this region of the pseudomolecule revealed that the 22K oligoarray contains genes located in the vicinity of the predicted mutation. Gene expression profiling of GR978 using this oligoarray can provide information on the behavior of the genes within the region of interest, helping us to better understand the nature of the mutation that causes the gain of resistance phenotype of GR978. Before focusing on the expression pattern in this particular region, the global gene expression dataset was analyzed, starting with the statistical determination of the set of significantly differentially expressed genes per experiment, which is further partitioned into subsets of CDE and DI genes as described in the Methodology section.

#### Data quality assessment

Prior to any analysis, experimental data was inspected for quality and appropriately processed. Assessing the utility of the 22K oligoarray, the range of number of genes detected as expressed per array is from 18,014 to 20,686, with an average of 19,656 genes. A 91% success rate is expected for any gene expression experiment using the Agilent Technologies 22K rice oligochip system (Table 7)

**Table 7.** The count of expressed genes in the 16 array experiments using the Agilent Technologies 22K rice oligoarray

| ARRAY<br>NUMBER | COMPARISON | BIOREP | NUMBER OF<br>EXPRESSED<br>GENES |
| --- | --- | --- | --- |
| 1 | GR978 - 24h inoc vs water | 1 | 20,605 |
| 2 | dye swap of (1) | 1 | 20,127 |
| 3 | GR978 - 24h inoc vs water dye swap | 2 | 19,203 |
| 4 | GR978 - 24h inoc vs water | 3 | 18,982 |
| 5 | GR978 - 48h inoc vs water | 1 | 19,612 |
| 6 | dye swap of (5) | 1 | 20,301 |
| 7 | GR978 - 48h inoc vs water ds | 2 | 19,342 |
| 8 | GR978 - 48h inoc vs water | 3 | 18,904 |
| 9 | IR64 - 24h inoc vs water | 1 | 20,355 |
| 10 | dye swap of (9) | 1 | 20,103 |
| 11 | IR64 - 24h inoc vs water | 2 | 19,097 |
| 12 | IR64 - 24h inoc vs water ds | 3 | 18,014 |
| 13 | IR64 - 48h inoc vs water | 1 | 20,486 |
| 14 | dye swap of (13) | 1 | 20,686 |
| 15 | IR64 - 48h inoc vs water | 2 | 19,600 |
| 16 | IR64 - 48h inoc vs water ds | 3 | 19,081 |
| Average number of genes detected as<br>expressed |  |  | **19,656** |
| % Expressed versus total spotted (21,495) |  |  | **91%** |

Ratio-intensity (RI) plots were done to diagnose for intensity-dependent dye-specific effects in the log2 ratio values of gene expression. These systematic errors often occur for the weakly fluorescing spots, and skew the mean of all log2 ratios away from zero (the expected mean if there were no intensity-specific artifacts). RI plots from experiments with these types of systematic errors show a curve of the scatter plot away from the horizontal line where mean is zero. The RI plots (using log2 ratios, no data transformation as data points) of the twelve distinct microarray experiments (in biorep 1, the dye-swap experiments were collapsed as single straight labeled experiment using a flip-dye consistency transformation filtered at +/− 2σ) show no such errors (Figures 3 to 6).

**Figure 3.**
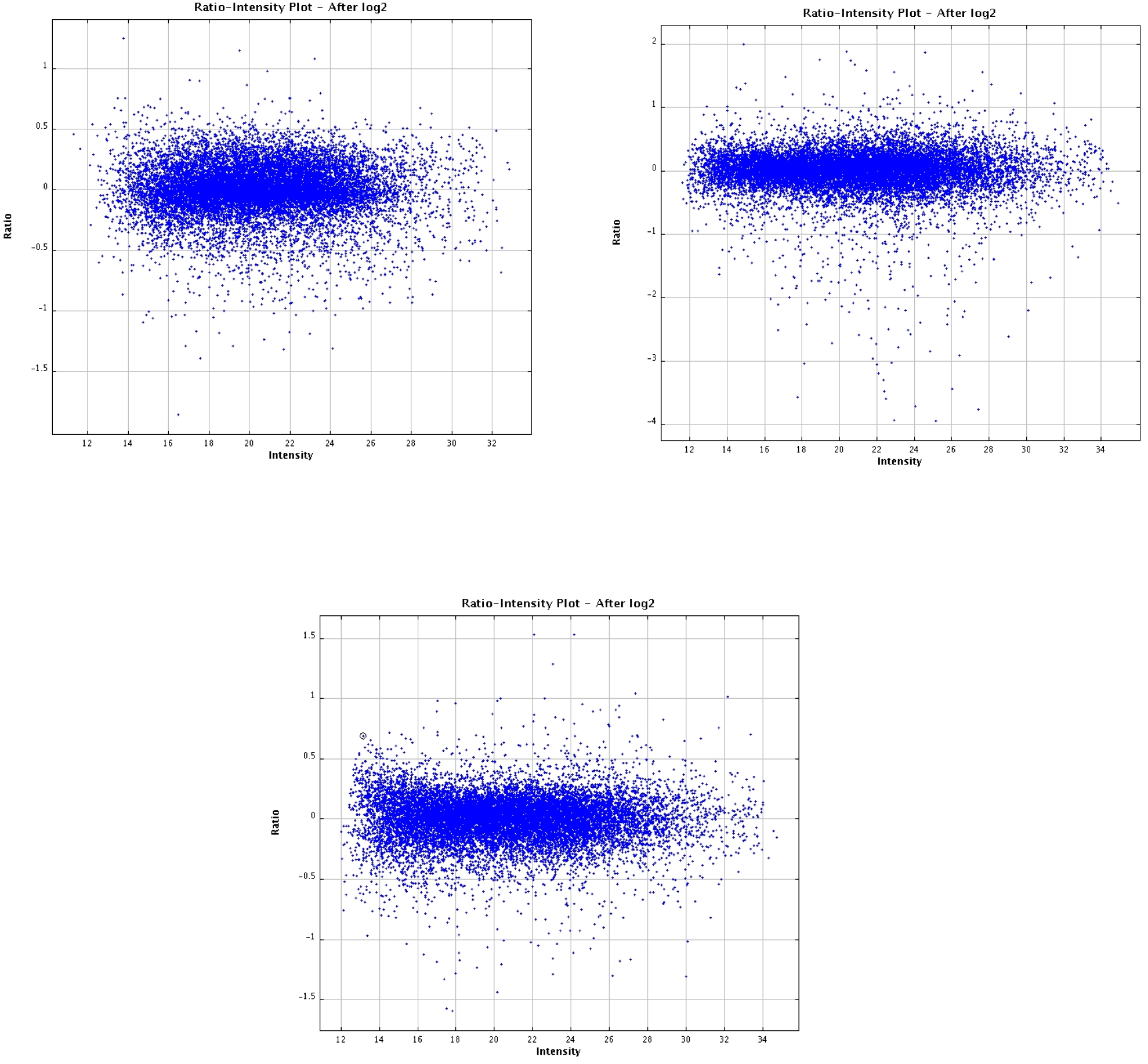
Experiments 1-3 Ratio-Intensity plot: GR978 inoculated vs water at 24h, bioreps 1 – 3 (clockwise from top left)

**Figure 4.**
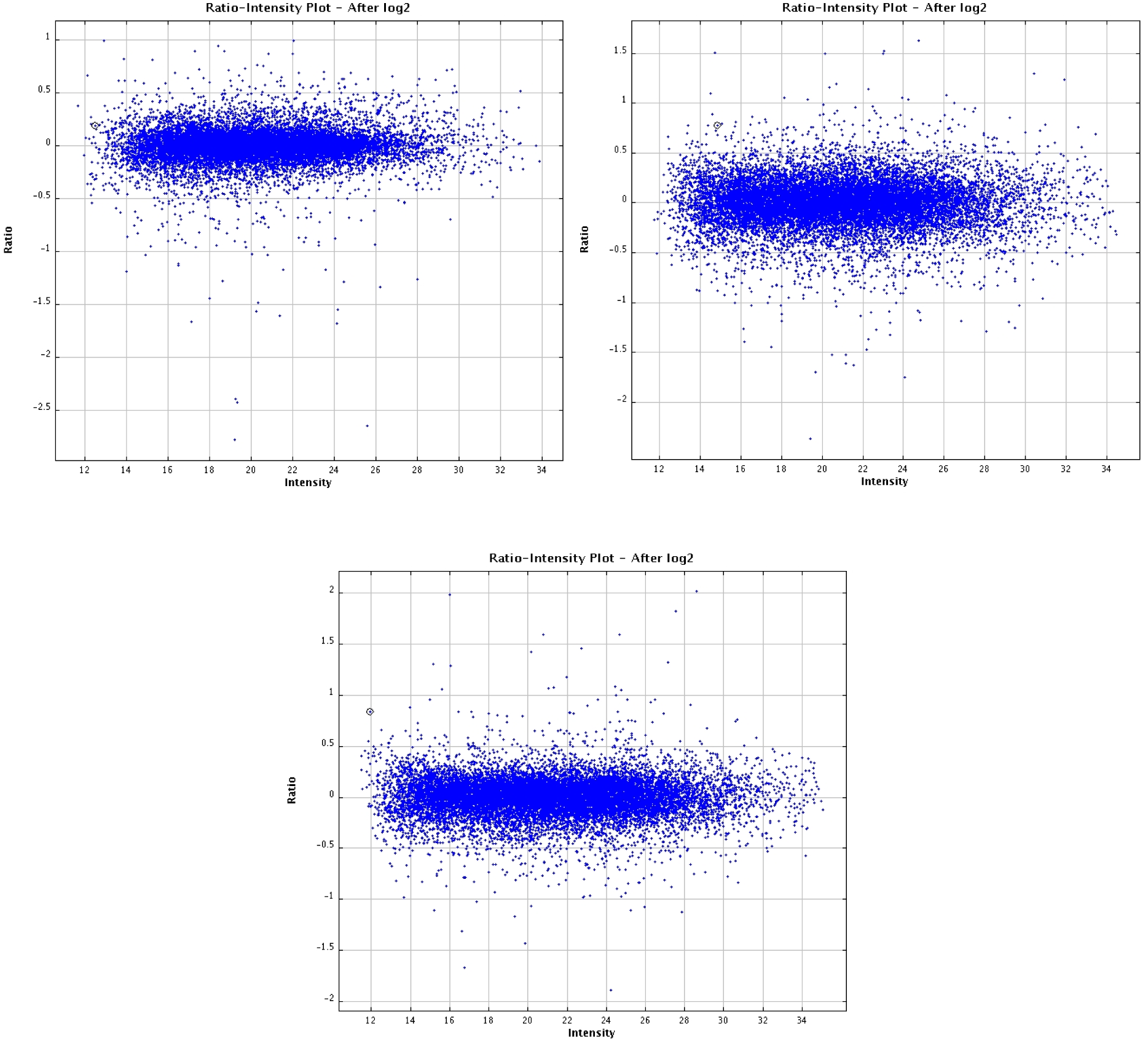
Experiment 4-6 RI plot: IR64 inoculated vs water at 24h, 3 bioreps (clockwise from top left)

**Figure 5.**
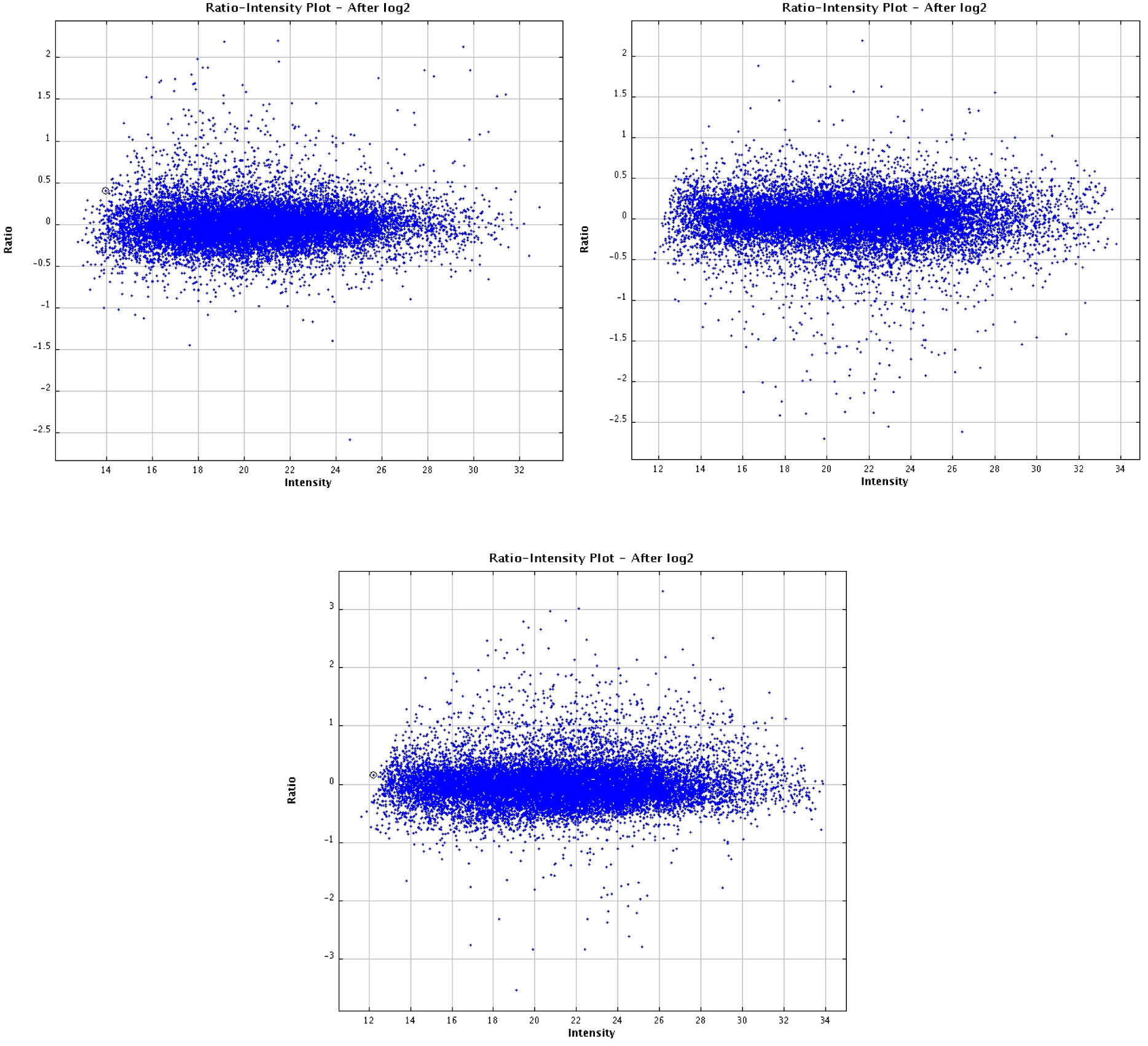
Experiments 7-9 RI plot: GR978 inoculated vs water at 48h, 3 bioreps (clockwise from top left)

**Figure 6.**
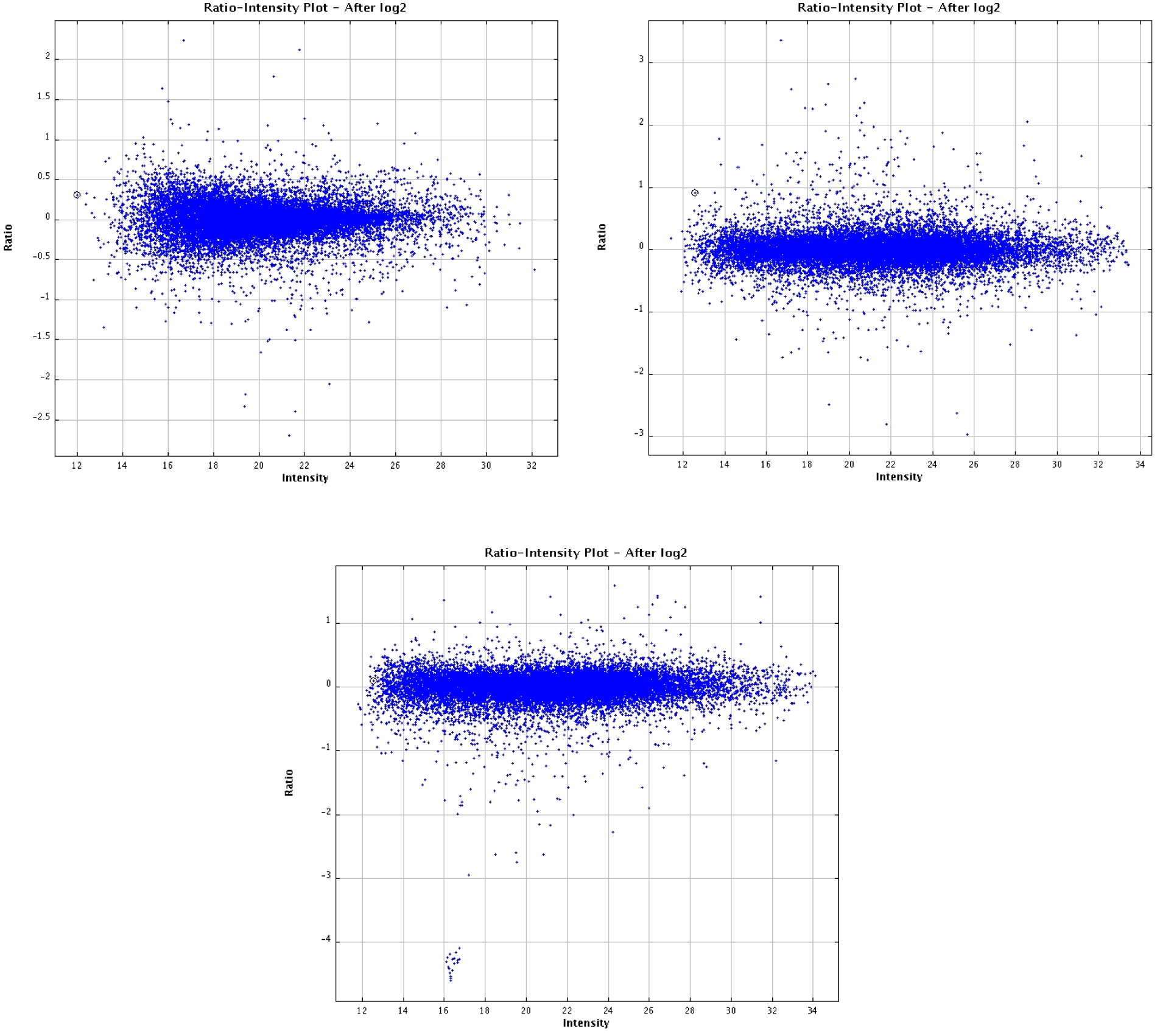
Experiment 10-12 RI plot: IR64 inoculated vs water at 48h, 3 bioreps (clockwise from top left)

This type of horizontal graph was expected since the software used to quantify expression (Agilent FE Software v 8.1) automatically performed signal processing via LOWESS. The normalization method is widely used in microarray experiments as it can remove intensity-dependent dye-specific effects in the log2 ratios (Yang et al., 2002). In terms of data quality, the twelve microarray experiments were deemed fit for expression analysis.

#### Statistical analysis for differentially expressed genes

To maximize the list of DEGs determined individually at each time point, two statistical methods were employed: (1) independent t-tests (Pan 2002), which is more sensitive in detecting significance for genes with low expression ratios, and (2) mixed model ANOVA using a split plot design implemented by MAANOVA (Wu et al., 2002), which is less sensitive and more conservative.

As expected, more genes were detected as expressed using the independent t-tests. For the CDE genes, 210 and 278 genes were detected at 24h and 48h after infection, respectively. There were 344 and 820 DI genes in the 24h and 48h time courses (either up or down regulated relative to IR64), respectively. The numbers of common and uniquely expressed genes in the two time points are shown in figures 7 and 8.

**Figure 7.**
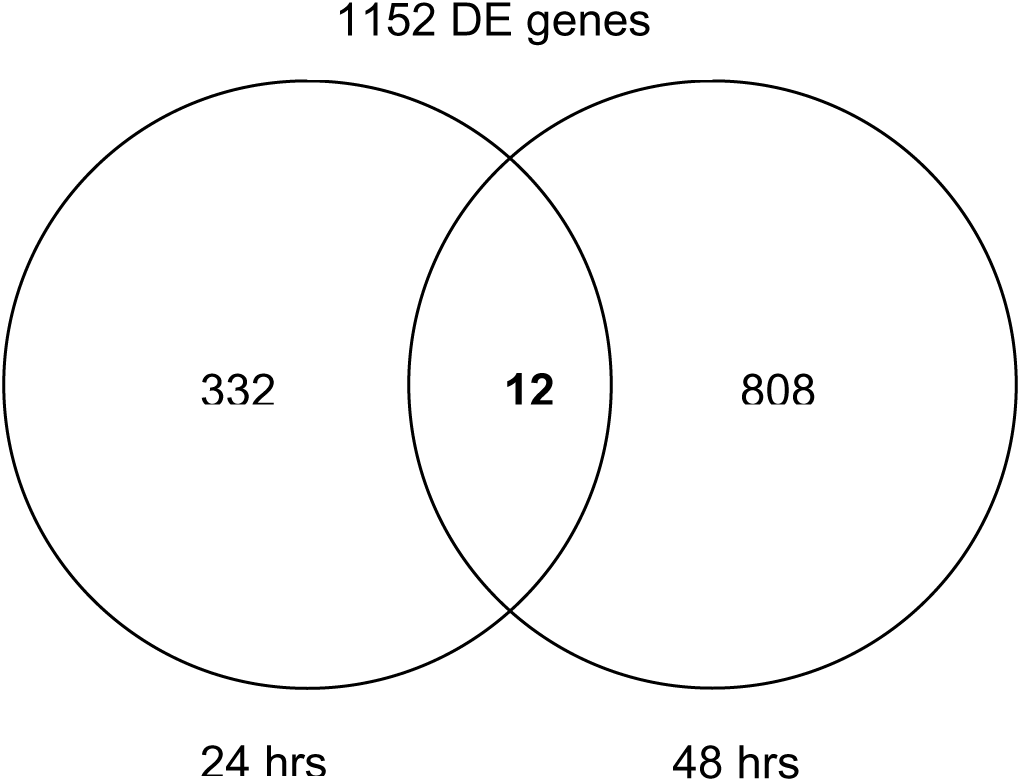
Count of DI genes in GR978 from joined independent t-tests

**Figure 8.**
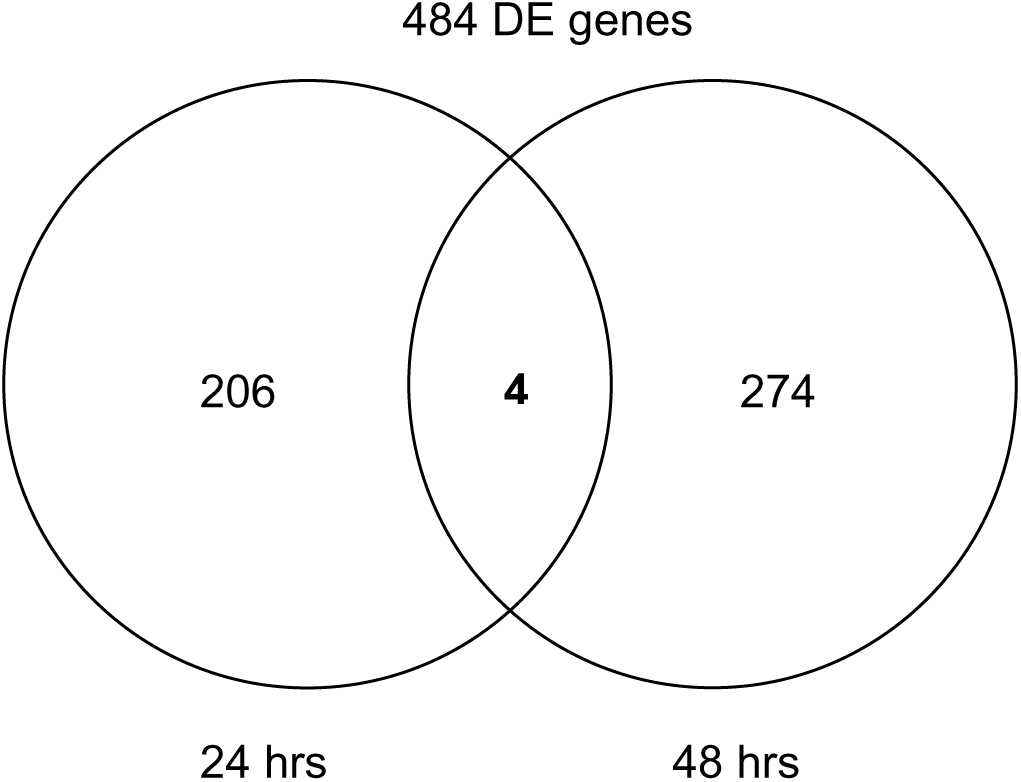
Count of CDE genes in GR978 from joined independent t-tests

#### Candidate genes conditioning broad-spectrum resistance in GR978 from the DEG list

It is suspected that the gene in GR978 that conditions moderate resistance against rice blast lies within the region defined by genetic analysis (Madamba et al. 2009). To make the search for candidate genes in this region as sensitive as possible, DEGs from GR978 during blast infection were identified using t-test. These DEGs were co-localized with the translated physical location of the mapped bacterial blight resistance gene in GR978. With this simple method, twelve interesting genes that were detected as differentially expressed in GR978 (4 CDE, 8 DI genes) were nominated as potential candidates for the observed gain of resistance phenotype (Table 8).

**Table 8.** Twelve candidate genes within the genomic region associated with resistance in GR978

| NO | KOME ID | DESCRIPTION | TU*_START | TU_END | LOG<sub>2</sub><br>RATIO<br>@24H | LOG<sub>2</sub><br>RATIO<br>@48H |
| --- | --- | --- | --- | --- | --- | --- |
| *Set 1. constitutively differentially expressed in GR978* |  |  |  |  |  |  |
| 1 | AK073343 | Oryza sativa unknown mRNA.\|PLN | 8,668,927 | 8,678,383 |  | -0.14 |
| 2 | AK102127 | Arabidopsis thaliana hypothetical protein (At2g33640/F4P9.41) mRNA, complete cds.\|PLN | 9,272,993 | 9,275,902 |  | -0.24 |
| 3 | AK058791 | Arabidopsis thaliana At1g08490/T27G7_14 mRNA, complete cds.\|PLN | 10,962,059 | 10,973,063 |  | -0.38 |
| 4 | AK061410 | Wheat mRNA for Rubisco subunit binding-protein alpha subunit.\|PLN | 10,292,469 | 10,296,815 | -0.32 |  |

\*Transcription unit, numbers are from location (in bases) along the TIGR rice pseudomolecule release 3.0

Table 8. continued.
| NO | KOME ID | DESCRIPTION | TU*_START | TU_END | LOG<sub>2</sub><br>RATIO<br>@24H | LOG<sub>2</sub><br>RATIO<br>@48H |
| --- | --- | --- | --- | --- | --- | --- |
| *Set 2. Differentially induced in GR978* |  |  |  |  |  |  |
| 5 | AK109396 | Unknown expressed protein | 7,873,181 | 7,874,146 | 0.07 |  |
| 6 | AK073908 | Arabidopsis thaliana<br>At1g80930/F23A5_23 mRNA, complete<br>cds.\|PLN | 8,800,446 | 8,806,666 | 0.25 |  |
| 7 | AK064488 | Arabidopsis thaliana selenium-binding<br>protein-like (MPO12.16) mRNA,<br>complete cds.\|PLN | 8,822,214 | 8,824,066 |  | -0.25 |
| 8 | AK099555 | Zea mays mRNA for putative glutathione<br>synthetase (gsh2 gene).\|PLN | 9,262,620 | 9,265,483 |  | -0.18 |
| 9 | AK068573 | Homo sapiens, Similar to hypothetical<br>protein FLJ11164, clone<br>IMAGE:2988291, mRNA, partial cds.\|PRI | 9,765,572 | 9,771,234 |  | -0.68 |
| 10 | AK102429 | Arabidopsis thaliana unknown protein<br>(At1g20780) mRNA, complete cds.\|PLN | 10,279,808 | 10,285,613 | -0.71 |  |
| 11 | AK068027 | Arabidopsis thaliana<br>AT5g40670/MNF13_190 mRNA,<br>complete cds.\|PLN | 10,416,568 | 10,424,741 |  | -0.45 |
| 12 | AK112053 | Unknown expressed protein | 11,192,535 | 11,203,710 | -0.31 |  |

\*Transcription unit, numbers are from location (in bases) along the TIGR rice pseudomolecule release 3.0

There were more DI genes (8) in the region than CDE genes (4). The 12 genes were differentially expressed either at 24h or 48h; none were differentially expressed at both time points. By inspection of functional annotation of the candidates, majority of the genes (9 out of 12) listed are yet of unknown function. Most of the candidates are down-expressed relative to IR64.

#### Characterizing the resistance transcriptome in GR978

Expression profile characterization of the resistance transcriptome and comparisons of profiles across other disease-resistant genotypes was done using the DEGs determined by MAANOVA, owing to the method’s more conservative approach in computing for statistical significance. The ANOVA model used was inspected for appropriateness to the microarray experiments conducted and looking at the variance components of random error terms, which are shown in Figure 9.

**Figure 9.**
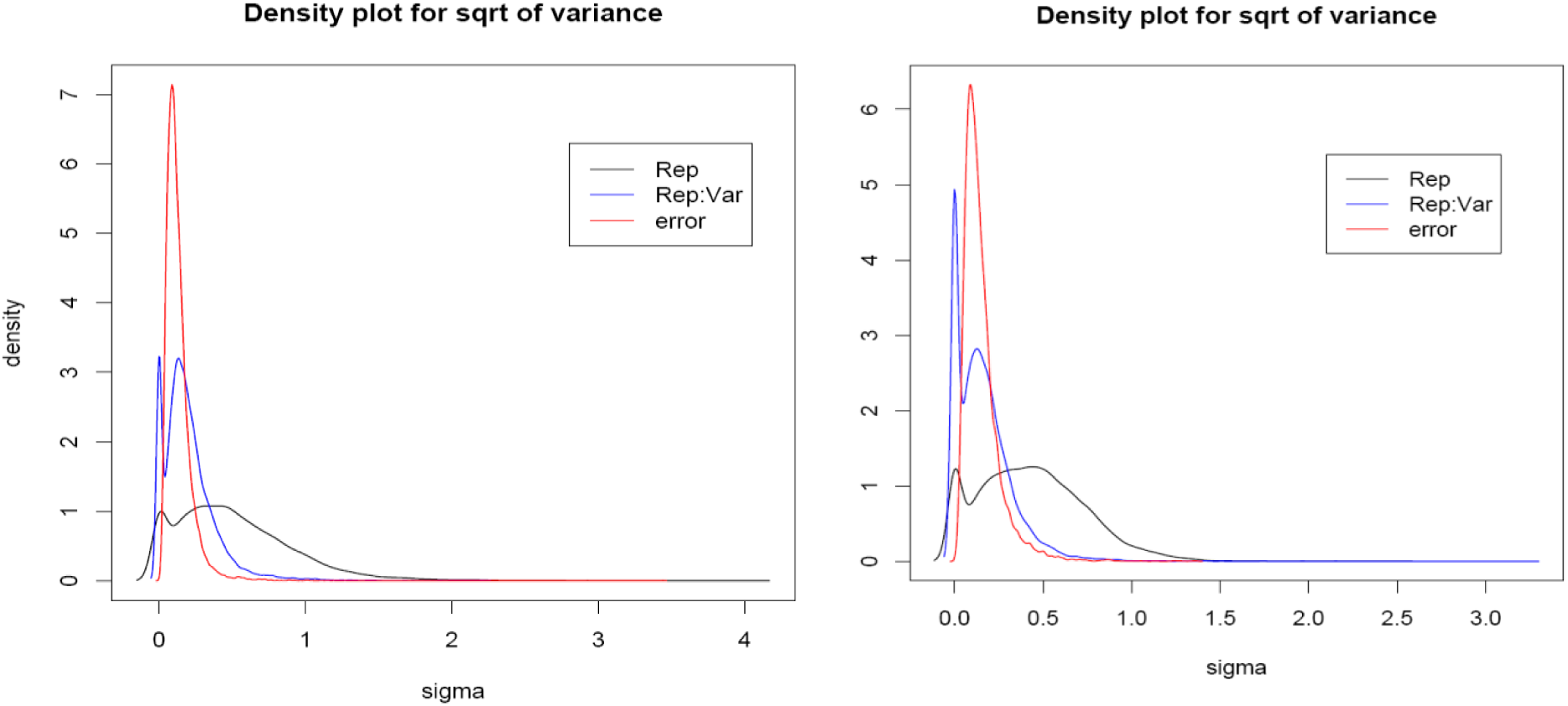
Variance components of random error for the ANOVA model used for analyzing 24h (left) and 48 h (right) time course data.

These error components estimates show that the models are able to separate and quantify the error variance components, thus are appropriate for the experimental design.

For the differentially expressed genes, there were 138 and 210 CDE genes at 24h and 48h after infection, respectively. The numbers of DI genes in GR978 (either up or down regulated relative to IR64) were 66 genes and 392 genes in the 24h and 48h time courses, respectively. For the numbers of common and uniquely expressed genes in the two time points, the following Venn diagrams (Figs. 10 and 11) summarizing these differences are shown.

**Figure 10.**
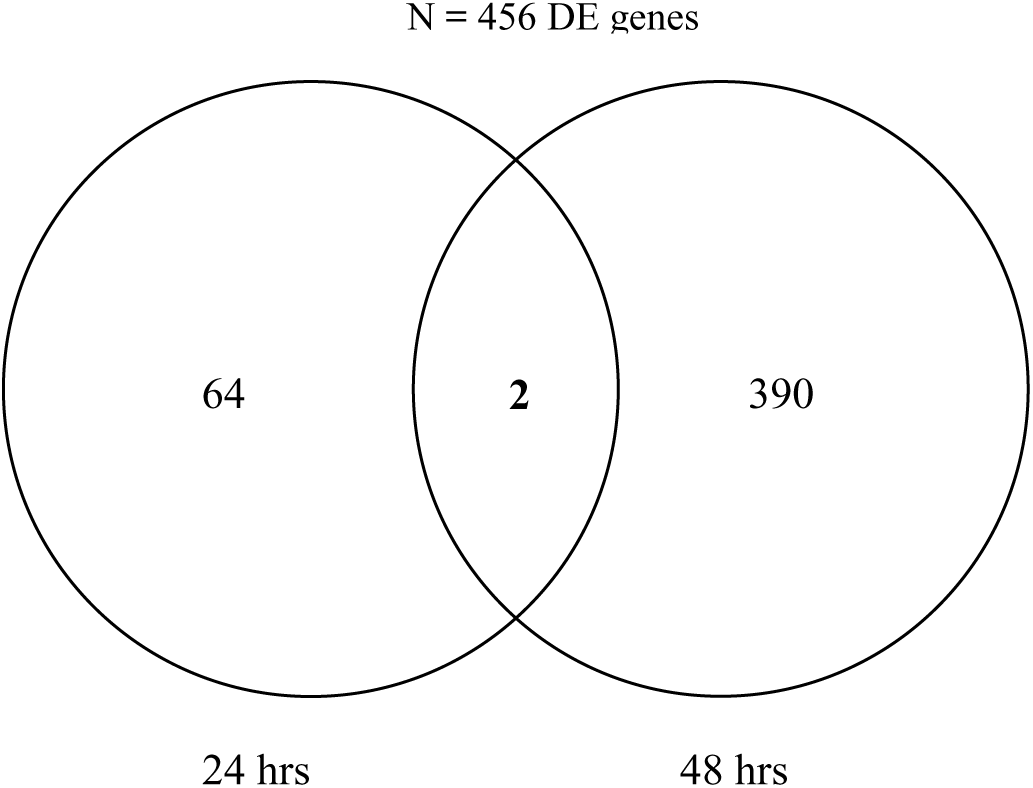
Count of DI genes in GR978 detected at 24h and 48h after infection

**Figure 11.**
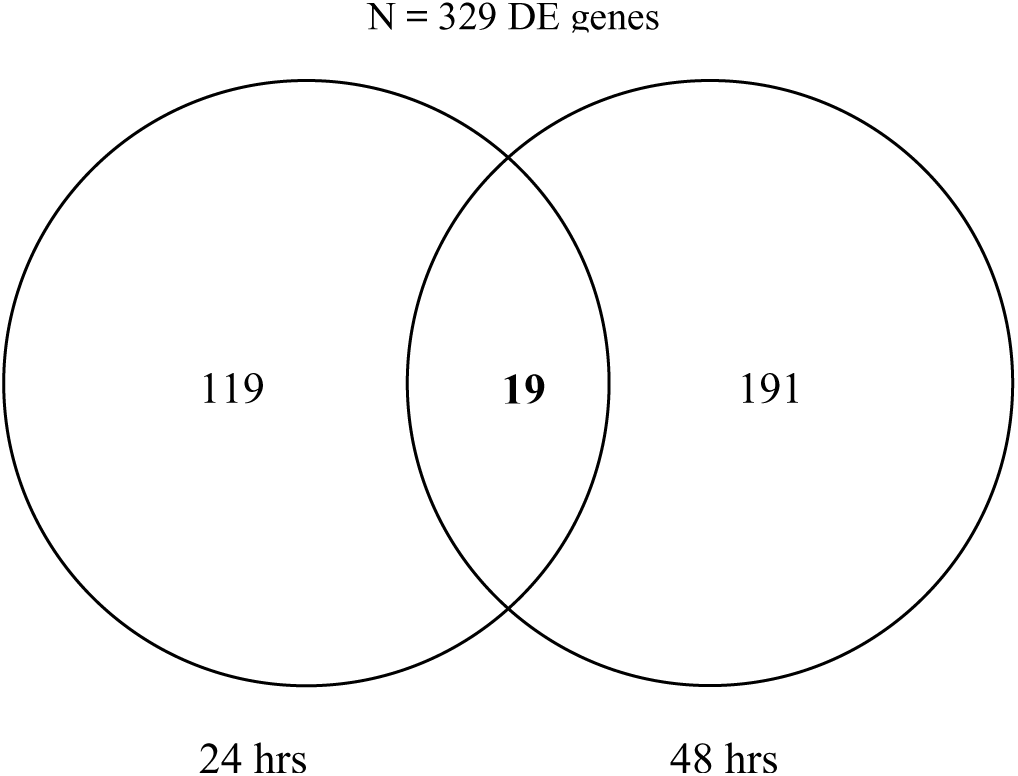
Count of CDE genes in GR978 detected at 24h and 48h after infection

Genes that were highly differentially expressed (≥ two-fold difference, where log2 ratio ≥ 1 or ≤ −1) at any of the two time points were filtered and examined for their function annotations. The short list shown in Table 9 is a union list of the genes from 24h and 48h with significant 2-fold ratio difference in at least one timepoint.

**Table 9.**
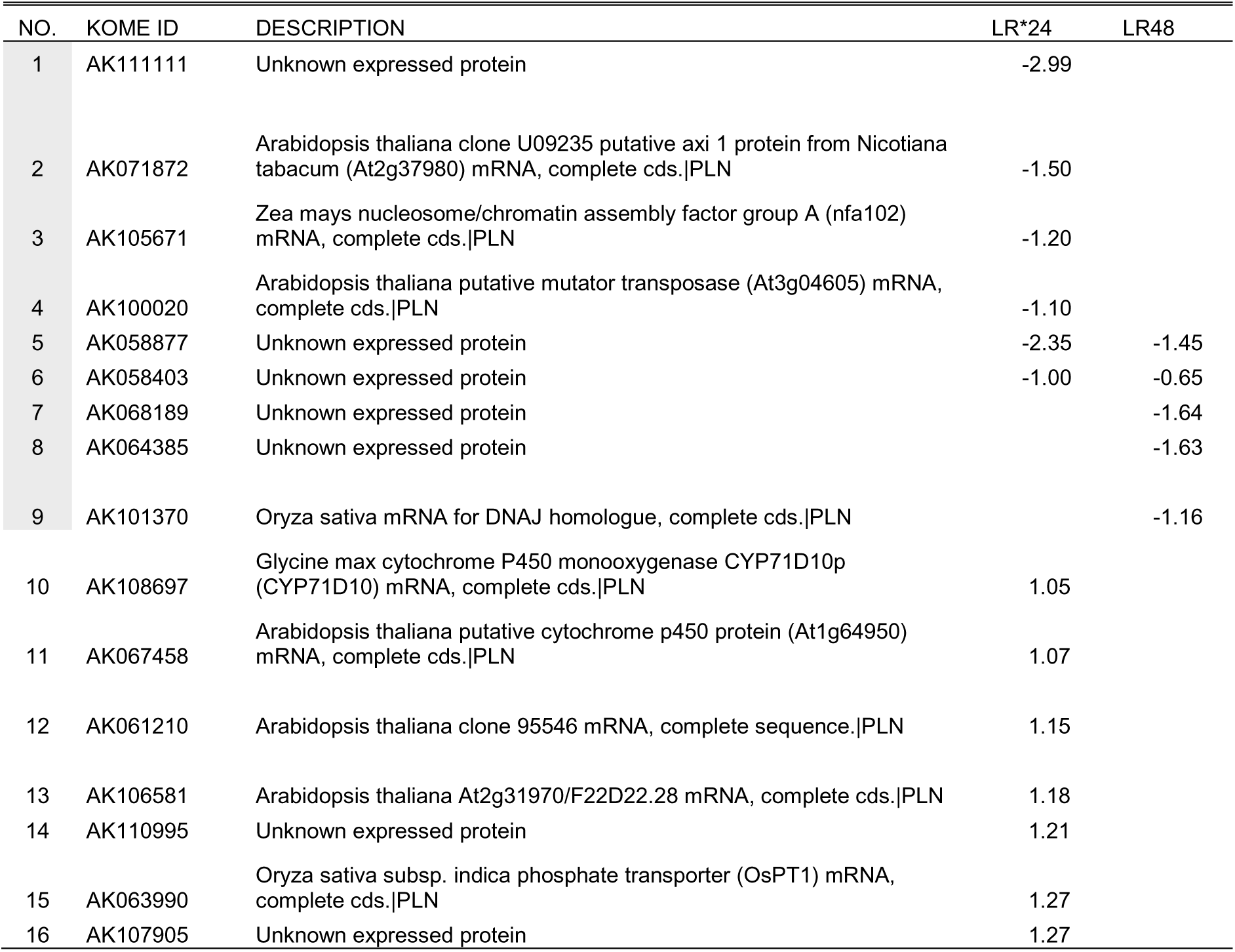
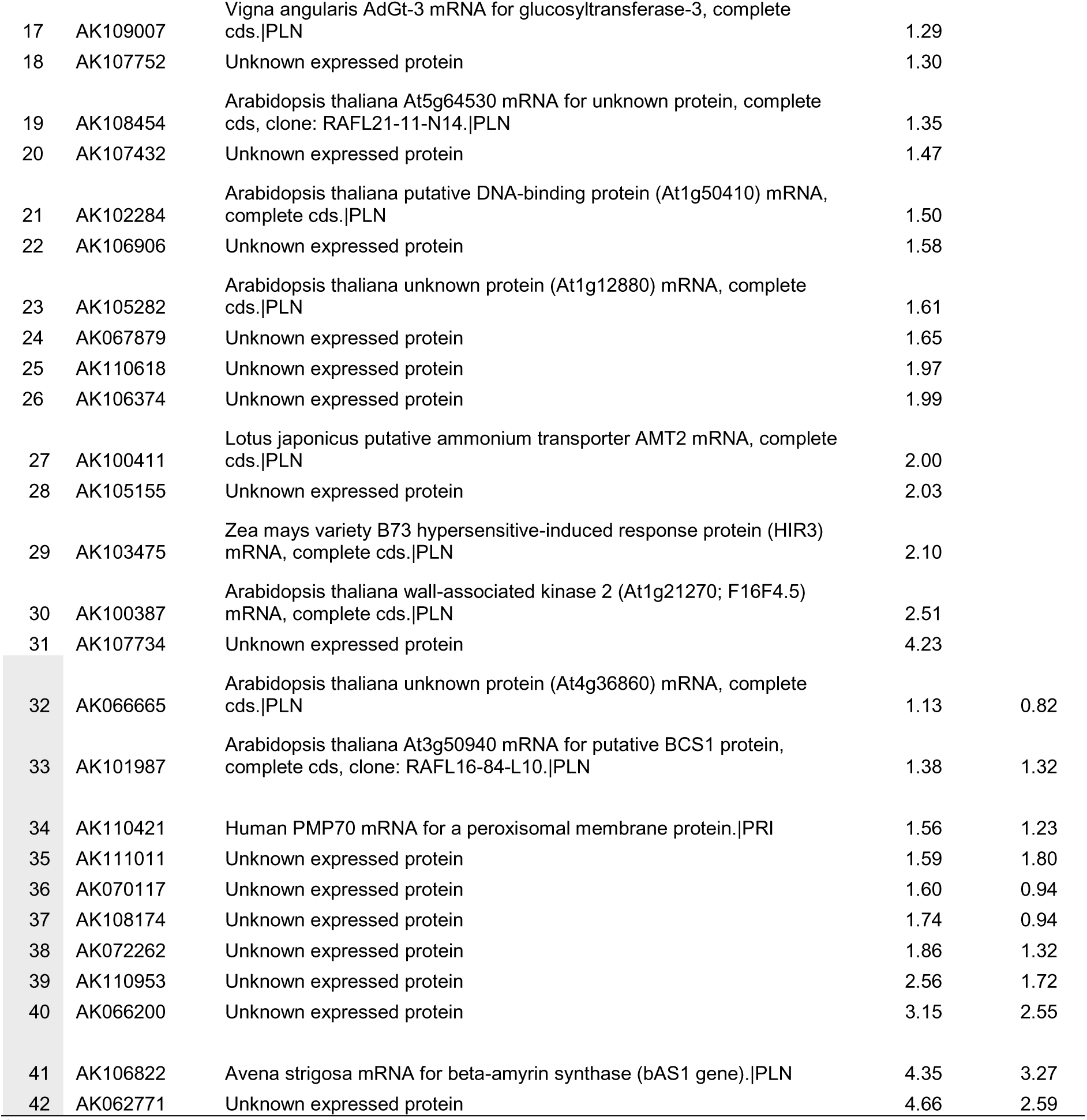
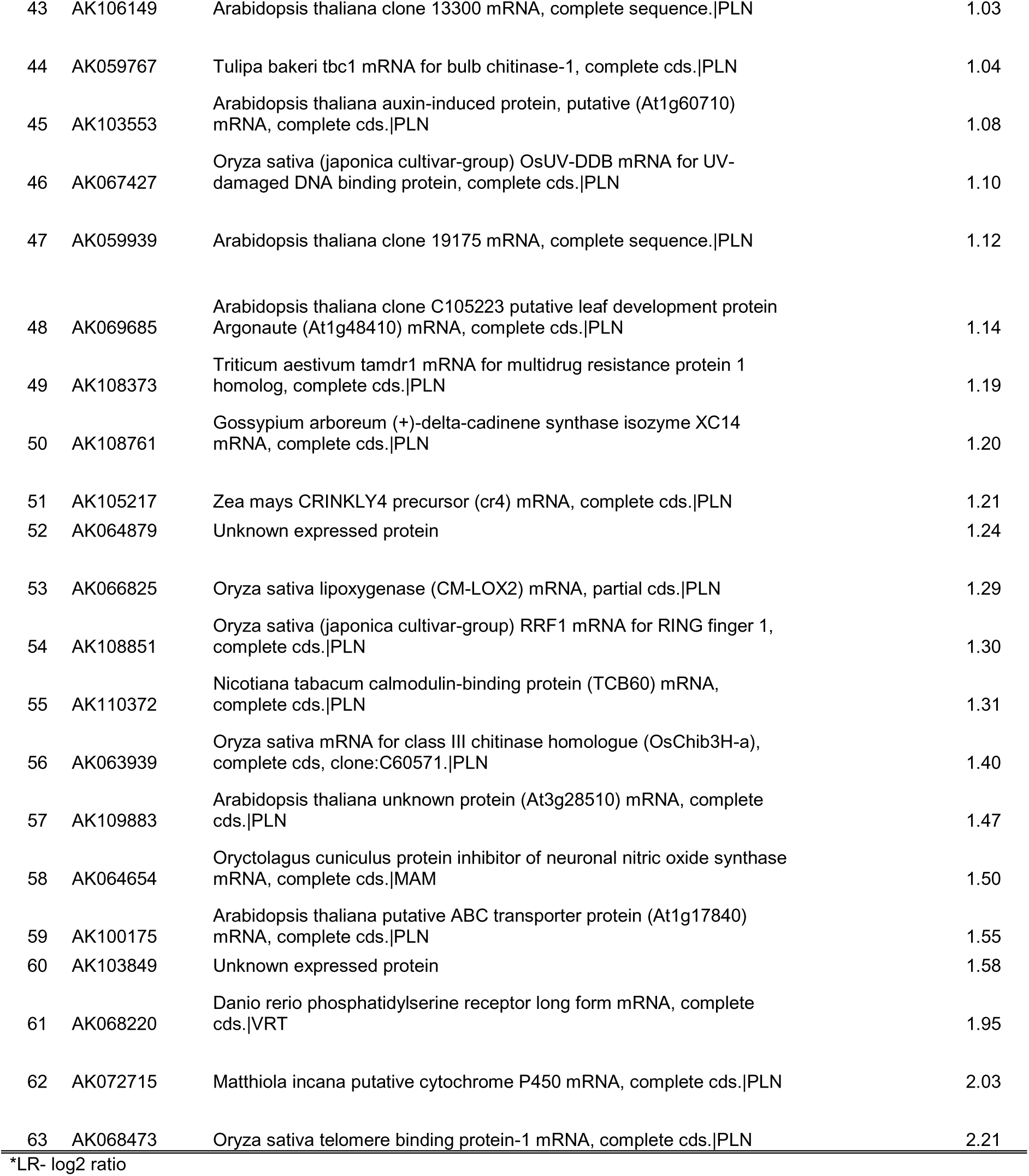
Highly expressed CDE genes (63 in all, filtered for ≥2 fold difference in at least one time point)

In this CDE gene subset, 31 genes are annotated of unknown function. The rest with function annotation represent a diverse group of genes.

The gene list was partitioned into groups in terms of significant expression within timepoints. There are 39 and 37 genes that are uniquely differentially expressed at the 24h and 48h timepoints, respectively, and 17 DEGs are in common at both timepoints. For the timepoint in which a gene is not significant in differential expression, no expression ratio value is provided. A group of 9 down-expressed genes (24h and/or 48h) can be seen; most are of unknown function and four of known function.

For the set of 22 up-expressed genes exclusive to 24h timepoint, 8 function annotations were observed, the rest are of unknown functions. Another group is composed of 11 up-expressed genes in both 24h and 48h, with 3 known function annotations. The last gene group is composed of 21 up-expressed genes exclusively in 48h. There are 16 known function annotations in this list.

For the set of 53 DI genes, almost 50% of the genes are of unknown function (22 genes), while the rest are annotated for putative function (Table 10).

**Table 10.** Fifty three highly expressed DI genes (showing a ≥2 fold difference in at least one time point)

| NO. | KOME ID | DESCRIPTION | LR*24 | LR48 |
| --- | --- | --- | --- | --- |
| 1 | AK072245 | Lolium temulentum mRNA for putative soluble acid invertase (inv1:4 gene).\|PLN | -1.18 |  |
| 2 | AK061798 | Arabidopsis thaliana clone 1368 mRNA, complete sequence.\|PLN | -1.12 |  |
| 3 | AK108477 | Pisum sativum polygalacturonase (Ppg1) mRNA, complete cds.\|PLN | -1.00 |  |
| 4 | AK105331 | Arabidopsis thaliana At5g12010/F14F18_180 mRNA, complete cds.\|PLN |  | -1.73 |
| 5 | AK066737 | Oryza sativa mRNA for lipoxygenase (rci-1 gene).\|PLN |  | -1.47 |
| 6 | AK109364 | Unknown expressed protein |  | -1.28 |
| 7 | AK064395 | Nicotiana tabacum UDP-glucose:salicylic acid glucosyltransferase (SA-GTase) mRNA, complete cds.\|PLN |  | -1.17 |
| 8 | AK065631 | Arabidopsis thaliana At2g38970/T7F6.14 mRNA, complete cds.\|PLN |  | -1.09 |
| 9 | AK106178 | C.arietinum mRNA for chitinase.\|PLN |  | -1.03 |
| 10 | AK069495 | Arabidopsis thaliana hypothetical protein (At2g23790) mRNA, complete cds.\|PLN | 1.04 |  |
| 11 | AK061399 | Medicago truncatula MTD1 mRNA, complete cds.\|PLN | 1.15 |  |
| 12 | AK071546 | Lolium rigidum clone Lol-5-v putative cytochrome P450 mRNA, complete cds.\|PLN | 1.30 |  |
| 13 | AK073487 | Rice alpha-amylase mRNA, complete cds, clone pOS137.\|PLN | 1.32 |  |
| 14 | AK063833 | Z.mays mRNA for subtilisin-chymotrypsin inhibitor.\|PLN | 1.36 |  |
| 15 | AK099157 | Rice mRNA for probenazole-inducible protein PBZ1, complete cds.\|PLN | 1.42 |  |
| 16 | AK104139 | Oryza sativa endo-1,3-beta-glucanase mRNA, complete cds.\|PLN | 1.42 |  |
| 17 | AK067892 | Arabidopsis thaliana AT4g27870/T27E11_110 mRNA, complete cds.\|PLN | 1.48 |  |
| 18 | AK069910 | Arabidopsis thaliana putative phosphatidylglycerotransferase (At4g04870) mRNA, complete cds.\|PLN | 1.70 |  |
| 19 | AK108679 | Nicotiana tabacum patatin-like protein 1 (PAT1) mRNA, complete cds.\|PLN | 1.54 | 0.51 |
| 20 | AK064892 | Oryza sativa abscisic acid- and stress-inducible protein (Asr1) mRNA, complete cds.\|PLN |  | 1.00 |
| 21 | AK058657 | Unknown expressed protein |  | 1.01 |
| 22 | AK062296 | Arabidopsis thaliana putative clathrin coat assembly protein (At4g24550) mRNA, complete cds.\|PLN |  | 1.04 |
| 23 | AK110899 | Arabidopsis thaliana SYNC1 protein mRNA, complete cds.\|PLN |  | 1.05 |
| 24 | AK058861 | Anthoceros formosae chloroplast psbM mRNA for photosystem II M-protein, complete cds.\|PLN |  | 1.11 |
| 25 | AK058289 | Oryza sativa chlorophyll a/b binding protein (kcd1895) mRNA, complete cds.\|PLN |  | 1.15 |
| 26 | AK106229 | Oryza sativa Cyt-P450 monooxygenase (PM-II) mRNA, complete cds.\|PLN |  | 1.16 |
| 27 | AK062864 | A.calamus chloroplast ndhB mRNA.\|PLN |  | 1.18 |
| 28 | AK108523 | Unknown expressed protein |  | 1.18 |
| 29 | AK107807 | Arabidopsis thaliana At2g16590/F1P15.3 mRNA sequence.\|PLN |  | 1.18 |

\*LR- log2 ratio

Table 10. continued.
| NO. | KOME ID | DESCRIPTION | LR*24 | LR48 |
| --- | --- | --- | --- | --- |
| 30 | AK072894 | Arabidopsis thaliana clone 151921 mRNA, complete sequence.\|PLN |  | 1.22 |
| 31 | AK104994 | Triticum aestivum clone CYP73-TA cytochrome P450 mRNA, complete cds.\|PLN |  | 1.26 |
| 32 | AK101002 | Arabidopsis thaliana clone 36276 mRNA, complete sequence.\|PLN |  | 1.28 |
| 33 | AK071769 | Unknown expressed protein |  | 1.30 |
| 34 | AK059110 | Oryza sativa early proembryo mRNA, complete sequence.\|PLN |  | 1.31 |
| 35 | AK108346 | Arabidopsis thaliana WRKY transcription factor 35 (WRKY35) mRNA, complete cds.\|PLN |  | 1.33 |
| 36 | AK059592 | Serenoa repens chloroplast partial mRNA for thylakoid structural protein (psbB gene).\|PLN |  | 1.36 |
| 37 | AK066307 | Marsilea quadrifolia abscisic acid-regulated heterophyly operon mRNA, partial sequence; chloroplast genes for chloroplast products.\|PLN |  | 1.41 |
| 38 | AK106319 | Oryza sativa (japonica cultivar-group) mRNA for chitinase, complete cds.\|PLN |  | 1.42 |
| 39 | AK066176 | Unknown expressed protein |  | 1.43 |
| 40 | AK109589 | N.sylvestris mitochondrial mRNA for NADH:ubiquinone oxydoreductase subunit 7.\|PLN |  | 1.46 |
| 41 | AK059143 | Mesembryanthemum crystallinum chloroplast cytochrome b-559 alpha subunit (psbE), cytochrome b-559 beta subunit (psbF), and photosystem II polypeptide (psbL) mRNA, complete cds.\|PLN |  | 1.50 |
| 42 | AK110820 | Arabidopsis thaliana AT3g57880/T10K17_90 mRNA, complete cds.\|PLN |  | 1.60 |
| 43 | AK068350 | Unknown expressed protein |  | 1.60 |
| 44 | AK064513 | N.sylvestris mitochondrial mRNA for NADH:ubiquinone oxydoreductase subunit 7.\|PLN |  | 1.63 |
| 45 | AK110458 | Drosophila melanogaster clone LD22726 BcDNA.LD22726 (BcDNA.LD22726) mRNA, complete cds.\|INV |  | 1.64 |
| 46 | AK101654 | Unknown expressed protein |  | 1.67 |
| 47 | AK062117 | O.sativa mRNA for glutaredoxin.\|PLN |  | 1.74 |
| 48 | AK108349 | Unknown expressed protein |  | 1.88 |
| 49 | AK106985 | Arabidopsis thaliana At3g41950 mRNA sequence.\|PLN |  | 2.00 |
| 50 | AK105407 | Arabidopsis thaliana At3g41950 mRNA sequence.\|PLN |  | 2.20 |
| 51 | AK058625 | Arabidopsis thaliana At2g07707 mRNA sequence.\|PLN |  | 2.21 |
| 52 | AK109648 | Oryza sativa early proembryo mRNA, complete sequence.\|PLN |  | 2.63 |
| 53 | AK060185 | Pea Chloroplast 4.5S, 5S, 16S and 23S mRNA.\|PLN |  | 2.70 |

\*LR- log2 ratio

The same timepoint partitioning was done for the 53 genes, and three groups are discernible. The first group consists of 9 down-expressed genes, 5 with known function annotation. The second group consists of 10 up-expressed genes at 24h, with 8 genes annotated for known function. The last group has 24 members, and 19 have known function annotation.

#### Gene expression clustering

Determining the relationships between individual DEGs and the groups of genes detected during the resistance response may ultimately lead to defining the resistance pathway against rice blast. Furthermore, inference of gene function for genes annotated as unknown in function can be made by determining their association of expression to genes with known function. Since a significant portion of the genes in the 22K oligoarray was annotated for known function, it is possible to characterize the function of the unknown genes in the same array by clustering genes of similar expression profiles.

For each gene set (DI genes, CDE genes), the filtered list for the highly expressed genes (≥ two-fold difference) was clustered by HCL using average linkage in order to determine gene groups with similar expression profiles across two time points. This method is applicable for a relatively short gene list and has been widely used in gene expression analysis, with a great degree of success in the transcriptional profiling of yeast using various independent time series expression data (Eisen et al. 1998).

For the CDE gene set, three obvious gene groups can be seen from HCL (Figure 12). CDE Group one is the largest, with 36 members, and it consists of moderately up-expressed genes (mean log2 ratio = 1.125) in the 24h and/or 48h time courses. There are 17 genes in this group annotated as unknown in function. Inspection of the annotation of the genes in this group show that many of functions annotated are related to stress responses related to disease resistance.

**Figure 12.**
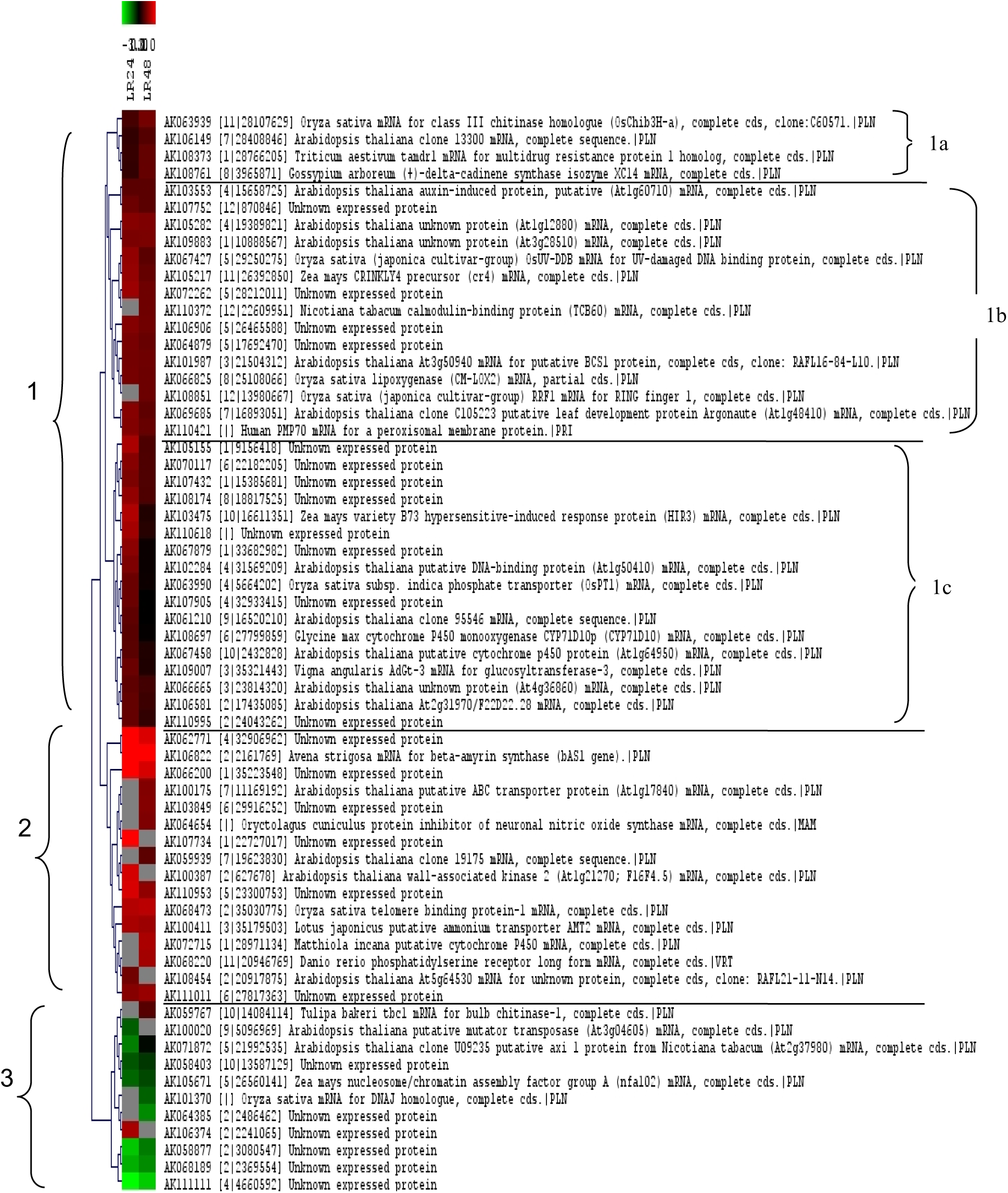
Co expressed gene clusters of highly expressed (two-fold difference) CDE genes from the 24h and 48h time courses; group assignment on the left side of the tree. Subgroups 1a, 1b, and 1c are indicated on right side of the tree

CDE Group two of the CDE gene set is a highly up-expressed cluster (mean log2 ratio = 2.356) in the 24h and/or 48h time courses with 16 members. In this group, eight genes have their function annotated as unknown.

A mostly down-expressed cluster (mean log2 ratio ≤-1.118), designated as CDE group three is also evident, with 11 gene members, 6 of which are annotated as unknown in function. Two members in CDE group 3, by annotation inspection, appear to be associated by literature to defense response.

For the DI gene set, HCL reveal three gene groups (Figure 13). In the mostly down-expressed DI group 1 (mean log2 ratio = −0.673) with 9 members, four genes were not annotated for any known function. Interestingly, all the rest of the genes with function annotation are associated by literature with defense response.

**Figure 13.**
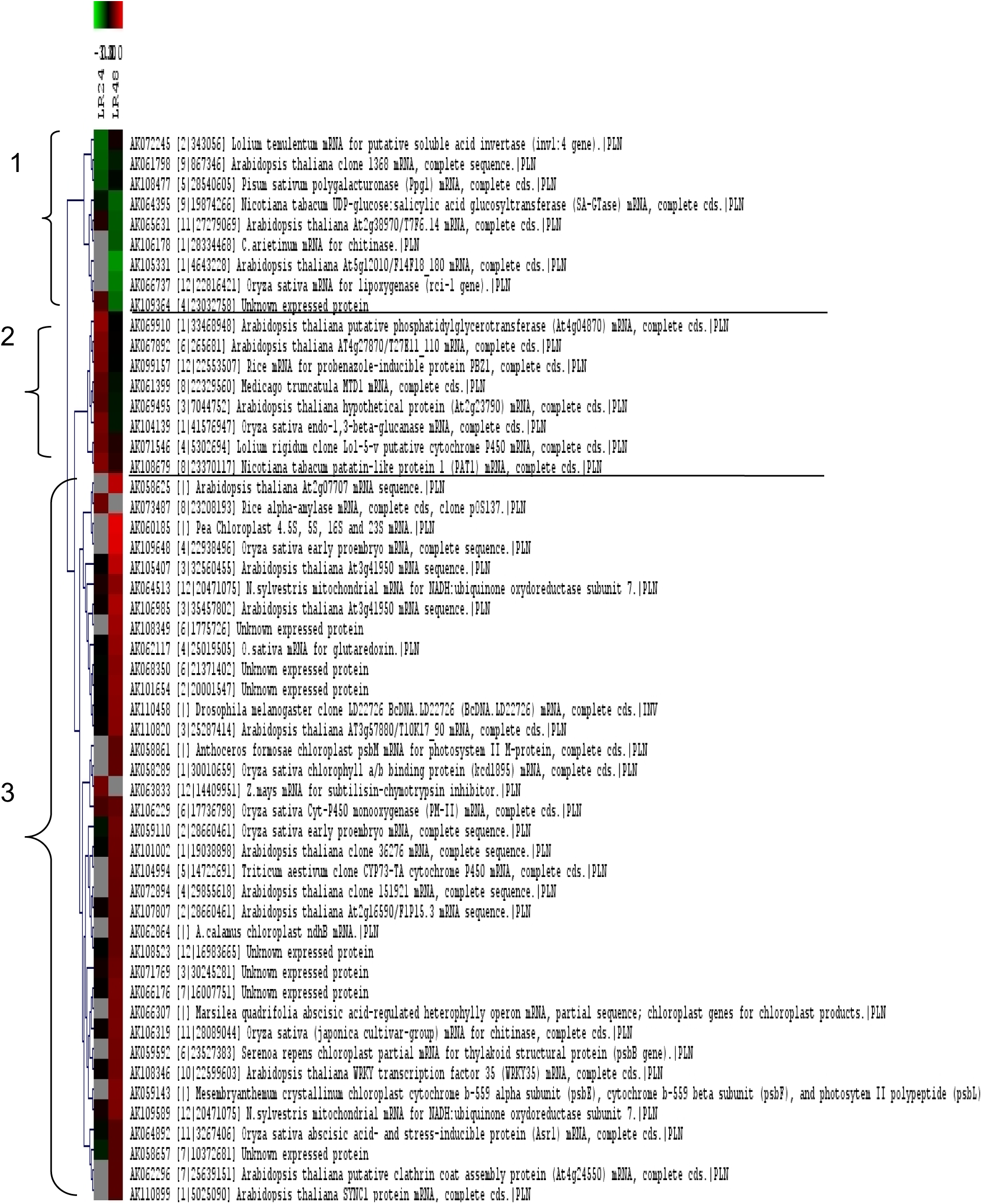
Co expressed gene clusters of highly expressed (two-fold difference) DI genes from the 24h and 48h time courses, group assignment on the left side of the tree.

DI group two, which are mostly up-expressed genes at 24h ((mean log2 ratio = 0.70), has 8 members, two of which are not annotated for known function. This group is also interesting because biotic stress response genes were found here.

DI group three is the largest with 36 gene members; most are up-expressed at 48h (mean log2 ratio = 0.98). A total of 14 members are not annotated for known function. Member genes annotated for known function include biotic stress and defense response related genes such as cytochrome p450-related genes (AK106229, AK104994), chitinase (AK106319), WRKY transcription factor (AK108346), as well as abscisic acid and stress-inducible protein (AK064892), among others. There are non-biotic stress response genes as well such as alpha-amylases and chloroplast-related genes. In all, a significant portion of the genes in all three groups is annotated as function unknown.

#### GO gene category enrichment analysis

Ontological analysis of gene expression data has proven to be useful in the biological interpretation of a list of DEGs, whether this be a list derived from significance analysis (t-test or ANOVA), or a subset of the DEG list from the gene clusters derived from HCL of the DEG list. Higher order interpretation of the list of DEGs as well as the gene clusters derived from HCL, identified during defense response, was done in order to have an insight on the underlying biological events/processes underlying disease resistance response in GR978. GO gene category enrichment analysis reveal interesting functional themes of the sets of differentially expressed genes that are implicated by literature in the defense response pathway.

For the CDE gene set, while there were few GO categories not explicitly related to biotic stress response (cell cycle processes such as microtubule binding and microtubule motor activity, chromosome segregation, cytoplasm), most of the categories listed are processes described by literature to be involved in resistance response (Table 11).

**Table 11.**
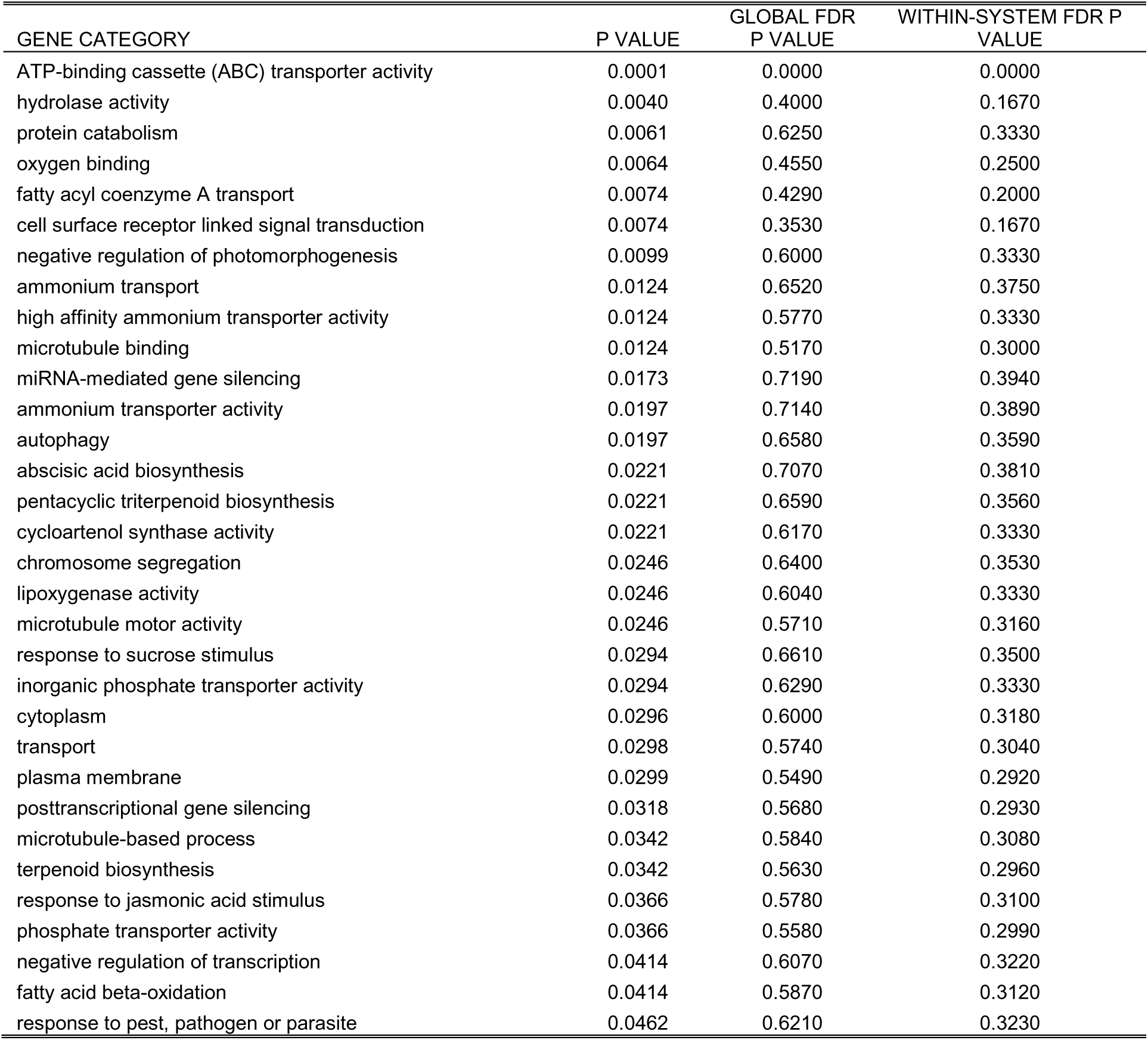
GO gene category enrichment analysis for the CDE gene set in both 24h and 48h time courses (cutoff at p < 0.05)

For the filtered DI gene set, a list of GO categories describing a more diverse range of defense response-related processes is evident. Gene categories such as response to wounding, response to jasmonic acid stimulus, response to pest, pathogen or parasite and systemic acquired resistance, among others, are explicitly related to resistance response (Table 12). Many other categories are involved in protein degradation and HR/PCD (microsome, arsenate reductase (glutaredoxin) activity, catalytic activity). A few categories, either non-defense related or indirectly related, are also listed (alpha-amylase activity, cuticle biosynthesis, thylakoid membrane).

**Table 12.** GO gene category enrichment analysis for DI gene set at 24h and 48h time course

| GENE CATEGORY | P VALUE | GLOBAL FDR P VALUE | WITHIN-SYSTEM FDR P VALUE |
| --- | --- | --- | --- |
| carbohydrate metabolism | 0.0001 | 0.0000 | 0.0000 |
| hydrolase activity, hydrolyzing O-glycosyl compounds | 0.0003 | 0.0000 | 0.0000 |
| response to wounding | 0.0032 | 0.1250 | 0.0000 |
| trans-cinnamate 4-monooxygenase activity | 0.0034 | 0.0909 | 0.0000 |
| cellular response to water deprivation | 0.0034 | 0.0714 | 0.0000 |
| CDP-diacylglycerol-glycerol-3-phosphate 3-phosphatidyltransferase activity | 0.0034 | 0.0588 | 0.0000 |
| thylakoid membrane | 0.0050 | 0.2000 | 0.0952 |
| arsenate reductase (glutaredoxin) activity | 0.0050 | 0.1740 | 0.0833 |
| photosynthesis, light harvesting in photosystem II | 0.0050 | 0.1540 | 0.0741 |
| phosphatidyltransferase activity | 0.0050 | 0.1380 | 0.0667 |
| fatty acid elongase activity | 0.0067 | 0.1880 | 0.0909 |
| cell fate determination | 0.0067 | 0.1710 | 0.0833 |
| phospholipase A2 activity | 0.0084 | 0.2630 | 0.1280 |
| lipid catabolism | 0.0134 | 0.3900 | 0.2140 |
| sucrose catabolism, using beta-fructofuranosidase | 0.0150 | 0.4550 | 0.2440 |
| abscisic acid biosynthesis | 0.0150 | 0.4260 | 0.2290 |
| lipoxygenase activity | 0.0167 | 0.4200 | 0.2350 |
| wax biosynthesis | 0.0167 | 0.3960 | 0.2220 |
| glycogen catabolism | 0.0167 | 0.3750 | 0.2110 |
| phenylpropanoid biosynthesis | 0.0167 | 0.3560 | 0.2000 |
| glucan 1,3-beta-glucosidase activity | 0.0183 | 0.3870 | 0.2220 |
| alpha-amylase activity | 0.0200 | 0.3850 | 0.2120 |
| light-harvesting complex | 0.0233 | 0.4260 | 0.2320 |
| response to jasmonic acid stimulus | 0.0249 | 0.4510 | 0.2500 |
| very-long-chain fatty acid metabolism | 0.0249 | 0.4320 | 0.2400 |
| systemic acquired resistance | 0.0265 | 0.4290 | 0.2310 |
| cellulase activity | 0.0282 | 0.4500 | 0.2470 |
| chlorophyll binding | 0.0282 | 0.4340 | 0.2380 |
| cuticle biosynthesis | 0.0298 | 0.4420 | 0.2300 |
| microsome | 0.0298 | 0.4270 | 0.2220 |
| response to pest, pathogen or parasite | 0.0314 | 0.4350 | 0.2370 |
| clathrin vesicle coat | 0.0314 | 0.4210 | 0.2290 |
| photosynthesis | 0.0412 | 0.5100 | 0.2730 |
| polygalacturonase activity | 0.0412 | 0.4950 | 0.2650 |
| catalytic activity | 0.0478 | 0.5380 | 0.2860 |

All in all, the lists of DEGs analyzed (filtered for two-fold difference) represent GO categories, majority of which are implicated by literature as defense-response related.

The list of genes in the clusters of co-expression that were defined by HCL for the DI and CDE genes was analyzed for GO category overrepresentation. In the CDE set, over-representation analysis for each of the 3 gene groups was done and results are summarized in the following tables, revealing the specific biological themes enriched for each gene group.

By inspection, most of the processes enriched in HCL CDE group 1 are resistance engagement events (Table 13).

**Table 13.** GO gene categories enriched in HCL Group 1 of the CDE gene set

| GENE CATEGORY | P VALUE | GLOBAL FDR<br>P VALUE | WITHIN-<br>SYSTEM<br>FDR P<br>VALUE |
| --- | --- | --- | --- |
| ATP-binding cassette (ABC) transporter activity | 0.0004 | 0.0000 | 0.0000 |
| hydrolase activity | 0.0008 | 0.0000 | 0.0000 |
| protein catabolism | 0.0026 | 0.2500 | 0.1110 |
| fatty acyl coenzyme A transport | 0.0048 | 0.4550 | 0.2500 |
| negative regulation of photomorphogenesis | 0.0064 | 0.4290 | 0.2670 |
| cytoplasm | 0.0066 | 0.3530 | 0.2220 |
| miRNA-mediated gene silencing | 0.0112 | 0.6500 | 0.3330 |
| abscisic acid biosynthesis | 0.0144 | 0.7390 | 0.4170 |
| chromosome segregation | 0.0160 | 0.7310 | 0.3700 |
| lipoxygenase activity | 0.0160 | 0.6550 | 0.3330 |
| zinc ion binding | 0.0189 | 0.7190 | 0.3940 |
| response to sucrose stimulus | 0.0192 | 0.6570 | 0.3610 |
| inorganic phosphate transporter activity | 0.0192 | 0.6050 | 0.3330 |
| posttranscriptional gene silencing | 0.0208 | 0.6100 | 0.3330 |
| terpenoid biosynthesis | 0.0223 | 0.6140 | 0.3330 |
| phosphate transporter activity | 0.0239 | 0.6170 | 0.3330 |
| response to jasmonic acid stimulus | 0.0239 | 0.5800 | 0.3140 |
| oxygen binding | 0.0250 | 0.5470 | 0.2960 |
| fatty acid beta-oxidation | 0.0271 | 0.5540 | 0.2980 |
| negative regulation of transcription | 0.0271 | 0.5250 | 0.2830 |
| response to pest, pathogen or parasite | 0.0302 | 0.5480 | 0.3020 |
| peroxisome | 0.0364 | 0.6150 | 0.3180 |
| transport | 0.0401 | 0.6180 | 0.3330 |
| leaf morphogenesis | 0.0411 | 0.6060 | 0.3190 |

HCL CDE group 2 (Table 14) consists of gene categories that are either not obviously involved in resistance response (microtubule-related categories) or those related directly to early resistance-engagement events (cell surface receptor-linked signal transduction, transport activities).

**Table 14.** GO gene categories enriched in HCL Group 2 of the CDE gene set

| GENE CATEGORY | P VALUE | GLOBAL FDR<br>P VALUE | WITHIN-<br>SYSTEM<br>FDR P<br>VALUE |
| --- | --- | --- | --- |
| cell surface receptor linked signal transduction | 0.0020 | 0.5000 | 0.0000 |
| plasma membrane | 0.0023 | 0.4000 | 0.1670 |
| ammonium transport | 0.0034 | 0.3750 | 0.2220 |
| high affinity ammonium transporter activity | 0.0034 | 0.2730 | 0.1670 |
| microtubule binding | 0.0034 | 0.2140 | 0.1330 |
| autophagy | 0.0054 | 0.2940 | 0.1670 |
| ammonium transporter activity | 0.0054 | 0.2500 | 0.1430 |
| pentacyclic triterpenoid biosynthesis | 0.0060 | 0.2610 | 0.1670 |
| cycloartenol synthase activity | 0.0060 | 0.2310 | 0.1480 |
| microtubule motor activity | 0.0067 | 0.2410 | 0.1330 |
| microtubule-based process | 0.0094 | 0.3130 | 0.1820 |
| microtubule associated complex | 0.0298 | 0.6570 | 0.3330 |
| polyamine transport | 0.0383 | 0.7110 | 0.3590 |
| nodulation | 0.0383 | 0.6590 | 0.3330 |
| cobalt ion transport | 0.0396 | 0.6140 | 0.3110 |
| organelle inner membrane | 0.0396 | 0.5740 | 0.2920 |
| nitrate transport | 0.0409 | 0.5600 | 0.2750 |
| cytochrome complex assembly | 0.0428 | 0.5470 | 0.2780 |

CDE group 3 (Table 15) represents categories not explicitly related to resistance response.

**Table 15.** GO gene categories enriched in HCL Group 3 of the CDE gene set

| GENE CATEGORY | P VALUE | GLOBAL FDR | WITHIN- |
| --- | --- | --- | --- |
|  |  | P VALUE | SYSTEM FDR |
| nucleosome assembly | 0.0082 | 4.0000 | 1.6700 |
| chaperone activity | 0.0287 | 3.8000 | 1.6700 |
| protein folding | 0.0358 | 2.6300 | 1.2200 |

The same analysis was made for the DI gene set, for each group determined by HCL. Majority of the categories enriched in HCL DI group 1 is early to late resistance-response related (Table 16).

**Table 16.** GO gene categories enriched in HCL Group 1 of the DI gene set

| GENE CATEGORY | P VALUE | GLOBAL FDR P | WITHIN-SYSTEM |
| --- | --- | --- | --- |
|  |  | VALUE | FDR P VALUE |
| carbohydrate metabolism | 0.0003 | 0.0000 | 0.0000 |
| phospholipase A2 activity | 0.0027 | 0.4000 | 0.1670 |
| lipid catabolism | 0.0043 | 0.5000 | 0.2220 |
| abscisic acid biosynthesis | 0.0048 | 0.4550 | 0.2500 |
| sucrose catabolism, using beta-fructofuranosidase | 0.0048 | 0.3570 | 0.2000 |
| hydrolase activity, hydrolyzing O-glycosyl compounds | 0.0050 | 0.2940 | 0.1670 |
| lipoxygenase activity | 0.0054 | 0.2500 | 0.1430 |
| response to jasmonic acid stimulus | 0.0080 | 0.3040 | 0.1670 |
| response to pest, pathogen or parasite | 0.0102 | 0.3460 | 0.1850 |
| polygalacturonase activity | 0.0134 | 0.3450 | 0.2000 |
| jasmonic acid biosynthesis | 0.0171 | 0.3750 | 0.1820 |
| response to water deprivation | 0.0171 | 0.3430 | 0.1670 |
| transferase activity, transferring hexosyl groups | 0.0239 | 0.3950 | 0.2050 |
| response to wounding | 0.0265 | 0.3900 | 0.1900 |
| ubiquitin-protein ligase activity | 0.0323 | 0.4090 | 0.2000 |
| metabolism | 0.0330 | 0.3830 | 0.1880 |
| UDP-glycosyltransferase activity | 0.0499 | 0.4600 | 0.2160 |

GO gene category enrichment analysis result for Group 2 of the DI set is shown in Table 17.

**Table 17.** GO gene categories enriched in HCL Group 2 of the DI gene set

| GENE CATEGORY | P VALUE | GLOBAL FDR P VALUE | WITHIN-SYSTEM FDR P VALUE |
| --- | --- | --- | --- |
| CDP-diacylglycerol-glycerol-3-phosphate 3-phosphatidyltransferase activity | 0.0008 | 0.0000 | 0.0000 |
| phosphatidyltransferase activity | 0.0012 | 0.0000 | 0.0000 |
| glucan 1,3-beta-glucosidase activity | 0.0044 | 0.5000 | 0.2220 |
| systemic acquired resistance | 0.0064 | 0.5450 | 0.2500 |
| cellulase activity | 0.0068 | 0.4290 | 0.2000 |
| phospholipid biosynthesis | 0.0152 | 0.6470 | 0.3330 |
| nutrient reservoir activity | 0.0224 | 0.7000 | 0.3330 |

Again, majority of the enriched categories describe defense-related responses.

GO gene category enrichment analysis of DI group 3 is shown in Table 18, with most of the enriched categories representing defense response processes. Many enriched processes are not explicitly defense-related, however.

**Table 18.** GO gene categories enriched in HCL Group 3 of the DI gene set

| GENE CATEGORY | P VALUE | GLOBAL FDR P VALUE | WITHIN-SYSTEM FDR P VALUE |
| --- | --- | --- | --- |
| trans-cinnamate 4-monooxygenase activity | 0.0015 | 0.0000 | 0.0000 |
| cellular response to water deprivation | 0.0015 | 0.0000 | 0.0000 |
| photosynthesis, light harvesting in photosystem II | 0.0022 | 0.0000 | 0.0000 |
| thylakoid membrane | 0.0022 | 0.0000 | 0.0000 |
| arsenate reductase (glutaredoxin) activity | 0.0022 | 0.0000 | 0.0000 |
| cell fate determination | 0.0030 | 0.1180 | 0.0556 |
| fatty acid elongase activity | 0.0030 | 0.1000 | 0.0476 |
| phenylpropanoid biosynthesis | 0.0074 | 0.3910 | 0.2080 |
| wax biosynthesis | 0.0074 | 0.3460 | 0.1850 |
| glycogen catabolism | 0.0074 | 0.3100 | 0.1670 |
| alpha-amylase activity | 0.0088 | 0.3130 | 0.1820 |
| light-harvesting complex | 0.0103 | 0.3710 | 0.1940 |
| very-long-chain fatty acid metabolism | 0.0110 | 0.3420 | 0.1790 |
| chlorophyll binding | 0.0125 | 0.3660 | 0.2140 |
| microsome | 0.0132 | 0.3640 | 0.2000 |
| cuticle biosynthesis | 0.0132 | 0.3400 | 0.1880 |

Table 18. continued.
| GENE CATEGORY | P VALUE | GLOBAL FDR P<br>VALUE | WITHIN-SYSTEM<br>FDR P VALUE |
| --- | --- | --- | --- |
| clathrin vesicle coat | 0.0139 | 0.3400 | 0.1760 |
| carbohydrate metabolism | 0.0171 | 0.3960 | 0.2040 |
| photosynthesis | 0.0183 | 0.3930 | 0.2110 |
| thiol-disulfide exchange intermediate activity | 0.0284 | 0.5760 | 0.3000 |
| toxin catabolism | 0.0313 | 0.5810 | 0.3020 |
| glutathione transferase activity | 0.0327 | 0.5690 | 0.3030 |
| response to wounding | 0.0363 | 0.5880 | 0.3040 |
| acyltransferase activity | 0.0370 | 0.5770 | 0.3060 |
| response to light stimulus | 0.0378 | 0.5680 | 0.2930 |

#### Inferring the transcription regulatory network of the resistance transcriptome

Computational methods for clustering co-regulated genes and inferring the mechanisms of global gene regulation using gene expression values, though well established, often ignore the underlying biological mechanisms governing the regulation of gene expression. Gene co-regulation is effected by *cis*-acting regulatory elements (motifs) in the upstream region shared among the co regulated genes and the transcription factors binding these motifs (Dolinski et al., 2005), and an approach to co-regulation clustering is the identification of motifs in the set of genes of interest, identifying shared motifs, and determining if these shared motifs are enriched in the gene set. Genes are then grouped according to shared *cis* elements or motifs enriched in the gene set under study.

The 1kb upstream sequences of all the KOME genes used in the 22K oligoarray were scanned for the occurrence of the 462 motif sequences deposited in the PLACE release 24database (Higo et al., 1999), and the computed distribution of each motif in the whole genome becomes the expected distribution of each motif. Motif enrichment analysis is a test for fixed ratio using Chi square test of the motifs within the subset under study against the global ratio (see *Methodology* section for details).

Owing to the inducible nature of the DI genes, motif enrichment analysis focused on the 53 DI gene set (with log2ratio ≥ 1.0 in either 24h or 48h timepoint) for motif enrichment analysis, using the cutoff criteria of p <.01 and motif ratio ≥ 1.5 in at least one timepoint. Forty motifs were detected as significantly enriched in the DI genes for the 24h and 48h time courses. By casual inspection of motif annotation, interesting motifs involved in regulating stress and defense response genes were observed, such as ABA responsive motifs (Table 19).

**Table 19.**
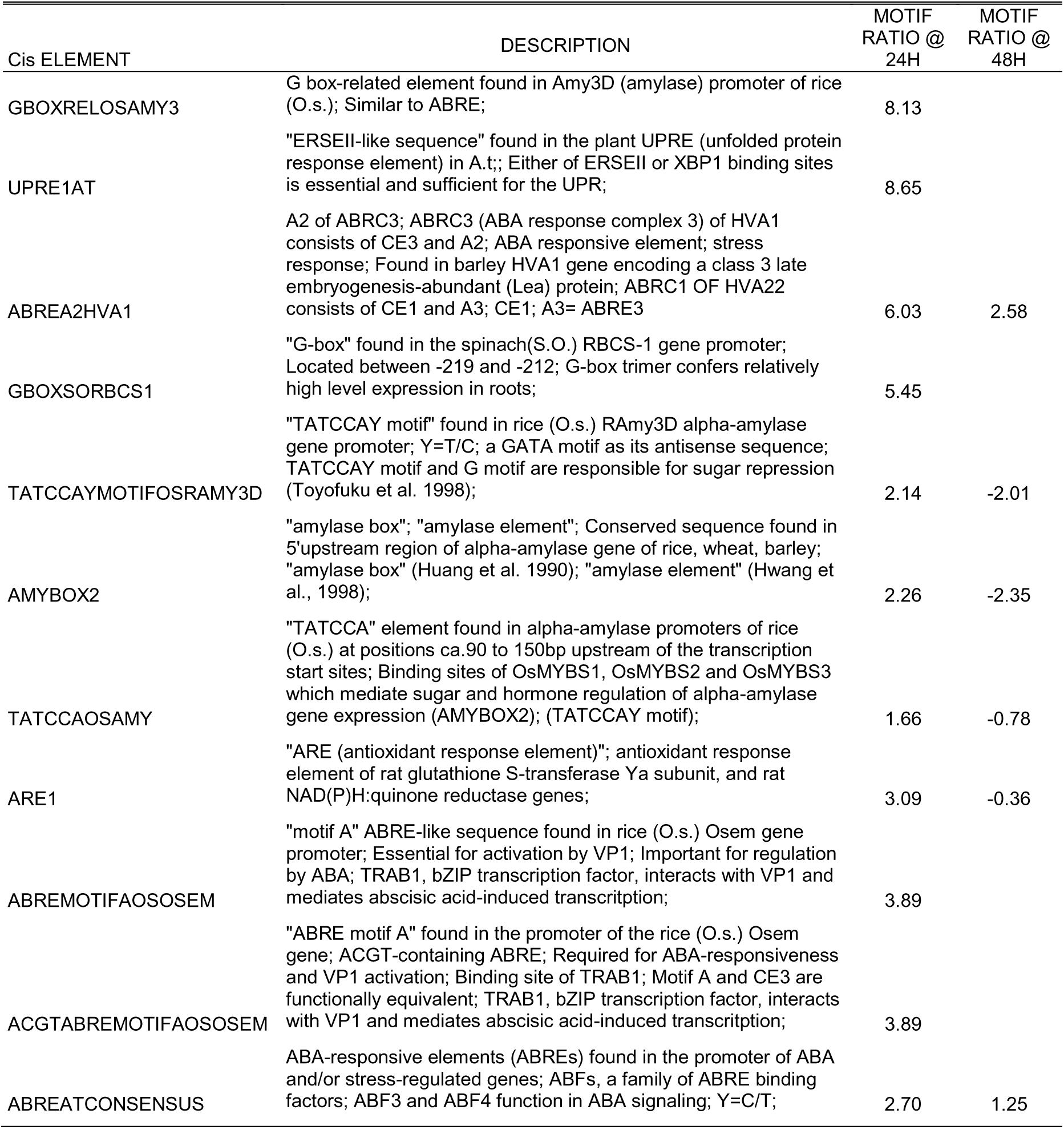
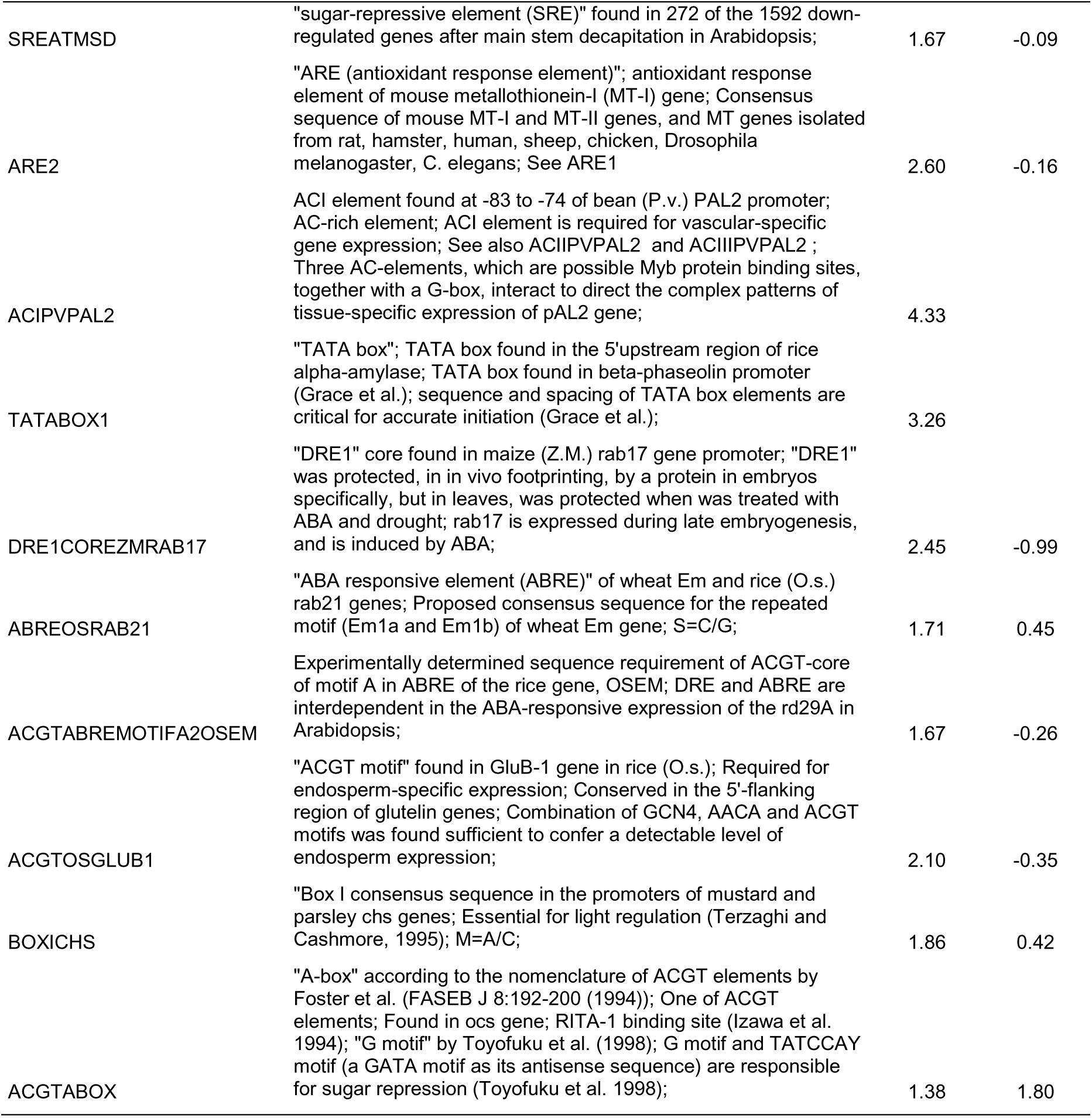
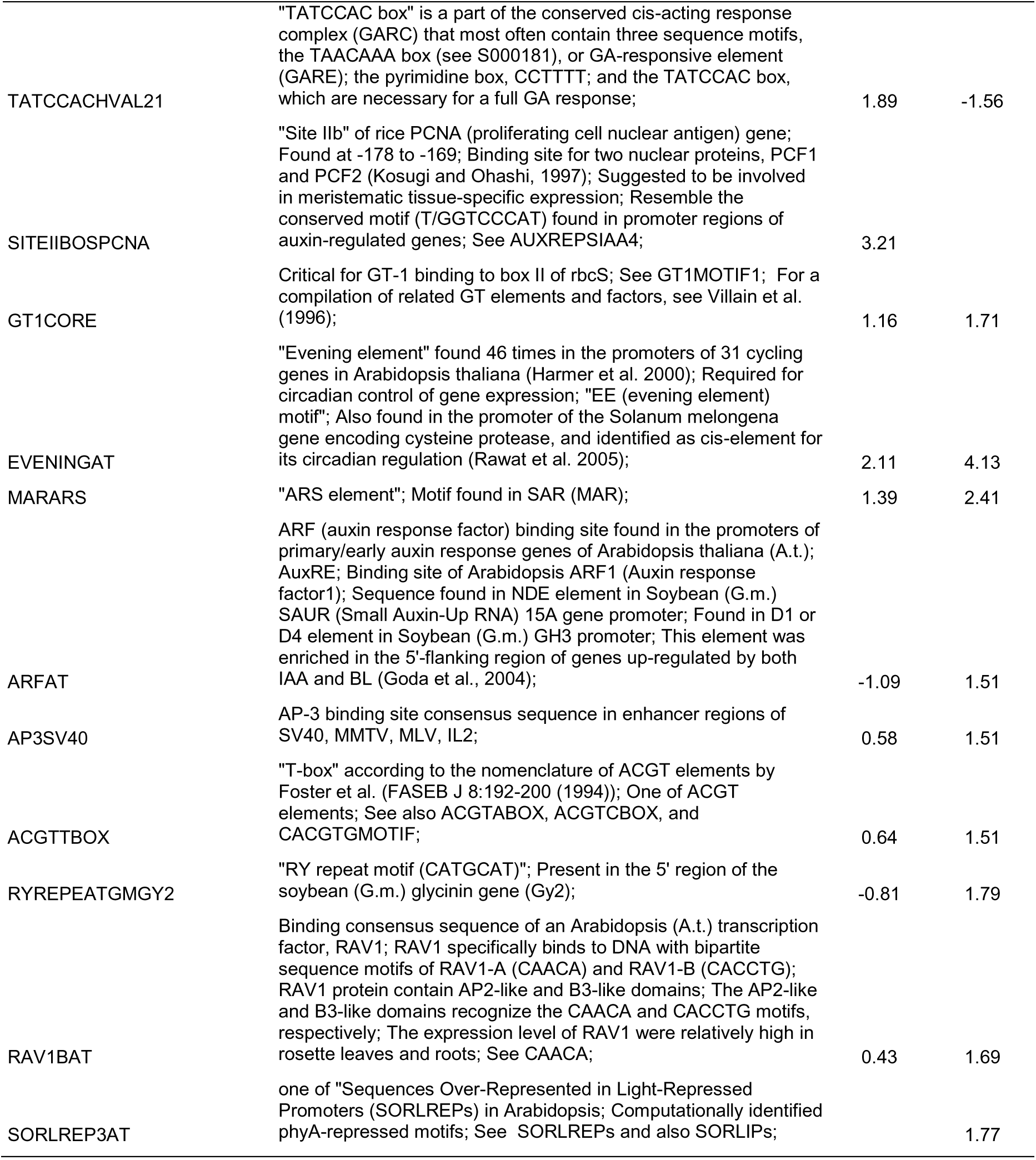
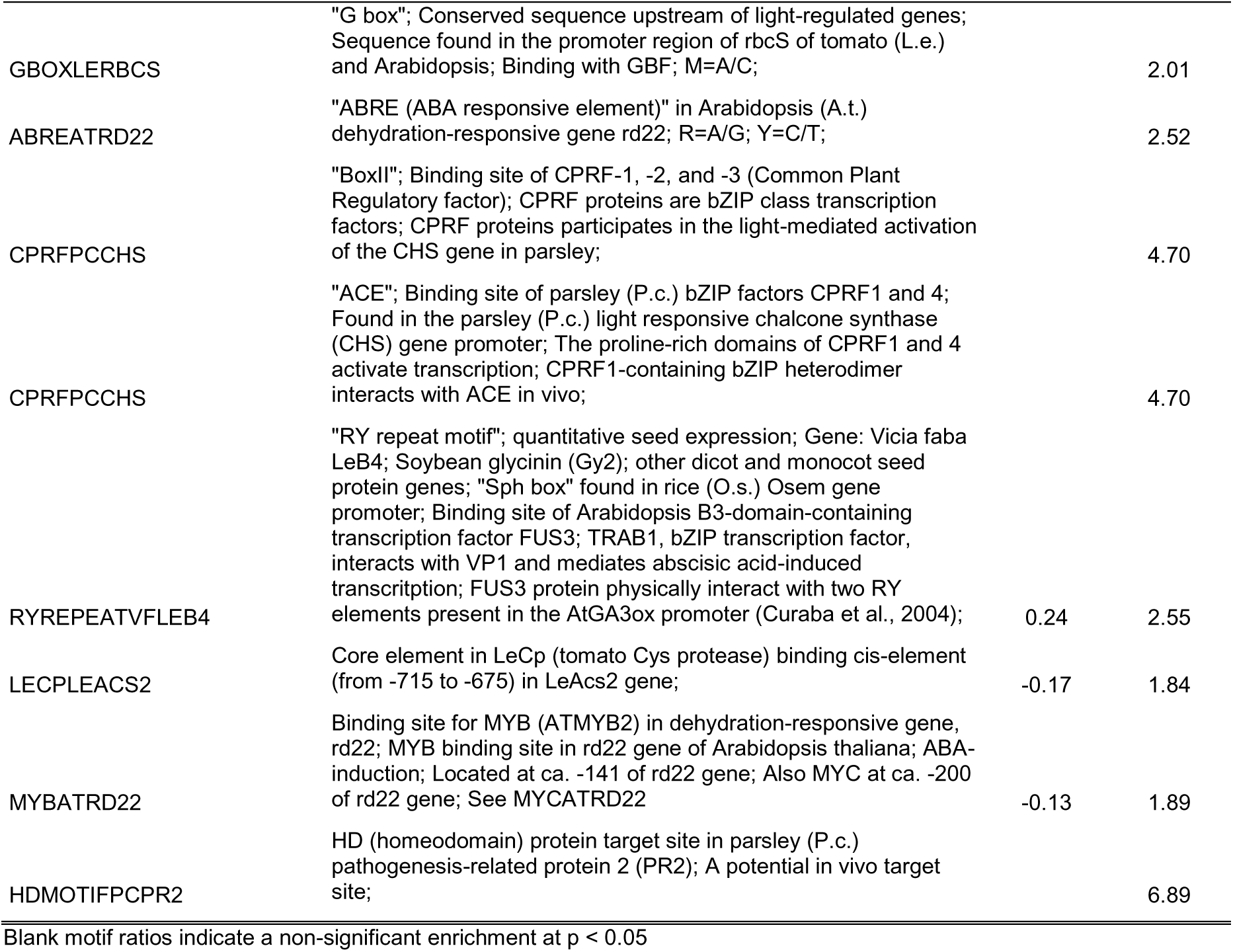
Forty interesting *cis*-element motifs enriched in highly expressed DI genes (p <0.01, motif ratio ≥1.5 significance)

Looking at the 37 DI gene set, a subset of 10 genes associated with the enriched motifs was further filtered at the 24h time point (Table 20).

**Table 20.** Subset 10 DI genes associated with motif enrichment at 24h time course

| NO. | KOME ID AND DESCRIPTION | LR* AT 24H |
| --- | --- | --- |
| 1 | AK072245 [2\|343056] *Lolium temulentum* mRNA for putative soluble acid invertase (*inv1:4* gene).\|PLN | -1.18 |
| 2 | AK061798 [9\|867346] *Arabidopsis thaliana* clone 1368 mRNA, complete sequence.\|PLN | -1.12 |
| 3 | AK069495 [3\|7044752] *Arabidopsis thaliana* hypothetical protein (*At2g23790*) mRNA, complete cds.\|PLN | 1.04 |
| 4 | AK061399 [8\|22329560] *Medicago truncatula* MTD1 mRNA, complete cds.\|PLN | 1.15 |
| 5 | AK071546 [4\|5302694] *Lolium rigidum* clone Lol-5-v putative cytochrome P450 mRNA, complete cds.\|PLN | 1.30 |
| 6 | AK073487 [8\|23208193] Rice alpha-amylase mRNA, complete cds, clone pOS137.\|PLN | 1.32 |
| 7 | AK104139 [1\|41576947] *Oryza sativa* endo-1,3-beta-glucanase mRNA, complete cds.\|PLN | 1.42 |
| 8 | AK067892 [6\|265681] *Arabidopsis thaliana* AT4g27870/T27E11_110 mRNA, complete cds.\|PLN | 1.48 |
| 9 | AK108679 [8\|23370117] *Nicotiana tabacum* patatin-like protein 1 (*PAT1*) mRNA, complete cds.\|PLN | 1.54 |
| 10 | AK069910 [1\|33468948] *Arabidopsis thaliana* putative phosphatidylglycerotransferase (*At4g04870*) mRNA, complete cds.\|PLN | 1.70 |

\*log2 ratio

In comparison, simple filtering of the DI genes for at least two-fold difference at 24h time course gives a list of 13 genes from the original 37 DI genes.

Nineteen interesting DI genes were found to be associated with enriched motifs in the 48h time course (Table 21).

**Table 21.** Nineteen DI genes associated with enriched motifs at 48 h

| NO. | KOME ID AND DESCRIPTION | LR* AT 48H |
| --- | --- | --- |
| 1 | AK066737 [12\|22816421] *Oryza sativa* mRNA for lipoxygenase (rci-1 gene).\|PLN | -1.47 |
| 2 | AK109364 [4\|23032758] Unknown expressed protein | -1.28 |
| 3 | AK064395 [9\|19874266] *Nicotiana tabacum* UDP-glucose:salicylic acid glucosyltransferase (SA-GTase) mRNA, complete cds.\|PLN | -1.17 |
| 4 | AK065631 [11\|27279069] *Arabidopsis thaliana* At2g38970/T7F6.14 mRNA, complete cds.\|PLN | -1.09 |
| 5 | AK064892 [11\|3267406] *Oryza sativa* abscisic acid- and stress-inducible protein (Asr1) mRNA, complete cds.\|PLN | 1.00 |
| 6 | AK058657 [7\|10372681] Unknown expressed protein | 1.01 |
| 7 | AK110899 [1\|5025090] *Arabidopsis thaliana* SYNC1 protein mRNA, complete cds.\|PLN | 1.05 |
| 8 | AK058289 [1\|30010659] *Oryza sativa* chlorophyll a/b binding protein (kcd1895) mRNA, complete cds.\|PLN | 1.15 |
| 9 | AK106229 [6\|17736798] *Oryza sativa* Cyt-P450 monooxygenase (PM-II) mRNA, complete cds.\|PLN | 1.16 |
| 10 | AK108523 [12\|16983665] Unknown expressed protein | 1.18 |
| 11 | AK072894 [4\|29855618] *Arabidopsis thaliana* clone 151921 mRNA, complete sequence.\|PLN | 1.22 |
| 12 | AK104994 [5\|14722691] *Triticum aestivum* clone CYP73-TA cytochrome P450 mRNA, complete cds.\|PLN | 1.26 |
| 13 | AK101002 [1\|19038898] *Arabidopsis thaliana* clone 36276 mRNA, complete sequence.\|PLN | 1.28 |
| 14 | AK071769 [3\|30245281] Unknown expressed protein | 1.30 |
| 15 | AK059592 [6\|23527383] *Serenoa repens* chloroplast partial mRNA for thylakoid structural protein (psbB gene).\|PLN | 1.36 |
| 16 | AK106319 [11\|28089044] *Oryza sativa* (japonica cultivar-group) mRNA for chitinase, complete cds.\|PLN | 1.42 |
| 17 | AK068350 [6\|21371402] Unknown expressed protein | 1.60 |
| 18 | AK101654 [2\|20001547] Unknown expressed protein | 1.67 |
| 19 | AK108349 [6\|1775726] Unknown expressed protein | 1.88 |

\*Log2 ratio

For the 48h time point, simple filtering of the genes for at least two-fold differences gives a list of 40 DI genes. Associating enriched motifs to the same DI gene list results in a shorter list of DI genes of interest.

The GO gene categories enriched in the 10 gene subset of the 24h DI gene list is interesting, with most of the gene categories being supported by literature as involved in pathways implicated in defense and biotic stress responses (Table 22).

**Table 22.**
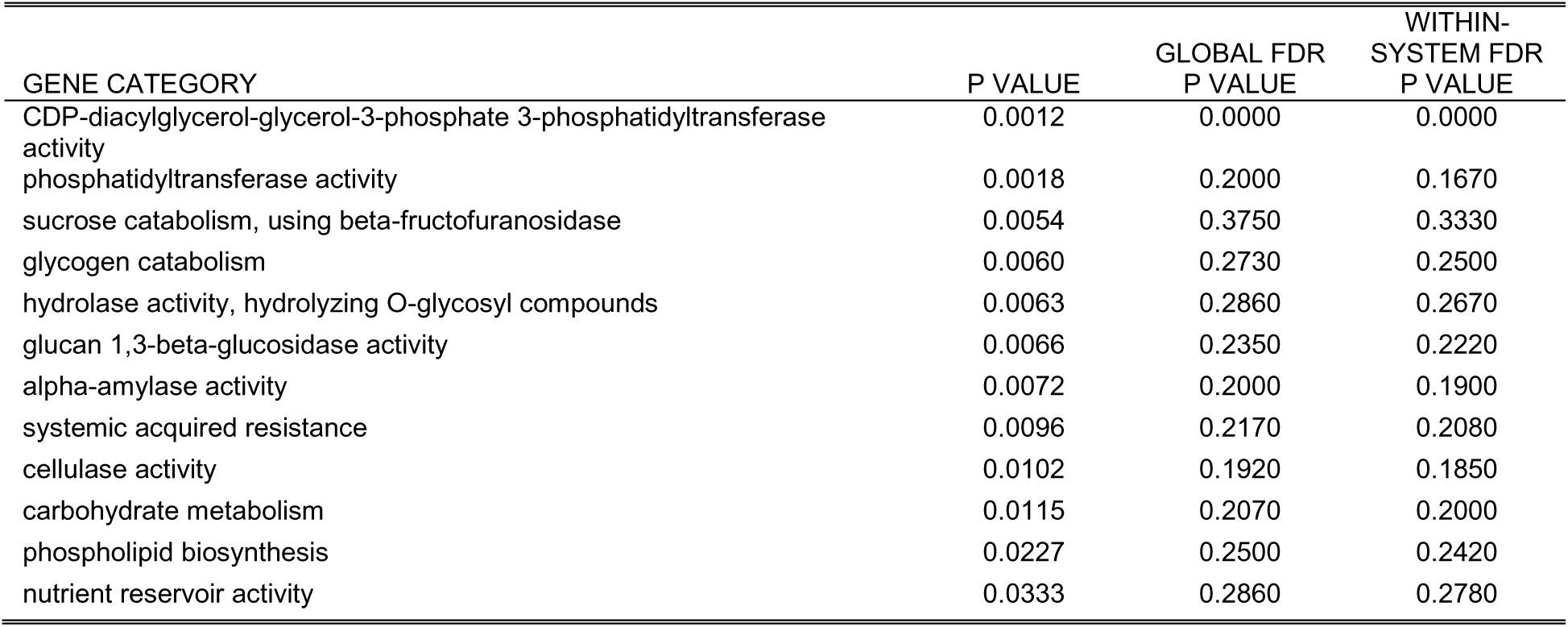
GO gene categories enriched in the 10 DI gene set associated with enriched motifs at 24h

The enriched GO gene categories of the 29 short-listed DI KOME genes associated with enriched motifs is also very interesting, with most of the gene categories associated with defense responses (Table 23).

**Table 23.** GO gene categories enriched in the 29 DI gene set associated with enriched motifs at 48h

| GENE CATEGORY | P VALUE | GLOBAL FDR P VALUE | WITHIN-SYSTEM FDR P VALUE |
| --- | --- | --- | --- |
| response to wounding | 0.0004 | 0.0000 | 0.0000 |
| trans-cinnamate 4-monooxygenase activity | 0.0012 | 0.0000 | 0.0000 |
| thylakoid membrane | 0.0018 | 0.1250 | 0.1110 |
| photosynthesis, light harvesting in photosystem II | 0.0018 | 0.0909 | 0.0833 |
| fatty acid elongase activity | 0.0024 | 0.0714 | 0.0667 |
| cell fate determination | 0.0024 | 0.0588 | 0.0556 |
| abscisic acid biosynthesis | 0.0054 | 0.2000 | 0.1900 |
| phenylpropanoid biosynthesis | 0.0060 | 0.1740 | 0.1670 |
| lipoxygenase activity | 0.0060 | 0.1540 | 0.1480 |
| wax biosynthesis | 0.0060 | 0.1380 | 0.1330 |

Table 23. continued.
| GENE CATEGORY | P VALUE | GLOBAL FDR P<br>VALUE | WITHIN-<br>SYSTEM FDR P<br>VALUE |
| --- | --- | --- | --- |
| light-harvesting complex | 0.0084 | 0.1560 | 0.1520 |
| very-long-chain fatty acid metabolism | 0.0090 | 0.1710 | 0.1670 |
| response to jasmonic acid stimulus | 0.0090 | 0.1580 | 0.1540 |
| chlorophyll binding | 0.0102 | 0.1710 | 0.1670 |
| cuticle biosynthesis | 0.0108 | 0.1590 | 0.1560 |
| microsome | 0.0108 | 0.1490 | 0.1460 |
| response to pest, pathogen or parasite | 0.0114 | 0.1600 | 0.1570 |
| photosynthesis | 0.0150 | 0.1700 | 0.1670 |
| response to water deprivation | 0.0192 | 0.1960 | 0.1930 |
| jasmonic acid biosynthesis | 0.0192 | 0.1860 | 0.1830 |
| transferase activity, transferring hexosyl groups | 0.0269 | 0.2100 | 0.2060 |
| acyltransferase activity | 0.0304 | 0.2150 | 0.2120 |
| response to light stimulus | 0.0310 | 0.2060 | 0.2030 |
| ubiquitin-protein ligase activity | 0.0363 | 0.2110 | 0.2080 |
| metabolism | 0.0414 | 0.2160 | 0.2130 |
| endoplasmic reticulum | 0.0444 | 0.2080 | 0.2050 |

In general, for the DI gene sets associated with enriched motifs, the enriched GO gene categories are associated to biotic stress and defense responses.

The enriched *cis*-elements can be classified into three distinct groups: Group 1, which is enriched at 24 h only; Group 2, enriched at 48 h only and Group 3, enriched in both timepoints, which consists only of the motif ACGTABOX. Connecting each *cis*-element with the corresponding gene(s) reveals the putative network of co-regulated gene expression in the subset of highly expressed inducible genes of the resistance transcriptome (Figures 14 to 16).

**Figure 14.**
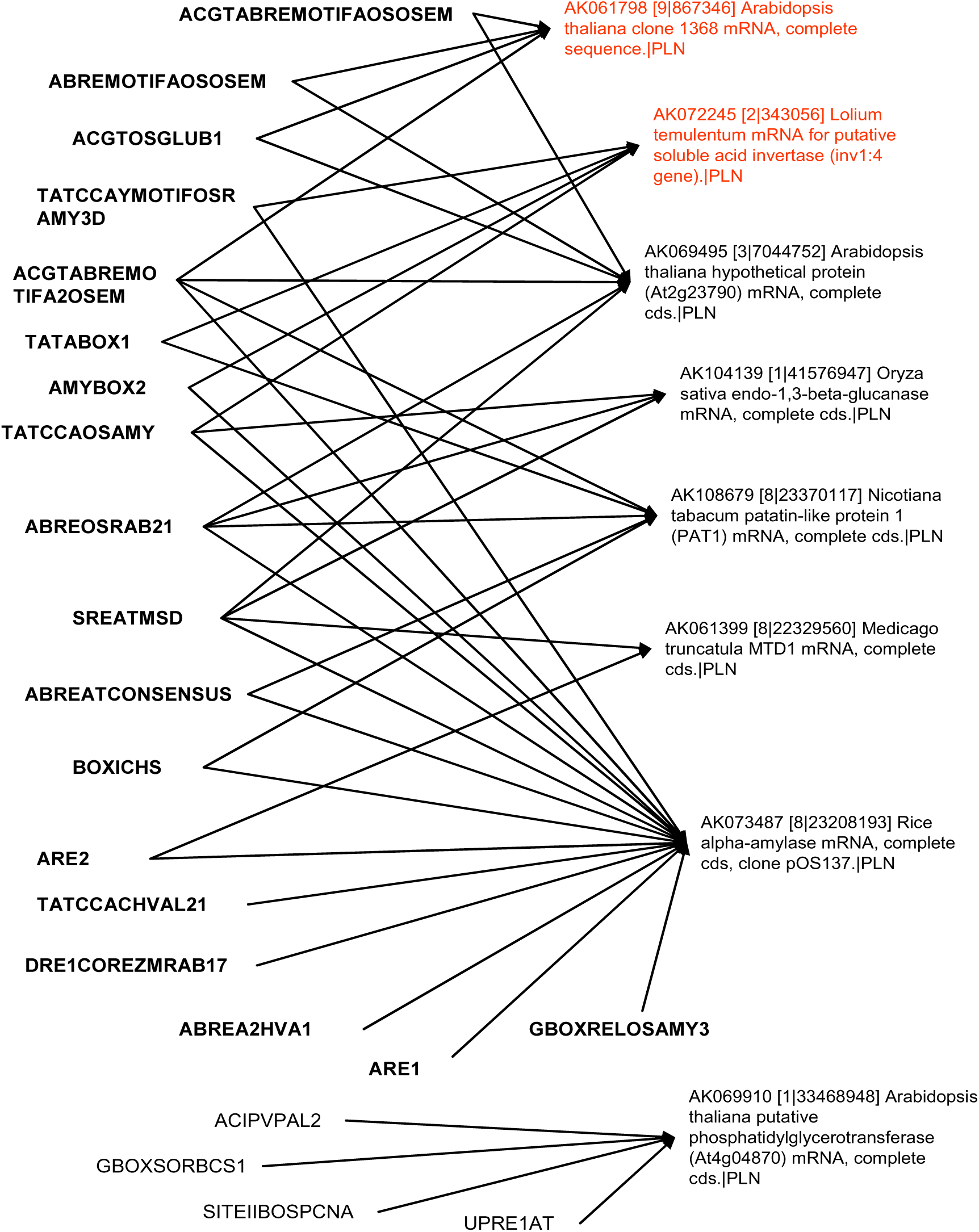
The inferred network of co-regulation of DI genes under the control of the associated enriched motifs at 24h. Genes in **red** font are down-expressed, in **black** font is up-expressed.

**Figure 15.**
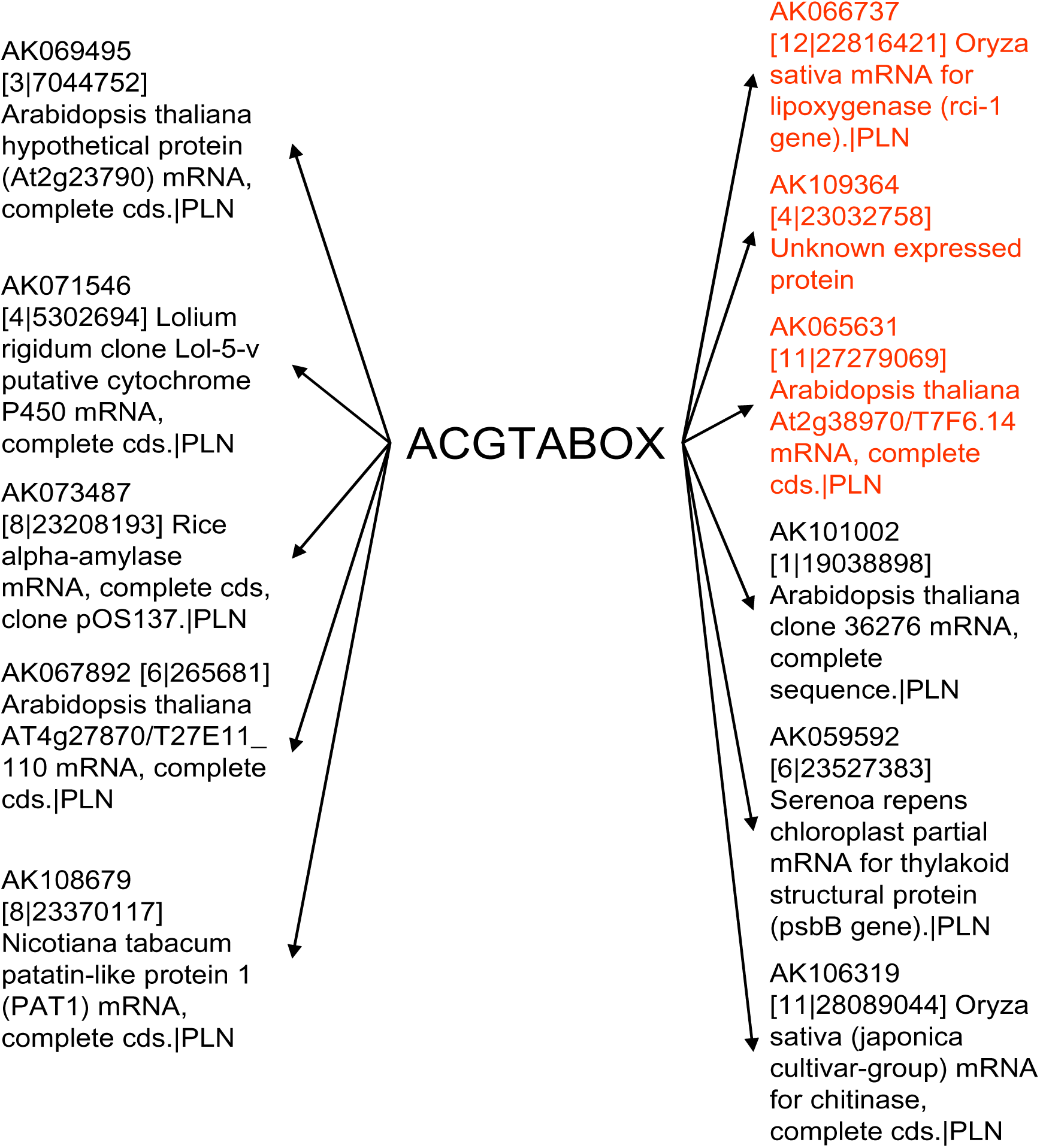
The enriched motif common at 24h and 48h and associated DI genes. The column on the left are 24h DI genes, on the right are 48h DI genes. Genes in **red** font are down-expressed, in **black** font is up-expressed.

**Figure 16.**
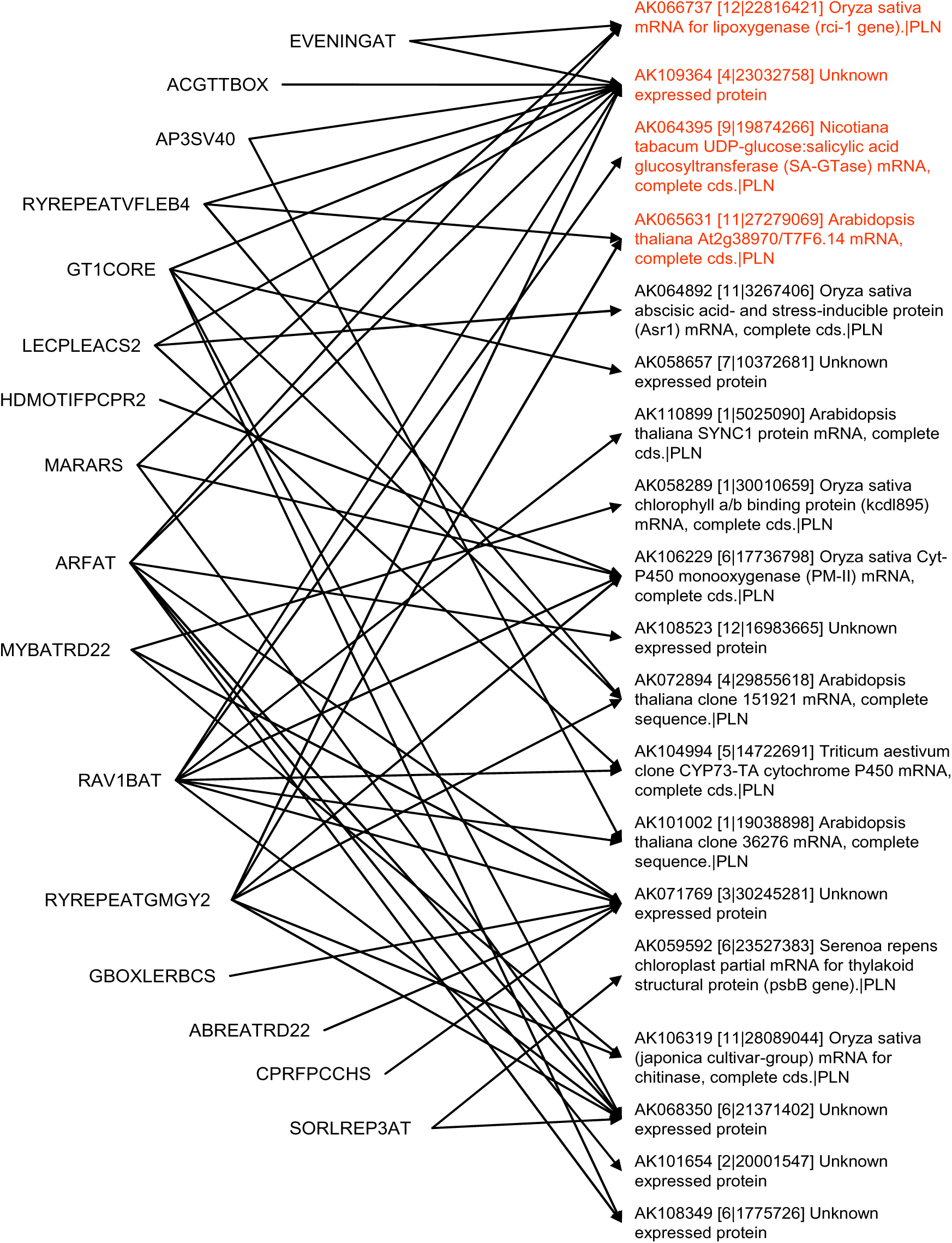
The inferred network of co-regulation of DI genes under the control of the associated enriched motifs at 48h. Genes in **red** font are down-expressed, in **black** font is up-expressed.

### Discussion

#### Assessment of the microarray experiments

The experimental results show the excellent quality of the gene expression data that can be obtained from the 22K oligoarray platform. Self-self hybridizations (same samples were labeled with cy5 and cy3, and then hybridized together, data not shown) indicate qualitatively that there is no discernible dye-gene interaction that can bias expression ratios on a gene per gene basis. Details of the signal processing algorithm by the Agilent FE software, while proprietary, is well described in general in the software manual (Agilent G2567AA manual), and all subsequent analyses utilized the processed intensities of the cy3 and cy5 channel. The proprietary use of the internal control spot intensities in the 22K oligoarray during data transformation by Agilent FE software implies that 3^rd^ party (non-Agilent) image analysis software may not be able to quantify spots into signal intensities accurately, therefore the use of an all Agilent hybridization/detection/image analysis system for this experiment appears to be the best recourse.

#### Characteristics of the GR978 disease resistance transcriptome

Looking at the set of DEGs (DI and CDE genes) for the two time points, relatively few genes were in common; essentially different sets of genes were expressed at the two time points, suggesting temporal change of genes expressed during the course of disease progress, even within a span of 24 hours. However, it is expected that many of the genes are not related to the resistance transcriptome but are involved in other plant growth, development, and maintenance processes as well.

While the physical boundaries defined by the genetic markers that were found to be associated with the resistance phenotype is fuzzy at best due to expected differences in the japonica vs. indica genomes, as well as the nature of quantitative trait mapping itself (which is statistical in nature), saturation mapping of the genetic region of interest with SSR markers revealed a very high association of the region with quantitative blast resistance, with the linkage order of markers showing a high degree of concordance to the physical genome map (Madamba et al, 2009). From the physically translated genetic region, a relatively small set of genes was found to be differentially expressed. Both up and down expressed genes were selected as candidates since it is not yet clear how the gain of resistance mechanism works in GR978; it may be over-expression of a disease response related gene or the knock-out of function of a negative regulatory gene that normally suppresses expression of either defense response or programmed cell death as in the *spl11* rice mutant reported by Zeng et al. (2004).

Since the location of the mutation is known, taking a liberal approach to identify candidate genes in the target region seems appropriate to avoid making a false negative declaration (committing Type II error). Of the three genes with known functional annotations listed, one (AK061410 – wheat mRNA for Rubisco subunit binding-protein alpha subunit) is not associated by literature to any disease response specific pathway. The annotation of AK099555 (*Zea mays* mRNA for putative glutathione synthetase (gsh2 gene) is related by literature to oxidative response, while the annotation of AK064488 (*Arabidopsis thaliana* selenium-binding protein-like (MPO12.16) mRNA) is associated by literature with metal binding; both are involved in SA and JA-induced defense response in sorghum (Salzman et al, 2005). The characterization of the 9 genes of unknown function may lead to the discovery of novel genes involved in the resistance response. In all, given the short list of genes, it may be worthwhile to do experiments (e.g. qRT-PCR, Northern hybridization) to validate the function of the twelve candidate genes.

Looking beyond the candidate genes in chromosome 12, a particular gene family of interest in our group is the oxalate oxidase family, which is highly associated with disease resistance in other studies (Leach, et al. pers comm., Colorado State University). Using the DI gene list obtained from the t-test based significance analysis, several instances of the oxalate oxidase gene were found as differentially expressed in GR978 (Table 24); it is highly up-expressed in chromosome 4.

**Table 24.** Oxalate oxidase genes significantly expressed in GR978 experiments as determined by the joined t-tests.

| KOME ID | CHR | DESCRIPTION | LOG2 RATIO<br>AT 24H | LOG2 RATIO<br>AT 48H |
| --- | --- | --- | --- | --- |
| AK061164 | 1 | Oryza sativa germin-like protein 4 (GER4)<br>mRNA, complete cds.\|PLN |  | 0.171 |
| AK108987 | 9 | Medicago truncatula germin-like protein 1<br>mRNA, complete cds.\|PLN |  | 0.088 |
| AK109057 | 4 | Oryza sativa germin-like protein 6 (GER6)<br>mRNA, complete cds.\|PLN | 1.73 |  |

#### Clustering co expressed genes

Informally partitioning the DI and CDE gene list into which timepoints they exhibit significant expression enables a quick view as to which genes are active during particular stages of disease resistance development. For the group of down-expressed genes at 24h and/or 48h, inspection of three known function annotation (axi 1 protein, nucleosome/chromatin assembly factor, mutator transposase) shows no apparent association to defense responses. Interestingly, AK101370 (DNAJ homologue) is implicated in defense response pathways (Marathe et al. 2004), but collectively, this group is likely not associated with the mutation causing the gain of resistance phenotype of GR978.

The constitutive behavior of GR978 (looking at known function annotation) during the early stage of disease resistance response (24h) appear to be the up-expression of genes involved in general disease reaction responses, as supported by literature. Reactive oxygen intermediate (ROI) related genes are implicated in HR and PCD, and genes such as cytochrome P450 proteins (AK067458), cytochrome P450 monooxygenase (AK108697) belong to this class (Marathe et al, 2004), as well as the hypersensitive-induced response protein (AK103475). DNA-binding proteins (AK102284) are involved in transcription reprogramming that occurs during early resistance reaction events, which associates this gene to the defense response (Dangl and Jones 2001). Resistance-related transporters such as phosphate transporter (AK063990) and ammonium transporter (AK100411) are also in this group. Genes for defense signaling (wall-associated kinase 2, AK100387) and phenylpropanoid/secondary metabolites (glucosyltransferase, AK109007) are also expressed. The constitutive expression at 24h of these disease resistance related genes imply that the GR978 mutation may have de-regulated gene(s) upstream involved in early disease resistance response of the resistance pathway. Other gene(s) controlling late resistance responses may have been expressed at 48h and caused the return to a normal expression state of these genes.

For the group of genes constitutively up-expressed at both time points, it is suspected that there is de-regulation of expression of defense related genes (HR-PCD related gene peroxisomal membrane protein, AK110421; BCS1 mitochondrial chaperone, AK101987; secondary metabolite product B-amyrin (phytoalexin) synthase, AK106822), may be associated with the mutation that caused the gain of resistance.

GR978 also shows constitutive expression of many genes in the later stage of the resistance response (48h), many of which are implicated by literature to be involved in plant defense. Constitutive expression at 48h suggests loss of late regulatory control of these genes. Among the genes implicated in the disease response are: (1) HR/HR-signaling/ PCD related genes such as AK064654 (protein inhibitor of neuronal nitric oxide), AK110372 (calmodulin-binding protein), and AK072715 (cytochrome P450), (2) Resistance/pathogenesis related genes such as AK059767 (bulb chitinase-1), AK063939 (class III chitinase homologue), and AK068220 (phosphatidylserine receptor) and AK108761 (delta-cadiene/phytoalexin synthase), (3) transcription regulation gene AK067427, (4) Fatty Acid (FA) metabolism/JA synthesis related gene AK066825 (lipoxygenase), (5) Defense signaling gene AK105217 (protein kinase CRINKLY 4 receptor), (6) transporter proteins AK100175 (ABC transporter) and AK108373 (tamdr1 ABC transporter), and (7) protease –related RRF1 RING finger gene (AK108851). There are genes not implicated by literature to the resistance response that are also constitutively up-expressed, such as auxin-induced protein (AK103553), leaf development protein (AK069685), and telomere-binding protein-1 (AK086473); the constitutive up-expression these genes are likely not caused by the mutation resulting in the gain of resistance phenotype of GR978.

The resistance phenotype of GR978 appears to be caused by the set of inducible genes as well. Genes with known function associations were inspected for the informal groups. Looking at the induced genes during the early resistance response (24h), a down-expressed group contains two genes implicated in defense response (AK066737, lipoxygenase, for FA/JA synthesis; chitinase AK106178). This is paradoxical considering that defense responses require up-regulation of the gene products. The exclusively 24h induced genes, all up-expressed, are interesting: it is composed of defense-related genes such as HR/PCD related gene AK071546 (cytochrome P450) and resistance/defense related genes AK099157 (probenazole-inducible protein, PBZ-1), AK104319 (endo-1,3-beta-glucanase) and AK108679 (PAT1 protein). The group of genes exclusively expressed at 48h is also composed of many defense-related genes such as AK064892 (ABA-stress inducible protein), HR/PCD-related genes such as AK106229 (Cytochrome P450 monoxygenase), AK104994 (Cytochrome P450 protein), AK059143 (cytochrome b-559 beta subunit) and AK062117 (glutaredoxin/glutathione reductase), SA-coregulation associated gene AK108346 (WRKY transcription factor), and resistance/defense gene AK106319 (chitinase). It appears that the mutation conditioning the gain of resistance phenotype in GR978 induces the resistance transcriptome composed of the defense-related genes discussed. It is also evident that many genes that are not implicated in defense are also induced in the experiments.

The collection of defense-related genes in the DI and CDE gene sets suggest that the gain of resistance phenotype is the result of both constitutive and inducible expression of genes implicated by literature to be belonging to pathways involved in basal plant defense response, which is consistent with broad-spectrum resistance phenotype shown by GR978. Notably absent in the DEGs detected, however, are the disease resistance genes of the NB-LRR class, which are involved in specific interactions between pathogens and host plants. The genetic overlap between specific resistance and basal resistance responses suggest that R-mediated signaling serves to activate defense mechanisms shared in both pathways more rapidly and effectively (Dangl and Jones 2001). The absence of such expressed R-genes in GR978, however, implies that non-R gene mediated signaling processes upstream to the basal defense pathways are disrupted by the mutation, causing enhanced or de-regulated expression of the genes in this pathway.

There is a huge gap in knowledge presented by the genes with unknown function annotation, which presents opportunity in the discovery of novel genes involved in defense response. Computational methods were employed in order to infer putative functions to the unknown genes, as well as determine the 2^nd^ order relationships among the genes composing the resistance transcriptome, results of which are presented in the next chapter.

Inspection of the function annotations of each gene from the CDE and DI genes list gives insight into the sets of genes that are associated with defense and biotic stress response. However, directly characterizing each of the significantly expressed genes is a tedious task. By doing hierarchical clustering of the highly expressed gene sets, genes of the resistance transcriptome (with or without function annotations) are clustered into groups exhibiting similar expression patterns across the two time points. Genes clustering to the same group imply their involvement in a common process or pathway.

HCL partitioned the defense-related genes into separate groups with a coherent defense response theme, implying the co-regulation of a particular gene group. For instance, in the CDE Group 1 are 16 genes with function annotations implicated by literature to be involved in defense response of plants, broadly categorized as HR-signaling, HR-PCD, and resistance-pathogenesis related. From this big group, three gene subgroups can be observed: (1a) a mostly 48h expressed group with 3 defense-related members, (1b) a group expressed at both 24h and 48h with 11 defense-related members, and (1c) a mostly 24h expressed group with six defense-related gene members (Table 25).

**Table 25.**
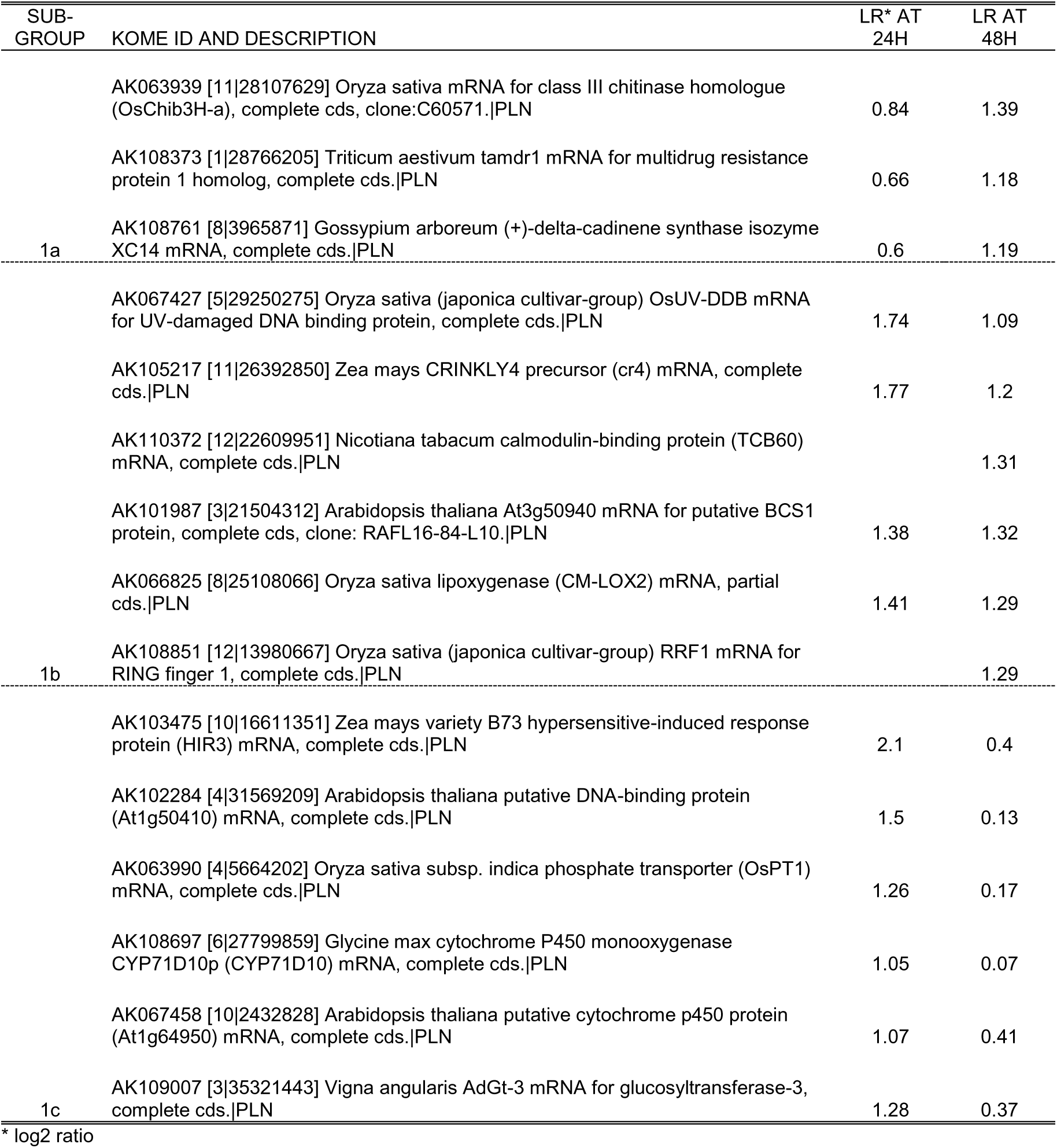
Disease resistance related genes in 3 subgroups of HCL group 1 CDE

In all these subgroups, the three broad classes of defense responses are represented. The relative time as to which each defense response category is preferably expressed is resolved by HCL.

In the second CDE group with highly expressed members, the three broad defense response classes are also represented also. The third CDE group is paradoxical, the two defense-related members are down-expressed, but literature shows that these genes should be up-expressed during defense response.

Some genes annotated with unknown function are clustered among genes of known function by HCL, thus providing insight on their probable function. For instance, AK072262 in CDE group 1c tightly clusters with genes whose annotated functions are involved in early resistance engagement events (AK067427 – DNA binding protein, AK105217 – CRINKLY4 kinase, AK110372 – calmodulin-binding protein), implying its involvement in early constitutive resistance activation. Many unknown genes do tend to cluster closely together, and the cluster formed is distinct from other genes of known function (i.e., CDE group 1c – AK105155, AK070117, AK107432, AK108174), thus failing to shed light on their probable role, but at the same time implying the participation of these genes in a common pathway. One unknown gene (AK107552) clusters with another gene (AK103553 – auxin-induced protein) whose annotation does not associate with defense at all, implying non-participation of the unknown gene in defense response. Most of the unknown genes are found everywhere among the clusters, however.

For the DI genes, HCL defines two very distinct groups in terms of defense response. The down-expression of the first DI group is contrary to expected up-expression of resistance/pathogenesis and FA/JA pathway genes, implying the non-involvement of this group in the resistance response. DI group 2 resolves into a mostly of 24h up-expressed genes, majority of which are resistance/pathogenesis-specific in function (AK099157-PBZ-1, AK104139-endo-glucanase, AK108679-PAT-1); only one is involved in HR/PCD (AK071546-cytochrome p450 protein). DI group 3 consists of genes mostly up-expressed at 48h, and a diverse group of defense-related functions are present, such as HR/PCD, resistance/pathogenesis, and SAR-coregulation. HCL clearly resolves which gene functions are induced during the early and later stages of the resistance reaction: early disease resistance appears to be characterized by resistance/pathogenesis responses, and the HR/PCD as well as the SAR responses subsequently being expressed.

The expression of WRKY transcription factor (AK108346) suggests the involvement of SAR-coregulated genes in the gain of resistance phenotype of GR978. Maleck et al. (2000) reported that the *cis*-element bound by the plant-specific WRKY transcription factors is the common element in a set of SAR co-regulated genes. The core binding site of WRKY transcription factor is the W box with sequence TTGAC (Eulgem et al 2000). A scan of the 1kb upstream region of the genes in DI group 3 reveal three genes with two or more W box elements (AK058657, AK068350, and AK071769); however, the W box elements are not enriched in the DI group 3 set (p > .01). All three genes with upstream W box elements are not annotated for known function; these may be novel genes involved in SAR response. Gene AK071769 in particular belongs to a tight cluster with two other unknown genes AK108523 and AK066176, suggesting that these two unknown genes may also be part of the SAR response as well. The GR978 mutation appears to have induced SAR responses ultimately leading to induction of SAR-coregulated genes.

Most of the other genes of unknown function appear in many of the clusters in DI group 3. However, the clustering of many unknown genes with genes of known function provides additional information as to whether an unknown gene participates in defense response or not. Hierarchical clustering is powerful in determining interesting grouping of the genes involved in the defense response. It provides inferences on the putative function of the genes with unknown function via association of function with genes of known function within the same cluster. Clearly, HCL was useful in partitioning the genes into biologically meaningful clusters that define the resistance response at particular time points, even with the limited set of independent time-course experiments available. However, it does not automatically reveal the biological relationships among the genes in the list among the clusters; the annotation of each member gene of an interesting cluster must be inspected one at a time and compared to prior studies in order to recognize a unifying functional theme for the particular cluster under study. Given that the set of genes in the 22K oligoarray is fairly well annotated, the inability to include biological information available for the genes under scrutiny is a shortcoming of HCL, as well for other purely computationally based methods such as k-means and self-organizing maps.

#### Biological themes in the GR978 disease resistance transcriptome

The GO gene category enrichment analysis method used to infer higher-order biological themes from the subset gene list quickly recapitulates known biological processes that are involved in plant defense response. For both DI and CDE gene lists, the gene category terms observed as enriched are mostly processes related to rapid plant defense events such as HR/PCD processes, as well as processes leading to enhanced resistance (SAR-related events), implying that both general defense response pathways are constitutively occurring as well as inducible in GR978. For CDE group 1, both rapid and general defense response processes were enriched, while disease response-related events are not observed in CDE group 3, implying that significant expression of the member genes of this cluster is not defense related DI groups 1 and 2 also show enriched gene categories implicated in rapid and long-term resistance responses. DI group 3 is not resolved by gene category enrichment as a purely defense response group even though many genes in the group are annotated for functions that are implicated by literature as defense-related; either this co-expression of defense and non-defense related genes is coincidental or there is a gene regulatory network that intersects both non-defense and defense-related processes. This phenomenon is not uncommon; Cooper et al (2003) reports such putative network of rice genes associated with both stress response and with seed and plant development.

While manual inspection of the annotation of each gene in the list is already informative of the probable biological processes underlying a selected list of genes, this requires the tedious process of researching literature about what higher-level process each gene is a member of. Furthermore, manual annotation inspection attaches no measure of significance of enrichment for a particular biological category observed in the list. There are several shortcomings of this method: the first lies in the amount of prior biological information available for the set of genes under study. In the KOME reference gene set spotted in the 22K oligoarray used (actual total of 21,495 genes), only 14,895 of them are annotated with GO terms; the rest are assigned as unknown. If a short gene list analyzed for GO term overrepresentation has an abundance of genes without GO annotations, no informative biological themes can be inferred from this gene list. A second shortcoming is that only the most significantly expressed set is considered; the subtly expressed genes, which may be the case for many transcription factor genes, are ignored. Furthermore, the order of gene expression (from the most to the least highly expressed genes) is discarded. The third shortcoming is that information that some genes are of correlated expression is also discarded. Nonetheless, this method is a de facto standard for the secondary analysis of differentially expressed genes in an expression experiment and as shown in the study, provides quick and useful inferences on the higher order biological process involved in defense response of GR978 against rice blast. Recently, there have been efforts by computational biologists and mathematicians to incorporate biological knowledge into distance-based clustering analysis methods, which may address the shortcomings mentioned (Huang and Pan 2006).

#### The inferred network of co regulated disease response genes

The simple statistical method of determining enriched motifs in a subset of induced genes enables the partitioning of the gene list into either of the following groups: (1) genes under the control of a shared set of cis elements bound by a common binding factor, implying co-regulation of this group by the same transcription factor/group of TFs, and (2) genes under the control of *cis* elements bound by many types of binding factors, implying the participation of these genes in other diverse pathways.

Several shortcomings of the method must be pointed out, such as that the upstream sequences used came from the Nipponbarre reference genome sequence, not from the actual genotype under investigation. Upstream sequence polymorphism between varieties/genotypes can change the occurrence and abundance of *cis* elements in genes. No information about what transcription factors bind which motifs in the context of the rice genome is given. The method also does not impose any of the constraints involved in transcription activation such as the distance between cis elements and to other promoter elements, as well as the interplay of different motifs (motif synergism/antagonism) that also affects transcription activation. Nevertheless, the method provides an insight to the putative network of interaction existing for the list of DEGs under study. From the motif enrichment analysis, it is interesting to note that a distinct set of motifs is enriched in a particular time point, implying that the set of genes coregulated at 24h and 48h is under the control of different mechanisms. The action of WRKY transcription factor was suspected since the GO gene category of systemic acquired resistance is significantly enriched in the 24h DI set, but the motif bound by WRKY (TTGAC -W box) is not enriched in the same gene set (data not shown).

Many of the enriched motifs are annotated as involved in abiotic stress responses (such ABRE/ABA-responsive elements, bZIP, DRE1, MYB, light responsive/regulated elements) as well as general function elements (amylase activity, auxin response factor binding site, common plant regulatory factor binding site). Interesting motifs implicated in resistance / pathogenesis responses were also found in the enriched motifs (PR2 binding site HDMOTIFPCPR2, Cys protease binding element LECPLEACS2). The lack of higher order categories makes the classification of motif elements a tedious process. Nonetheless, partitioning genes into putative co-regulated groups via enriched motifs offers another way of viewing and analyzing microarray experimental data. Furthermore, the basis of grouping by enriched motifs is not purely computational but relies on biological information, that is, *cis*-element-gene association and the motif’s relative abundance in the gene set of interest.

## III. GLOBAL CHARACTERISTICS OF THE RESISTANCE TRANSCRIPTOME WITHIN AND ACROSS RESISTANT VARIETIES

### Results

The characteristics of the resistance transcriptome within and beyond a genotype was examined by mining gene expression data from the mutant GR978 and other rice varieties (SHZ-2, Nipponbarre) undergoing disease resistance response. Resistance transcriptome characterization was carried out at different levels; the first is comparison of individual DEGs between the varieties/genotypes and examining the co-expression profile, if possible, of the common DEGs. The second level of characterization computed for the correlation structure of gene expression along the genome in order to find Regions of Correlated Gene Expression (RCE) within each genotype/variety. The similarities and differences of the RCEs between GR978 and SHZ-2 was established by (a) characterizing the gene members of these regions, (b) determining the genomic distribution of these RCEs, and (c) statistical determination for enrichment of genotype-specific and common DEGs within the RCEs. In the third level of characterization, rice blast resistance-specific QTL mapping data was used to partition the genome into Bl-QTL meta-regions, and the association of these Bl-QTL regions with the genotype-specific and common DEGs and RCEs was determined.

#### The common DEGs across varieties/genotypes

The list of DEGs from GR978 were combined with DEGs obtained from two studies: the first is from the blast resistant variety SHZ-2 undergoing disease resistance response (B. Liu et al., unpublished data), and the second study from a japonica variety, Nipponbarre, artificially induced for defense response with BTH (Hirochika et al., unpublished data).

In the SHZ-2 study, the experimental design (split-plot microarray design) allowed for the computation of separate DI and CDE gene sets; each list was compared to the corresponding DI and CDE list from GR978. The common genes are analyzed for co-expression using HCL and the result is summarized in the figures 14 and 15. For the common CDE genes (Fig. 17), the most similar expression profiles are within variety-specific experiments (GR978 24h and 48h experiments; SHZ-2 24h and 48h experiments). The expression profiles of common DI genes appear to be influenced by the variety/genotype and the time of sampling used, with the GR978-24h and SHZ-2-48h profiles being most similar; the GR978-48h expression profile is less similar to the two profiles, and the SHZ-2-24h expression profile is the least similar among the experiments compared (Fig 18).

**Figure 17.**
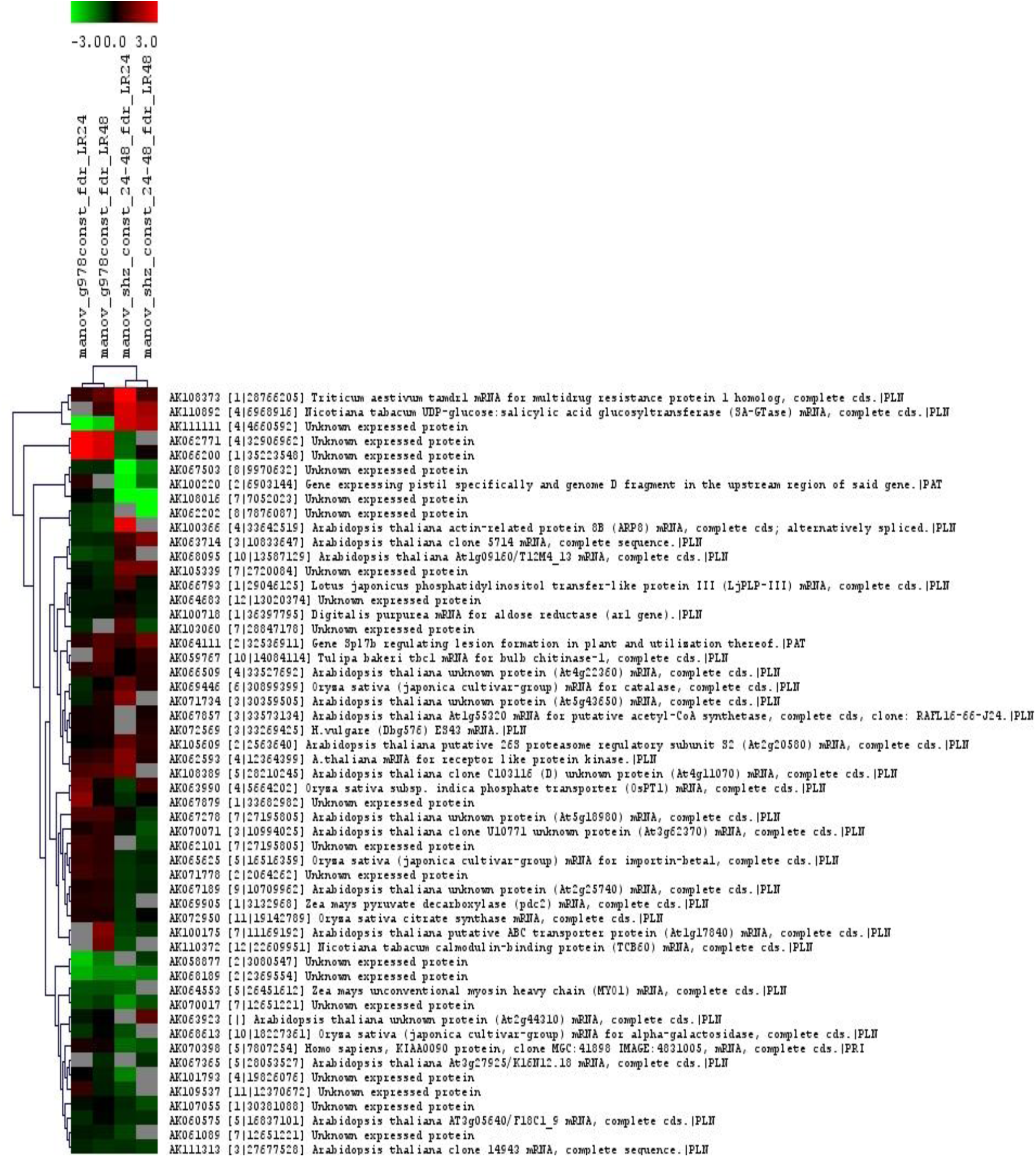
Two-way HCL of 53 common CDE genes between GR978 and SHZ-2. The tree on top shows similarity of the experiments; the tree at the left shows similarity of genes.

**Figure 18.**
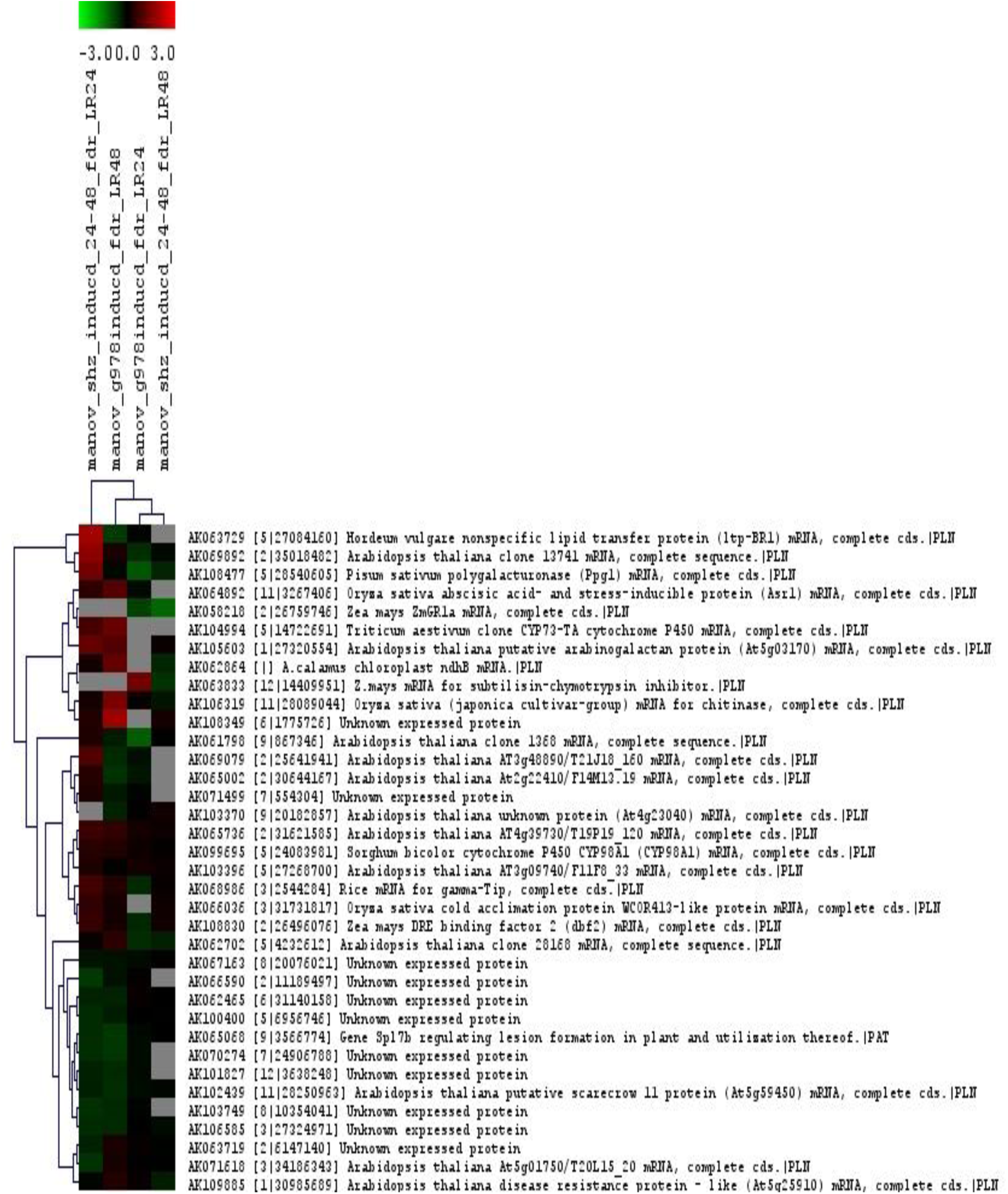
Two-way HCL of 36 common DI genes between GR978 and SHZ-2. Clustering of experiments and genes shown in top and left side of expression image, respectively.

The biological themes of the common CDE gene lists from GO category enrichment analysis are shown in Table 26.

**Table 26.**
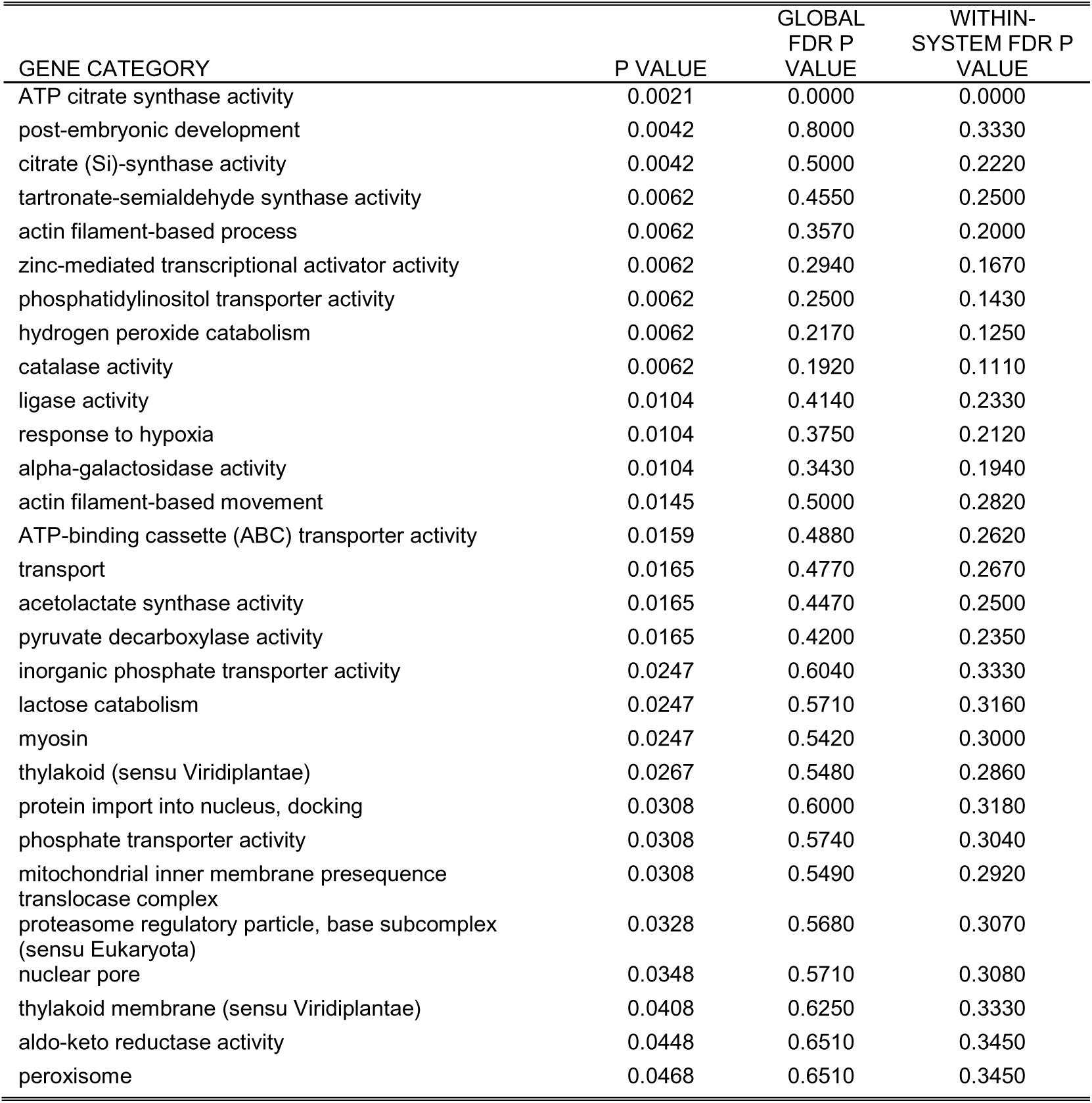
GO gene category enrichment analysis for the CDE genes common between SHZ-2 and GR978

For the common DI genes, the result of GO category enrichment analysis is shown in Table 27.

**Table 27.** GO gene category enrichment analysis of DI genes in common between GR978 and SHZ-2

| GENE CATEGORY | P VALUE | GLOBAL FDR P VALUE | WITHIN-SYSTEM FDR P VALUE |
| --- | --- | --- | --- |
| phenylpropanoid biosynthesis | 0.0001 | 0.0000 | 0.0000 |
| p-coumarate 3-hydroxylase activity | 0.0015 | 0.0000 | 0.0000 |
| trans-cinnamate 4-monooxygenase activity | 0.0031 | 0.2500 | 0.1110 |
| lipid transport | 0.0031 | 0.1820 | 0.0833 |
| lipid binding | 0.0033 | 0.1430 | 0.0667 |
| vesicle-mediated transport | 0.0199 | 1.4700 | 0.7780 |
| SOS response | 0.0229 | 1.4000 | 0.7140 |
| endomembrane system | 0.0273 | 1.4300 | 0.7500 |
| Microsome | 0.0274 | 1.2700 | 0.6670 |
| ATP-dependent DNA helicase activity | 0.0320 | 1.2800 | 0.6670 |
| regulation of translation | 0.0349 | 1.2200 | 0.6360 |
| vacuolar membrane (sensu Magnoliophyta) | 0.0364 | 1.1400 | 0.6110 |
| polygalacturonase activity | 0.0379 | 1.0800 | 0.5640 |
| water channel activity | 0.0439 | 1.1200 | 0.5710 |

Similar to the results of the GR978-only experiments, enriched GO gene categories directly or indirectly involved in defense response are evident in both CDE and DI genes common between GR978 and SHZ-2.

The set of DEGs from GR978 was compared to 158 DEGs detected from BTH-induced Nipponbarre. No distinction between DI and CDE genes were available for the Nipponbarre-BTH DEGs, therefore comparisons were made to union of the CDE and DI genes from GR978. There were 14 DEGs found in common between the two genotypes, shown in Table 28.

**Table 28.** The 14 DEGs in common with GR978 and BTH-induced Nipponbarre (BTH-Npb) and their respective log2ratios at the timepoints under study

| NO | KOME ID AND DESCRIPTION | BTH-NPB<br>LR*24H | BTH-NPB<br>LR72H | GR978<br>CDE<br>LR24H | GR978<br>CDE<br>LR48H | GR978<br>DI<br>LR24H | GR978<br>DI<br>LR48H |
| --- | --- | --- | --- | --- | --- | --- | --- |
| 1 | AK059154 [7\|26704202] *Oryza sativa* 10 kDa chaperonin mRNA, complete cds.\|PLN | -0.09 |  | 0.39 |  |  |  |
| 2 | AK064395 [9\|19874266] *Nicotiana tabacum* UDP-glucose:salicylic acid glucosyltransferase (SA-GTase) mRNA, complete cds.\|PLN | -0.41 |  |  |  |  | -1.17 |
| 3 | AK065631 [11\|27279069] *Arabidopsis thaliana* At2g38970/T7F6.14 mRNA, complete cds.\|PLN | 0.02 |  |  |  |  | -1.09 |
| 4 | AK066255 [5\|14814283] *Oryza sativa* (indica variety-group) ABA (chilling-regulated) mRNA, partial sequence.\|PLN | 0.05 | -0.16 |  |  |  | -0.65 |
| 5 | AK066737 [12\|22816421] *Oryza sativa* mRNA for lipoxygenase (rci-1 gene).\|PLN | 0.04 | -0.06 |  |  |  | -1.47 |
| 6 | AK066760 [3\|31384665] *Lolium rigidum* clone FHH-t putative cytochrome P450 mRNA, complete cds.\|PLN | -0.15 |  |  | 0.65 |  |  |
| 7 | AK068652 [4\|15546041] *Helianthus annuus* auxin-induced protein (HaAC1) mRNA, complete cds.\|PLN | 0.01 |  |  |  |  | -0.49 |
| 8 | AK099157 [12\|22553507] Rice mRNA for probenazole-inducible protein PBZ1, complete cds.\|PLN | -0.42 |  |  |  | 1.42 |  |
| 9 | AK103553 [4\|15658725] *Arabidopsis thaliana* auxin-induced protein, putative (At1g60710) mRNA, complete cds.\|PLN | 0.42 |  |  | 1.08 |  | -0.98 |
| 10 | AK103699 [10\|20123161] *Oryza sativa* subsp. japonica clone S14511 putative glutathione S-transferase OsGSTU6 mRNA, complete cds.\|PLN | -0.25 |  |  |  | -0.53 |  |
| 11 | AK106178 [1\|28334468] *C. arietinum* mRNA for chitinase.\|PLN | -0.18 |  |  |  |  | -1.03 |
| 12 | AK108373 [1\|28766205] *Triticum aestivum* tamdr1 mRNA for multidrug resistance protein 1 homolog, complete cds.\|PLN | -0.17 |  |  | 1.19 |  |  |
| 13 | AK108389 [5\|28210245] *Arabidopsis thaliana* clone C103116 (D) unknown protein (At4g11070) mRNA, complete cds.\|PLN | 0.11 |  |  | 0.97 |  |  |
| 14 | AK109364 [4\|23032758] Unknown expressed protein | -0.02 |  |  |  |  | -1.28 |

\*LR- log2 ratio

The common DEGs are interesting as it is composed mostly of either disease resistance or abiotic stress response genes. The enriched GO gene categories from the 14 common DEGs shown in Table 29, provides further support that the common DEGs are mostly resistance response genes.

**Table 29.** GO gene categories enriched in DEGs common in GR978 and BTH-induced Nipponbarre

| GENE CATEGORY | P VALUE | GLOBAL<br>FDR P<br>VALUE | WITHIN-<br>SYSTEM FDR P<br>VALUE |
| --- | --- | --- | --- |
| cytoplasm | 0.0044 | 2.0000 | 1.0000 |
| abscisic acid biosynthesis | 0.0066 | 1.4000 | 0.6670 |
| lipoxygenase activity | 0.0074 | 0.8750 | 0.4440 |
| response to sucrose stimulus | 0.0088 | 0.9090 | 0.4170 |
| voltage-gated potassium channel activity | 0.0088 | 0.7140 | 0.3330 |
| regulation of transcription, DNA-dependent | 0.0106 | 0.6470 | 0.3330 |
| response to jasmonic acid stimulus | 0.0110 | 0.5500 | 0.2860 |
| response to pest, pathogen or parasite | 0.0139 | 0.5650 | 0.2920 |
| aldo-keto reductase activity | 0.0161 | 0.5770 | 0.2960 |
| transcription factor activity | 0.0225 | 0.6210 | 0.3330 |
| response to water deprivation | 0.0234 | 0.5940 | 0.3030 |
| jasmonic acid biosynthesis | 0.0234 | 0.5430 | 0.2780 |
| DNA methylation | 0.0263 | 0.5260 | 0.2560 |
| toxin catabolism | 0.0313 | 0.5610 | 0.2620 |
| glutathione transferase activity | 0.0327 | 0.5230 | 0.2670 |
| transferase activity, transferring hexosyl groups | 0.0327 | 0.4890 | 0.2500 |
| response to wounding | 0.0363 | 0.5000 | 0.2550 |
| ubiquitin-protein ligase activity | 0.0442 | 0.5280 | 0.2590 |

#### The correlation structure of the resistance transcriptome within and across genotypes

The genome-wide behavior of the resistance transcriptome was characterized using the method described by Spellman and Rubin (2002) for Drosophila transcription analysis, to determine if there were regions of the rice genome showing condition-specific correlated expression against blast infection. Correlation in expression between adjacent genes was calculated in a moving window starting from 2 genes all the way to 15 genes per window, sliding one gene at a time. Using expression data from six experiments comparing un-inoculated with blast-inoculated GR978, regions of correlated expression were determined. Looking at the net number of correlated genes (difference between the number of correlated genes in the genome-ordered and randomized datasets) expressed from a window size of two to 15 genes, the net number of correlated genes was at maximum at a window size of ten genes (Figure 19). This indicates that correlated expression groups during resistance response in GR978 include about ten gene models as resolved by the 22K oligoarray used, and subsequent correlation analyses at different significance p-values were done using the ten gene window. The other gene windows, though showing correlated groups, were ignored since using other gene windows result in the inclusion of more gene groups that are correlated by chance.

**Figure 19.**
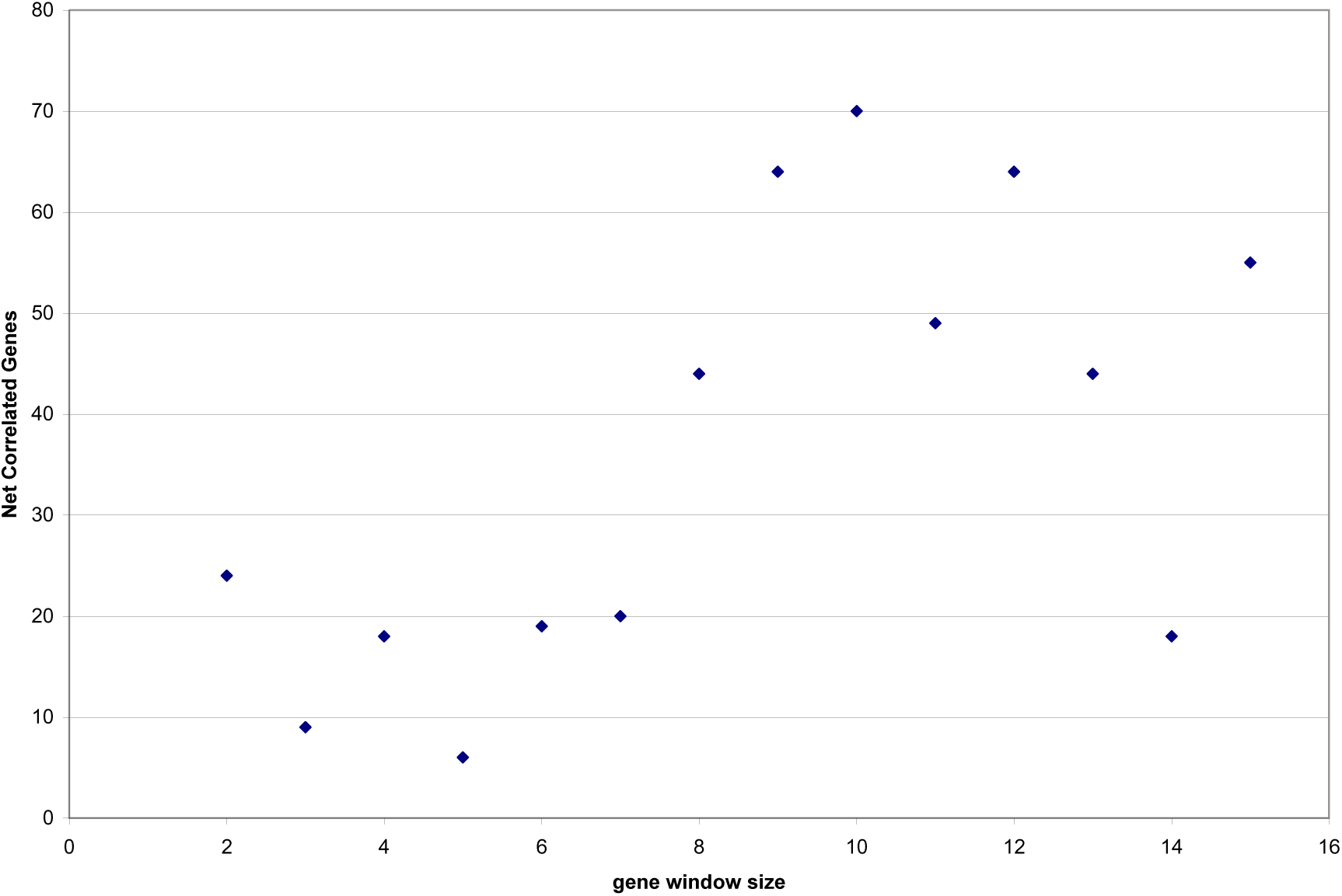
The net number of genes in a correlated group based on significance of correlation (p < 0.0001) at different gene window sizes for GR978-only experiments

There were either overlapping groups of the ten-adjacent genes of RCEs that are very near each other; these were collapsed as a single group with more than 10 genes, resulting in the detection of 175 gene members in 16 merged RCEs at p value <0.0005 (Appendix Table 1), with a minimum ten genes per group. Computation of the average size of the gene models and their distance from each other, for each group of adjacent genes consisting the 16 RCEs, gives an estimated average size of 286.9 kb per RCE; this is ∼2.87 times the size of the ten-gene, 100kb correlated window reported by Ma et al. (2005) when they profiled the genome-wide transcription in indica rice using a high density oligoarray. The frequency distribution of the average sizes of RCEs show that the most frequent correlated window size is ∼246 kb (Figure 20). The total size of all RCEs is ∼4.9 mb, which is ∼1.3% of the total genome. These results suggest that specific regions of the rice genome shows a correlated expression pattern during biotic stress response and is, therefore, likely organized into so-called co-expressed chromosomal regions of the resistance transcriptome.

**Figure 20.**
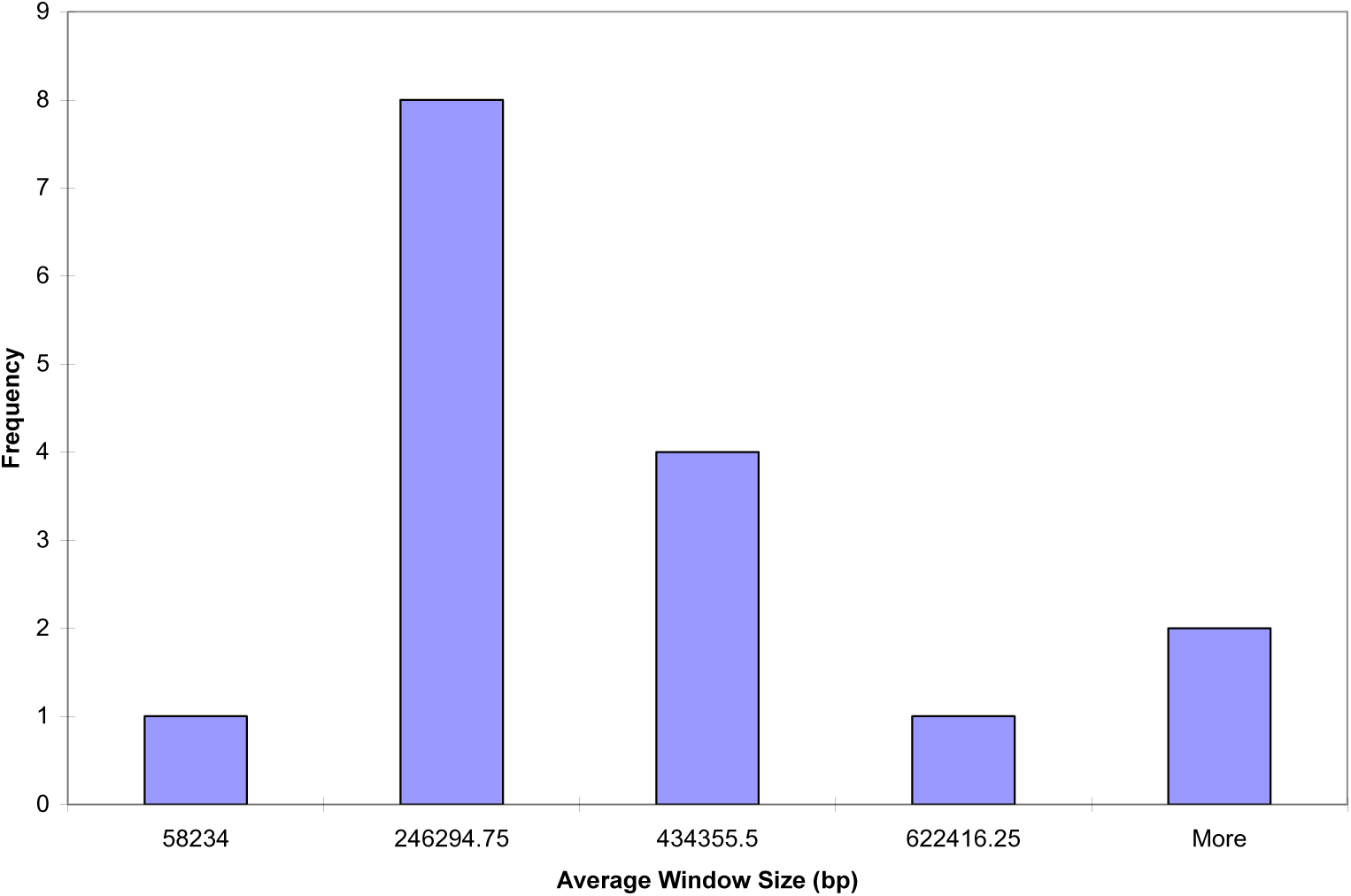
The distribution of the average sizes of RCEs in the GR978 resistance transcriptome.

In terms of correlation, Table 30 summarizes the count of correlated gene and gene groups in the GR978-only dataset.

**Table 30.** The gene and merged gene group counts of GR978 at a window of 10 genes

| SIGNIFICANCE<br>(P VALUE) | GENE COUNT<br>Ordered dataset | GENE COUNT<br>Randomized<br>dataset | GROUP COUNT<br>Ordered dataset | GROUP COUNT<br>Randomized<br>dataset |
| --- | --- | --- | --- | --- |
| 0.01 | 1,396 | 920 | 115 | 81 |
| 0.001 | 259 | 77 | 23 | 7 |
| 0.0005 | 175 | 37 | 16 | 3 |
| 0.0001 | 80 | 10 | 7 | 1 |

In order to compare resistance transcriptomes across genotypes, the same analysis was done with corresponding SHZ-2 experiments undergoing similar disease response. Figure 21 shows the gene window with the maximum net number of correlated genes for the SHZ-2 only experiments.

**Figure 21.**
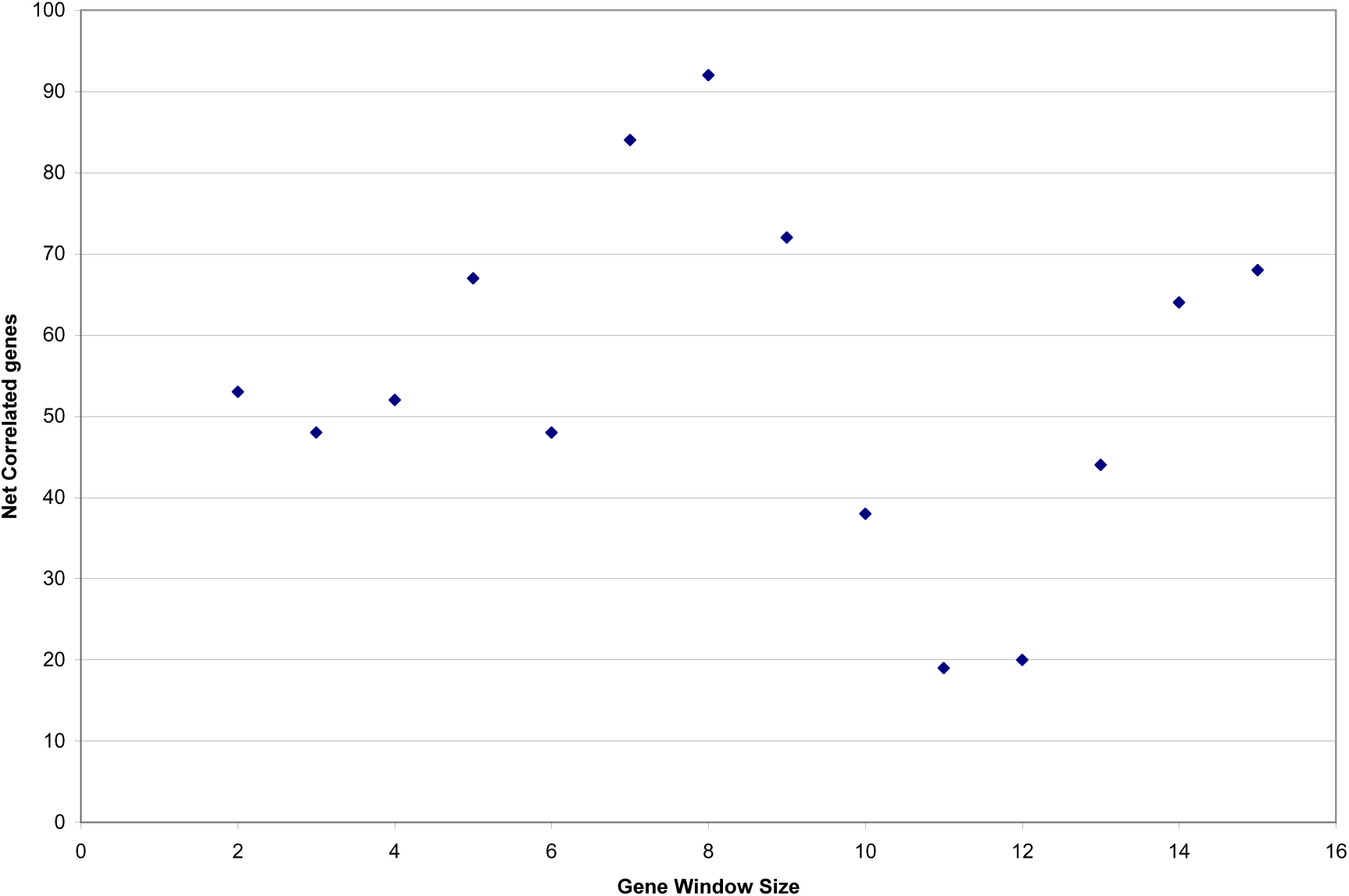
The net number of genes in a correlated group based on significance of correlation (p < 0.0005) at different gene window sizes for SHZ-2-only experiments

In the SHZ-2 only experiments, the maximum net number of correlated genes occurs at an 8-gene window. The counts of correlated genes and gene groups in the ordered and random SHZ-2 datasets are shown in Table 31.

**Table 31.** The gene and merged gene group counts of SHZ-2 at a window of 8 genes

| SIGNIFICANCE<br>(P VALUE) | GENE COUNT<br>Ordered dataset | GENE COUNT<br>Randomized<br>dataset | GROUP COUNT<br>Ordered dataset | GROUP COUNT<br>Randomized<br>dataset |
| --- | --- | --- | --- | --- |
| 0.01 | 940 | 534 | 101 | 58 |
| 0.001 | 184 | 79 | 21 | 9 |
| 0.0005 | 127 | 35 | 15 | 4 |
| 0.0001 | 33 | 16 | 4 | 2 |

There are 15 merged groups (at P<0.0005) with 127 gene members comprising the RCEs in SHZ-2 resistance transcriptome, having a computed average size of ∼295kb per 8-gene RCE, with a total size of ∼4.68 mb, which is similar to the GR978-only RCE total size.

To complete the RCE characterization of the resistance transcriptome, correlation of gene expression profiles were computed across resistant rice genotypes as well. Gene expression data from SHZ-2 were combined with the GR978 expression data obtained at the same time points (24h and 48h after blast infection) and analyzed using the same correlation method as before. At a p-value < 0.0005 level of significance of correlation, the net number of genes of correlated expression per window size shows an initial peak at the two gene window, which may be real but was not considered since two gene groups are less informative of the expression correlation structure of the genome; after the 2-gene peak, the graph of the net correlated gene count is similar in trend with the GR978-only experiment graph, with the highest net number of correlated genes in groups also occurring at the 10-gene window (Figure 22)

**Figure 22.**
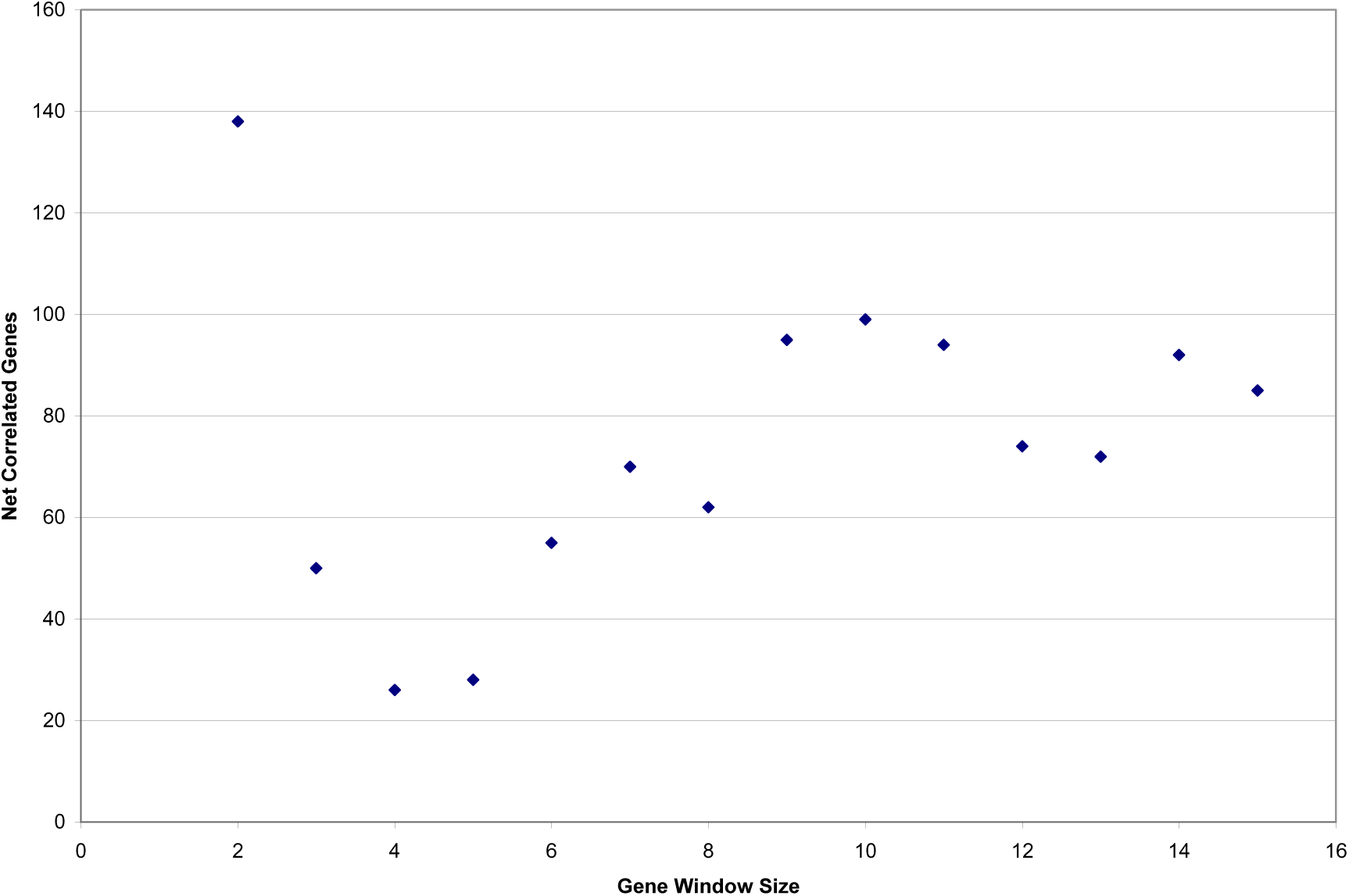
The net number of genes identified as grouped based on significance of correlation (p < 0.0005) at different gene window sizes for GR978-SHZ-2 regions of correlated expression

Table 32 shows the number of 10-gene groups of correlated expression from the ordered as well as the randomized data set. As with the GR978-only correlated region analysis, groups with overlapping genes were merged as one RCE. At p value cut-off < 0.0005, there were 156 genes in 14 RCEs.

**Table 32.** The number of 10 gene groups and the total number of merged genes in groups that are identified at various levels of significance (p values) for GR978-SHZ-2

| SIGNIFICANCE<br>(P VALUE) | GENE COUNT<br>Ordered dataset | GENE COUNT<br>Randomized dataset | GROUP COUNT<br>Ordered dataset | GROUP COUNT<br>Randomized<br>dataset |
| --- | --- | --- | --- | --- |
| 0.01 | 1,093 | 545 | 87 | 47 |
| 0.001 | 236 | 91 | 21 | 9 |
| 0.0005 | 156 | 57 | 14 | 5 |
| 0.0001 | 54 | 22 | 5 | 2 |

Using the 10-gene window of correlated expression, the estimated average size of a correlated window was approximately 332.7 kb per group for the 14 merged groups, and a total of 5.64 mb, which is bigger than the 287kb – 295kb average size range of RCE in GR978-only and SHZ-2 only experiments. These results show that the total counts of RCEs of the resistance transcriptome across the two disease-resistant rice genotypes are comparable.

A comparison of RCEs was made by comparing the genes comprising each RCE across genotypes; this reveals that in terms of the gene composing them, majority of the RCEs are distinct between GR978-only, GR978-SHZ-2 combined, and SHZ-2 only experiments. For the distinct RCEs, 155 out of 175 genes are unique to GR978 only, while 110 out of 127 genes are unique to SHZ-2 only. To determine if the distinct RCEs are different in general functions associated with each, GO gene category enrichment analysis of the member genes of the distinct GR978 RCEs and SHZ-2 RCEs was done and comparison of the GO categories in each list show that the enriched gene functions in the SHZ-2 RCEs are distinct from the enriched gene functions in GR978 RCEs (Tables 33 and 34).

**Table 33.** GO gene categories enriched in the GR978-only RCEs.

| GENE CATEGORY | P VALUE | GLOBAL<br>FDR P<br>VALUE | WITHIN-<br>SYSTEM<br>FDR P<br>VALUE |
| --- | --- | --- | --- |
| ribulose biphosphate carboxylase complex (sensu Magnoliophyta) | 0.0002 | 0.0000 | 0.0000 |
| carbon utilization by fixation of carbon dioxide | 0.0002 | 0.0000 | 0.0000 |
| ribulose-biphosphate carboxylase activity | 0.0002 | 0.0000 | 0.0000 |
| eukaryotic translation initiation factor 2B complex | 0.0013 | 0.0000 | 0.0000 |
| oxygen binding | 0.0013 | 0.0000 | 0.0000 |
| 3'-5'-exoribonuclease activity | 0.0021 | 0.0588 | 0.0556 |
| auxin:hydrogen symporter activity | 0.0038 | 0.0500 | 0.0476 |
| nitrogen compound metabolism | 0.0060 | 0.0870 | 0.0833 |
| acid phosphatase activity | 0.0068 | 0.0769 | 0.0741 |
| terpene metabolism | 0.0079 | 0.1030 | 0.1000 |
| nicotinamide-nucleotide adenyltransferase activity | 0.0079 | 0.0938 | 0.0909 |
| hexitol metabolism | 0.0079 | 0.0857 | 0.0833 |
| auxin polar transport | 0.0151 | 0.2630 | 0.2560 |
| response to copper ion | 0.0156 | 0.2440 | 0.2380 |
| CDP-diacylglycerol-inositol 3-phosphatidyltransferase activity | 0.0156 | 0.2270 | 0.2220 |
| proteasome regulatory particle (sensu Eukaryota) | 0.0156 | 0.2130 | 0.2080 |
| histone acetyltransferase activity | 0.0156 | 0.2000 | 0.1960 |
| copper chaperone activity | 0.0156 | 0.1890 | 0.1850 |
| ent-kaurene synthase activity | 0.0234 | 0.3570 | 0.3510 |
| glycine dehydrogenase complex (decarboxylating) | 0.0234 | 0.3390 | 0.3330 |
| 4-hydroxyphenylpyruvate dioxygenase activity | 0.0234 | 0.3230 | 0.3170 |
| nitrilase activity | 0.0234 | 0.3080 | 0.3030 |
| protein neddylation | 0.0234 | 0.2940 | 0.2900 |
| phosphatidylinositol biosynthesis | 0.0234 | 0.2820 | 0.2780 |
| two-component sensor molecule activity | 0.0311 | 0.3920 | 0.3870 |
| vitamin E biosynthesis | 0.0311 | 0.3770 | 0.3720 |
| phospholipase A2 activity | 0.0387 | 0.3750 | 0.3700 |
| protein tagging activity | 0.0387 | 0.3610 | 0.3570 |
| carotenoid biosynthesis | 0.0387 | 0.3490 | 0.3450 |
| phosphoserine phosphatase activity | 0.0387 | 0.3370 | 0.3330 |
| peroxisome organization and biogenesis | 0.0387 | 0.3260 | 0.3230 |
| cell surface (sensu Magnoliophyta) | 0.0387 | 0.3160 | 0.3130 |
| calcium ion binding | 0.0403 | 0.3060 | 0.3030 |
| glycine decarboxylation via glycine cleavage system | 0.0462 | 0.3760 | 0.3730 |
| glycine dehydrogenase (decarboxylating) activity | 0.0462 | 0.3650 | 0.3620 |

**Table 34.** GO gene categories enriched in the SHZ-2 only RCEs.

| GENE CATEGORY | P VALUE | GLOBAL<br>FDR P<br>VALUE | WITHIN-<br>SYSTEM<br>FDR P<br>VALUE |
| --- | --- | --- | --- |
| endoplasmic reticulum membrane | 0.0033 | 0.5000 | 0.3330 |
| regulation of phosphorylation | 0.0058 | 0.4000 | 0.3330 |
| neurotransmitter:sodium symporter activity | 0.0058 | 0.2500 | 0.2220 |
| physiological process | 0.0058 | 0.1820 | 0.1670 |
| phytoene dehydrogenase activity | 0.0058 | 0.1430 | 0.1330 |
| positive regulation of cell proliferation | 0.0115 | 0.3530 | 0.3330 |
| ferredoxin:thioredoxin reductase activity | 0.0115 | 0.3000 | 0.2860 |
| UDP-N-acetylgalactosamine biosynthesis | 0.0115 | 0.2610 | 0.2500 |
| ferredoxin reductase activity | 0.0115 | 0.2310 | 0.2220 |
| mediator complex | 0.0115 | 0.2070 | 0.2000 |
| water transport | 0.0115 | 0.1880 | 0.1820 |
| water channel activity | 0.0121 | 0.1710 | 0.1670 |
| rhodopsin-like receptor activity | 0.0137 | 0.1840 | 0.1790 |
| response to water deprivation | 0.0146 | 0.1950 | 0.1900 |
| calcium ion transporter activity | 0.0172 | 0.2950 | 0.2890 |
| dihydropteroate synthase activity | 0.0172 | 0.2770 | 0.2710 |
| nucleotide phosphatase activity | 0.0172 | 0.2600 | 0.2550 |
| phosphoglucomutase activity | 0.0172 | 0.2450 | 0.2410 |
| glycine cleavage complex | 0.0229 | 0.3570 | 0.3510 |
| photosynthesis, light reaction | 0.0229 | 0.3390 | 0.3330 |
| adenine salvage | 0.0285 | 0.3230 | 0.3170 |
| adenine phosphoribosyltransferase activity | 0.0285 | 0.3080 | 0.3030 |
| hydrogen-translocating pyrophosphatase activity | 0.0285 | 0.2940 | 0.2900 |
| beta-glucuronidase activity | 0.0285 | 0.2820 | 0.2780 |
| glycine catabolism | 0.0285 | 0.2700 | 0.2670 |
| calcium-dependent protein serine/threonine kinase activity | 0.0285 | 0.2600 | 0.2560 |
| phosphatidate phosphatase activity | 0.0342 | 0.3250 | 0.3210 |
| beta-amylase activity | 0.0342 | 0.3130 | 0.3100 |
| polysaccharide catabolism | 0.0342 | 0.3020 | 0.2990 |
| response to abiotic stimulus | 0.0342 | 0.2920 | 0.2890 |
| Microsporogenesis | 0.0342 | 0.2830 | 0.2800 |
| glycine dehydrogenase (decarboxylating) activity | 0.0342 | 0.2740 | 0.2710 |
| regulation of stomatal movement | 0.0397 | 0.3270 | 0.3230 |
| intramolecular transferase activity, phosphotransferases | 0.0397 | 0.3170 | 0.3140 |
| ion channel activity | 0.0397 | 0.3080 | 0.3050 |
| GTPase activator activity | 0.0397 | 0.2990 | 0.2960 |
| abscisic acid mediated signaling | 0.0453 | 0.3270 | 0.3240 |

The non-overlap of the enriched gene categories between the genotype-specific RCEs imply that different gene functions are attributed to the distinct RCEs that were determined for each genotype.

Only one RCE found in chromosome 11 is common across GR978 and SHZ-2 (Figure 23). This RCE is particularly interesting due to the function annotation of disease resistance (RPR1, NBS-LRR) for two genes in this region, as well as the enriched gene categories in this gene subset that recapitulates disease resistance response events accurately (Tables 35 and 36). Moreover, no DEGs from either GR978 or SHZ-2 experiments co-localize with this RCE.

**Figure 23.**
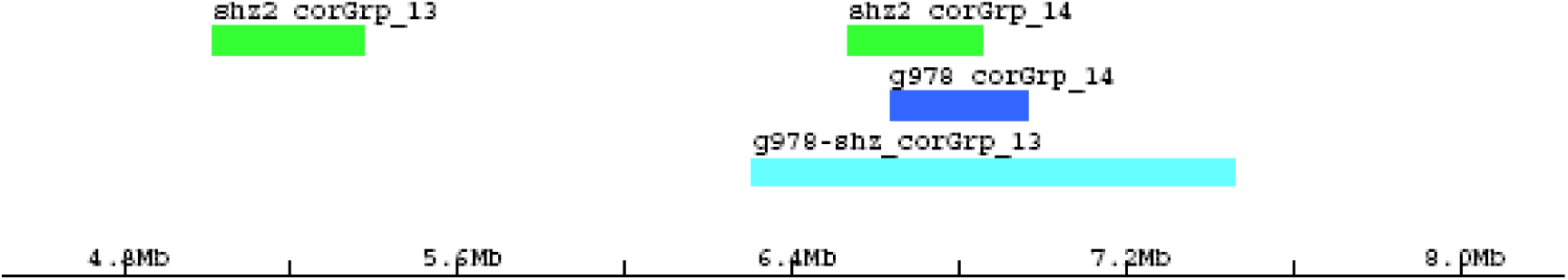
The co-localization in chromosome 11 of RCEs determined from GR978 only, GR978-SHZ-2 combined, and SHZ-2 only expression data sets

**Table 35.**
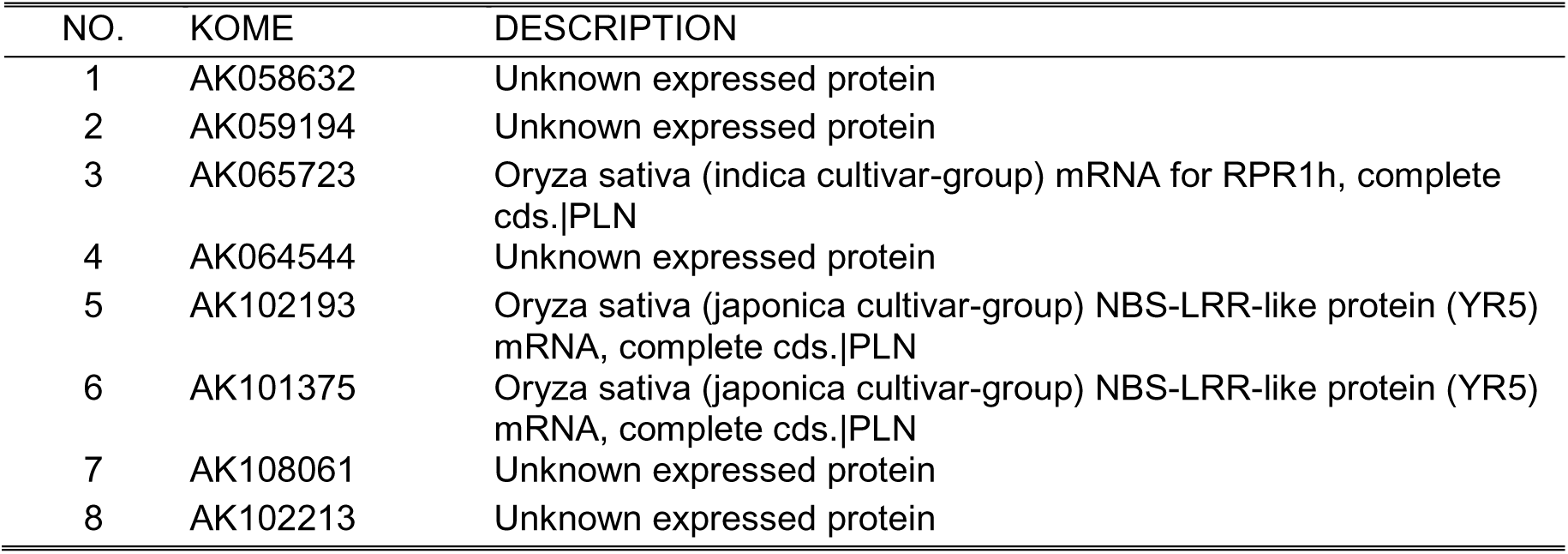
Eight genes in the GR978 - SHZ-2 common RCE found at chromosome 11 (6,637,634 bp to 6,854,148 bp)

**Table 36.** Enriched GO gene categories in the GR978 - SHZ-2 common RCE found at chromosome 11 (6,637kb to 6,854 kb)

| GENE CATEGORY | P VALUE | GLOBAL<br>FDR P<br>VALUE | WITHIN-<br>SYSTEM<br>FDR P<br>VALUE |
| --- | --- | --- | --- |
| extrinsic to plasma membrane | 0.0028 | 0.5000 | 0.3330 |
| hypersensitive response | 0.0044 | 0.4000 | 0.3330 |
| nucleotide binding | 0.0113 | 0.6250 | 0.5560 |
| defense response | 0.0132 | 0.4550 | 0.4170 |
| protein binding | 0.0329 | 0.6430 | 0.6000 |

#### Are differentially expressed genes within and across genotypes enriched within the regions of correlated gene expression?

In order to infer more possible functions associated with the RCEs detected during blast infection of GR978, SHZ-2, and in common between GR978 and SHZ-2, DEGs from GR978-only experiments (GR978-DEG), SHZ-2-only experiments, and the common DEGs from GR978 and SHZ-2 (GR978-SHZ-2 DEG) were tested for association with these respective correlated regions. In order to establish association and enrichment, the test for fixed ratio using Fisher exact test is employed extensively, as described in the methodology section. Table 37 summarizes the counts of features that were used for testing for enrichment of DEGs within RCEs.

**Table 37.** Count of DEGs that bin against the RCE specified in the three genotypes studied

| GENE GROUPS | BINNING WITHIN THE REGIONS OF |  |  | Global |
| --- | --- | --- | --- | --- |
|  | GR978<br>RCE | GR978-SHZ2<br>RCE | SHZ2<br>RCE |  |
| GR978 DEG | 2 | 6 | 8 | 797 |
| GR978-SHZ2<br>DEG | 0 | 2 | 1 | 89 |
| SHZ DEG | 46 | 45 | 35 | 4,326 |
| ALL GENES | 175 | 156 | 127 | 20,870 |

To test if DEGs associate with RCEs, each DEG group was tested for enrichment against each RCE as shown in Tables 38 to 40. For each study, the test for fixed ratio was between the actual DEG count within:outside ratio against the expected (adjusted) DEG count within:outside ratio. For all DEGs (GR978-DEGs, GR978-SHZ2 DEGs, and SHZ2-DEGs) tested for association against all RCEs (GR978, GR978-SHZ2, SHZ2 RCEs), all Fisher exact test results were not significant (p > 0.05); for all nine DEG-RCE comparisons (the actual to expected ratio of DEGs to non-DEGs was not significantly different. DEGs are not enriched within RCE regions. The same association analysis was repeated on a per chromosome basis for the same DEG-RCE comparisons, with the same non-significant results for all chromosomes (Appendix tables 5 to 7).

**Table 38.**
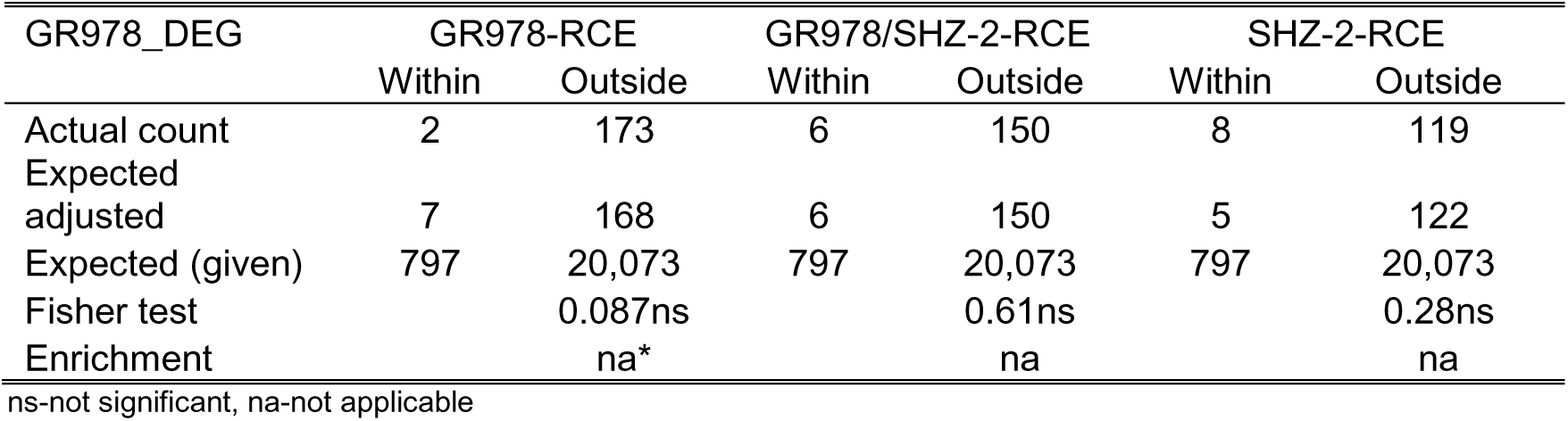
Association of GR978-specific DEGs to all RCEs defined by each genotype and genotype combination

| GR978_DEG | GR978-RCE |  | GR978/SHZ-2-RCE |  | SHZ-2-RCE |  |
| --- | --- | --- | --- | --- | --- | --- |
|  | Within | Outside | Within | Outside | Within | Outside |
| Actual count | 2 | 173 | 6 | 150 | 8 | 119 |
| Expected adjusted | 7 | 168 | 6 | 150 | 5 | 122 |
| Expected (given) | 797 | 20,073 | 797 | 20,073 | 797 | 20,073 |
| Fisher test |  | 0.087ns |  | 0.61ns |  | 0.28ns |
| Enrichment |  | na* |  | na |  | na |

ns-not significant, na-not applicable

**Table 39.**
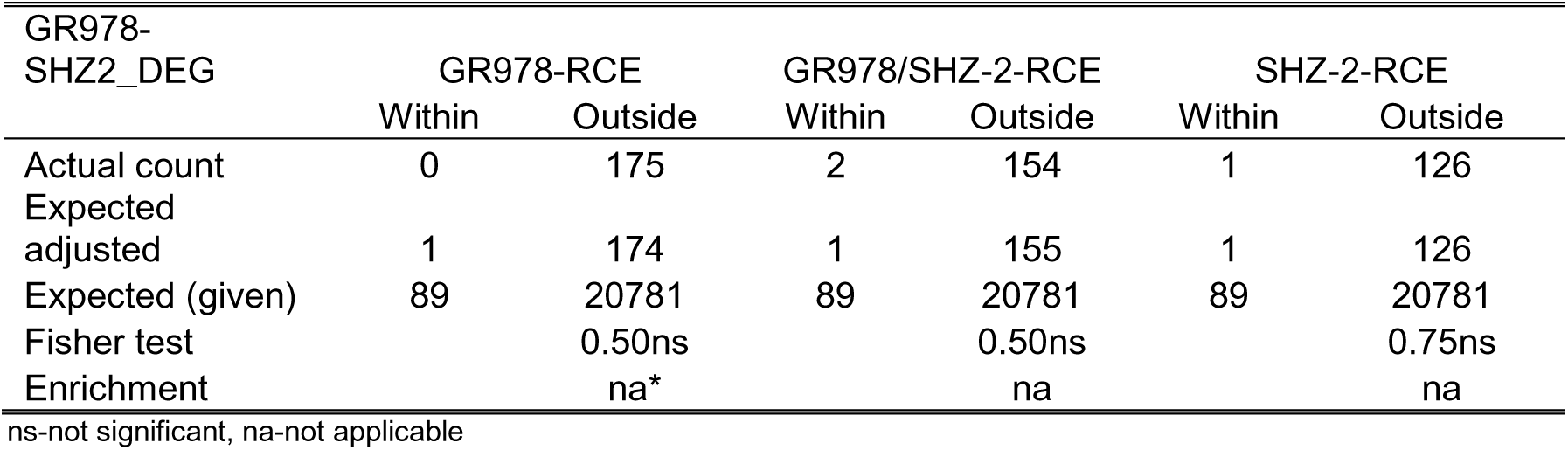
Association of GR978-SHZ2 DEGs to all RCEs defined by each genotype and genotype combination

**Table 40.** Association of SHZ2 DEGs to all RCEs defined by each genotype and genotype combination

| SHZ2_DEG | GR978-RCE |  | GR978/SHZ-2-RCE |  | SHZ-2-RCE |  |
| --- | --- | --- | --- | --- | --- | --- |
|  | Within | Outside | Within | Outside | Within | Outside |
| Actual count | 46 | 129 | 45 | 111 | 35 | 92 |
| Expected adjusted | 36 | 139 | 32 | 124 | 26 | 101 |
| Expected (given) | 4326 | 16544 | 4326 | 16544 | 4326 | 16544 |
| Fisher test |  | 0.13ns |  | 0.057ns |  | 0.12ns |
| Enrichment |  | na* |  | na |  | na |

ns-not significant, na-not applicable

#### The association of DEGs and RCEs to published rice blast resistance QTL regions

Biological evidence such as QTL location data from previous blast resistance studies might be helpful in characterizing the resistance transcriptome: for this third level of characterization of the resistance transcriptome, we hypothesize that disease response-specific gene expression are co-localized within known QTL regions. To test this hypothesis, the association of the resistance transcriptome with regions of QTLs controlling blast disease resistance was examined. The physical locations of the 40 Bl-QTLs (as described in the methodology section) were used as bins for co-localizing the resistance transcriptome. The spatial distribution in the genome of all DEGs, RCEs from all the studies analyzed, and all Bl-QTLs are shown in Figures 24 to 27. The count of genes from each study that were used in association analysis is given in Table 41.

**Figure 24.**
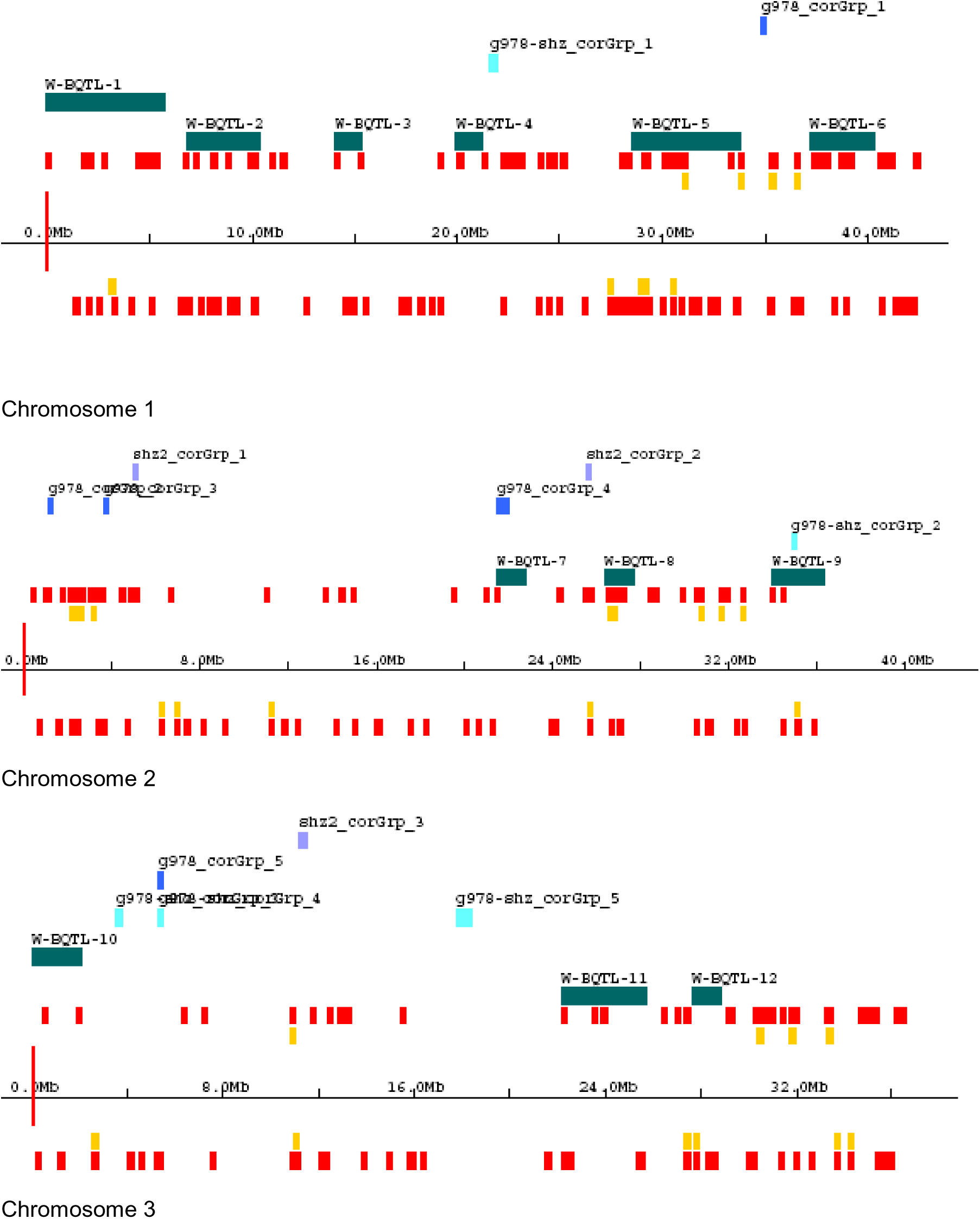
Distribution along the genome of Bl-QTLs, RCEs, and DEGs for all genotypes studied (chromosomes 1 – 3). Orange blocks are GR978 DEGs, yellow blocks are GR978-SHZ DEGs

**Figure 25.**
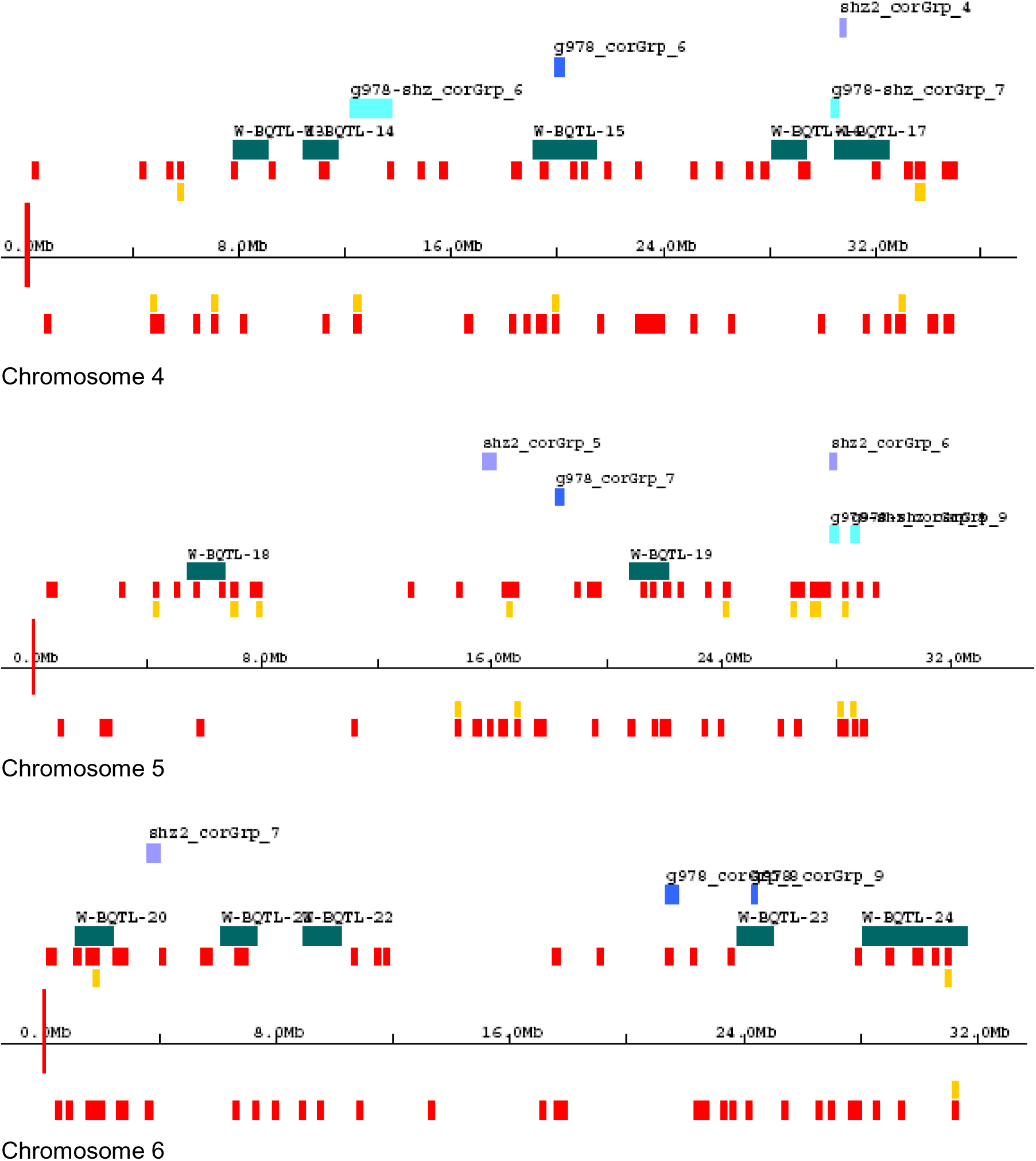
Distribution along the genome of Bl-QTLs, RCEs, and DEGs for all genotypes studied (chromosomes 4 – 6). Orange blocks are GR978 DEGs, yellow blocks are GR978-SHZ DEGs, cont’d..

**Figure 26.**
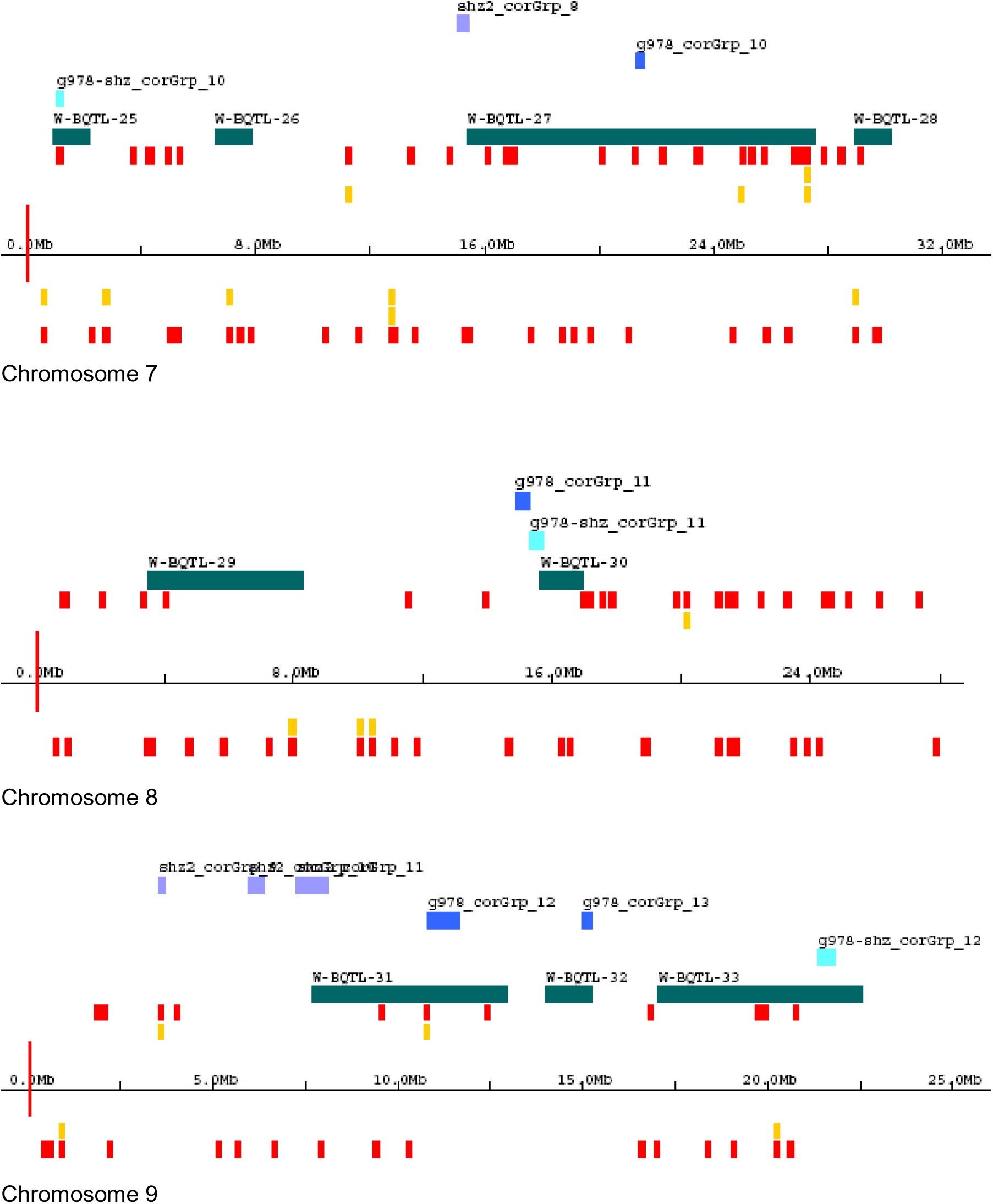
Distribution along the genome of Bl-QTLs, RCEs, and DEGs for all genotypes studied (chromosomes 7 – 9). Orange blocks are GR978 DEGs, yellow blocks are GR978-SHZ DEGs, contd

**Figure 27.**
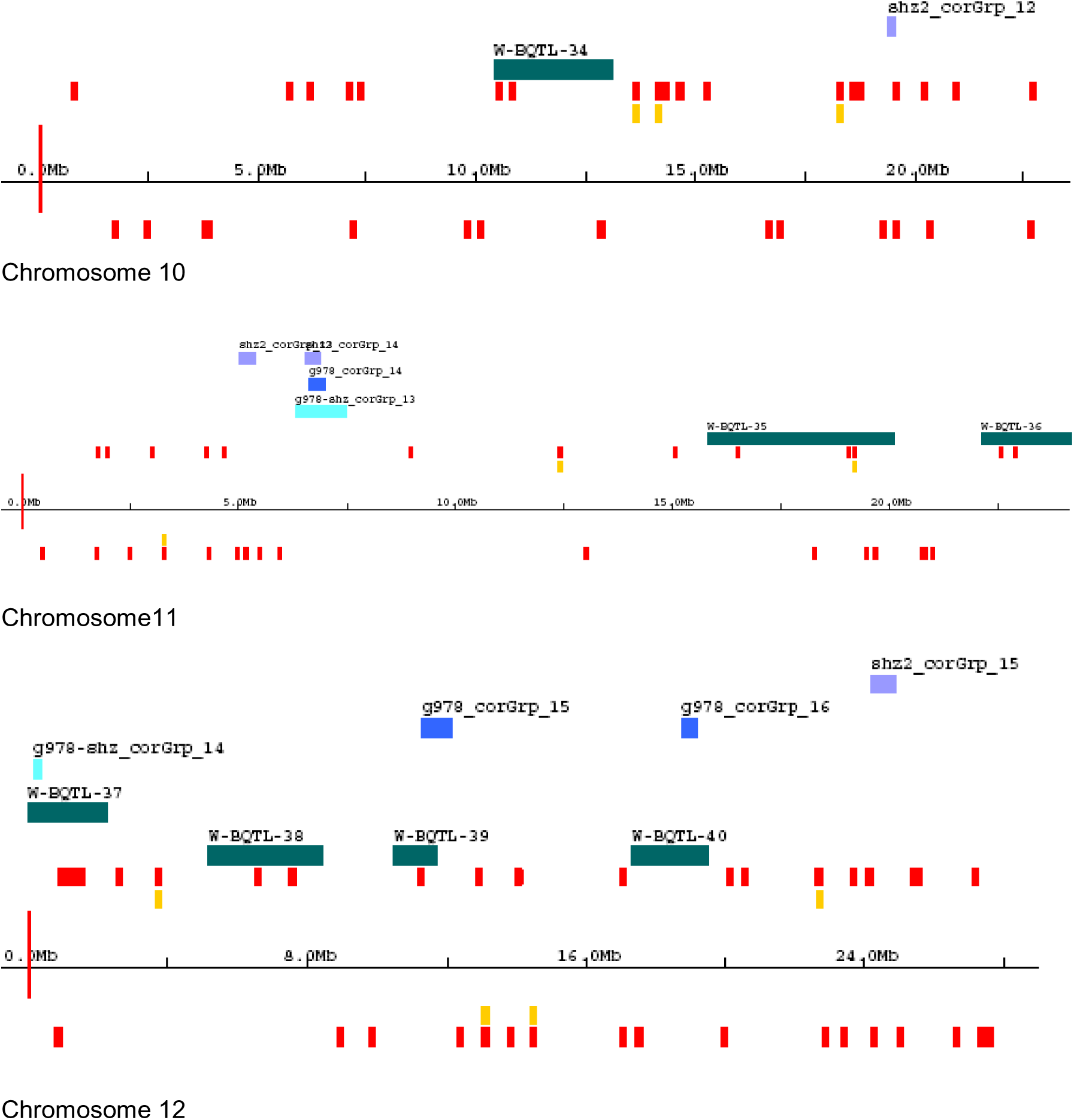
Distribution along the genome of Bl-QTLs, RCEs, and DEGs for all genotypes studied (chromosomes 9 – 12). Orange blocks are GR978 DEGs, yellow blocks are GR978-SHZ DEGs

**Table 41.** Count of DEGs and RCEs binning within Bl-QTLs for the 3 studies analyzed

| GENE SET | BINNING WITHIN |  |
| --- | --- | --- |
|  | BL-QTL | Global Count |
| GR978 DEG | 226 | 797 |
| GR978-SHZ2 DEG | 21 | 89 |
| SHZ DEG | 1,237 | 4,326 |
| GR978-RCE | 84 | 175 |
| GR978-SHZ2-RCE | 43 | 156 |
| SHZ2-RCE | 16 | 127 |
| All Genes | 6,187 | 20,870 |

Using the enrichment analysis method described in the previous section, the association of DEGs and RCEs from each study with the Bl-QTL regions was computed in the following tables (Table 42 to 43). For the association of DEGs with Bl-QTLs, for each study, the ratio of actual (within Bl-QTL : outside Bl-QTL) was tested against the expected (within Bl-QTL : outside Bl-QTL) ratio by Fisher exact test. Repeating the same association analysis on a per chromosome basis for the same DEG-Bl-QTL comparisons yielded the same non-significant results (Appendix table 8).

**Table 42.** Association of DEGs computed from GR978, SHZ2, and common GR978-SHZ2 with Bl-QTL regions

|  | GR978-DEG |  | GR978/SHZ-2_DEG |  | SHZ-2-DEG |  |
| --- | --- | --- | --- | --- | --- | --- |
|  | Within BL-QTL | Outside BL-QTL | Within BL-QTL | Outside BL-QTL | Within BL-QTL | Outside BL-QTL |
| Actual count | 226 | 5961 | 21 | 6166 | 1237 | 4950 |
| Expected adjusted | 236 | 5951 | 26 | 6161 | 1282 | 4905 |
| Expected (given) | 797 | 20,073 | 89 | 20,781 | 4,326 | 16,544 |
| Fisher test |  | 0.335ns |  | 0.280ns |  | 0.163ns |
| Enrichment |  | na* |  | na |  | na |

ns-not significant, na-not applicable

All DEGs did not associate with Bl-QTL regions, as shown by the Fisher exact test (Table 42). There is no enrichment of DEGs within the Bl-QTL regions.

The same tests to analyze for the association of RCE to Bl-QTLs were done separately for each study. For the RCEs, there is significant association of GR978 RGCEs with Bl-QTLs, with enrichment (Table 43). The common GR978-SHZ2 RGCEs are not associated with Bl-QTLs. In the case of SHZ-RCE, although there was significant difference in the ratio of SHZ-RCE:non-RCE within the Bl-QTL region as compared to the global ratio, enrichment value was less than 1.0, thereby it was considered as non-enrichment.

**Table 43.**
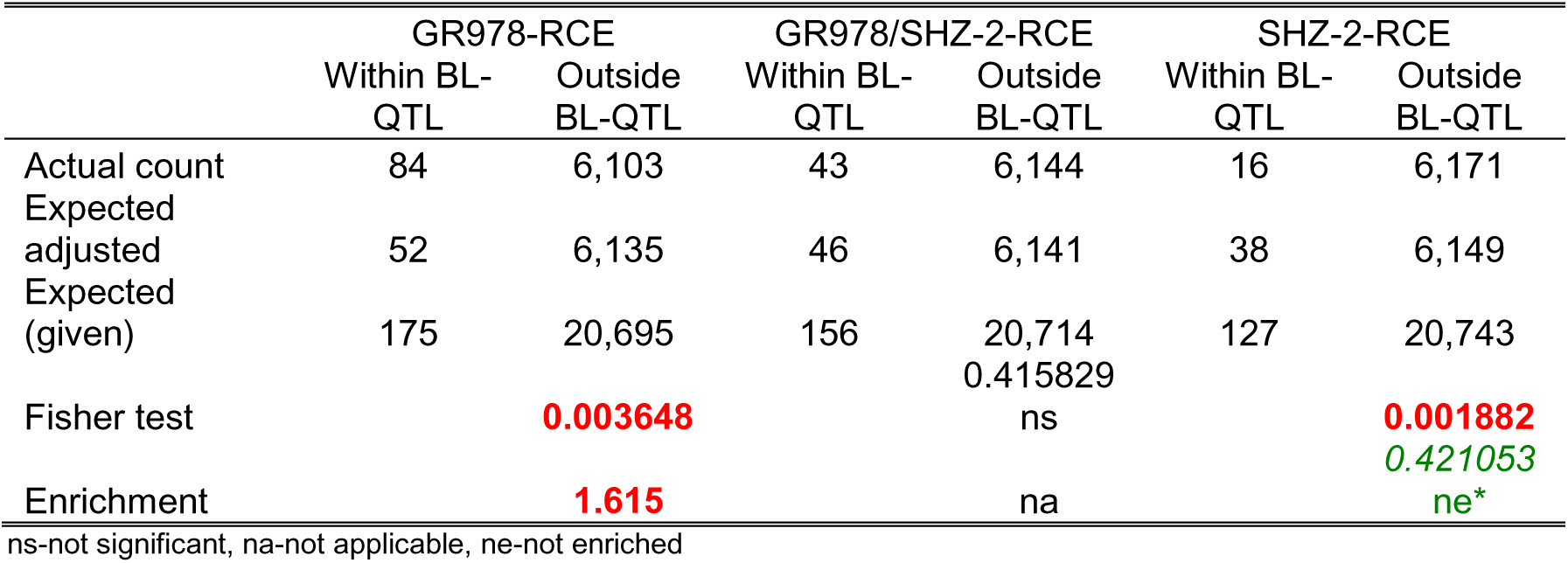
Association of RCE in the 2 genotypes analyzed with Bl-QTL regions

The GR978-RCEs associated with Bl-QTLs are listed on Table 44, showing 8 Bl-QTLs in chromosomes 2, 4, 6, 7, 9, and 12 aligning with 8 GR978-RCEs. There are 84 interesting genes within these aligned regions (Appendix Table 4). None of the 84 genes are DEGs.

**Table 44.** Alignment of GR978-RCEs with Bl-QTLs

| CHROMOSOME | GR978 RCE GROUP NUMBER | BL-QTL NUMBER |
| --- | --- | --- |
| 2 | 4 | 7 |
| 4 | 6 | 15 |
| 6 | 9 | 23 |
| 7 | 10 | 27 |
| 9 | 12 | 31 |
| 9 | 13 | 32 |
| 12 | 15 | 39 |
| 12 | 16 | 40 |

The GO gene categories enriched for these 84 genes show interesting resistance-response themes (Table 45).

**Table 45.** Enriched GO gene categories in the RCEs enriched in Bl-QTL regions.

| GENE CATEGORY | P VALUE | GLOBAL<br>FDR P<br>VALUE | WITHIN-<br>SYSTEM<br>FDR P<br>VALUE |
| --- | --- | --- | --- |
| oxygen binding | 0.0000 | 0.0000 | 0.0000 |
| ribulose biphosphate carboxylase complex (sensu Magnoliophyta) | 0.0001 | 0.0000 | 0.0000 |
| carbon utilization by fixation of carbon dioxide | 0.0001 | 0.0000 | 0.0000 |
| ribulose-bisphosphate carboxylase activity | 0.0001 | 0.0000 | 0.0000 |
| eukaryotic translation initiation factor 2B complex | 0.0004 | 0.0000 | 0.0000 |
| electron transport | 0.0038 | 0.0588 | 0.0556 |
| nicotinamide-nucleotide adenyltransferase activity | 0.0042 | 0.0500 | 0.0476 |
| histone acetyltransferase activity | 0.0084 | 0.1740 | 0.1670 |
| proteasome regulatory particle (sensu Eukaryota) | 0.0084 | 0.1540 | 0.1480 |
| copper chaperone activity | 0.0084 | 0.1380 | 0.1330 |
| response to copper ion | 0.0084 | 0.1250 | 0.1210 |
| ent-kaurene synthase activity | 0.0126 | 0.2570 | 0.2500 |
| protein neddylation | 0.0126 | 0.2370 | 0.2310 |
| glycine dehydrogenase complex (decarboxylating) | 0.0126 | 0.2200 | 0.2140 |
| nitrilase activity | 0.0126 | 0.2050 | 0.2000 |
| two-component sensor molecule activity | 0.0168 | 0.2340 | 0.2290 |
| translational initiation | 0.0175 | 0.3000 | 0.2940 |
| phosphoserine phosphatase activity | 0.0210 | 0.2830 | 0.2780 |
| phospholipase A2 activity | 0.0210 | 0.2680 | 0.2630 |
| protein tagging activity | 0.0210 | 0.2540 | 0.2500 |
| glycine dehydrogenase (decarboxylating) activity | 0.0251 | 0.3230 | 0.3170 |
| glycine decarboxylation via glycine cleavage system | 0.0251 | 0.3080 | 0.3030 |
| cell recognition | 0.0292 | 0.3530 | 0.3480 |
| hydrolase activity, acting on carbon-nitrogen (but not peptide) bonds | 0.0292 | 0.3380 | 0.3330 |
| lipid catabolism | 0.0333 | 0.3240 | 0.3200 |
| signal transducer activity | 0.0335 | 0.3510 | 0.3460 |
| translation initiation factor activity | 0.0372 | 0.3380 | 0.3330 |
| defense response to pathogen, incompatible interaction | 0.0374 | 0.3250 | 0.3210 |
| glycine metabolism | 0.0415 | 0.3490 | 0.3450 |
| organic anion transport | 0.0456 | 0.3820 | 0.3780 |
| response to pathogenic bacteria | 0.0496 | 0.3800 | 0.3760 |

### Discussion

#### Common DEGs across genotypes undergoing the resistance response

Many of the defense-responsive genes detected in the GR978 experiments were found in common with the DE genes observed in SHZ-2. Hierarchical clustering of the expression profiles of the common DEGs using the merged SHZ-2 and GR978 data show that the clusters are different as compared to the GR978 only experiment; this may be due to the inherent difference in resistance mechanisms between the two genotypes. By visual inspection of the genes in the CDE clusters, there is, as in the case of DEGs from GR978-only experiments, the expression of genes involved in early resistance engagement and HR/PCD, as well as general resistance genes. Looking at the enriched gene categories in the common CDE genes show the enrichment of resistance responsive, as well as non-resistance related gene categories, indicating that the gene expression experiments for each respective variety captures the variety-specific differential expression of genes involved in general metabolic processes aside from the resistance-specific expression.

The list of common DI genes appears more related to resistance-specific genes. It implies the conservation of many inducible genes from each variety during blast disease response, and looking at the results of GO gene category enrichment analysis, it shows abundance of general resistance response themes observed during early resistance engagement which is similar to that observed in DI genes from GR978-only experiments. Some abiotic stress-related genes are also found expressed such as cold acclimation protein WCOR13-like (AK066036) and DRE binding factor (AK108830), indicating possible intersection of these genes with the biotic stress response pathway. It is likely that these common induced genes are the ones effecting the resistance phenotype but are not the causal genes for the resistance phenotype in GR978 and SHZ-2.

The treatment of Nipponbarre with BTH induces the systemic activation of SAR pathway. It is interesting to note that while most of the differential expression occurs at 24h in Nipponbarre, most of the corresponding gene induction responses occur at 48h in GR978. Another intriguing observation is that the common defense response genes are mostly down-expressed in Nipponbarre-BTH treated, which is paradoxical for the SAR response; no clear pattern of expression of the common DE genes can be discerned between GR978 and Nipponbarre-BTH during resistance response. HCL could not be utilized for clustering co-expressed genes since it is not possible to determine the constitutively different and induced genes from the Nipponbarre DE gene set. Generally, the level of gene expression is higher in GR978 than Nipponbarre for the common DE genes, but the down-expression of many resistance-related genes in GR978 is opposite of what is expected during resistance response. Further characterization of the transcripts in GR978 for these genes using other experimental methods may provide a better insight as to the role of these common DEGs in the resistance response of GR978.

In general, characterizing the resistance transcriptome using individual DEGs does not appear to be informative in providing a list of candidate genes with strong evidence supporting direct involvement of this gene list with the resistance phenotype.

#### Regions of the genome showing correlated expression during the resistance response

Taking the expression profile of mutant GR978 and resistant variety SHZ-2 undergoing disease response, around 15 groups of adjacent genes that are expressed together are found in the resistance transcriptome, with 8 to 16 genes comprising each group. The average sizes of the RCEs of ∼290kb are almost three times larger than the correlated regions reported by Ma et al. (2005) using a 70-mer oligo set representing ∼41,000 rice gene models. This computed size may be the consequence of lower gene density in the 22K array system used; in the set of ∼21,000 spotted genes, the distribution of these genes along the genome are not random, and many of the detected RCEs may in fact be two or more RCEs in close proximity. The sizes of ten-gene RCEs actually vary from 58kb to 800kb as computed based on the position of the KOME genes in the TIGR rice pseudomolecule release 3.0. The use of a higher density array will probably resolve the sizes of the RCEs more accurately.

Most of the RCEs determined from GR978-only, SHZ-only, and GR978-SHZ combined data do not co-localize with each other, and these distinct RCEs most likely represent different sets of functions, as shown by GO gene category enrichment analysis. There is, however, the exception of the common RCE found in chromosome 11. In this group are three genes that are annotated as disease resistance genes, as well as five other genes of unknown function; the correlated expression of the 5 unknown genes in this region with the 3 R genes strongly implies the involvement of these unknown genes with the resistance response. A naturally resistant cultivar such as SHZ-2 most likely utilizes these types of R genes for its resistant phenotype. Intuitively, the gain of resistance in a mutant (the wildtype IR64 having no known active blast-resistance gene) should be of a different mechanism; it is likely caused by de-regulation or alteration of signaling that triggers basal defense-related genes, as the case of the *spl11* lesion mimic mutant. However, the correlated expression this common gene group between GR978 and SHZ-2 suggests the involvement of disease-R genes in this region with the resistance responses of GR978 and SHZ-2. It is interesting to pursue experiments to confirm if this region contributes significantly to the resistance phenotype of SHZ-2 and GR978.

The determination and comparison of RCEs within and across genotypes is a novel way of mining microarray expression data to uncover potential candidate genes for blast resistance in regions that would otherwise be declared as consisting of not-significantly expressed genes if standard methods (ANOVA, t-test) for establishing significance-of-expression were employed.

#### Are regions of correlated expression associated with DEGs?

Although gene expression experiments involving microarrays are able to give information on the behavior of many genes all at once, the ultimate goal of finding which gene(s) are responsible for the phenotype under study remains elusive. In this experiment, while many genes were detected as differentially expressed only during the resistance response, selecting candidates by virtue of differential expression alone is not enough to guarantee successful candidate gene selection. We tried to explore the possibility that the more interesting DEGs are enriched within RCEs, assuming that DEGs responsible for the phenotype under study should also be co-regulated along with other, more subtly expressed genes. Association analysis show that DEGs consistently failed to significantly associate with RCEs, suggesting that condition-specific DEGs are not necessarily located within regions of correlated expression, and are probably under a different type of regulatory control than RCEs. This does not mean that the two sets of genes (DEGs and RCEs) are not related to each other in terms of controlling the resistance phenotype. RCEs are interesting since the concept of region-specific co-expression agrees nicely with results of genetic studies implicating genetic regions (such as QTLs) with quantitative resistance phenotypes.

The occurrence of DEGs all over the genome also suggests the complexity of the resistance mechanism: while numerous studies already clearly explain the events from early resistance engagement until the onset of long-lasting systemic resistance, identifying the specific genes in the rice genome that are directly involved in these events remains a daunting task. From this experiment we see so many instances of chitinases, cytochromes, peroxidases, and other resistance-responsive genes that are all differentially expressed. We need to explore more robust analytical approaches to microarray analysis in order to sift through the noise of general non-resistance related expression as well as the non-genotype specific common basal defense responses in order to find the most relevant gene(s) that ultimately cause the resistance phenotype.

#### Association of known blast resistance QTLs with DEGs and RCEs

The significant association of GR978-specific regions of correlated expression to blastR QTL meta-regions is very interesting; it provides another inference to the biological relevance of condition-specific and genotype-specific correlated expression of adjacent gene groups, in this case in the context of disease resistance response of GR978 to rice blast. From these observations, a working hypothesis is that there is genotype-specific correlated expression of adjacent genes during defense response localizing to known blast resistance QTL regions. A caveat to this method is that it is highly dependent on the quality of annotation behind QTL regions. QTL locations are notoriously imprecise due to the translation of genetic distances and locations into physical coordinates on the reference genome; translated QTL intervals often cover a big portion of the genome such that the probability of significant association is higher than expected, leading to false positive associations. Another weakness of the QTL association method is that all QTLs are equal in the genome: a QTL with a weak phenotypic effect appears the same as a QTL with a strong effect, and association tests do not care if the QTL region has a strong or weak phenotypic effect. This case is similar to the false positive scenario discussed before.

Looking at the GR978 experiment results, it is inferred that there is genotype-specific correlated expression of adjacent genes during defense response that localizes to known blast resistance QTL regions. Although GR978 is a gain-of resistance mutant and the likely cause of quantitative resistance may be entirely different from the multi-genic quantitative resistance expressed by field-resistant rice cultivars, the conservation of basal defense mechanisms as seen from hierarchical clustering results of GR978-SHZ-2 common DEGs indicates that similar expression events may be happening in naturally disease-resistant cultivars.

These findings have implications in QTL cloning studies because it puts forward a hypothesis that QTLs are regions of adjacent co-expressed genes in the genome and not single gene with quantitative effects. In order to obtain the phenotype associated with QTLs, regions of 100-300 kbs might be needed. Marker-assisted selection for regions of correlated expression may be a useful approach to maintain the gene complexes together.

These results highlight the fact that selecting candidate genes by filtering for differential expression is not sufficient anymore for microarray experiment results. A case in point in this experiment is the identification of a candidate region with 8 interesting genes in chromosome 11 where there are no known QTLs, and as further shown, overlaying biologically relevant information such as QTL positions, with the correlation structure of gene expression data led to the identification of 84 genes that are otherwise missed by significance analysis since genes in this region were not declared as differentially expressed. We put forward the idea that in the selection of candidate genes for resistance characterization or for breeding purposes from microarray-based gene expression experiments, it is necessary to establish the correlation structure of the expression data set within the genotype. The subsequent selection of good candidate genes for a trait of interest should be made by comparing genotype-specific RCEs across the genotypes of interest. Informatically, we propose to analyze more public gene expression data sets involving disease or biotic stress in order to find if the same or other RCEs are associated with disease resistance QTLs. Experimentally, we can focus on these common RCEs from a breeding perspective: genotyping mapping populations with markers flanking common RCEs and determining phenotype association with these RCEs. Since the regions of concern are relatively large (100kb to 300kb), a marker-based approach is practical and inexpensive.

## GENERAL DISCUSSION

The development and use of a specialized mutant phenotype CV enables the accurate and unambiguous characterization of the IR64 mutant collection at IRRI. Furthermore, through the use of the CV, gene function discovery is possible by linking information from other mutant collections that are well characterized for gene mutations, with the IR64 mutant collection, which is mostly characterized for phenotype. One use case is in reverse genetics: a researcher might be interested in a particular gene characterized in Tos 17 mutants and would like to study the same gene in an indica variety such as IR64. By querying the Tos17 database for the gene sequence, the corresponding phenotype description for the gene mutation can be retrieved from the same database, and if this phenotype is mapped by the CV to the IR64 mutant database, the corresponding IR64 mutants with the same phenotypes can be selected and these mutants might have the same gene functions altered and therefore can be further characterized. The use of mutant phenotype CV by various rice mutant resources can enrich information contained within each database by enabling cross-queries implemented seamlessly via web services.

Focusing on a particular mutant from the IRRI collection, GR978 was selected for its gain of broad-spectrum disease resistance; the gene expression profile of this genotype was characterized during blast-resistance response. The disease resistance transcriptome of GR978 is composed of sets of differentially expressed genes located within and outside the genetically mapped broad spectrum resistance gene of GR978, as well as of adjacent gene groups showing correlated expression, where most of the individual gene functions and functional categories are supported by literature to be involved in defense / disease resistance responses in *Arabidopsis* and other plants. Second-order analysis of the parameters of disease resistance transcriptome, using data from GR978 and another blast-resistant genotype, SHZ-2, revealed that there are genotype non-specific sets of genes that probably constitute the basal defense response pathway of rice plants undergoing a general response to disease infection, as well as genotype-specific gene sets that may be defining the resistance phenotype unique to each variety under study. We found that there were subtly expressed regions of correlated expression that significantly associated with known blast resistance QTLs mapped by other studies. In order to have a stronger, more focused list of candidate genes extracted from microarray experiments for further characterization, we propose a combined first and second-order analytical approach using the parameters of the disease resistance transcriptome (DEGs, RCEs) and statistically associating these parameters with each other and with existing biological information from other related studies (major gene and QTL positions).

## SUMMARY AND CONCLUSION

The gain of blast resistance phenotype of a rice mutant, GR978, originally generated by gamma-irradiation of wild-type indica cultivar IR64, was characterized at the gene expression level using a high density rice oligoarray that represents ∼22,000 genes of the rice genome (Agilent Technologies 22K oligoarray). GR978 was obtained from the IR64 mutant collection at IRRI (Leung et al. 2004), which were generated by various types of mutagens. In the course of describing the mutant phenotypes in the mutant collection, a set of controlled vocabularies documenting the mutant traits observed in ∼3,700 entries was developed. Through a collaboration with the Tos17 rice mutant group (Hirochika, et al., 2004; Miyao, pers comm..), the CV set describing TOs 17 and IR64 mutants were merged and a CV set with 91 descriptions is now available at the IR64 mutant website (http://www.iris.irri.org/action/mutant?method=viewTerm). The table is maintained by IRRI and each CV maps onto public ontology databases (PO, TO, OBO).

The disease resistance transcriptome of rice was characterized using gene expression data from GR978, and another cultivar, SHZ-2, which shows durable resistance to rice blast. The disease resistance transcriptome parameters elucidated were the differentially expressed genes (DEGs), regions of correlated gene expression (RCEs), and the associations between DEGs and RCEs. Differential gene expression was determined statistically within each genotype using MAANOVA, and RCEs were determined within and across genotypes by gene expression correlation analysis along a moving 2 to 15 gene window for the whole genome.

Focusing on the resistance gene locus established by genetic analysis of GR978, candidate genes involved in the resistance phenotype of GR978 were determined by co-localizing the DEGs with the location of the SSR markers that flank the r-gene locus. Twelve candidate genes within the locus were located and listed for further study.

DEGs found within GR978 and SHZ-2 and in common between the two genotypes were filtered for high expression (≥2-fold difference) and clustered for coexpression using hierarchical clustering and association with a common set of enriched cis-elements. Most of the gene clusters determined were associated with defense-response, from inspection of the gene function annotations as well as from results of GO gene category enrichment analysis of the respective DEG clusters. This suggests that most of the highly expressed DEGs effect the resistance phenotype; these DEGs, however, are not necessarily the ones that explain the genetic behavior of the resistance phenotype of GR978.

Comparisons of RCEs between SHZ-2 and GR978 show that although most of the RCEs between genotypes were not located in common regions of the genome, an 8-gene RCE in chromosome 11 was found to be common between SHZ2 and GR978. Inspection of the gene annotations and GO enrichment analysis show that this cluster is highly associated with the resistance response. More interestingly, this region is not populated by DEGs nor is it associated with known blast resistance QTLs. This suggests that subtle co-expression of R genes is involved in the resistance phenotypes of both GR978 and SHZ-2. Comparison of RCEs across genotypes is a novel way of finding interesting candidate genes for disease resistance. Associations between RCEs and DEGs were determined statistically and it was found that there was no enrichment of DEGs in the RCEs within a genotype and across genotypes as well.

QTL data from blast-resistance studies (Bl-QTLs, courtesy of R. Wisser, pers comm., Cornell University) was used to further characterize the disease resistance transcriptome. It was found that Bl-QTLs are not associated with DEGs. Bl-QTLs, however, are significantly associated to genotype-specific RCEs; in GR978, RCEs are in fact enriched within Bl-QTL regions. We put forward a hypothesis that QTLs with small to moderate effects are represented by genome regions in which adjacent genes show correlated expression, suggesting the control of this region by a common regulatory mechanism. We propose cosegregation analysis of RCEs and the resistance phenotype in well-characterized backcross and recombinant inbred lines to test this hypothesis.

## LIST OF ABBREVIATIONS

Bl-QTL: Rice-Blast specific QTL
CDE: Constitutively Differentially Expressed
DEG: Differentially Expressed Gene
DI: Differentially Induced
FL-cDNA: Full-Length complementary DNA
GO: Gene Ontology
HCL: Hierarchical Clustering
KOME: Knowledge-based Oryza Molecular biological Encyclopedia
MAANOVA: Microarray Analysis of Variance
QTL: Quantitative Trait Locus
RCE: Regions of Correlated gene Expression
SHZ-2: Sanhuangzhan 2

