## Supplementary tables for "Identifying Novel Candidate Defense Genes Against Rice Blast By Disease-Resistance Transcriptome Analysis"

Supplementary Table 1. Groups of adjacent genes of correlated expression in GR978 (10 gene window, p < 0.0005)

| GROUP | CHR | TU_START | KOME ID | DESCRIPTION |
|---|---|---|---|---|
| 1 | 1 | 34814238 | AK070194 | *Arabidopsis thaliana* unknown protein (At1g71090) mRNA, complete cds.\|PLN |
| 1 | 1 | 34814238 | AK069848 | *Arabidopsis thaliana* unknown protein (At1g71090) mRNA, complete cds.\|PLN |
| 1 | 1 | 34821626 | AK103555 | Unknown expressed protein |
| 1 | 1 | 34831972 | AK063274 | *Arabidopsis thaliana* At3g08620 mRNA for unknown protein, complete cds, clone: RAFL17-04-D01.\|PLN |
| 1 | 1 | 34842151 | AK059910 | *Populus tremula* X *Populus tremuloides* mRNA for Homeodomain protein HB2, partial.\|PLN |
| 1 | 1 | 34842151 | AK102788 | *Populus tremula* X *Populus tremuloides* mRNA for Homeodomain protein HB2, partial.\|PLN |
| 1 | 1 | 34842151 | AK068585 | *Populus tremula* X *Populus tremuloides* mRNA for Homeodomain protein HB2, partial.\|PLN |
| 1 | 1 | 34867977 | AK102393 | Unknown expressed protein |
| 1 | 1 | 34867977 | AK073454 | *Mus musculus*, clone MGC:30456 IMAGE:3980155, mRNA, complete cds.\|ROD |
| 1 | 1 | 34867977 | AK073866 | Unknown expressed protein |
| 2 | 2 | 1131745 | AK105698 | Unknown expressed protein |
| 2 | 2 | 1145778 | AK110925 | *Quercus ilex* mRNA for putative chloroplast terpene synthase (16 gene).\|PLN |
| 2 | 2 | 1151804 | AK103260 | Unknown expressed protein |
| 2 | 2 | 1154673 | AK109567 | Arabidopsis thaliana clone U10254 unknown protein (At3g04760) mRNA, complete cds.\|PLN |
| 2 | 2 | 1193366 | AK062255 | Arabidopsis thaliana putative arginine/serine-rich splicing factor RSP41 homolog (At5g52040) mRNA, complete cds.\|PLN |
| 2 | 2 | 1193366 | AK065148 | Arabidopsis thaliana putative arginine/serine-rich splicing factor RSP41 homolog (At5g52040) mRNA, complete cds.\|PLN |
| 2 | 2 | 1198100 | AK073966 | Arabidopsis thaliana At1g01860 mRNA for putative dimethyladenosine transferase, complete cds, clone: RAFL17-01-K17.\|PLN |
| 2 | 2 | 1213994 | AK065876 | Arabidopsis thaliana shrunken seed protein (SSE1) mRNA, complete cds.\|PLN |
| 2 | 2 | 1227307 | AK070168 | Arabidopsis thaliana putative NADPH-dependent mannose 6-phosphate reductase (At2g21250) mRNA, complete cds.\|PLN |
| 2 | 2 | 1231020 | AK072529 | Oryza sativa mRNA for phosphatidylinositol synthase.\|PLN |
| 3 | 2 | 3619836 | AK061659 | Oryza sativa (japonica cultivar-group) mRNA for beta-tubulin, complete cds.\|PLN |
| 3 | 2 | 3659089 | AK069128 | Arabidopsis thaliana unknown protein (At2g02560) mRNA, complete cds.\|PLN |
| 3 | 2 | 3674678 | AK106654 | Glycine max urease accessory protein UreF mRNA, complete cds.\|PLN |
| 3 | 2 | 3677462 | AK058890 | Hordeum vulgare mRNA for 4-hydroxyphenylpyruvate dioxygenase.\|PLN |
| 3 | 2 | 3687915 | AK103301 | Arabidopsis thaliana unknown protein (At1g06660) mRNA, complete cds.\|PLN |
| 3 | 2 | 3702927 | AK073338 | Oryza sativa clone Osh16 unknown protein mRNA, partial cds.\|PLN |
| 3 | 2 | 3705882 | AK067291 | Arabidopsis thaliana clone 124466 mRNA, complete sequence.\|PLN |
| 3 | 2 | 3711140 | AK061562 | Arabidopsis thaliana mRNA for FKBP like protein (fkbp25I gene).\|PLN |
| 3 | 2 | 3715271 | AK102265 | Triticum aestivum porphobilinogen deaminase mRNA, partial cds.\|PLN |
| 3 | 2 | 3718322 | AK102098 | Unknown expressed protein |
| 4 | 2 | 21506959 | AK068282 | Unknown expressed protein |
| 4 | 2 | 21508582 | AK066393 | Arabidopsis thaliana unknown protein (At1g07700) mRNA, complete cds.\|PLN |
| 4 | 2 | 21514322 | AK069840 | Arabidopsis thaliana unknown protein (At1g47380) mRNA, complete cds.\|PLN |

Appendix Table 1. continued…

| GROUP | CHR | TU_START | KOME ID | DESCRIPTION |
|---|---|---|---|---|
| 4 | 2 | 21532878 | AK105793 | Arabidopsis thaliana clone 111056 mRNA, complete sequence.\|PLN |
| 4 | 2 | 21543249 | AK060161 | Nicotiana plumbaginifolia partial mRNA for RNA Binding Protein 47 (rbp47 gene).\|PLN |
| 4 | 2 | 21607719 | AK059010 | Oryza sativa Cyt-P450 monooxygenase (PM-II) mRNA, complete cds.\|PLN |
| 4 | 2 | 21635489 | AK060722 | Oryza sativa Cyt-P450 monooxygenase (PM-II) mRNA, complete cds.\|PLN |
| 4 | 2 | 21685415 | AK105913 | Oryza sativa Cyt-P450 monooxygenase (PM-II) mRNA, complete cds.\|PLN |
| 4 | 2 | 21710122 | AK068310 | Oryza sativa OsDTC1 mRNA for putative diterpene cyclase, complete cds.\|PLN |
| 4 | 2 | 21733281 | AK107418 | Solanum tuberosum mRNA for cytochrome P450 (CYP71D4 gene).\|PLN |
| 4 | 2 | 21755452 | AK070167 | Triticum aestivum N-1 mRNA for cytochrome P450, complete cds.\|PLN |
| 4 | 2 | 21849467 | AK101003 | Oryza sativa Cyt-P450 monooxygenase (PM-II) mRNA, complete cds.\|PLN |
| 4 | 2 | 21872680 | AK109940 | Oryza sativa early proembryo mRNA, complete sequence.\|PLN |
| 4 | 2 | 21926092 | AK066007 | Arabidopsis thaliana clone 142216 mRNA, complete sequence.\|PLN |
| 4 | 2 | 21955857 | AK099415 | Oryza sativa OsMST1 mRNA for monosaccharide transporter 1, complete cds.\|PLN |
| 5 | 3 | 5252913 | AK107086 | Unknown expressed protein |
| 5 | 3 | 5256593 | AK069251 | Oryza sativa mRNA for cyc07, complete cds.\|PLN |
| 5 | 3 | 5265126 | AK103392 | Arabidopsis thaliana Surfeit 1 (SURF1) mRNA, complete cds.\|PLN |
| 5 | 3 | 5269348 | AK112007 | Arabidopsis thaliana ADP-ribosylation factor-like protein (At3g49870) mRNA, complete cds.\|PLN |
| 5 | 3 | 5276190 | AK069273 | Arabidopsis thaliana AT5g54540/MRB17_4 mRNA, complete cds.\|PLN |
| 5 | 3 | 5279001 | AK102337 | Arabidopsis thaliana At5g67530/K9I9_9 mRNA, complete cds.\|PLN |
| 5 | 3 | 5302441 | AK065564 | Mus musculus, Similar to hypothetical protein FLJ23476, clone MGC:38336 IMAGE:5343817, mRNA, complete cds.\|ROD |
| 5 | 3 | 5302441 | AK067825 | Mus musculus, Similar to hypothetical protein FLJ23476, clone MGC:38336 IMAGE:5343817, mRNA, complete cds.\|ROD |
| 5 | 3 | 5306289 | AK102580 | Unknown expressed protein |
| 5 | 3 | 5354651 | AK071140 | Unknown expressed protein |
| 5 | 3 | 5375600 | AK065462 | Dactylis glomerata somatic embryogenesis related protein mRNA, complete cds.\|PLN |
| 5 | 3 | 5382627 | AK102234 | Unknown expressed protein |
| 5 | 3 | 5415581 | AK100415 | Medicago truncatula type IIB calcium ATPase MCA5 (MCA5) mRNA, complete cds.\|PLN |
| 6 | 4 | 19935234 | AK108077 | Zea mays ZmRCP2 mRNA for root cap protein 2, complete cds.\|PLN |
| 6 | 4 | 19965028 | AK070396 | Unknown expressed protein |
| 6 | 4 | 19993750 | AK102032 | Arabidopsis thaliana unknown protein (At3g48410) mRNA, complete cds.\|PLN |
| 6 | 4 | 20013514 | AK108223 | Arabidopsis thaliana AT3g52740/F3C22_140 mRNA, complete cds.\|PLN |
| 6 | 4 | 20020484 | AK062016 | C.sinensis mRNA for non-photosynthetic ferredoxin.\|PLN |
| 6 | 4 | 20027785 | AK110902 | Arabidopsis thaliana putative beta-1,3-glucanase (At2g27500) mRNA, complete cds.\|PLN |
| 6 | 4 | 20033597 | AK111482 | Unknown expressed protein |
| 6 | 4 | 20046859 | AK108347 | Unknown expressed protein |
| 6 | 4 | 20052966 | AK073418 | Arabidopsis thaliana Unknown protein (At2g27460; F10A12.14) mRNA, complete cds.\|PLN |
| 6 | 4 | 20098163 | AK100104 | Unknown expressed protein |

Supplementary Table 1. continued...

| GROUP | CHR | TU_START | KOME ID | DESCRIPTION |
|---|---|---|---|---|
| 6 | 4 | 20139031 | AK058617 | Unknown expressed protein |
| 6 | 4 | 20161656 | AK109638 | Unknown expressed protein |
| 7 | 5 | 18203542 | AK106900 | Arabidopsis thaliana unknown protein mRNA, complete cds.\|PLN |
| 7 | 5 | 18275043 | AK105553 | Unknown expressed protein |
| 7 | 5 | 18278625 | AK072704 | Arabidopsis thaliana putative GTPase-activating protein (At3g07940) mRNA, complete cds.\|PLN |
| 7 | 5 | 18305994 | AK068231 | Cicer arietinum mRNA for annexin.\|PLN |
| 7 | 5 | 18312340 | AK061728 | Arabidopsis thaliana clone 3294 mRNA, complete sequence.\|PLN |
| 7 | 5 | 18314669 | AK058345 | Arabidopsis thaliana AT3g57090/F24I3_170 mRNA, complete cds.\|PLN |
| 7 | 5 | 18328398 | AK107536 | Unknown expressed protein |
| 7 | 5 | 18387336 | AK111176 | Arabidopsis thaliana clone U10087 putative ABC transporter protein (At2g13610) mRNA, complete cds.\|PLN |
| 7 | 5 | 18398666 | AK069987 | Arabidopsis thaliana zinc finger protein (PRAF1) mRNA, complete cds.\|PLN |
| 7 | 5 | 18440962 | AK109014 | Unknown expressed protein |
| 8 | 6 | 21371402 | AK068350 | Unknown expressed protein |
| 8 | 6 | 21372999 | AK069355 | Tomato acid phosphatase (aps1) mRNA, complete cds.\|PLN |
| 8 | 6 | 21372999 | AK060112 | Tomato acid phosphatase (aps1) mRNA, complete cds.\|PLN |
| 8 | 6 | 21396008 | AK063490 | Mus musculus iron-regulated transporter IREG1 (Ireg1) mRNA, complete cds.\|ROD |
| 8 | 6 | 21430260 | AK108074 | Arabidopsis thaliana AT4g17486/AT4g17486 mRNA, complete cds.\|PLN |
| 8 | 6 | 21597235 | AK067114 | A.sativa (Pewi) ASTCP-K36 mRNA for t complex polypeptide 1.\|PLN |
| 8 | 6 | 21613168 | AK109753 | Arabidopsis thaliana Unknown protein (T18E12.18; At2g03150) mRNA, partial cds.\|PLN |
| 8 | 6 | 21636463 | AK099669 | Arabidopsis thaliana AT5g56750/MIK19_22 mRNA, complete cds.\|PLN |
| 8 | 6 | 21653899 | AK111862 | Oryza sativa protein phosphatase 2A 55 kDa B regulatory subunit mRNA, complete cds.\|PLN |
| 8 | 6 | 21653908 | AK111494 | Oryza sativa protein phosphatase 2A 55 kDa B regulatory subunit mRNA, complete cds.\|PLN |
| 9 | 6 | 24228506 | AK072533 | Danio rerio fast skeletal muscle troponin C (tnnc) mRNA, complete cds.\|VRT |
| 9 | 6 | 24232599 | AK067228 | Arabidopsis thaliana At4g17150 mRNA for unknown protein, complete cds, clone: RAFL21-18-K13.\|PLN |
| 9 | 6 | 24267674 | AK062635 | Unknown expressed protein |
| 9 | 6 | 24287219 | AK109906 | Zea mays seven transmembrane protein Mlo7 mRNA, complete cds.\|PLN |
| 9 | 6 | 24299115 | AK063128 | Unknown expressed protein |
| 9 | 6 | 24303827 | AK066754 | Nicotiana tabacum mRNA for CND41, chloroplast nucleoid DNA binding protein with aspartic protease activity, complete cds.\|PLN |
| 9 | 6 | 24303827 | AK062719 | Unknown expressed protein |
| 9 | 6 | 24314444 | AK067236 | Nicotiana tabacum mRNA for microtubule-associated protein MAP65-1a (map65-1a gene).\|PLN |
| 9 | 6 | 24319205 | AK109238 | Unknown expressed protein |
| 9 | 6 | 24328419 | AK107800 | Unknown expressed protein |
| 9 | 6 | 24339229 | AK072708 | Z.mays mRNA for polygalacturonase (clone PG2).\|PLN |
| 9 | 6 | 24354750 | AK111644 | Avena sativa victorin binding protein mRNA, complete cds.\|PLN |
| 10 | 7 | 21297936 | AK105928 | Arabidopsis thaliana AT3g08780/F17O14_25 mRNA, complete cds.\|PLN |

Supplementary Table 1. continued...

| GROUP | CHR | TU_START | KOME ID | DESCRIPTION |
|---|---|---|---|---|
| 10 | 7 | 21357958 | AK100722 | Arabidopsis thaliana mRNA for receptor-like protein kinase.\|PLN |
| 10 | 7 | 21383094 | AK111550 | Arabidopsis thaliana At4g23180 mRNA for putative receptor-like protein kinase 4 (RLK4), complete cds, clone: RAFL17-45-D12.\|PLN |
| 10 | 7 | 21389808 | AK110523 | Unknown expressed protein |
| 10 | 7 | 21389808 | AK064568 | Unknown expressed protein |
| 10 | 7 | 21393500 | AK105128 | Arabidopsis thaliana At4g23140 mRNA for putative receptor-like protein kinase 5 (RLK5), complete cds, clone: RAFL19-72-N19.\|PLN |
| 10 | 7 | 21406209 | AK102941 | Arabidopsis thaliana At4g23180 mRNA for putative receptor-like protein kinase 4 (RLK4), complete cds, clone: RAFL17-45-D12.\|PLN |
| 10 | 7 | 21432948 | AK111865 | Phaseolus vulgaris receptor-like protein kinase homolog RK20-1 mRNA, complete cds.\|PLN |
| 10 | 7 | 21465080 | AK068361 | Arabidopsis thaliana Unknown protein (At5g54680) mRNA, complete cds.\|PLN |
| 10 | 7 | 21518679 | AK105710 | Unknown expressed protein |
| 11 | 8 | 14843167 | AK060562 | Pisum sativum xyloglucan fucosyltransferase mRNA, complete cds.\|PLN |
| 11 | 8 | 14960824 | AK064536 | Unknown expressed protein |
| 11 | 8 | 14982168 | AK099602 | Unknown expressed protein |
| 11 | 8 | 14982168 | AK072420 | Homo sapiens mRNA for TRX5 protein.\|PRI |
| 11 | 8 | 15030770 | AK100576 | Arabidopsis thaliana clone 100460 mRNA, complete sequence.\|PLN |
| 11 | 8 | 15070350 | AK068124 | Homo sapiens, general transcription factor IIH, polypeptide 1 (62kD subunit), clone MGC:4300 IMAGE:2819217, mRNA, complete cds.\|PRI |
| 11 | 8 | 15094469 | AK109191 | Unknown expressed protein |
| 11 | 8 | 15100069 | AK059052 | Arabidopsis thaliana clone 35284 mRNA, complete sequence.\|PLN |
| 11 | 8 | 15123189 | AK073476 | Zea mays enhancer of polycomb-like protein (epc101) mRNA, complete cds.\|PLN |
| 11 | 8 | 15140714 | AK073244 | Unknown expressed protein |
| 11 | 8 | 15221379 | AK068279 | Arabidopsis thaliana At5g03800 mRNA for unknown protein, complete cds, clone: RAFL16-75-K17.\|PLN |
| 11 | 8 | 15258731 | AK072532 | Lotus japonicus phosphatidylinositol transfer-like protein III (LjPLP-III) mRNA, complete cds.\|PLN |
| 12 | 9 | 10844762 | AK068895 | Unknown expressed protein |
| 12 | 9 | 10861935 | AK100597 | Triticum aestivum Sec61p (sec61) mRNA, complete cds.\|PLN |
| 12 | 9 | 10869048 | AK101995 | Zea mays histone acetyl transferase (hac106) mRNA, complete cds.\|PLN |
| 12 | 9 | 10875106 | AK071224 | Arabidopsis thaliana unknown protein (At5g55810) mRNA, complete cds.\|PLN |
| 12 | 9 | 10924313 | AK063411 | Zea mays transposon Doppia transposase DOPD and transposase DOPA mRNA, complete cds.\|PLN |
| 12 | 9 | 11183249 | AK065335 | Arabidopsis thaliana putative bystin (At1g31660) mRNA, complete cds.\|PLN |
| 12 | 9 | 11321555 | AK069410 | Unknown expressed protein |
| 12 | 9 | 11333855 | AK105685 | Oryza sativa mRNA for OsD305, complete cds.\|PLN |
| 12 | 9 | 11395157 | AK072578 | Unknown expressed protein |
| 12 | 9 | 11649601 | AK059689 | Arabidopsis thaliana arginine methyltransferase pam1 (At4g29510) mRNA, complete cds.\|PLN |
| 13 | 9 | 15011730 | AK068557 | Unknown expressed protein |
| 13 | 9 | 15022954 | AK107198 | Hordeum vulgare cultivar Orthega cinnamoyl CoA reductase mRNA, complete cds.\|PLN |

Supplementary Table 1. continued…

| GROUP | CHR | TU_START | KOME ID | DESCRIPTION |
|---|---|---|---|---|
| 13 | 9 | 15037373 | AK073786 | Arabidopsis thaliana clone 108517 mRNA, complete sequence.\|PLN |
| 13 | 9 | 15041211 | AK109320 | Arabidopsis thaliana clone 108517 mRNA, complete sequence.\|PLN |
| 13 | 9 | 15072960 | AK068795 | Arabidopsis thaliana clone 108517 mRNA, complete sequence.\|PLN |
| 13 | 9 | 15100475 | AK072629 | Arabidopsis thaliana clone 3836 mRNA, complete sequence.\|PLN |
| 13 | 9 | 15103705 | AK111264 | Arabidopsis thaliana Cu-chaperone (COX17) mRNA, complete cds.\|PLN |
| 13 | 9 | 15107602 | AK099513 | Deschampsia antarctica clone Dacor 1.0 polyubiquitin 2 mRNA, complete cds.\|PLN |
| 13 | 9 | 15109795 | AK065475 | Arabidopsis thaliana unknown protein (At4g31820) mRNA, complete cds.\|PLN |
| 13 | 9 | 15198704 | AK101900 | Nicotiana tabacum mRNA for hydroxycinnamoyl transferase (hct gene).\|PLN |
| 14 | 11 | 6637634 | AK058632 | Unknown expressed protein |
| 14 | 11 | 6637634 | AK059194 | Unknown expressed protein |
| 14 | 11 | 6657155 | AK065723 | Oryza sativa (indica cultivar-group) mRNA for RPR1h, complete cds.\|PLN |
| 14 | 11 | 6674313 | AK064544 | Unknown expressed protein |
| 14 | 11 | 6683962 | AK102193 | Oryza sativa (japonica cultivar-group) NBS-LRR-like protein (YR5) mRNA, complete cds.\|PLN |
| 14 | 11 | 6683962 | AK101375 | Oryza sativa (japonica cultivar-group) NBS-LRR-like protein (YR5) mRNA, complete cds.\|PLN |
| 14 | 11 | 6841012 | AK108061 | Unknown expressed protein |
| 14 | 11 | 6850392 | AK102213 | Unknown expressed protein |
| 14 | 11 | 6876697 | AK111665 | Oryza sativa (japonica cultivar-group) mRNA for RPR1, complete cds.\|PLN |
| 14 | 11 | 6957275 | AK108047 | Unknown expressed protein |
| 15 | 12 | 11275929 | AK099574 | Oryza sativa (japonica cultivar-group) mRNA for the small subunit of ribulose-1,5-bisphosphate carboxylase, complete cds, clone pOSSS2106.\|PLN |
| 15 | 12 | 11318956 | AK068266 | Oryza sativa (japonica cultivar-group) mRNA for the small subunit of ribulose-1,5-bisphosphate carboxylase, complete cds, clone pOSSS1139.\|PLN |
| 15 | 12 | 11348497 | AK066177 | Arabidopsis thaliana At1g67840/F12A21_3 mRNA, complete cds.\|PLN |
| 15 | 12 | 11418673 | AK111535 | Arabidopsis thaliana unknown protein (At2g47790/F17A22.18) mRNA, complete cds.\|PLN |
| 15 | 12 | 11714578 | AK061658 | Unknown expressed protein |
| 15 | 12 | 11726084 | AK072331 | Solanum tuberosum starch associated protein R1 mRNA, complete cds.\|PLN |
| 15 | 12 | 11903236 | AK103895 | Unknown expressed protein |
| 15 | 12 | 11914119 | AK071145 | Oryza sativa mRNA for SAR DNA binding protein, partial cds.\|PLN |
| 15 | 12 | 12073829 | AK064189 | Arabidopsis thaliana At3g07750 mRNA for putative 3 exoribonuclease, complete cds, clone: RAFL21-33-K24.\|PLN |
| 15 | 12 | 12073829 | AK059940 | Arabidopsis thaliana At3g07750 mRNA for putative 3 exoribonuclease, complete cds, clone: RAFL21-33-K24.\|PLN |
| 16 | 12 | 18725053 | AK110307 | Triticum aestivum stripe rust resistance protein Yr10 (Yr10) mRNA, complete cds.\|PLN |
| 16 | 12 | 18822849 | AK105366 | Arabidopsis thaliana putative transcription co-activator (SYT3) mRNA, complete cds.\|PLN |
| 16 | 12 | 18846515 | AK071243 | Arabidopsis thaliana putative translation initiation factor EIF-2B alpha subunit (At1g72340) mRNA, complete cds.\|PLN |
| 16 | 12 | 18846515 | AK068908 | Arabidopsis thaliana putative translation initiation factor EIF-2B alpha subunit (At1g72340) mRNA, complete cds.\|PLN |
| 16 | 12 | 18880405 | AK070044 | Arabidopsis thaliana clone RAFL15-27-J23 (R20860) unknown protein (At1g69390) mRNA, complete cds.\|PLN |
| 16 | 12 | 18899996 | AK073582 | Homo sapiens, clone MGC:8856 IMAGE:3890907, mRNA, complete cds.\|PRI |

Supplementary Table 1. continued.

| GROUP | CHR | TU_START | KOME ID | DESCRIPTION |
|---|---|---|---|---|
| 16 | 12 | 18905820 | AK067767 | Arabidopsis thaliana clone RAFL14-85-F19 (R20222) unknown protein (At5g55610) mRNA, complete cds.\|PLN |
| 16 | 12 | 19088840 | AK060063 | Arabidopsis thaliana clone 30535 mRNA, complete sequence.\|PLN |
| 16 | 12 | 19105480 | AK106653 | Lycopersicon esculentum mRNA for cyclin A2 (CycA2 gene).\|PLN |
| 16 | 12 | 19114474 | AK111001 | Arabidopsis thaliana putative 3-phosphoserine phosphatase (At1g18640) mRNA, complete cds.\|PLN |
| 16 | 12 | 19118282 | AK069786 | Arabidopsis thaliana At4g08790/T32A17_100 mRNA, complete cds.\|PLN |

Supplementary Table 2. Groups of adjacent genes of correlated expression in GR978-SHZ (10 gene window, p < 0.0005)

| GROUP | CHR | TU_START | GENENAME | DESCRIPTION |
|---|---|---|---|---|
| 1 | 1 | 21606816 | AK060571 | Oryza sativa abscisic acid- and stress-inducible protein (Asr1) mRNA, complete cds.\|PLN |
| 1 | 1 | 21610840 | AK065822 | Unknown expressed protein |
| 1 | 1 | 21613634 | AK107516 | Unknown expressed protein |
| 1 | 1 | 21616693 | AK065173 | Unknown expressed protein |
| 1 | 1 | 21641070 | AK106499 | Unknown expressed protein |
| 1 | 1 | 21645342 | AK058473 | Arabidopsis thaliana clone U19580 unknown protein (At4g32810) mRNA, complete cds.\|PLN |
| 1 | 1 | 21678614 | AK100223 | Arabidopsis thaliana clone 26655 mRNA, complete sequence.\|PLN |
| 1 | 1 | 21693394 | AK071488 | Unknown expressed protein |
| 1 | 1 | 21699523 | AK069327 | Oryza sativa OsMST3 mRNA for monosaccharide transporter 3, complete cds.\|PLN |
| 1 | 1 | 21876276 | AK061023 | Nicotiana tabacum mRNA for putative carbamoyl phosphate synthase large subunit (CT4 gene).\|PLN |
| 2 | 2 | 34879650 | AK059429 | Arabidopsis thaliana unknown protein (At5g62030) mRNA, complete cds.\|PLN |
| 2 | 2 | 34890103 | AK102246 | Arabidopsis thaliana clone U19358 putative protein kinase (At2g20300) mRNA, complete cds.\|PLN |
| 2 | 2 | 34908735 | AK058331 | Arabidopsis thaliana F25I18.1/F25I18.1 mRNA, complete cds.\|PLN |
| 2 | 2 | 34912707 | AK060243 | Arabidopsis thaliana clone 34612 mRNA, complete sequence.\|PLN |
| 2 | 2 | 34927286 | AK071879 | Arabidopsis thaliana unknown protein (At2g20195) mRNA, complete cds.\|PLN |
| 2 | 2 | 34950025 | AK058624 | Arabidopsis thaliana AT3g24190/MUJ8_17 mRNA, complete cds.\|PLN |
| 2 | 2 | 34956503 | AK059572 | Unknown expressed protein |
| 2 | 2 | 34958679 | AK102271 | Arabidopsis thaliana putative NADH-ubiquinone oxireductase (At2g20360) mRNA, complete cds.\|PLN |
| 2 | 2 | 34986184 | AK099758 | Unknown expressed protein |
| 2 | 2 | 35012312 | AK110894 | Oryza sativa potassium channel beta subunit protein (KOB1) mRNA, complete cds.\|PLN |
| 3 | 3 | 3452852 | AK063366 | Arabidopsis thaliana unknown protein (At4g38180) mRNA, complete cds.\|PLN |
| 3 | 3 | 3521247 | AK062304 | Arabidopsis thaliana hypothetical protein (At2g27640/F15K20.26) mRNA, complete cds.\|PLN |
| 3 | 3 | 3543176 | AK072498 | Arabidopsis thaliana At1g76010/T4O12_22 mRNA, complete cds.\|PLN |
| 3 | 3 | 3638494 | AK069884 | Arabidopsis thaliana AT3g58500/F14P22_90 mRNA, complete cds.\|PLN |
| 3 | 3 | 3655822 | AK072898 | Arabidopsis thaliana AT3g60070/T2O9_50 mRNA, complete cds.\|PLN |
| 3 | 3 | 3662775 | AK099530 | Arabidopsis thaliana AT4g27310/M4I22_120 mRNA, complete cds.\|PLN |
| 3 | 3 | 3695167 | AK107189 | Arabidopsis thaliana clone RAFL15-01-M05 (R20306) putative axi 1 protein (At2g03280) mRNA, complete cds.\|PLN |
| 3 | 3 | 3709437 | AK068500 | Unknown expressed protein |
| 3 | 3 | 3717713 | AK066912 | Cucurbita maxima mRNA for AOBP (ascorbate oxidase promoter-binding protein), complete cds.\|PLN |
| 3 | 3 | 3720543 | AK060900 | Cucurbita maxima mRNA for AOBP (ascorbate oxidase promoter-binding protein), complete cds.\|PLN |
| 3 | 3 | 3726121 | AK068278 | Arabidopsis thaliana At1g18680 mRNA for unknown protein, complete cds, clone: RAFL19-32-I16.\|PLN |
| 3 | 3 | 3729608 | AK063222 | Cicer arietinum partial mRNA for hypothetical protein (ORF1), clone Can74-2.\|PLN |
| 3 | 3 | 3742195 | AK060801 | Unknown expressed protein |
| 4 | 3 | 5252913 | AK107086 | Unknown expressed protein |

Supplementary Table 2. continued…

| GROUP | CHR | TU_START | GENENAME | DESCRIPTION |
|---|---|---|---|---|
| 4 | 3 | 5256593 | AK069251 | Oryza sativa mRNA for cyc07, complete cds.\|PLN |
| 4 | 3 | 5265126 | AK103392 | Arabidopsis thaliana Surfeit 1 (SURF1) mRNA, complete cds.\|PLN |
| 4 | 3 | 5269348 | AK112007 | Arabidopsis thaliana ADP-ribosylation factor-like protein (At3g49870) mRNA, complete cds.\|PLN |
| 4 | 3 | 5276190 | AK069273 | Arabidopsis thaliana AT5g54540/MRB17_4 mRNA, complete cds.\|PLN |
| 4 | 3 | 5279001 | AK102337 | Arabidopsis thaliana At5g67530/K9I9_9 mRNA, complete cds.\|PLN |
| 4 | 3 | 5302441 | AK067825 | Mus musculus, Similar to hypothetical protein FLJ23476, clone MGC:38336 IMAGE:5343817, mRNA, complete cds.\|ROD |
| 4 | 3 | 5306289 | AK102580 | Unknown expressed protein |
| 4 | 3 | 5354651 | AK071140 | Unknown expressed protein |
| 4 | 3 | 5382627 | AK102234 | Unknown expressed protein |
| 4 | 3 | 5415581 | AK100415 | Medicago truncatula type IIB calcium ATPase MCA5 (MCA5) mRNA, complete cds.\|PLN |
| 5 | 3 | 17788216 | AK099511 | Arabidopsis thaliana At1g22050 mRNA for unknown protein, complete cds, clone: RAFL21-14-A13.\|PLN |
| 5 | 3 | 17804045 | AK069737 | Arabidopsis thaliana AT5g39360/MUL8_40 mRNA, complete cds.\|PLN |
| 5 | 3 | 17814537 | AK111492 | Arabidopsis thaliana clone RAFL15-02-A01 (R20221) unknown protein (At3g29270) mRNA, complete cds.\|PLN |
| 5 | 3 | 17898814 | AK111273 | Festuca pratensis partial mRNA for expansin (exp5 gene).\|PLN |
| 5 | 3 | 17913307 | AK102768 | Arabidopsis thaliana clone 31680 mRNA, complete sequence.\|PLN |
| 5 | 3 | 17980142 | AK068917 | Arabidopsis thaliana At5g67610 mRNA for unknown protein, complete cds, clone: RAFL21-02-I18.\|PLN |
| 5 | 3 | 17990778 | AK059145 | Arabidopsis thaliana unknown protein (At4g00990) mRNA, complete cds.\|PLN |
| 5 | 3 | 18074464 | AK110714 | Unknown expressed protein |
| 5 | 3 | 18117188 | AK065739 | Oryza sativa subsp. indica mRNA for cytosolic pyruvate orthophosphate dikinase.\|PLN |
| 5 | 3 | 18353811 | AK107787 | Arabidopsis thaliana clone RAFL15-11-J08 (R20460) unknown protein (At1g73790) mRNA, complete cds.\|PLN |
| 6 | 4 | 12215681 | AK064584 | Unknown expressed protein |
| 6 | 4 | 12364399 | AK062593 | A.thaliana mRNA for receptor like protein kinase.\|PLN |
| 6 | 4 | 12457651 | AK063164 | Unknown expressed protein |
| 6 | 4 | 12667645 | AK111082 | Arabidopsis thaliana clone RAFL15-20-M04 (R20457) putative short chain alcohol dehydrogenase (At5g06060) mRNA, complete cds.\|PLN |
| 6 | 4 | 12782046 | AK105141 | Hordeum vulgare mRNA for expressed sequence tag.\|PLN |
| 6 | 4 | 12826224 | AK107143 | Unknown expressed protein |
| 6 | 4 | 12913521 | AK061424 | Unknown expressed protein |
| 6 | 4 | 13349714 | AK072639 | Arabidopsis thaliana putative S-receptor kinase (At4g32300) mRNA, complete cds.\|PLN |
| 6 | 4 | 13355877 | AK069467 | Arabidopsis thaliana clone sps843 unknown mRNA.\|PLN |
| 6 | 4 | 13437894 | AK108051 | Arabidopsis thaliana unknown protein (At3g59980) mRNA, complete cds.\|PLN |
| 6 | 4 | 13632217 | AK103443 | Arabidopsis thaliana clone 3116 mRNA, complete sequence.\|PLN |
| 6 | 4 | 13651623 | AK067583 | Arabidopsis thaliana clone 33530 mRNA, complete sequence.\|PLN |
| 6 | 4 | 13664679 | AK070019 | Arabidopsis thaliana unknown protein (At4g21180) mRNA, partial cds.\|PLN |
| 6 | 4 | 13693163 | AK067041 | Arabidopsis thaliana At1g79670/F20B17_27 mRNA, complete cds.\|PLN |
| 7 | 4 | 30359231 | AK099648 | Arabidopsis thaliana 26S proteasome regulatory subunit (At2g32730) mRNA, complete cds.\|PLN |

Supplementary Table 2. continued…

| GROUP | CHR | TU_START | GENENAME | DESCRIPTION |
|---|---|---|---|---|
| 7 | 4 | 30367077 | AK067687 | Arabidopsis thaliana putative receptor kinase (At4g31240) mRNA, complete cds.\|PLN |
| 7 | 4 | 30381973 | AK065298 | Arabidopsis thaliana At1g55500/T5A14_10 mRNA, complete cds.\|PLN |
| 7 | 4 | 30413178 | AK103652 | Arabidopsis thaliana AT3g13050/MGH6_16 mRNA, complete cds.\|PLN |
| 7 | 4 | 30436061 | AK070129 | Arabidopsis thaliana putative protein phosphatase 2C (At4g28400) mRNA, complete cds.\|PLN |
| 7 | 4 | 30448996 | AK058541 | Unknown expressed protein |
| 7 | 4 | 30448996 | AK101763 | Homo sapiens mRNA for FLJ00173 protein.\|PRI |
| 7 | 4 | 30475060 | AK060380 | Unknown expressed protein |
| 7 | 4 | 30481407 | AK062284 | Unknown expressed protein |
| 7 | 4 | 30513340 | AK100672 | Arabidopsis thaliana putative potassium transporter AtKT5p (At4g33530) mRNA, complete cds.\|PLN |
| 8 | 5 | 27815824 | AK111480 | Arabidopsis thaliana putative protein (At5g44170) mRNA, complete cds.\|PLN |
| 8 | 5 | 27823563 | AK065766 | Oryza sativa hypothetical protein mRNA, partial cds.\|PLN |
| 8 | 5 | 27831280 | AK109344 | Arabidopsis thaliana At5g60860 mRNA for putative GTP-binding protein, complete cds, clone: RAFL19-94-D08.\|PLN |
| 8 | 5 | 27863654 | AK067090 | Solanum tuberosum mRNA for urease accessory protein G (ureG gene) isoform R3.\|PLN |
| 8 | 5 | 27921570 | AK099641 | Arabidopsis thaliana unknown protein (At5g46420) mRNA, complete cds.\|PLN |
| 8 | 5 | 27927209 | AK100389 | Oryza sativa (japonica cultivar-group) mRNA for putative mitogen-activated protein kinase wjumk1 (wjumk1 gene).\|PLN |
| 8 | 5 | 27927209 | AK061645 | Oryza sativa (japonica cultivar-group) mRNA for putative mitogen-activated protein kinase wjumk1 (wjumk1 gene).\|PLN |
| 8 | 5 | 27943924 | AK058475 | Arabidopsis thaliana AT5g64130/MHJ24_11 mRNA, complete cds.\|PLN |
| 8 | 5 | 27952409 | AK065748 | Oryza sativa (japonica cultivar-group) cold acclimation protein COR413-TM1 mRNA, complete cds.\|PLN |
| 8 | 5 | 27970240 | AK066453 | Rice mRNA for aspartic protease, complete cds.\|PLN |
| 8 | 5 | 27970240 | AK100749 | Rice mRNA for aspartic protease, complete cds.\|PLN |
| 9 | 5 | 28540605 | AK108477 | Pisum sativum polygalacturonase (Ppg1) mRNA, complete cds.\|PLN |
| 9 | 5 | 28557879 | AK068594 | Beta vulgaris integral membrane protein mRNA, complete cds.\|PLN |
| 9 | 5 | 28593248 | AK064624 | Arabidopsis thaliana clone 32047 mRNA, complete sequence.\|PLN |
| 9 | 5 | 28602504 | AK063417 | Arabidopsis thaliana mRNA for auxilin-like protein.\|PLN |
| 9 | 5 | 28610237 | AK100910 | Oryza sativa (japonica cultivar-group) mRNA for ADP glucose pyrophosphorylase large subunit, complete cds.\|PLN |
| 9 | 5 | 28627985 | AK102000 | Arabidopsis thaliana mRNA for cohesin.\|PLN |
| 9 | 5 | 28671725 | AK108181 | Unknown expressed protein |
| 9 | 5 | 28678584 | AK102861 | Arabidopsis thaliana unknown protein (At1g29370) mRNA, complete cds.\|PLN |
| 9 | 5 | 28698668 | AK058748 | Arabidopsis thaliana putative protein (At5g46340) mRNA, complete cds.\|PLN |
| 9 | 5 | 28723051 | AK066579 | Arabidopsis thaliana At SPS mRNA for solanesyl diphosphate synthase, complete cds.\|PLN |
| 10 | 7 | 1100145 | AK103259 | Unknown expressed protein |
| 10 | 7 | 1104151 | AK072975 | Unknown expressed protein |
| 10 | 7 | 1104151 | AK073714 | Unknown expressed protein |
| 10 | 7 | 1112267 | AK102706 | Unknown expressed protein |
| 10 | 7 | 1141850 | AK064136 | Unknown expressed protein |

Supplementary Table 2. continued…

| GROUP | CHR | TU_START | GENENAME | DESCRIPTION |
|---|---|---|---|---|
| 10 | 7 | 1148041 | AK102508 | Unknown expressed protein |
| 10 | 7 | 1151566 | AK072846 | Unknown expressed protein |
| 10 | 7 | 1197514 | AK108490 | Arabidopsis thaliana unknown protein (At2g16070) mRNA, complete cds.\|PLN |
| 10 | 7 | 1253344 | AK063647 | Oryza sativa mRNA for putative phytosulfokine peptide precursor (psk4 gene).\|PLN |
| 10 | 7 | 1267339 | AK111259 | Unknown expressed protein |
| 11 | 8 | 15306493 | AK073790 | Zea mays (clone pAKHSDH1) aspartate kinase-homoserine dehydrogenase mRNA, complete cds.\|PLN |
| 11 | 8 | 15343716 | AK068357 | Oryza sativa S-locus receptor-like kinase RLK13 (RLK13) mRNA, complete cds.\|PLN |
| 11 | 8 | 15361398 | AK067775 | Arabidopsis thaliana unknown protein (At1g03280) mRNA, complete cds.\|PLN |
| 11 | 8 | 15421299 | AK111861 | Zea mays histone deacetylase (HDA108) mRNA, partial cds.\|PLN |
| 11 | 8 | 15459419 | AK066618 | Zea mays plastid phosphate/phosphoenolpyruvate translocator precursor (MZPPT1) mRNA, complete cds.\|PLN |
| 11 | 8 | 15459419 | AK059591 | Oryza sativa phosphoenolpyruvate/phosphate translocator mRNA, complete cds.\|PLN |
| 11 | 8 | 15525653 | AK099939 | Citrus X paradisi pyrophosphate-dependent phosphofructokinase alpha subunit (PPi-PFKa) mRNA, complete cds.\|PLN |
| 11 | 8 | 15533361 | AK058551 | O.sativa ADP-glucose pyrophosphorylase 51kD subunit mRNA, complete cds.\|PLN |
| 11 | 8 | 15558744 | AK103873 | Arabidopsis thaliana AT3g24120/MUJ8_3 mRNA, complete cds.\|PLN |
| 11 | 8 | 15622921 | AK058279 | Unknown expressed protein |
| 12 | 9 | 21375364 | AK103727 | Unknown expressed protein |
| 12 | 9 | 21475741 | AK066732 | Arabidopsis thaliana clone U19722 unknown protein (At4g26600) mRNA, complete cds.\|PLN |
| 12 | 9 | 21564624 | AK104344 | Oryza sativa (japonica cultivar-group) UGP mRNA for UDP-glucose pyrophosphorylase, complete cds.\|PLN |
| 12 | 9 | 21564624 | AK065780 | Oryza sativa (japonica cultivar-group) UGP mRNA for UDP-glucose pyrophosphorylase, complete cds.\|PLN |
| 12 | 9 | 21578554 | AK068240 | Arabidopsis thaliana At2g01350/F10A8.23 mRNA, complete cds.\|PLN |
| 12 | 9 | 21593505 | AK099586 | Arabidopsis thaliana unknown protein (At1g42550) mRNA, complete cds.\|PLN |
| 12 | 9 | 21610065 | AK070682 | Arabidopsis thaliana clone 105072 mRNA, complete sequence.\|PLN |
| 12 | 9 | 21711022 | AK108154 | Oryza sativa (indica cultivar-group) phytoene synthase radicle isoform mRNA, partial cds.\|PLN |
| 12 | 9 | 21733505 | AK101950 | Arabidopsis thaliana putative C2H2-type zinc finger protein (At2g02070) mRNA, complete cds.\|PLN |
| 12 | 9 | 21767688 | AK105452 | Unknown expressed protein |
| 12 | 9 | 21776324 | AK059357 | Arabidopsis thaliana At4g00370 mRNA for unknown protein, complete cds, clone: RAFL19-65-D16.\|PLN |
| 13 | 11 | 6309915 | AK060902 | O.sativa SC34 mRNA for tumor suppressor.\|PLN |
| 13 | 11 | 6408492 | AK101439 | Hordeum vulgare CC-NBS-LRR resistance protein MLA13 mRNA, complete cds; alternatively spliced.\|PLN |
| 13 | 11 | 6535559 | AK072355 | Hordeum vulgare mRNA for MLA6 protein.\|PLN |
| 13 | 11 | 6637634 | AK058632 | Unknown expressed protein |
| 13 | 11 | 6637634 | AK059194 | Unknown expressed protein |
| 13 | 11 | 6657155 | AK065723 | Oryza sativa (indica cultivar-group) mRNA for RPR1h, complete cds.\|PLN |
| 13 | 11 | 6674313 | AK064544 | Unknown expressed protein |
| 13 | 11 | 6683962 | AK102193 | Oryza sativa (japonica cultivar-group) NBS-LRR-like protein (YR5) mRNA, complete cds.\|PLN |

Supplementary Table 2. continued.

| GROUP | CHR | TU_START | GENENAME | DESCRIPTION |
|---|---|---|---|---|
| 13 | 11 | 6683962 | AK101375 | Oryza sativa (japonica cultivar-group) NBS-LRR-like protein (YR5) mRNA, complete cds.\|PLN |
| 13 | 11 | 6841012 | AK108061 | Unknown expressed protein |
| 13 | 11 | 6850392 | AK102213 | Unknown expressed protein |
| 13 | 11 | 6957275 | AK108047 | Unknown expressed protein |
| 13 | 11 | 7000165 | AK111558 | Arabidopsis thaliana putative receptor protein kinase (At1g28440) mRNA, complete cds.\|PLN |
| 13 | 11 | 7113638 | AK108764 | Arabidopsis thaliana unknown protein (At1g33420) mRNA, complete cds.\|PLN |
| 13 | 11 | 7181374 | AK100802 | Arabidopsis thaliana putative oligopeptide transporter protein (At1g59740) mRNA, complete cds.\|PLN |
| 13 | 11 | 7431982 | AK106466 | Unknown expressed protein |
| 14 | 12 | 135199 | AK068622 | Arabidopsis thaliana putative mitochondrial carrier protein (At2g35800) mRNA, complete cds.\|PLN |
| 14 | 12 | 219455 | AK065998 | Arabidopsis thaliana unknown protein (At2g29970) mRNA, complete cds.\|PLN |
| 14 | 12 | 225238 | AK071185 | Oryza sativa (japonica cultivar-group) mRNA for w-3 fatty acid desaturase, partial cds.\|PLN |
| 14 | 12 | 254447 | AK058259 | Unknown expressed protein |
| 14 | 12 | 254447 | AK058226 | Glycine max clathrin heavy chain mRNA, complete cds.\|PLN |
| 14 | 12 | 272612 | AK072914 | Arabidopsis thaliana CHUP1 mRNA for actin binding protein, complete cds.\|PLN |
| 14 | 12 | 295658 | AK107297 | Arabidopsis thaliana At2g38300 mRNA for unknown protein, complete cds, clone: RAFL17-35-G17.\|PLN |
| 14 | 12 | 300276 | AK067668 | Arabidopsis thaliana unknown protein (MOP10.6) mRNA, complete cds.\|PLN |
| 14 | 12 | 305336 | AK070979 | Arabidopsis thaliana At2g40270/T7M7.15 mRNA, complete cds.\|PLN |
| 14 | 12 | 319214 | AK102242 | Oryza sativa (japonica cultivar-group) ferritin (Fer2) mRNA, complete cds.\|PLN |

Supplementary Table 3. Groups of adjacent genes of correlated expression in SHZ-2 (8 gene window, p < 0.0005)

| GROUP | CHR | TU_START | KOME ID | DESCRIPTION |
|---|---|---|---|---|
| 1 | 2 | 4939675 | AK111066 | Unknown expressed protein |
| 1 | 2 | 4945755 | AK060691 | Arabidopsis thaliana putative APG protein (At4g26790) mRNA, complete cds.\|PLN |
| 1 | 2 | 4950266 | AK108100 | Unknown expressed protein |
| 1 | 2 | 4969015 | AK110945 | Gossypium hirsutum P-glycoprotein (CMDR1) mRNA, complete cds.\|PLN |
| 1 | 2 | 5039526 | AK099853 | Arabidopsis thaliana unknown protein (At3g01690) mRNA, complete cds.\|PLN |
| 1 | 2 | 5046514 | AK111037 | Arabidopsis thaliana clone 31276 mRNA, complete sequence.\|PLN |
| 1 | 2 | 5064204 | AK064615 | Arabidopsis thaliana AT5g15820/F14F8_200 mRNA, complete cds.\|PLN |
| 1 | 2 | 5122748 | AK101757 | Arabidopsis thaliana unknown protein (not annotated) mRNA, complete cds.\|PLN |
| 2 | 2 | 25524566 | AK106548 | Unknown expressed protein |
| 2 | 2 | 25535037 | AK107831 | Arabidopsis thaliana clone 31366 mRNA, complete sequence.\|PLN |
| 2 | 2 | 25537113 | AK060633 | Arabidopsis thaliana At1g15130/F9L1_7 mRNA, complete cds.\|PLN |
| 2 | 2 | 25564132 | AK111517 | Arabidopsis thaliana AT3g15470/MJK13_13 mRNA, complete cds.\|PLN |
| 2 | 2 | 25571859 | AK067293 | Unknown expressed protein |
| 2 | 2 | 25610124 | AK068974 | Arabidopsis thaliana PRL1-associated protein-like protein (At5g58720) mRNA, complete cds.\|PLN |
| 2 | 2 | 25616663 | AK105980 | Populus x canescens putative RING protein (RING) mRNA, complete cds.\|PLN |
| 2 | 2 | 25616663 | AK070367 | Populus x canescens putative RING protein (RING) mRNA, complete cds.\|PLN |
| 3 | 3 | 11167537 | AK068580 | Arabidopsis thaliana AT5g52840/MXC20_6 mRNA, complete cds.\|PLN |
| 3 | 3 | 11224297 | AK060813 | Arabidopsis thaliana At2g47250/T8I13.9 mRNA, complete cds.\|PLN |
| 3 | 3 | 11228773 | AK100769 | Arabidopsis thaliana putative protein (At5g46340) mRNA, complete cds.\|PLN |
| 3 | 3 | 11253332 | AK065474 | Arabidopsis thaliana unknown protein (At1g21680) mRNA, complete cds.\|PLN |
| 3 | 3 | 11258348 | AK069184 | Arabidopsis thaliana AT4g01940/T7B11_20 mRNA, complete cds.\|PLN |
| 3 | 3 | 11269852 | AK103334 | Oryza sativa (indica cultivar-group) invertase (INV) mRNA, complete cds.\|PLN |
| 3 | 3 | 11337942 | AK065115 | Spinacia oleracea chloroplast ribosomal protein S1 (rpS1) mRNA, complete cds.\|PLN |
| 3 | 3 | 11382493 | AK069044 | Arabidopsis thaliana clone 21701 mRNA, complete sequence.\|PLN |
| 4 | 4 | 30634571 | AK071230 | Arabidopsis thaliana At1g18900 mRNA for unknown protein, complete cds, clone: RAFL17-06-O22.\|PLN |
| 4 | 4 | 30650118 | AK109490 | Arabidopsis thaliana AT4g20040/F18F4_140 mRNA, complete cds.\|PLN |
| 4 | 4 | 30656477 | AK065653 | Arabidopsis thaliana At1g16320/F3O9_12 mRNA, complete cds.\|PLN |
| 4 | 4 | 30669771 | AK105086 | Unknown expressed protein |
| 4 | 4 | 30669771 | AK064783 | Arabidopsis thaliana At1g31070/F17F8_1 mRNA, complete cds.\|PLN |
| 4 | 4 | 30689127 | AK072123 | Arabidopsis thaliana putative potassium transporter (At2g35060) mRNA, complete cds.\|PLN |
| 4 | 4 | 30700031 | AK073265 | Arabidopsis thaliana unknown protein (At3g12790) mRNA, complete cds.\|PLN |

Supplementary Table 3.    continued…

| GROUP | CHR | TU_START | KOME ID | DESCRIPTION |
|---|---|---|---|---|
| 4 | 4 | 30752560 | AK058602 | Unknown expressed protein |
| 5 | 5 | 15709191 | AK111022 | Unknown expressed protein |
| 5 | 5 | 15839579 | AK111385 | Homo sapiens, KIAA0118 protein, clone MGC:16965 IMAGE:4342334, mRNA, complete cds.\|PRI |
| 5 | 5 | 15878730 | AK111106 | Unknown expressed protein |
| 5 | 5 | 15928267 | AK064169 | Arabidopsis thaliana clone 14329 mRNA, complete sequence.\|PLN |
| 5 | 5 | 15998886 | AK060058 | Unknown expressed protein |
| 5 | 5 | 16073482 | AK073748 | Unknown expressed protein |
| 5 | 5 | 16087885 | AK066614 | Arabidopsis thaliana clone 1391 mRNA, complete sequence.\|PLN |
| 5 | 5 | 16087885 | AK073761 | Arabidopsis thaliana clone 1391 mRNA, complete sequence.\|PLN |
| 5 | 5 | 16109937 | AK066689 | Arabidopsis thaliana putative protein (At5g58440) mRNA, complete cds.\|PLN |
| 5 | 5 | 16117370 | AK071850 | Hordeum vulgare dehydration-responsive AP2 domain transcriptional activator (DRF1.3) mRNA, complete cds.\|PLN |
| 6 | 5 | 27815824 | AK111480 | Arabidopsis thaliana putative protein (At5g44170) mRNA, complete cds.\|PLN |
| 6 | 5 | 27823563 | AK065766 | Oryza sativa hypothetical protein mRNA, partial cds.\|PLN |
| 6 | 5 | 27831280 | AK109344 | Arabidopsis thaliana At5g60860 mRNA for putative GTP-binding protein, complete cds, clone: RAFL19-94-D08.\|PLN |
| 6 | 5 | 27863654 | AK067090 | Solanum tuberosum mRNA for urease accessory protein G (ureG gene) isoform R3.\|PLN |
| 6 | 5 | 27921570 | AK099641 | Arabidopsis thaliana unknown protein (At5g46420) mRNA, complete cds.\|PLN |
| 6 | 5 | 27927209 | AK100389 | Oryza sativa (japonica cultivar-group) mRNA for putative mitogen-activated protein kinase wjumk1 (wjumk1 gene).\|PLN |
| 6 | 5 | 27927209 | AK061645 | Oryza sativa (japonica cultivar-group) mRNA for putative mitogen-activated protein kinase wjumk1 (wjumk1 gene).\|PLN |
| 6 | 5 | 27940571 | AK099052 | Arabidopsis thaliana putative translation initiation factor eIF3 (At4g20980) mRNA, complete cds.\|PLN |
| 6 | 5 | 27943924 | AK058475 | Arabidopsis thaliana AT5g64130/MHJ24_11 mRNA, complete cds.\|PLN |
| 7 | 6 | 3639786 | AK068738 | Arabidopsis thaliana unknown protein (At3g06430) mRNA, complete cds.\|PLN |
| 7 | 6 | 3648517 | AK105920 | Lotus japonicus mRNA for calcium-binding protein (cbp1 gene).\|PLN |
| 7 | 6 | 3688672 | AK065612 | Oryza sativa (japonica cultivar-group) OsRPT5b mRNA for 26S proteasome regulatory particle triple-A ATPase subunit5b, partial cds.\|PLN |
| 7 | 6 | 3772667 | AK065311 | Arabidopsis thaliana AT4g32150/F10N7_40 mRNA, complete cds.\|PLN |
| 7 | 6 | 3780123 | AK099635 | Unknown expressed protein |
| 7 | 6 | 3800201 | AK100481 | Arabidopsis thaliana unknown protein (At5g35200) mRNA, complete cds.\|PLN |
| 7 | 6 | 3918501 | AK066933 | Oryza sativa mRNA for vacuolar H+-pyrophosphatase, complete cds.\|PLN |
| 7 | 6 | 3923681 | AK065263 | Arabidopsis thaliana putative protein (At5g61250) mRNA, complete cds.\|PLN |
| 8 | 7 | 15067684 | AK070141 | Arabidopsis thaliana unknown protein (At3g24180) mRNA, complete cds.\|PLN |

Supplementary Table 3. continued…

| GROUP | CHR | TU_START | KOME ID | DESCRIPTION |
|---|---|---|---|---|
| 8 | 7 | 15202399 | AK074000 | Arabidopsis thaliana CIP7 mRNA for COP1-Interacting Protein 7, complete cds.\|PLN |
| 8 | 7 | 15235478 | AK100927 | Arabidopsis thaliana AT3g23280/K14B15_17 mRNA, complete cds.\|PLN |
| 8 | 7 | 15337218 | AK064872 | Arabidopsis thaliana AT5g17530/K10A8_10 mRNA, complete cds.\|PLN |
| 8 | 7 | 15357265 | AK072632 | Zea mays plasma membrane integral protein ZmPIP2-6 mRNA, complete cds.\|PLN |
| 8 | 7 | 15375221 | AK107700 | Zea mays plasma membrane integral protein ZmPIP2-6 mRNA, complete cds.\|PLN |
| 8 | 7 | 15404747 | AK103938 | Oryza sativa aquaporin (PIP2a) mRNA, complete cds.\|PLN |
| 8 | 7 | 15419975 | AK062392 | Arabidopsis thaliana unknown protein (At4g22310) mRNA, complete cds.\|PLN |
| 9 | 9 | 3566774 | AK065068 | Gene Spl7b regulating lesion formation in plant and utilization thereof.\|PAT |
| 9 | 9 | 3601703 | AK105777 | Arabidopsis thaliana At4g12730/T20K18_80 mRNA, complete cds.\|PLN |
| 9 | 9 | 3603479 | AK103826 | Arabidopsis thaliana putative protein (At5g48390) mRNA, complete cds.\|PLN |
| 9 | 9 | 3616690 | AK099919 | Arabidopsis thaliana putative protein (At5g48390) mRNA, complete cds.\|PLN |
| 9 | 9 | 3679109 | AK072805 | Gene of 36kDa protein appearing in iron-deficient barley root.\|PAT |
| 9 | 9 | 3692424 | AK063577 | Arabidopsis thaliana AT4g31460/F3L17_30 mRNA, complete cds.\|PLN |
| 9 | 9 | 3697771 | AK068062 | Arabidopsis thaliana At1g28060/F13K9_16 mRNA, complete cds.\|PLN |
| 9 | 9 | 3704719 | AK069757 | Oryza sativa mRNA for protein phosphatase 2A regulatory A subunit (RPA1 gene).\|PLN |
| 10 | 9 | 5984593 | AK102936 | Arabidopsis thaliana mRNA for struwwelpeter 1 protein (swp1 gene).\|PLN |
| 10 | 9 | 6141869 | AK069582 | Arabidopsis thaliana clone 108517 mRNA, complete sequence.\|PLN |
| 10 | 9 | 6189891 | AK059004 | Unknown expressed protein |
| 10 | 9 | 6192948 | AK110658 | Arabidopsis thaliana clone 35284 mRNA, complete sequence.\|PLN |
| 10 | 9 | 6198587 | AK072349 | Zea mays enhancer of polycomb-like protein (epc101) mRNA, complete cds.\|PLN |
| 10 | 9 | 6269965 | AK071578 | Unknown expressed protein |
| 10 | 9 | 6321038 | AK072114 | Lycopersicon esculentum mRNA for Na+/H+ antiporter (NHX2 gene).\|PLN |
| 10 | 9 | 6334602 | AK060090 | Prunus armeniaca AP2 domain containing protein (AP2DCP) mRNA, partial cds.\|PLN |
| 10 | 9 | 6351193 | AK106057 | Solanum tuberosum cip353 mRNA for AP2/ERF-domain protein, complete cds.\|PLN |
| 10 | 9 | 6386337 | AK059944 | Arabidopsis thaliana clone 4326 mRNA, complete sequence.\|PLN |
| 11 | 9 | 7262786 | AK069336 | Arabidopsis thaliana AT3g24120/MUJ8_3 mRNA, complete cds.\|PLN |
| 11 | 9 | 7818300 | AK064903 | Unknown expressed protein |
| 11 | 9 | 7872271 | AK100856 | Arabidopsis thaliana unknown protein (At5g51230:At5g51240) mRNA, complete cds.\|PLN |
| 11 | 9 | 7985887 | AK065670 | Zea mays SET domain protein 110 (sdg110) mRNA, complete cds.\|PLN |
| 11 | 9 | 8079080 | AK058754 | Unknown expressed protein |

Supplementary Table 3. continued…

| GROUP | CHR | TU_START | KOME ID | DESCRIPTION |
|---|---|---|---|---|
| 11 | 9 | 8098518 | AK062084 | Arabidopsis thaliana clone 37546 mRNA, complete sequence.\|PLN |
| 11 | 9 | 8098518 | AK063079 | Vigna unguiculata phosphatidic acid phosphatase beta mRNA, complete cds.\|PLN |
| 11 | 9 | 8112688 | AK102522 | Arabidopsis thaliana At1g60560/F8A5_10 mRNA, complete cds.\|PLN |
| 12 | 10 | 19386866 | AK062851 | Oryza sativa H protein subunit of glycine decarboxylase mRNA, complete cds.\|PLN |
| 12 | 10 | 19400516 | AK065934 | electron transfer flavoprotein-ubiquinone oxidoreductase [human, fetal liver, mRNA, 2124 nt].\|PRI |
| 12 | 10 | 19421118 | AK059055 | Unknown expressed protein |
| 12 | 10 | 19454864 | AK070811 | Arabidopsis thaliana AT5g23450/K19M13_8 mRNA, complete cds.\|PLN |
| 12 | 10 | 19463927 | AK107070 | Unknown expressed protein |
| 12 | 10 | 19517047 | AK068787 | Arabidopsis thaliana AT5g53570/MNC6_11 mRNA, complete cds.\|PLN |
| 12 | 10 | 19520805 | AK058780 | Rice mRNA for Cytochrome b5.\|PLN |
| 12 | 10 | 19529747 | AK102374 | Unknown expressed protein |
| 13 | 11 | 5016540 | AK064282 | Arabidopsis thaliana putative histone deacetylase (RPD3A) mRNA, complete cds.\|PLN |
| 13 | 11 | 5141584 | AK063427 | Unknown expressed protein |
| 13 | 11 | 5167147 | AK063307 | Stevia rebaudiana UDP-glucosyltransferase mRNA, complete cds.\|PLN |
| 13 | 11 | 5171128 | AK102530 | Arabidopsis thaliana clone 37855 mRNA, complete sequence.\|PLN |
| 13 | 11 | 5246760 | AK071941 | Unknown expressed protein |
| 13 | 11 | 5265073 | AK099643 | Oryza sativa (japonica cultivar-group) mRNA for ADP glucose pyrophosphorylase large subunit, complete cds.\|PLN |
| 13 | 11 | 5270714 | AK100488 | Triticum aestivum (wali7) mRNA, 3 end, partial cds.\|PLN |
| 13 | 11 | 5361291 | AK109707 | Arabidopsis thaliana clone 22350 mRNA, complete sequence.\|PLN |
| 13 | 11 | 5361291 | AK108795 | Arabidopsis thaliana clone 22350 mRNA, complete sequence.\|PLN |
| 14 | 11 | 6535559 | AK072355 | Hordeum vulgare mRNA for MLA6 protein.\|PLN |
| 14 | 11 | 6637634 | AK058632 | Unknown expressed protein |
| 14 | 11 | 6637634 | AK059194 | Unknown expressed protein |
| 14 | 11 | 6657155 | AK065723 | Oryza sativa (indica cultivar-group) mRNA for RPR1h, complete cds.\|PLN |
| 14 | 11 | 6674313 | AK064544 | Unknown expressed protein |
| 14 | 11 | 6683962 | AK102193 | Oryza sativa (japonica cultivar-group) NBS-LRR-like protein (YR5) mRNA, complete cds.\|PLN |
| 14 | 11 | 6683962 | AK101375 | Oryza sativa (japonica cultivar-group) NBS-LRR-like protein (YR5) mRNA, complete cds.\|PLN |
| 14 | 11 | 6841012 | AK108061 | Unknown expressed protein |
| 14 | 11 | 6850392 | AK102213 | Unknown expressed protein |
| 15 | 12 | 24180206 | AK071898 | Arabidopsis thaliana putative protein (At4g37590) mRNA, complete cds.\|PLN |
| 15 | 12 | 24208080 | AK065030 | Arabidopsis thaliana putative membrane transporter (At2g26510; T9J22.18) mRNA, complete cds.\|PLN |
| 15 | 12 | 24235702 | AK071424 | Unknown expressed protein |
| 15 | 12 | 24420981 | AK069697 | Arabidopsis thaliana protein kinase-like protein (At4g33950) mRNA, complete cds.\|PLN |

Supplementary Table 3. continued.

| GROUP | CHR | TU_START | KOME ID | DESCRIPTION |
|---|---|---|---|---|
| 15 | 12 | 24447113 | AK066259 | Medicago truncatula type IIB calcium ATPase MCA5 (MCA5) mRNA, complete cds.\|PLN |
| 15 | 12 | 24598131 | AK073627 | Triticum aestivum adenine phosphoribosyltransferase form 1 (APT1) mRNA, complete cds.\|PLN |
| 15 | 12 | 24659642 | AK070028 | Unknown expressed protein |
| 15 | 12 | 24833595 | AK064058 | Unknown expressed protein |

Supplementary Table 4. The 84 genes in the GR978-RCE Groups associated with Bl-QTLs

| KOME ID AND DESCRIPTION | GROUP | CHR | TU_START |
|---|---|---|---|
| AK059010 [2\|21607719] Oryza sativa Cyt-P450 monooxygenase (PM-II) mRNA, complete cds.\|PLN | 4 | 2 | 21607719 |
| AK060161 [2\|21543249] Nicotiana plumbaginifolia partial mRNA for RNA Binding Protein 47 (rbp47 gene).\|PLN | 4 | 2 | 21543249 |
| AK060722 [2\|21635489] Oryza sativa Cyt-P450 monooxygenase (PM-II) mRNA, complete cds.\|PLN | 4 | 2 | 21635489 |
| AK066007 [2\|21926092] Arabidopsis thaliana clone 142216 mRNA, complete sequence.\|PLN | 4 | 2 | 21926092 |
| AK066393 [2\|21508582] Arabidopsis thaliana unknown protein (At1g07700) mRNA, complete cds.\|PLN | 4 | 2 | 21508582 |
| AK068282 [2\|21506959] Unknown expressed protein | 4 | 2 | 21506959 |
| AK068310 [2\|21710122] Oryza sativa OsDTC1 mRNA for putative diterpene cyclase, complete cds.\|PLN | 4 | 2 | 21710122 |
| AK069840 [2\|21514322] Arabidopsis thaliana unknown protein (At1g47380) mRNA, complete cds.\|PLN | 4 | 2 | 21514322 |
| AK070167 [2\|21755452] Triticum aestivum N-1 mRNA for cytochrome P450, complete cds.\|PLN | 4 | 2 | 21755452 |
| AK099415 [2\|21955857] Oryza sativa OsMST1 mRNA for monosaccharide transporter 1, complete cds.\|PLN | 4 | 2 | 21955857 |
| AK101003 [2\|21849467] Oryza sativa Cyt-P450 monooxygenase (PM-II) mRNA, complete cds.\|PLN | 4 | 2 | 21849467 |
| AK105793 [2\|21532878] Arabidopsis thaliana clone 111056 mRNA, complete sequence.\|PLN | 4 | 2 | 21532878 |
| AK105913 [2\|21685415] Oryza sativa Cyt-P450 monooxygenase (PM-II) mRNA, complete cds.\|PLN | 4 | 2 | 21685415 |
| AK107418 [2\|21733281] Solanum tuberosum mRNA for cytochrome P450 (CYP71D4 gene).\|PLN | 4 | 2 | 21733281 |
| AK109940 [2\|21872680] Oryza sativa early proembryo mRNA, complete sequence.\|PLN | 4 | 2 | 21872680 |
| AK058617 [4\|20139031] Unknown expressed protein | 6 | 4 | 20139031 |
| AK062016 [4\|20020484] C.sinensis mRNA for non-photosynthetic ferredoxin.\|PLN | 6 | 4 | 20020484 |
| AK070396 [4\|19965028] Unknown expressed protein | 6 | 4 | 19965028 |
| AK073418 [4\|20052966] Arabidopsis thaliana Unknown protein (At2g27460; F10A12.14) mRNA, complete cds.\|PLN | 6 | 4 | 20052966 |
| AK100104 [4\|20098163] Unknown expressed protein | 6 | 4 | 20098163 |
| AK102032 [4\|19993750] Arabidopsis thaliana unknown protein (At3g48410) mRNA, complete cds.\|PLN | 6 | 4 | 19993750 |
| AK108077 [4\|19935234] Zea mays ZmRCP2 mRNA for root cap protein 2, complete cds.\|PLN | 6 | 4 | 19935234 |
| AK108223 [4\|20013514] Arabidopsis thaliana AT3g52740/F3C22_140 mRNA, complete cds.\|PLN | 6 | 4 | 20013514 |
| AK108347 [4\|20046859] Unknown expressed protein | 6 | 4 | 20046859 |
| AK109638 [4\|20161656] Unknown expressed protein | 6 | 4 | 20161656 |
| AK110902 [4\|20027785] Arabidopsis thaliana putative beta-1,3-glucanase (At2g27500) mRNA, complete cds.\|PLN | 6 | 4 | 20027785 |
| AK111482 [4\|20033597] Unknown expressed protein | 6 | 4 | 20033597 |
| AK062635 [6\|24267674] Unknown expressed protein | 9 | 6 | 24267674 |
| AK062719 [6\|24303827] Unknown expressed protein | 9 | 6 | 24303827 |
| AK063128 [6\|24299115] Unknown expressed protein | 9 | 6 | 24299115 |
| AK066754 [6\|24303827] Nicotiana tabacum mRNA for CND41, chloroplast nucleoid DNA binding protein with aspartic protease activity, complete cds.\|PLN | 9 | 6 | 24303827 |
| AK067228 [6\|24232599] Arabidopsis thaliana At4g17150 mRNA for unknown protein, complete cds, clone: RAFL21-18-K13.\|PLN | 9 | 6 | 24232599 |

Supplementary Table 4. continued…

| KOME ID AND DESCRIPTION | GROUP | CHR | TU_START |
|---|---|---|---|
| AK067236 [6\|24314444] Nicotiana tabacum mRNA for microtubule-associated protein MAP65-1a (map65-1a gene).\|PLN | 9 | 6 | 24314444 |
| AK072533 [6\|24228506] Danio rerio fast skeletal muscle troponin C (tnnc) mRNA, complete cds.\|VRT | 9 | 6 | 24228506 |
| AK072708 [6\|24339229] Z.mays mRNA for polygalacturonase (clone PG2).\|PLN | 9 | 6 | 24339229 |
| AK107800 [6\|24328419] Unknown expressed protein | 9 | 6 | 24328419 |
| AK109238 [6\|24319205] Unknown expressed protein | 9 | 6 | 24319205 |
| AK109906 [6\|24287219] Zea mays seven transmembrane protein Mlo7 mRNA, complete cds.\|PLN | 9 | 6 | 24287219 |
| AK111644 [6\|24354750] Avena sativa victorin binding protein mRNA, complete cds.\|PLN | 9 | 6 | 24354750 |
| AK064568 [7\|21389808] Unknown expressed protein | 10 | 7 | 21389808 |
| AK068361 [7\|21465080] Arabidopsis thaliana Unknown protein (At5g54680) mRNA, complete cds.\|PLN | 10 | 7 | 21465080 |
| AK100722 [7\|21357958] Arabidopsis thaliana mRNA for receptor-like protein kinase.\|PLN | 10 | 7 | 21357958 |
| AK102941 [7\|21406209] Arabidopsis thaliana At4g23180 mRNA for putative receptor-like protein kinase 4 (RLK4), complete cds, clone: RAFL17-45-D12.\|PLN | 10 | 7 | 21406209 |
| AK105128 [7\|21393500] Arabidopsis thaliana At4g23140 mRNA for putative receptor-like protein kinase 5 (RLK5), complete cds, clone: RAFL19-72-N19.\|PLN | 10 | 7 | 21393500 |
| AK105710 [7\|21518679] Unknown expressed protein | 10 | 7 | 21518679 |
| AK105928 [7\|21297936] Arabidopsis thaliana AT3g08780/F17O14_25 mRNA, complete cds.\|PLN | 10 | 7 | 21297936 |
| AK110523 [7\|21389808] Unknown expressed protein | 10 | 7 | 21389808 |
| AK111550 [7\|21383094] Arabidopsis thaliana At4g23180 mRNA for putative receptor-like protein kinase 4 (RLK4), complete cds, clone: RAFL17-45-D12.\|PLN | 10 | 7 | 21383094 |
| AK111865 [7\|21432948] Phaseolus vulgaris receptor-like protein kinase homolog RK20-1 mRNA, complete cds.\|PLN | 10 | 7 | 21432948 |
| AK059689 [9\|11649601] Arabidopsis thaliana arginine methyltransferase pam1 (At4g29510) mRNA, complete cds.\|PLN | 12 | 9 | 11649601 |
| AK063411 [9\|10924313] Zea mays transposon Doppia transposase DOPD and transposase DOPA mRNA, complete cds.\|PLN | 12 | 9 | 10924313 |
| AK065335 [9\|11183249] Arabidopsis thaliana putative bystin (At1g31660) mRNA, complete cds.\|PLN | 12 | 9 | 11183249 |
| AK068895 [9\|10844762] Unknown expressed protein | 12 | 9 | 10844762 |
| AK069410 [9\|11321555] Unknown expressed protein | 12 | 9 | 11321555 |
| AK071224 [9\|10875106] Arabidopsis thaliana unknown protein (At5g55810) mRNA, complete cds.\|PLN | 12 | 9 | 10875106 |
| AK072578 [9\|11395157] Unknown expressed protein | 12 | 9 | 11395157 |
| AK100597 [9\|10861935] Triticum aestivum Sec61p (sec61) mRNA, complete cds.\|PLN | 12 | 9 | 10861935 |
| AK101995 [9\|10869048] Zea mays histone acetyl transferase (hac106) mRNA, complete cds.\|PLN | 12 | 9 | 10869048 |
| AK105685 [9\|11333855] Oryza sativa mRNA for OsD305, complete cds.\|PLN | 12 | 9 | 11333855 |
| AK065475 [9\|15109795] Arabidopsis thaliana unknown protein (At4g31820) mRNA, complete cds.\|PLN | 13 | 9 | 15109795 |
| AK068557 [9\|15011730] Unknown expressed protein | 13 | 9 | 15011730 |
| AK068795 [9\|15072960] Arabidopsis thaliana clone 108517 mRNA, complete sequence.\|PLN | 13 | 9 | 15072960 |
| AK072629 [9\|15100475] Arabidopsis thaliana clone 3836 mRNA, complete sequence.\|PLN | 13 | 9 | 15100475 |
| AK073786 [9\|15037373] Arabidopsis thaliana clone 108517 mRNA, complete sequence.\|PLN | 13 | 9 | 15037373 |
| AK099513 [9\|15107602] Deschampsia antarctica clone Dacor 1.0 polyubiquitin 2 mRNA, complete cds.\|PLN | 13 | 9 | 15107602 |

Supplementary Table 4. continued.

| KOME ID AND DESCRIPTION | GROUP | CHR | TU_START |
|---|---|---|---|
| AK101900 [9\|15198704] Nicotiana tabacum mRNA for hydroxycinnamoyl transferase (hct gene).\|PLN | 13 | 9 | 15198704 |
| AK107198 [9\|15022954] Hordeum vulgare cultivar Orthega cinnamoyl CoA reductase mRNA, complete cds.\|PLN | 13 | 9 | 15022954 |
| AK109320 [9\|15041211] Arabidopsis thaliana clone 108517 mRNA, complete sequence.\|PLN | 13 | 9 | 15041211 |
| AK111264 [9\|15103705] Arabidopsis thaliana Cu-chaperone (COX17) mRNA, complete cds.\|PLN | 13 | 9 | 15103705 |
| AK066177 [12\|11348497] Arabidopsis thaliana At1g67840/F12A21_3 mRNA, complete cds.\|PLN | 15 | 12 | 11348497 |
| AK068266 [12\|11318956] Oryza sativa (japonica cultivar-group) mRNA for the small subunit of ribulose-1,5-bisphosphate carboxylase, complete cds, clone pOSSS1139.\|PLN | 15 | 12 | 11318956 |
| AK099574 [12\|11275929] Oryza sativa (japonica cultivar-group) mRNA for the small subunit of ribulose-1,5-bisphosphate carboxylase, complete cds, clone pOSSS2106.\|PLN | 15 | 12 | 11275929 |
| AK111535 [12\|11418673] Arabidopsis thaliana unknown protein (At2g47790/F17A22.18) mRNA, complete cds.\|PLN | 15 | 12 | 11418673 |
| AK060063 [12\|19088840] Arabidopsis thaliana clone 30535 mRNA, complete sequence.\|PLN | 16 | 12 | 19088840 |
| AK067767 [12\|18905820] Arabidopsis thaliana clone RAFL14-85-F19 (R20222) unknown protein (At5g55610) mRNA, complete cds.\|PLN | 16 | 12 | 18905820 |
| AK068908 [12\|18846515] Arabidopsis thaliana putative translation initiation factor EIF-2B alpha subunit (At1g72340) mRNA, complete cds.\|PLN | 16 | 12 | 18846515 |
| AK069786 [12\|19118282] Arabidopsis thaliana At4g08790/T32A17_100 mRNA, complete cds.\|PLN | 16 | 12 | 19118282 |
| AK070044 [12\|18880405] Arabidopsis thaliana clone RAFL15-27-J23 (R20860) unknown protein (At1g69390) mRNA, complete cds.\|PLN | 16 | 12 | 18880405 |
| AK071243 [12\|18846515] Arabidopsis thaliana putative translation initiation factor EIF-2B alpha subunit (At1g72340) mRNA, complete cds.\|PLN | 16 | 12 | 18846515 |
| AK073582 [12\|18899996] Homo sapiens, clone MGC:8856 IMAGE:3890907, mRNA, complete cds.\|PRI | 16 | 12 | 18899996 |
| AK105366 [12\|18822849] Arabidopsis thaliana putative transcription co-activator (SYT3) mRNA, complete cds.\|PLN | 16 | 12 | 18822849 |
| AK106653 [12\|19105480] Lycopersicon esculentum mRNA for cyclin A2 (CycA2 gene).\|PLN | 16 | 12 | 19105480 |
| AK110307 [12\|18725053] Triticum aestivum stripe rust resistance protein Yr10 (Yr10) mRNA, complete cds.\|PLN | 16 | 12 | 18725053 |
| AK111001 [12\|19114474] Arabidopsis thaliana putative 3-phosphoserine phosphatase (At1g18640) mRNA, complete cds.\|PLN | 16 | 12 | 19114474 |

Supplementary Table 5. Ratios tested for association analysis of GR978 DEGs in all RCEs for the genotypes GR978 and SHZ-2, done per chromosome (all not significant using Fisher exact test)

| | COUNTS | | | |
|---|---|---|---|---|
| | ACTUAL | | EXPECTED | |
| CHR | DEG AND RCE | NOT DEG AND RCE | DEG AND RCE | NOT DEG AND RCE |
| *GR978 RCE* | | | | |
| 1 | | 10 | 0 | 10 |
| 2 | | 35 | 1 | 34 |
| 3 | 1 | 12 | 0 | 13 |
| 4 | | 12 | 0 | 12 |
| 5 | | 10 | 0 | 10 |
| 6 | 1 | 21 | 1 | 21 |
| 7 | | 10 | 0 | 10 |
| 8 | | 12 | 0 | 12 |
| 9 | | 20 | 1 | 19 |
| 10 | | 0 | 0 | 0 |
| 11 | | 10 | 0 | 10 |
| 12 | | 21 | 1 | 20 |
| *GR978-SHZ-2 RCE* | | | | |
| 1 | | 10 | 0 | 10 |
| 2 | | 10 | 0 | 10 |
| 3 | 1 | 33 | 1 | 33 |
| 4 | 2 | 22 | 1 | 23 |
| 5 | 1 | 20 | 1 | 20 |
| 6 | | 0 | 0 | 0 |
| 7 | 2 | 8 | 0 | 10 |
| 8 | | 10 | 0 | 10 |
| 9 | | 11 | 0 | 11 |
| 10 | | 0 | 0 | 0 |
| 11 | | 16 | 1 | 15 |
| 12 | | 10 | 0 | 10 |
| *SHZ-2 RCE* | | | | |
| 1 | | 0 | 0 | 0 |
| 2 | 1 | 15 | 1 | 15 |
| 3 | | 8 | 0 | 8 |
| 4 | | 8 | 0 | 8 |
| 5 | 1 | 18 | 1 | 18 |
| 6 | | 8 | 0 | 8 |
| 7 | 1 | 7 | 0 | 8 |
| 8 | | 0 | 0 | 0 |
| 9 | 2 | 24 | 1 | 25 |
| 10 | 2 | 6 | 0 | 8 |
| 11 | 1 | 17 | 1 | 17 |
| 12 | | 8 | 0 | 8 |

Supplementary Table 6. Ratios tested for association analysis of GR978-SHZ2 DEGs in all RCEs for the genotypes GR978 and SHZ-2, done per chromosome (all not significant using Fisher exact test)

| | COUNTS | | | |
|---|---|---|---|---|
| | ACTUAL | | EXPECTED | |
| CHR | DEG AND RCE | NOT DEG AND RCE | DEG AND RCE | NOT DEG AND RCE |
| *GR978 RCE* | | | | |
| 1 | | 10 | 0 | 10 |
| 2 | | 10 | 0 | 10 |
| 3 | | 34 | 0 | 34 |
| 4 | | 24 | 0 | 24 |
| 5 | | 21 | 0 | 21 |
| 6 | | 0 | 0 | 0 |
| 7 | | 10 | 0 | 10 |
| 8 | | 10 | 0 | 10 |
| 9 | | 11 | 0 | 11 |
| 10 | | 0 | 0 | 0 |
| 11 | | 16 | 0 | 16 |
| 12 | | 10 | 0 | 10 |
| *GR978-SHZ-2 RCE* | | | | |
| 1 | | 10 | 0 | 10 |
| 2 | | 35 | 0 | 35 |
| 3 | | 13 | 0 | 13 |
| 4 | 1 | 11 | 0 | 12 |
| 5 | 1 | 9 | 0 | 10 |
| 6 | | 22 | 0 | 22 |
| 7 | | 10 | 0 | 10 |
| 8 | | 12 | 0 | 12 |
| 9 | | 20 | 0 | 20 |
| 10 | | 0 | 0 | 0 |
| 11 | | 10 | 0 | 10 |
| 12 | | 21 | 0 | 21 |
| *SHZ-2 RCE* | | | | |
| 1 | | 0 | 0 | 0 |
| 2 | | 16 | 0 | 16 |
| 3 | | 8 | 0 | 8 |
| 4 | | 8 | 0 | 8 |
| 5 | | 19 | 0 | 19 |
| 6 | | 8 | 0 | 8 |
| 7 | | 8 | 0 | 8 |
| 8 | | 0 | 0 | 0 |
| 9 | 1 | 25 | 0 | 26 |
| 10 | | 8 | 0 | 8 |
| 11 | | 18 | 0 | 18 |
| 12 | | 8 | 0 | 8 |

Supplementary Table 7. Ratios tested for association analysis of SHZ2 DEGs in all RCEs for the genotypes GR978 and SHZ-2, done per chromosome (all not significant using Fisher exact test)

| | COUNTS | | | |
|---|---|---|---|---|
| | ACTUAL | | EXPECTED | |
| CHR | DEG AND RCE | NOT DEG AND RCE | DEG AND RCE | NOT DEG AND RCE |
| *GR978 RCE* | | | | |
| 1 | 1 | 9 | 2 | 8 |
| 2 | 8 | 27 | 7 | 28 |
| 3 | 3 | 10 | 3 | 10 |
| 4 | 5 | 7 | 2 | 10 |
| 5 | 4 | 6 | 2 | 8 |
| 6 | 8 | 14 | 5 | 17 |
| 7 | 2 | 8 | 2 | 8 |
| 8 | | 12 | 2 | 10 |
| 9 | 6 | 14 | 4 | 16 |
| 10 | | 0 | 0 | 0 |
| 11 | 1 | 9 | 2 | 8 |
| 12 | 8 | 13 | 4 | 17 |
| *GR978-SHZ-2 RCE* | | | | |
| 1 | 1 | 9 | 2 | 8 |
| 2 | 3 | 7 | 2 | 8 |
| 3 | 12 | 22 | 7 | 27 |
| 4 | 7 | 17 | 5 | 19 |
| 5 | 8 | 13 | 5 | 16 |
| 6 | | 0 | 0 | 0 |
| 7 | 2 | 8 | 2 | 8 |
| 8 | 2 | 8 | 2 | 8 |
| 9 | 4 | 7 | 2 | 9 |
| 10 | | 0 | 0 | 0 |
| 11 | 2 | 14 | 3 | 13 |
| 12 | 4 | 6 | 2 | 8 |
| *SHZ-2 RCE* | | | | |
| 1 | | 0 | 0 | 0 |
| 2 | | 16 | 3 | 13 |
| 3 | 3 | 5 | 2 | 6 |
| 4 | 2 | 6 | 2 | 6 |
| 5 | 7 | 12 | 4 | 15 |
| 6 | 3 | 5 | 2 | 6 |
| 7 | 2 | 6 | 2 | 6 |
| 8 | | 0 | 0 | 0 |
| 9 | 7 | 19 | 5 | 21 |
| 10 | 1 | 7 | 1 | 7 |
| 11 | 4 | 14 | 4 | 14 |
| 12 | 6 | 2 | 2 | 6 |

Supplementary Table 8. Ratios tested for association analysis of all DEGs for the genotypes GR978 and SHZ-2 in all Bl-QTLs, done per chromosome (all not significant using Fisher exact test)

| | COUNTS | | | |
|---|---|---|---|---|
| | ACTUAL | | EXPECTED | |
| CHR | DEG IN QTL | NOT DEG IN QTL | DEG IN QTL | NOT DEG IN QTL |
| *GR978DEGs* | | | | |
| 1 | 67 | 1515 | 68 | 1514 |
| 2 | 16 | 365 | 15 | 366 |
| 3 | 15 | 454 | 14 | 455 |
| 4 | 16 | 473 | 18 | 471 |
| 5 | 9 | 121 | 5 | 125 |
| 6 | 23 | 541 | 22 | 542 |
| 7 | 37 | 879 | 36 | 880 |
| 8 | 8 | 227 | 8 | 227 |
| 9 | 14 | 698 | 18 | 694 |
| 10 | 4 | 89 | 3 | 90 |
| 11 | 8 | 250 | 9 | 249 |
| 12 | 9 | 349 | 13 | 345 |
| *GR978-SHZ-2 DEGS* | | | | |
| 1 | 5 | 1577 | 5 | 1577 |
| 2 | 3 | 378 | 2 | 379 |
| 3 | 1 | 468 | 2 | 467 |
| 4 | 1 | 488 | 2 | 487 |
| 5 | | 130 | 1 | 129 |
| 6 | 3 | 561 | 1 | 563 |
| 7 | 4 | 912 | 6 | 910 |
| 8 | 1 | 234 | 1 | 234 |
| 9 | 2 | 710 | 3 | 709 |
| 10 | | 93 | 0 | 93 |
| 11 | 1 | 257 | 1 | 257 |
| 12 | | 358 | 1 | 357 |
| *SHZ-2 DEGs* | | | | |
| 1 | 333 | 1249 | 332 | 1250 |
| 2 | 68 | 313 | 78 | 303 |
| 3 | 103 | 366 | 100 | 369 |
| 4 | 105 | 384 | 99 | 390 |
| 5 | 29 | 101 | 28 | 102 |
| 6 | 109 | 455 | 116 | 448 |
| 7 | 168 | 748 | 182 | 734 |
| 8 | 37 | 198 | 45 | 190 |
| 9 | 144 | 568 | 144 | 568 |
| 10 | 12 | 81 | 17 | 76 |
| 11 | 56 | 202 | 52 | 206 |
| 12 | 73 | 285 | 72 | 286 |

### SUPPLEMENTARY PROGRAM/SCRIPT CODES

1. SQL Script used for flagging bad spots in the gene expression data output from Agilent FE software, using a data table name 'data_source'.

```
SELECT 22k_annot.ID AS UID, `data_source`.gProcessedSignal AS IA,
`data_source`.rProcessedSignal AS IB, 22k_annot.Row AS R,
22k_annot.Col AS C, 22k_annot.Row AS MR, 22k_annot.Col AS MC,
If((gIsSaturated=0) And (gIsFeatNonUnifOL=0) And (gIsBGNonUnifOL=0)
And (gIsFeatPopnOL=0) And (gIsBGPopnOL=0) And (gIsPosAndSignif=1)
And (gIsWellAboveBG=1),'C','X') AS FlagA, If(rIsSaturated=0 And
rIsFeatNonUnifOL=0 And rIsBGNonUnifOL=0 And rIsFeatPopnOL=0 And
rIsBGPopnOL=0 And rIsPosAndSignif=1 And rIsWellAboveBG=1,'C','X') AS
FlagB, `data_source`.gMedianSignal AS MedA,
`data_source`.rMedianSignal AS MedB
FROM `data_source` INNER JOIN 22k_annot ON `data_source`.ID =
22k_annot.ID
WHERE (((22k_annot.ControlType)=0));
```

2. R script for MAANOVA analysis (template)

```
rm(list=ls(all=T))

# load library and read in data
library(maanova)

# read data
shz.raw <- read.madata("g978_24h.txt", designfile="
g978_design.txt",
       genename=2, pmt=7, spotflag=F)

shz <- createData(shz.raw, n.rep=1, avgreps=0)

# make several model objects and fit ANOVA

# full model. var x trt interaction - for split plot design
model.full.mix <- makeModel(data=shz,
      formula=~Rep+Var+Rep:Var+Trt+Var:Trt,
          random=~Rep+Rep:Var)
# fit anova
anova.full.mix <- fitmaanova(shz, model.full.mix)

# no interaction model
model.noint.mix <- makeModel(data=shz,
      formula=~Rep+Var+Rep:Var+Trt,
          random=~Rep+Rep:Var)
# fit anova
anova.noint.mix <- fitmaanova(shz, model.noint.mix)
#######

# use model without interaction to test var and trt effect
```

```
ftest.var.mix <- matest(data=shz, model=model.noint.mix,
term="Var",
      n.perm=100, test.method=c(1,0,1,1), nnodes=16)

# volcano plot
pdf(file="g978-24_Var.pdf");
idx.Var.mix <- volcano(ftest.var.mix, title="Var test - mixed
model")
graphics.off();

# var mean effect for all genes
write.table(anova.noint.mix$Var,"g978-
24_var.txt",append=FALSE,sep="\t",
  quote=FALSE,row.names=shz$genename,
col.names=anova.noint.mix$Var.level)

# genes with sig Var
gene.sig.V <- shz$genename[idx.Var.mix$idx.Fs]
prob.sig.V <- ftest.var.mix$Fs$Pvalperm[idx.Var.mix$idx.Fs]
nrows<-NROW(gene.sig.V)
sig.V<- matrix(0,nrows,2)
sig.V[,1]<-gene.sig.V
sig.V[,2]<-prob.sig.V

 write.table(sig.V,"g978-24_sigV.txt",append=FALSE,sep="\t",
   quote=FALSE,row.names=FALSE,col.names=FALSE)

ftest.trt.mix <- matest(data=shz, model=model.noint.mix,
term="Trt",
      n.perm=100, test.method=c(1,0,1,1), nnodes=16)

# volcano plot
pdf(file="g978-24_trt.pdf");
idx.trt.mix <- volcano(ftest.trt.mix, title="Trt test - mixed
model")
graphics.off();

# trt mean effect for all genes
write.table(anova.noint.mix$Trt,"g978-
24_trt.txt",append=FALSE,sep="\t",
  quote=FALSE,row.names=shz$genename,
col.names=anova.noint.mix$Trt.level)

# genes with sig trt
gene.sig.T <- shz$genename[idx.trt.mix$idx.Fs]
prob.sig.T <- ftest.trt.mix$Fs$Pvalperm[idx.trt.mix$idx.Fs]
nrows<-NROW(gene.sig.T)
sig.T<- matrix(0,nrows,2)
sig.T[,1]<-gene.sig.T
sig.T[,2]<-prob.sig.T
 write.table(sig.T,"g978-24_sigT.txt",append=FALSE,sep="\t",
   quote=FALSE,row.names=FALSE,col.names=FALSE)
```

```
# test interaction effect
ftest.VxT.mix <- matest(data=shz, model=model.full.mix,
term="Var:Trt",n.perm=100,
      test.method=c(1,0,1,1), nnodes=16)

# volcano plot
pdf(file="g978-24_VxT.pdf");
idx.VxT.mix <- volcano(ftest.VxT.mix, title="Var x Trt test -
mixed model")
graphics.off();

# varxt mean effect for all genes
write.table(anova.full.mix$"Var:Trt","g978-
24_VarxTrt.txt",append=FALSE,sep="\t",
  quote=FALSE,row.names=shz$genename,
col.names=anova.full.mix$"Var:Trt.level")

# genes with sig VarxT
gene.sig.VxT <- shz$genename[idx.VxT.mix$idx.Fs]
prob.sig.VxT <- ftest.VxT.mix$Fs$Pvalperm[idx.VxT.mix$idx.Fs]
nrows<-NROW(gene.sig.VxT)
sig.VxT<- matrix(0,nrows,2)
sig.VxT[,1]<-gene.sig.VxT
sig.VxT[,2]<-prob.sig.VxT

write.table(sig.VxT,"g978-
24_sigVxT.txt",append=FALSE,sep="\t",
 quote=FALSE,row.names=FALSE,col.names=FALSE)

save.image("g978-24.RData")
```

3. Perl script implementing the Spellman-Rubin correlation analysis

```
#!/usr/bin/perl

use strict;
my $line = 0;

srand();

my %mod;
while ($ARGV[0] =~ "-") {
    my $mod = shift;
    my @mod = split(/\=/, $mod);
    $mod{$mod[0]} = $mod[1];
}

if ((! $mod{'-pval'} || ! $mod{'-pfile'}) andand ! $mod{'-
iter'}) {
    die "missing arguments\n";
}

my $coords = shift;
```

```
my $data = shift;
#my $hist = shift;
if (! $data) {die "<script> [coords] [data] \n";}
open (COORD, "$coords");
open (DATA, $data);

my %chr;
my %coord;
while (<COORD>) {
    chomp;
    my @line = split(/\t/, $_);
    #print "@line\n";
    $chr{$line[0]} = $line[1];
    $coord{$line[0]} = $line[2];
}

my %data;
my %line_data;
my %cg;
my %chrl;
my @order;
my @exclude;
my $header;
while (<DATA>) {
    if ($line > 0) {
      s/\r//g;
      chomp;
      my $string = $_;
      my @data = split(/\t/, $string);
      my @name = @data[0..1];
      @data = @data[2..(@data -1)];
      @{$data{$line}} = @data;
      $line_data{$line} = $string;
      $chrl{$line} = $chr{$name[0]};
      push (@order, $line);
    }
    else {
      $header = $_;
    }
    $line++;
    #print "line $line\n";
}

my %hist;
my $window = 10;
my $iterations = $mod{'-iter'};
my $min_p_val = $mod{'-pval'};
if ($mod{'-iter'}) {
    my $p_arrayref = prob_group();
    my $string = join("\t", @$p_arrayref);
    print "$string\n"; #redirect output to a pfile for later
use...
}
elsif ($mod{'-pval'} andand $mod{'-pfile'}) {
```

```
    open (P, "$mod{'-pfile'}");
    my $sig = "$data.p_$mod{'-pval'}";
    open (SIG, ">$sig");
    print SIG "$header";

    my $bin = "$data.bin_p_$mod{'-pval'}";
    open (BIN, ">$bin");

    my $p_arrayref;
    while (<P>) {
      chomp;
      push (@$p_arrayref, (split(/\t/, $_)));
    }

    @$p_arrayref = sort bynum (@$p_arrayref);
    my $cut_off_index = int (@$p_arrayref * $mod{'-pval'});
    my $cut_off = $$p_arrayref[(@$p_arrayref - $cut_off_index)
-1];

    my $corr_hashref = corr_hash(\%data, \@order);

    my $pi;
    for (my $i = 1; $i < $line - $window -1; $i++) {
      #print "$i\n";
      if ($chrl{$i} eq $chrl{$i + $window -1}) {

          if (! @{$data{$i}} || ! @{$data{$i+$window}})  {
            print "gene $i\n";
          }
          my $corr_ave = corr_group($corr_hashref, $i);
          if ($corr_ave > $cut_off) {
            print BIN "gene $i\tcorr_ave $corr_ave\n";
            my $k;
            if ($pi < $i - $window) {
                $k = $i;
                print SIG "Break\tBreak\n";
            }
            else {
                $k = $pi + $window;
            }
            foreach (my $j = $k; $j < $i + $window; $j++) {
                print SIG "$line_data{$j}\n";
            }
            $pi = $i;
          }
      }
    }
}

sub correlation {
    my $data1 = $_[0];
    my $data2 = $_[1];
    my $product_sum = 0;
```

```
    my $x_sq_sum = 0;
    my $y_sq_sum = 0;
    my $y_sum = 0;
    if (@$data1 > 0) {
      for (my $i = 0; $i < @$data1; $i++) {
          my $val1 = $$data1[$i];
          my $val2 = $$data2[$i];
          $product_sum += $val1 * $val2;
          $x_sq_sum += $val1 * $val1;
          $y_sq_sum += $val2 * $val2;
          $y_sum += $val1;
      }
      my $sx = sqrt ($x_sq_sum / @$data1);
      my $sy = sqrt ($y_sq_sum / @$data1);
      #printf ("\nysum\t$y_sum");
      #printf ("%f\t%f\t%f\t%f\n", $product_sum, $sx, $sy);
      my $correlation = $product_sum / (@$data1 * $sx * $sy);
      return ($correlation);
    }
    else {return (0);}
}

sub prob_group {
    my @p_val;
    my @rand;
    for (my $iter = 0; $iter < $iterations + $window; $iter++)
{
      $rand[$iter] = int (rand() * ($line -1)) + 1;
    }
    my $corr_hashref = corr_hash(\%data, \@rand);
    for (my $iter = 1; $iter < $iterations +1; $iter++) {
      my $corr_ave = corr_group($corr_hashref, $iter);
      #print "p_val $iter\t$corr_ave\n";
      push (@p_val, $corr_ave);
    }
    return (\@p_val);
}

sub corr_hash {
    my $data_hashref = $_[0];
    my $order_arrayref = $_[1];
    if (! $order_arrayref) {die "missing arg corr hash\n";}
    my %corr;

    my $k = 1;
    for (my $i = $k; $i < @$order_arrayref; $i++) {
      #print "$i\n";
      if ($chrl{$k} eq $chrl{$k+$window -1}) {
          for (my $j = $i+1; $j < $i + $window; $j++) {
            my $io = $$order_arrayref[$i];
            my $jo = $$order_arrayref[$j];
            if ($$data_hashref{$io} andand
$$data_hashref{$jo}) {
```

```
                $corr{$i}{$j} =
(correlation(\@{$$data_hashref{$io}},

\@{$$data_hashref{$jo}}));
           }
           #printf ("%d\t%d\t%f\n", $i, $j, $corr{$i}{$j});
         }
         #print "\n";
     }
    }
    return (\%corr);
}

sub corr_group {
    #correlation hash, starting index
    my $corr_hashref = $_[0];
    my $i = $_[1];

    if (! $i) {die "missing arg corr_group $i\n";}

    my $sum = 0;
    my $num = 0;
    my $ave;
    for (my $j = $i; $j < $i + $window -1; $j++) {
      for (my $k = $j +1; $k < $i + $window; $k++) {
          #if ($data{$j} andand $data{$k} andand
$$corr_hashref{$j}{$k}){
          if ($$corr_hashref{$j}{$k}) {
            $sum += $$corr_hashref{$j}{$k};
            $num++;
            #print "if $$corr_hashref{$j}{$k}\t$sum\t$num\n";
          }
          else {
            print "el
$chrl{$i}\t$i\t$j\t$k\t$$corr_hashref{$j}{$k}\t$sum\t$num\n";
          }
      }
    }
    if ($num) {
      $ave = $sum / $num;
        return ($ave);
    }
    else {

      die "Corr Ave was null\t$i\n";
    }
}

sub bynum {
    $a ≤> $b;
}
```
